# Systematic Review of Aptamer Sequence Reporting in the Literature Reveals Widespread Unexplained Sequence Alterations

**DOI:** 10.1101/2021.11.02.466945

**Authors:** Alexandra A. Miller, Abhijit S. Rao, Sujana R. Nelakanti, Christopher Kujalowicz, Ted Shi, Ted Rodriguez, Andrew D. Ellington, Gwendolyn M. Stovall

## Abstract

Aptamers have been the subject of more than 144,000 papers to date. However, there has been a growing concern that errors in reporting aptamer research limit the reliability of these reagents for research and other applications. These observations noting inconsistencies in the use of our RNA anti-lysozyme aptamer served as an impetus for our systematic review of the reporting of aptamer sequences in the literature. Our detailed examination of literature citing the RNA anti-lysozyme aptamer revealed that 93% of the 61 publications reviewed reported unexplained altered sequences with 86% of those using DNA variants. The ten most cited aptamers were examined using a standardized methodology in order to categorize the extent to which the sequences themselves were apparently improperly reproduced, both in the literature and presumably in experiments beyond their discovery. Our review of 800 aptamer publications spanned decades, multiple journals, and research groups, and revealed that 44% of the papers reported unexplained sequence alterations. We identified ten common categories of sequence alterations including deletions, substitutions, additions, among others. The robust data set we have produced elucidates a source of irreproducibility and unreliability in our field and can be used as a starting point for building evidence-based best practices in publication standards to elevate the rigor and reproducibility of aptamer research.

---

The irreproducibility of research has become increasingly evident in several fields in recent years ^1^ and poses a serious threat to the efficacy and validity of the research and the general public’s perception of science. The field of aptamer research may be particularly susceptible to this reproducibility crisis.^2,3,4,5^ In part, the ability to develop affinity reagents quickly and simply based on molecular biology manipulations alone provides a ready entry point for a variety of researchers with varying backgrounds. The development of aptamers for use stands in stark contrast to the development of monoclonal antibodies via hybridoma and other technologies, where there is a much larger and longer trail of publications and standards, both academic and commercial, and where there is much larger use, allowing greater cross-validation. Because of the lack of adherence to suggested standards in aptamer research^2,6^, the reliability of aptamers has been drawn into question, undermining the credibility of aptamers.^3,7^

As a particularly egregious example of the issues relating to the reproducibility and reliability of aptamer research, we first noted the erred interpretation of an 80-mer RNA antilysozyme binding aptamer (Clone 1).^2,8^ As we now report in greater detail, subsequent works not only altered the aptamer sequence (using DNA as opposed to the original RNA aptamer), but also in a truncated the aptamer, which ultimately may have compromised affinity and specificity.^4,9^ Given the paucity of publication standards for the field ^6,8^, the sequence alteration of the anti-lysozyme binding aptamer was likely not an isolated event, but rather a systemic and widespread issue of reproducibility and reliability, stemming in part from the rapid increase in application-based publications.^6^

To elucidate the extent of *in silico* aptamer sequence alterations, we sampled a large swath of aptamer literature to gauge the fidelity of aptamer sequence reporting. We examined aptamers against the most frequently used aptamer targets: thrombin, adenosine triphosphate (ATP), vascular endothelial growth factor (VEGF), platelet derived growth factor (PDGF) BB, cocaine, theophylline, lysozyme, nucleolin, immunoglobulin E (IgE), and ochratoxin A (OTA) (top ten targets listed by application-based publication frequency.^6^ We identified original aptamer sequences and followed their subsequent descriptions in cited literature, characterizing apparent sequence alterations as adequately described, omitted, or unexplained. The unexplained aptamer sequence findings were then organized into phylogenic trees that aided in our identification of common sources and types of apparent error (**Figure 2**).

In our review of 800 publications, we provide evidence of widespread unexplained aptamer sequence alteration over time. Aptamers appear to ‘devolve’ in the literature. These findings strongly suggest the need for stringent publication standards to prevent future sequence errors that could compromise function and reproducibility of aptamers.

## METHODS

### Examination of the Reported Sequences for the RNA Antilysozyme Aptamer, Clone 1^8^

An anti-lysozyme aptamer selected in 2001 had previously been reported to ‘mutate’ in the literature over time, and we therefore used this aptamer as a starting point for establishing a broader methodology.^2,4^ As part of this methodology, we identified publications in Google Scholar that cited the original work using the “cited by” option for this reference. From the top literature results, those that were written in English, primary literature (i.e., not reviews, etc.), and used the aptamer *in vitro* or *vivo* were analyzed for reported sequence information. The aptamer sequence(s) were extracted with notes on any alterations to the sequence, using direct quotes from the text.

Analysis of the collected sequences then included the following:

1. Comparison of the reported sequences to the original aptamer sequence (Clone 1)^8^ and identification of alterations.
2. If alterations were present, further examination of the cited publication led to an assessment of whether the variants were adequately explained or unexplained.
3. In those cases where there were unexplained sequence alterations, they were classified into ten categories: (i) deletion, (ii) insertion, (iii) substitution, (iv) complete inversion, (v) 5’/3’ addition, (vi) partial inversion, (vii) complementary sequence, (viii) incorrect sequence attribution (i.e., in a figure or table), (ix) sequence omitted, and (x) unknown (i.e., reported an entirely different sequence).

Using phylogenetic trees as a model, we organized the aptamers reported in the literature similarly using sequence homology to visualize sequence fidelity in our sample. We loosely dubbed this organization an aptamer “phylogenetic” analysis. The nodes in the artificial phylogenetic trees represent groupings of aptamers according to unexplained sequence alterations. Further unexplained derivations were depicted as branching from a previous sequence alteration (i.e., **Figure 1, node 4A** was derived from **Figure 1, node 3A**). Publications that correctly reported the original aptamer sequence, including those with fully explained or described additions (e.g., 5’-terminal polyT linker or primary amine for conjugation), were given the same identifying number as the original aptamer.

**Figure 1.**
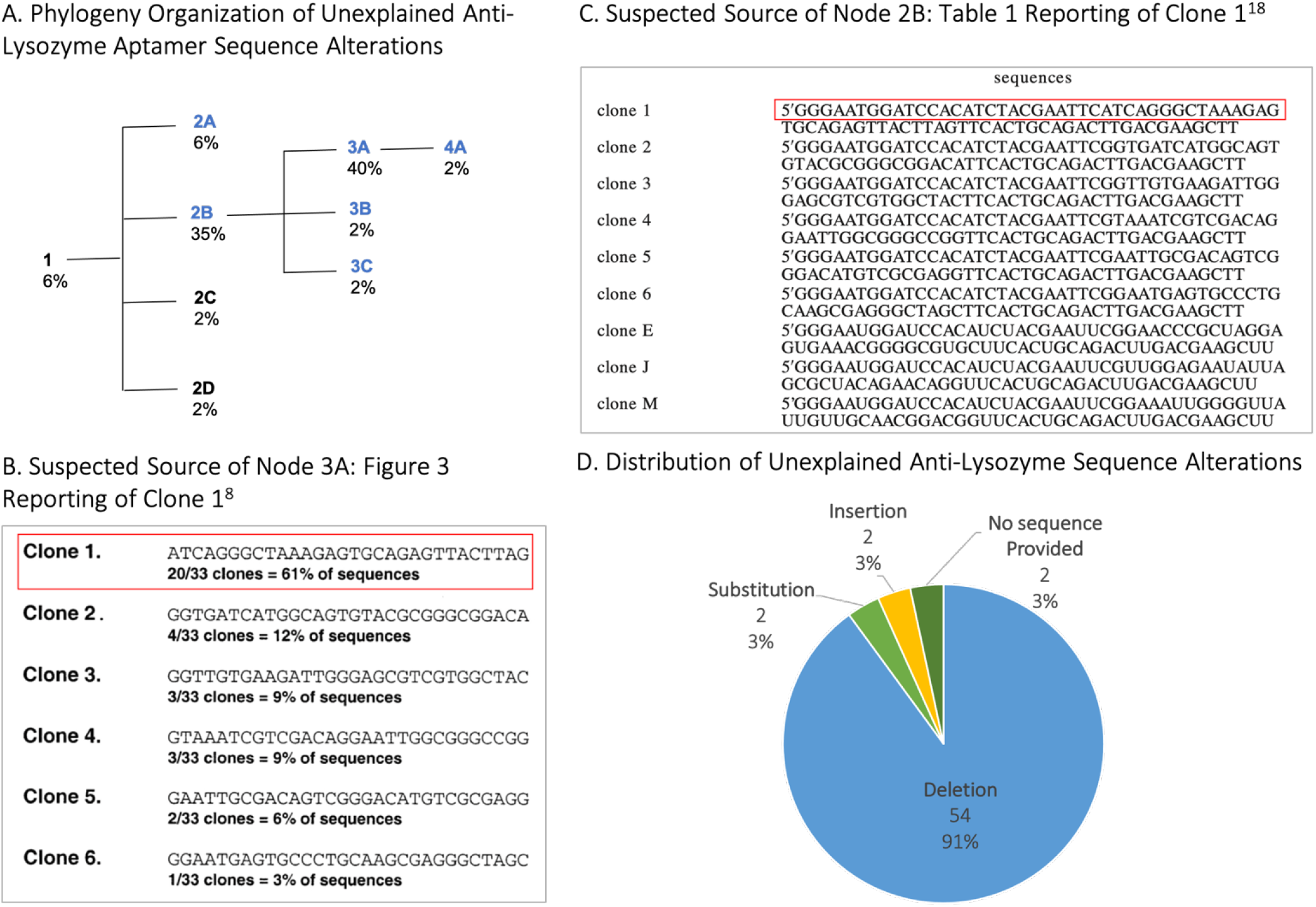
Phylogeny depicting unexplained sequence alterations introduced to the Clone 1 anti-Lysosome RNA Aptamer^8^ collected 01/01/2020. A. We loosely dibbed this sequence organization and aptamer “phylogenetic” analysis. The nodes in the artificial phylogenetic trees represent groupings of aptamers according to unexplained sequence alterations. The 79 references reviewed (18 papers excluded) can be found in the Supplemental 1 section. B. Reprinted from Cox, J. C., & Ellington, A. D. (2001). Automated selection of anti-protein aptamers. Bioorganic & medicinal chemistry, 9(10), 2525-2531 with permission from Elsevier. C. Reprinted in part with permission from Kirby, R.; Cho, E.J.; Gehrke, B.; Bayer, T.; Park, Y.S.; Neikirk, D.P.; McDevitt, J.T.; Ellington, A.D. Analytical Chemistry, 2004. 76 (14), 4066-4075. Copyright 2004 American Chemical Society. D. Unexplained sequence alterations were classified into categories.

### Applying the Standardized Method to Aptamer Sequences Across the Literature

To provide broader insights into the quality of the aptamer literature, we reviewed aptamers against the 10 most-used targets^6^ (**Table S1**), and again determined a primary (original) aptamer sequence, identified derivatives using the Google Scholar “cited by” option (i.e., works citing the original). From the results generated, publications were further sieved by identifying only those that reported aptamers against the target using the “search within citing articles” function and the search terms “[target]” “aptamer” (**Table S1**). Without this second search, extraneous papers -- for example those citing the anti-VEGF aptamer in the introduction -- would have been included across multiple reviews or other experimental papers. We then categorized the unexplained sequence alterations according to apparent type and organized the reported sequence derivatives into “phylogenies” according to sequence homology.

The literature search for each aptamer excluded publications that did not use the aptamer sequence experimentally or that were not in the English language (i.e., 172 publications were excluded); however, no exclusions were made based on the type of journal or date of publication. Publications were searched from some of the original selection experiments in 1990 ^10-11^ through March 2020.

The sample size for each aptamer examined varied between 22-171 publications, although we aimed to acquire at least 50 papers for each phylogeny. Cases where there was a smaller sample size included the Tran et al. anti-lysozyme aptamer^12^ and the Jellinek et al. anti-vascular endothelial growth factor (VEGF) aptamer^13^, whose phylogenies contained 22 and 25 papers respectively. The majority of the literature citing these originating aptamers did not use the sequence *in vitro* or *in vivo*, and therefore were excluded from their respective phylogenies **(Supplement 2 Raw Data Drive)**.

## RESULTS AND DISCUSSION

### A Study of the RNA Anti-Lysozyme Aptamer, Clone 1^8^

Lysozyme is one of the most utilized targets in the field of aptamer research and is amongst the top 10 in terms of citations.^6^ This popularity was driven in large measure by the publication of an RNA anti-lysozyme aptamer by Cox et al. (2001)^8^. We previously noted^2^ that while the original selection had been for an RNA aptamer, subsequent uses of this aptamer had sometimes involved the synthesis of a DNA version that had not been verified as binding to lysozyme; in other words, the aptamer had been mutated *in silico*. This error was perhaps understandable, given that the figure in the original paper showed the DNA template for the production of the RNA aptamer (although the text clearly describes the generation of RNA transcripts and Figures 2 and 4 note the use of RNA). This misreading and misattribution of the original work led to the *in silico* alterations propagated by subsequent groups (**Table S2**).

**Figure 2.**
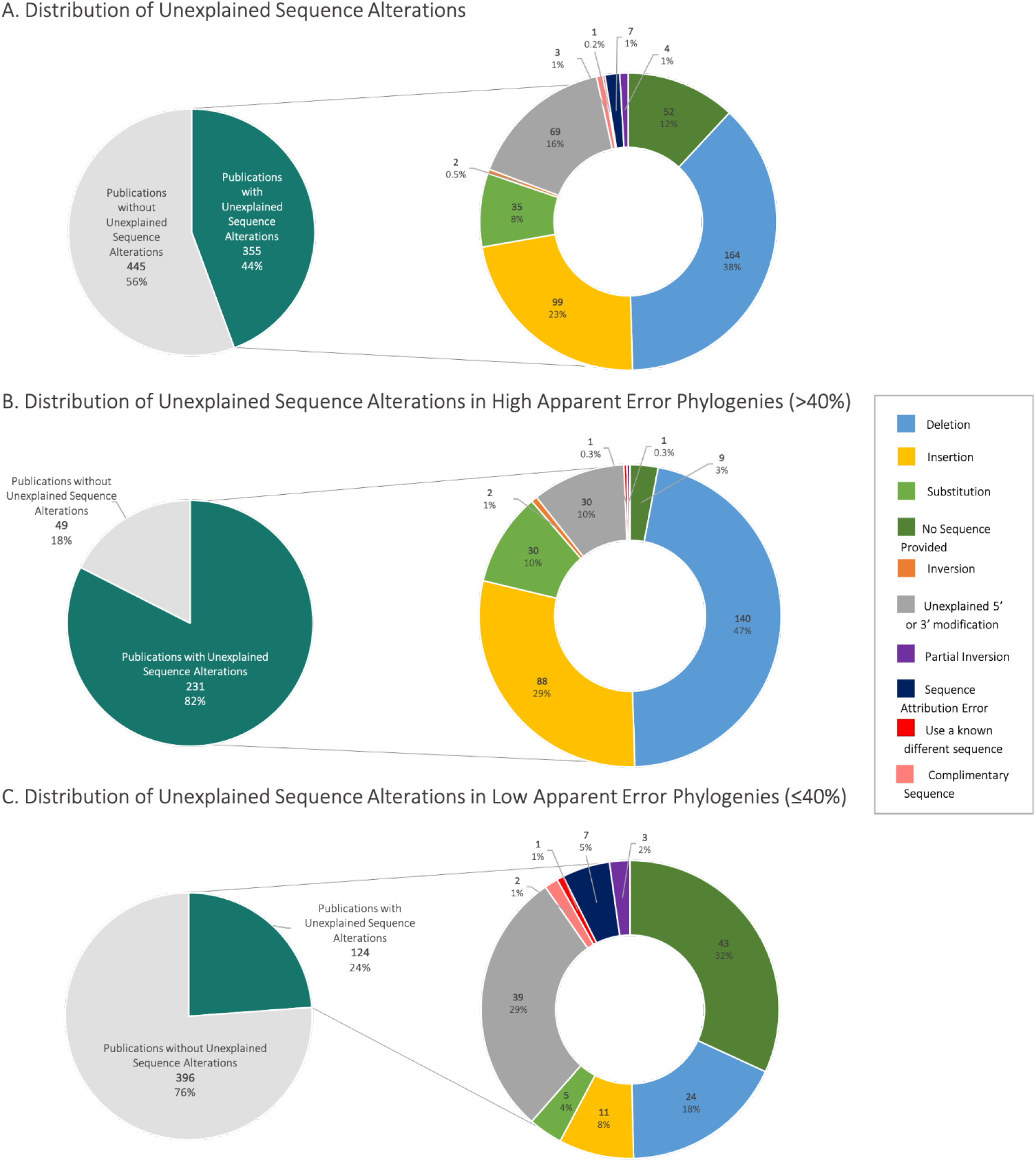
Distribution of Field-wide Unexplained Sequence Alterations. **A**. The breakdown of all publications examined (n = 800 publications, 169 excluded, 23 phylogenies grouped by 11 root publications) **B**. The distributions of phylogenies that contained greater than the median of 40% internal sequence alteration (5 phylogenies: Cox lysozyme, Tran lysozyme, ATP, PDGF BB, cocaine). **C**. The distribution of phylogenies that contained less than or equal to the median of 40% internal error (6 phylogenies: nucleolin, IgE, theophylline, VEGF, ochratoxin A, thrombin). The percentage of unexplained sequence alterations within each phylogeny can be found in Table S3.

To examine how *in silico* alterations can be propagated, we reviewed 61 primary research papers that cited the antilysozyme aptamer, Clone 1^8^. Of those reports that varied from the original RNA aptamer, the papers were reviewed to ascertain if the reported alterations were adequately described, omitted, or unexplained. Sequence alterations found include deletions, insertions, substitutions, inversions or partial inversions, improper description of complementary sequences, incorrect sequence attribution (i.e., boxing the incorrect aptamer sequence in a Figure), and/or reporting an entirely different sequence than the sequence cited.

In an effort to illustrate and organize the aptamer sequence data as it “evolved” in the literature, an aptamer phylogeny was created. The phylogeny includes the original/parental 80-mer anti-lysozyme RNA aptamer (Clone 1)^8^, also called the “root” aptamer, and nodes are organized by sequence homology (i.e., those that more greatly differ from the root aptamer are further away and have a larger node number). As shown in **Figure 1A**, the phylogeny includes 4 branch points and 9 nodes in all (including the root aptamer) with DNA variants indicated as blue nodes. While we made our best attempts to ascertain what was “reported” versus what was actually “used,” in most cases, we could not distinguish one from the other and thus settled on the liberal use of what was “reported.” Further, in an attempt to ascertain the source of the unexplained sequence modifications, we examined altered sequences and the publications they cited and report our hypotheses here. The anti-lysozyme aptamer sequence phylogeny alignment is shown in **Table 1**.

**Table 1.**
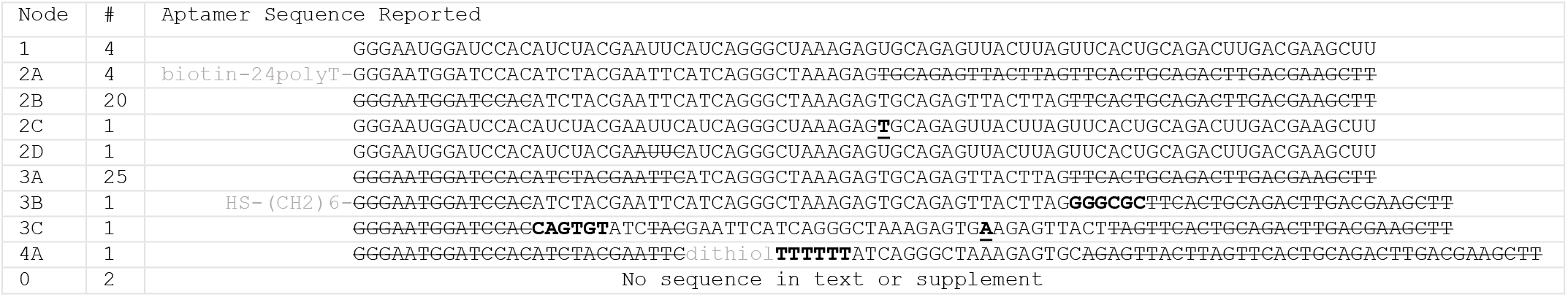
Unexplained sequence alterations introduced to the Clone 1 anti-lysosome RNA aptamer.^**8**^. Unexplained insertions are bolded, unexplained substitutions are bolded and underlined, unexplained deletions are struck out and justified or explained alterations are in light grey. The number (#) column indicates the number of publications found reporting each sequence in our analysis

After the root anti-lysozyme RNA aptamer node (1) in **Figure 1**, the first branch point leads to four nodes. The first node, 2A, is an unexplained truncated DNA sequence with a 39 nt 3’ deletion, as well as terminal additions (5’ biotinylation and a 24 nt oligo(T) linker). This aptamer was primarily used by a single group. ^14-17^ We hypothesize that the alteration was due to a misinterpretation of the Kirby et al. (Table 1)^18^, shown in **Figure 1C**, which reported the DNA template of the six original anti-lysozyme aptamers^8^ and three novel RNA aptamers with a line break in the sequences. Node 2A includes the first line of this sequence. A later 2015 publication by the same group reported the KD of the altered sequence (2A) without discussing its origin.^15^

The 2B node (**Figure 1**) is an unexplained DNA sequence with a 24 nt deletion from the 3’ end and a 14 nt deletion from the 5’ end and was found in 35% of papers reviewed. The 2C node is the RNA aptamer sequence, but with a single U to T substitution reported in the paper, an *in silico* error but likely not in the molecule used.^19^ Finally, the 2D node is a full-length RNA sequence but with a deletion of 4 internal nucleotides^20^, likely again an error in the paper but not the molecule itself, as the manuscript notes the acquisition of the original aptamer material from the lab where the aptamer originated.

All further unexplained sequence modifications branch from node 2B, the citation of an all-DNA variant (**Figure 1A**). Organized by sequence homology, the 5’ and 3’ cropped variant at the 2B node leads to a branch point with three nodes: 3A, 3B, and 3C. Node 3A, which makes up 40% of the altered sequences in this phylogeny, is a DNA variant with primer-binding regions deleted (i.e., 5’ 26 nt deletion and 3’ 24 nt deletion). This sequence is equivalent to what was reported in Figure 3 of the 2001 paper^8^, although this sequence was not in fact used for experiments. We suspect that the node 3A sequence alteration arose due to a misunderstanding of Figure 3 in the original paper, which reported the 30 nt DNA template for the random sequence region of the selected RNA aptamer, without also including accompanying primer-binding regions (although the remainder of the paper indicates the sequence of the entire RNA aptamer).^8^ Presumably, groups that mistakenly used the DNA template of the random region obtained it by using this figure.

Variant 3B differs from 3A in that it has an unexplained 3’-terminal addition of GGGCGC, the partial (13 nt) return of the 5’ primer region, and an adequately explained 5’-thiol modification (5’-HS-(CH2)6) for covalent conjugation to gold nanoparticles.^21^ Variant 3C contains a number of otherwise unexplained modifications: a C to A internal substitution, a 6 nt 5’-terminal addition, a 3 nt 3’-terminal deletion, and a 5’ – terminal 3 nt deletion.^22^

Finally, the 4A sequence (**Figure 1**) again builds off of the deletion of 5’ and 3’ primer regions described in node 3A and adds a well-described 6 nt polyT linker and thiol modification at the 5’ end for conjugation to silica particles, but also includes an unexplained additional 13 nt 3’ deletion.^23^

Notably, in some cases, the designation of “adequately described” was debatable and thus up to our discernment. For example, Truong et al.^24^ call their Cox anti-lysozyme aptamer the “5’ thiol modified DNA template of lysozyme RNA aptamer,” referencing Cox et al. (2001)^8^ and Kirby et al. (2004)^18^, but do not provide a rationale for the use of the DNA template rather than the selected RNA aptamer. Therefore, we categorized it as inadequately explained use of the DNA variant. The complete raw data for our classifications has been included (**Supplement 2 Raw Data Drive**) in an effort to increase the transparency of our findings.

Surprisingly, in all the reviewed publications, there were only 4 papers of the 61 reviewed that reported the correct antilysozyme aptamer, Clone 1 sequence and/or adequately explained sequence alterations. Of those that contained unexplained sequence alterations, 87% reported a DNA variant, 91% contained a deletion, 3% contained a substitution, 3% contained an insertion, and 3% failed to include sequence information in the manuscript or supplement. The original error in the interpretation of the anti-lysozyme aptamer sequence, Clone 1 was propagated *in silico* through multiple generations of researchers (**Figure 1**).

The question thus becomes not only why so many researchers misinterpreted the primary source of information, but also how the derivatives themselves work as reported? The answer is two-fold: first, hen egg-white lysozyme was originally chosen as a target for selection in part because it is extremely basic (pI = 11.35), and thus was very likely to yield high-affinity aptamers. But second, precisely because polyanions will bind oligocations, the DNA aptamer may still bind lysozyme. In an illuminating study, Potty et al.^4^ examined the binding properties of both the RNA aptamer and DNA of anti-lysozyme aptamer Clone 3 (an entirely different aptamer sequence than the Clone 1 sequence that is generally used). These authors found the DNA variant bound non-specifically to lysozyme through electrostatic interactions.^4^

This further raises the question of what constitutes specific binding. In general, aptamer studies should contain non-cognate binding experiments, in which variants of the aptamers (mutant or scrambled) are shown not to bind to the selection target, and the selected aptamer is shown not to bind to targets other than the selection target. Such studies are rarely carried out.

Overall, only 4 of the 61 examined RNA anti-lysozyme Clone 1 aptamer sequences recorded were correct or had modifications that were adequately explained. As we have previously suggested^2^, the lack of detail and level of apparent error in the reporting of aptamer experiments in general requires a reckoning in the field as a whole. To this end, we broadly sampled the aptamer literature to describe apparent sequence error in the field.

### Description of Error in the Field of Aptamer Research

Based on the extensive set of unexplained aptamer sequence alterations that were found in literature citing the antilysozyme aptamer^8^, we hypothesized that the incorrect reporting of aptamer sequences could be a more pervasive problem, potentially due to the rapid expansion of research into aptasensors^6^, and the lack of standardized guidelines for aptamer reporting.^2,6^ While a full review of the 135,000+ “aptamer” papers on Google Scholar was not feasible, a review of aptamers against the ten most used targets^6^ was carried out. This review systematically sampled a broad swath of the aptamer literature, spanning years, researchers, labs, and locations.

Similar to the analysis of the Cox anti-lysozyme binding aptamer literature, we first identified an originating aptamer and then examined the literature citing the sequence of this originating aptamer. In cases where multiple clones selected in a paper were used in the literature, all clones were examined as “root” aptamers (23 in all). Unexplained sequence alterations were again categorized, and phylogenies constructed. In all, 800 publications from 23 originating/root aptamers were reviewed using this standardized sampling methodology (**Table S3, Figures 1-10**). The 23 phylogenies were also grouped based on an originating publication (e.g., three different anti-PDGF aptamer clones from Green and Jellinek, et al. (1996)^25^ are grouped in **Figure S4**).

Overall, only 56% of the 800 publications reviewed correctly reported the aptamer sequence(s), while 44% (355 papers containing 362 altered sequences) contained one or more *in silico* sequences that were categorized as unexplained. We identified 137 novel sequences that according to our criteria were not adequately explained (**Table S4, Supplement 1, Supplement 2 Raw Data**). Some (38%) of these 137 sequences identified in 362 considered altered sequences contained deletions, 23% contained insertions, 16% contained 5’ or 3’ modifications, 12% did not provide the sequence at all, 8% contained substitutions, and less than 5% contained sequence attribution errors (i.e., boxing the incorrect aptamer sequence, etc.), inversions, the complementary sequence, or an entirely different sequence (**Figure 2**). Anti-VEGF aptamers were a notable outlier in these analyses because we did not find altered sequences, explained or not, using our methods. Instead, we provide a figure describing all novel VEGF aptamer sequences described and indicate papers that did not provide sequence information (**Figure S3**).

To further categorize the types of apparent *in silico* errors that are perpetuated in the literature, we used a “median split” to divide the dataset into two groups: “low”/few apparent errors and “high”/more apparent errors.^26-27^ The median percentage of unexplained sequence alterations within each phylogeny was 40%, and this value was used for the division into groups (**Table S2**, see also **Supplement 2 Raw Data**), (**Figure 2B and 2C**). There was a larger contribution of deletions (47%) and insertions (29%) in the high-level of unexplained sequence alteration group relative to the low-level of unexplained sequence alteration group, which had 18% deletions and 8% insertions (**Figure 2B and 2C**). In contrast, the low-level unexplained sequence alteration group contained a higher contribution of unexplained 5’/3’ modifications (29%) compared to the high sequence alteration group (10%). This observation generally suggests that deletions and insertions lead to subsequent reporting errors in the literature, while 5’/3’ modifications that may have unique, researcher-specific purpose do not. Since the deletions/insertion errors are most likely copy-edit errors, particularly where sequences could not be copied digitally from a figure or secondary structure directly, we believe the inclusion of a text file in manuscripts that contains relevant information for experimental replication will help ameliorate copy-edit errors.

## CONCLUSIONS

Our review of 800 publications reveals widespread sequence alterations that are poorly explained or unjustified in reported sequences (44%). This observed systemic lack of sequence fidelity, while not exhaustive, buttresses the conclusions of Yan and Levy^3^ that aptamers have been unreliable as reagents. The reasons underlying the large apparent error rates we report here can only be speculated on, but likely include the relative ease of creating aptamers based on literature reports alone, the rapid expansion of aptamer research, the rapid increase in application-based publications^6^ and a lack of adherence to coherent publication guidelines.^2,6^ Further, once unexplained sequence alterations propagate in the literature these may dilute the citation of accurate sequences, making it more difficult to discern the original, correct sequence.

Echoing previous work^2,3,6^, we assert that there is an urgent need both for the collaborative construction of evidence-based publication guidelines and standardized documentation of aptamer sequence use in the literature (e.g.,similar to CiteAb.com). Standardized sequence reporting could be modelled after Nature’s “Life Science Reporting Summary” and Cell’s “STAR Methods,” which create standardized templates for detailed descriptions of Experimental Design, Statistical Tests, and Materials and Reagents. At a minimum, we suggest that three categories of information should be required in aptamer publications: complete sequence and secondary structure information (i.e., mFOLD), including consistent sequence identifiers; detailed descriptions of binding and experimental conditions; and extensive use of negative and positive controls, for both aptamers and targets. As Yan and Levy^3^ suggest, we also support validation practices such as using an environment that is similar to the downstream application, limited incubation times and using blocking reagents to minimize nonspecific binding to targets, and multiple validations, ideally by different researchers.

In the future, collaborative evidence-based aptamer publication and validation guidelines should be encouraged by journals, including a checklist for the peer review process. Ultimately, we believe that the standardization of aptamer publication guidelines and increased availability of raw data will lead to a more open, nuanced discussion of the data presented, fewer wasted resources, and greater success in the translation of aptamer research to the clinic and industry.

## Supporting information

Raw Data

## ASSOCIATED CONTENT

### Supporting Information

The Supporting Information is available free of charge on the ACS Publications website.

Table S1 lists the top ten targets used and the search terms used during the Google Scholar literature search. Table S2 lists the publications reporting each sequence identified in the antilysozyme Clone 1 aptamer literature review. Table S3 lists the percentage of publications with unexplained sequence alterations within each phylogeny. Table S4 lists the number of unique sequences and number of sequences with unexplained alterations. Figures S1-S10 describe the extend of publication-induced error in the top ten aptamer targets using the method described here as well as the initial data collection using various methods (PDF).

The raw data collected for all 23 phylogenies grouped by originating paper (11 originating publications) and data analysis are included (.xls).

The database collected in this work with any additions or corrections is available, https://drive.google.com/drive/folders/17UIczwFHXwPOtJilqoTH6rhm1Lmw9As8?usp=sharing

## AUTHOR INFORMATION

### Author Contributions

A.A.M., S.R.N., A.S.R., C.K., T.R., and T.S. performed initial data collection. A.A.M. collected all data and analyzed the data.

A.A.M., G.M.S., and A.D.E. reviewed and interpreted the results. The methods were drafted by S.R.N. The final manuscript was consolidated and written by A.A.M. and edited by G.M.S., A.A.M., A.S.R., and A.D.E. All authors have approved the final version of the manuscript.

## ACKNOWLEDGMENT

Apart from Gwendolyn Stovall and Andrew Ellington, the authors began this work as undergraduate researchers in the UT Freshman Research Initiative and would like to thank the FRI’s Aptamer Stream and the 2018 TIDES Science Sprints at the University of Texas at Austin for supporting this project. The authors would also like to thank Jolyn Frnka, Haley Wolf, Shreya Thiagarajan, Eric Lumanog, Christine Mai, Hung Le, Kathy Tran, and Nidhi Pawate for their assistance in the cursory data collection and development of the method.

This research was funded by The University of Texas Freshman Research Initiative, which was supported by Howard Hughes Medical Institute (#52008124) and the College of Natural Sciences. Any opinions, findings, and conclusions or recommendations expressed in this material are those of the author(s) and do not necessarily reflect the views of The University of Texas at Austin or the Howard Hughes Medical Institute.

---

Authors are required to submit a graphic entry for the Table of Contents (TOC) that, in conjunction with the manuscript title, should give the reader a representative idea of one of the following: A key structure, reaction, equation, concept, or theorem, etc., that is discussed in the manuscript. Consult the journal’s Instructions for Authors for TOC graphic specifications.

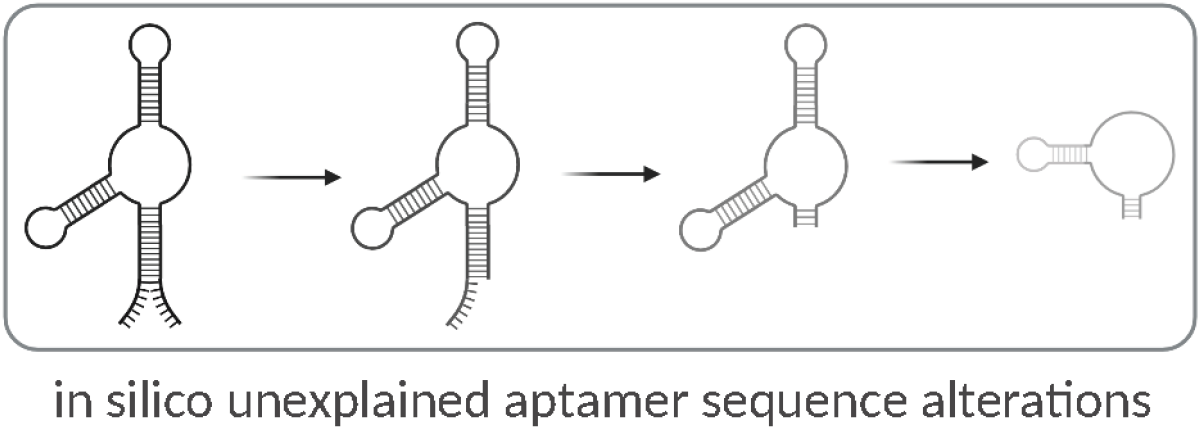

## REFERENCES

1. Miyakawa, T. (2020). No raw data, no science: Another possible source of the reproducibility crisis. Molecular Brain, 13(1), 1–6. https://doi.org/10.1186/s13041-020-0552-2

2. Cho, E. J.,, Lee, J.-W.; Ellington, A. D. (2009). Applications of Aptamers as Sensors. Annual Review of Analytical Chemistry, 2(1), 241–264. https://doi.org/10.1146/annurev.anchem.1.031207.11285

3. Yan, A. C.; Levy, M. (2018). Aptamer-Mediated Delivery and Cell-Targeting Aptamers: Room for Improvement. Nucleic Acid Therapeutics, 28(3), 194–199. https://doi.org/10.1089/nat.2018.0732

4. Potty, A. S. R.; Kourentzi, K.; Fang, H.; Schuck, P.; Willson, R. C. (2011). Biophysical characterization of DNA and RNA aptamer interactions with hen egg lysozyme. International Journal of Biological Macromolecules, 48(3), 392–397. https://doi.org/10.1016/j.ijbiomac.2010.12.007

5. Haßel, S. K.; Mayer, G. (2019). Aptamers as Therapeutic Agents: Has the Initial Euphoria Subsided? Molecular Diagnosis and Therapy, 23(3), 301–309. https://doi.org/10.1007/s40291-019-00400-

6. Dunn, M. R.; Jimenez, R. M.; Chaput, J. C. (2017). Analysis of aptamer discovery and technology. Nature Reviews Chemistry, 1. https://doi.org/10.1038/s41570017-007

7. Li, N.; Ebright, J. N.; Stovall, G. M.; Chen, X.; Nguyen, H. H.; Singh, A.; … Ellington, A. D. (2009). Technical and biological issues relevant to cell typing with aptamers. Journal of Proteome Research, 8(5), 2438–2448. https://doi.org/10.1021/pr801048z

8. Cox, J. C.; Ellington, A. D. (2001). Automated Selection of Anti-Protein Aptamers. Bioorganic and Medicinal Chemistry, 9, 2525– 2531. https://doi.org/10.1016/s0968-0896(01)00028-

9. Padlan, C. S.; Malashkevich, V. N.; Almo, S. C.; Levy, M.; Brenowitz, M.; Girvin, M. E. (2014). An RNA aptamer possessing a novel monovalent cation-mediated fold inhibits lysozyme catalysis by inhibiting the binding of long natural substrates. Rna, 20(4), 447–461. https://doi.org/10.1261/rna.043034.113

10. Ellington, A. D.; Szostak, J. W. (1990). In vitro selection of RNA molecules that bind specific ligands. Nature, 346(6287), 818–822. https://doi.org/10.1038/346818

11. Tuerk, C.; Gold, L. (1990). Systematic evolution of ligands by exponential enrichment: RNA ligands to bacteriophage T4 DNA polymerase. Science, 249(4968), 505–510. https://doi.org/10.1126/science.2200121

12. Tran, D. T.; Janssen, K. P.; Pollet, J.; Lammertyn, E.; Anné, J.; Van Schepdael, A.; Lammertyn, J. (2010). Selection and characterization of DNA aptamers for egg white lysozyme. Molecules, 15(3), 1127–1140.

13. Jellinek, D.; Green, L. S.; Bell, C.; Janjić, N. (1994). Inhibition of Receptor Binding by High-Affinity RNA Ligands to Vascular Endothelial Growth Factor. Biochemistry, 33(34), 10450–10456. https://doi.org/10.1021/bi00200a028

14. Mihai, I.; Vezeanu, A.; Polonschii, C.; David, S.; Gáspár, S.; Bucur, B.; … Vasilescu, A. (2014). Low-fouling SPR detection of lysozyme and its aggregates. Analytical Methods, 6(19), 7646– 7654. https://doi.org/10.1039/c4ay01237b

15. Mihai, I.; Vezeanu, A.; Polonschii, C.; Albu, C.; Radu, G. L.; Vasilescu, A. (2015). Label-free detection of lysozyme in wines using an aptamer based biosensor and SPR detection. Sensors and Actuators, B: Chemical, 206, 198–204. https://doi.org/10.1016/j.snb.2014.09.050

16. Vasilescu, A.; Gaspar, S.; Mihai, I.; Tache, A.; Litescu, S. C. (2013). Development of a label-free aptasensor for monitoring the self-association of lysozyme. Analyst, 138(12), 3530–3537. https://doi.org/10.1039/c3an00229b

17. Vasilescu, A.; Purcarea, C.; Popa, E.; Zamfir, M.; Mihai, I.; Litescu, S.; … Marty, J. L. (2016). Versatile SPR aptasensor for detection of lysozyme dimer in oligomeric and aggregated mixtures. Biosensors and Bioelectronics, 83, 353–360. https://doi.org/10.1016/j.bios.2016.04.080

18. Kirby, R.; Cho, E. J.; Gehrke, B.; Bayer, T.; Park, Y. S.; Neikirk, D. P.; … Ellington, A. D. (2004). Aptamer-based sensor arrays for the detection and quantitation of proteins. Analytical Chemistry, 76(14), 4066–4075. https://doi.org/10.1021/ac049858n

19. Cho, E. J.; Collett, J. R.; Szafranska, A. E.; Ellington, A. D. (2006). Optimization of aptamer microarray technology for multiple protein targets. Analytica Chimica Acta, 564(1), 82–90. https://doi.org/10.1016/j.aca.2005.12.038

20. Hybarger, G.; Bynum, J.; Williams, R. F.; Valdes, J. J.; Chambers, J. P. (2006). A microfluidic SELEX prototype. Analytical and Bioanalytical Chemistry, 384(1), 191–198. https://doi.org/10.1007/s00216-005-0089-3

21. Zuo, L.; Qin, G.; Lan, Y.; Wei, Y.; Dong, C. (2019). A turn-on phosphorescence aptasensor for ultrasensitive detection of lysozyme in humoral samples. Sensors and Actuators, B: Chemical, 289(December 2018), 100–105. https://doi.org/10.1016/j.snb.2019.03.088

22. Shamsipur, M.; Farzin, L.; Tabrizi, M. A. (2016). Ultrasensitive aptamer-based on-off assay for lysozyme using a glassy carbon electrode modified with gold nanoparticles and electrochemically reduced graphene oxide. Microchimica Acta, 183(10), 2733–2743. https://doi.org/10.1007/s00604-016-1920-6

23. Truong, P. L.; Choi, S. P.; Sim, S. J. (2013). Amplification of resonant rayleigh light scattering response using immunogold colloids for detection of lysozyme. Small, 9(20), 3485–3492. https://doi.org/10.1002/smll.201202638

24. Green, L. S.; Jellinek, D.; Jenison, R.; Östman, A.; Heldin, C. H.; Janjic, N. (1996). Inhibitory DNA ligands to platelet-derived growth factor B-chain. Biochemistry, 35(45), 14413–14424. https://doi.org/10.1021/bi961544+

25. Iacobucci, D.; Posavac, S. S.; Kardes, F. R.; Schneider, M. J.; Popovich, D. L. (2015). The median split: Robust, refined, and revived. Journal of Consumer Psychology, 25(4), 690–704. https://doi.org/10.1016/j.jcps.2015.06.01

26. Bayramoglu, G.; Ozalp, V. C.; Yilmaz, M.; Guler, U.; Salih, B.; Arica, M. Y. (2015). Lysozyme specific aptamer immobilized MCM-41 silicate for single-step purification and quartz crystal microbalance (QCM)-based determination of lysozyme from chicken egg white. Microporous and Mesoporous Materials, 207, 95–104. https://doi.org/10.1016/j.micromeso.2015.01.009

27. DeCoster, J.; Gallucci, M.; Iselin, A.-M. R. (2011). Best Practices for Using Median Splits, Artificial Categorization, and their Continuous Alternatives. Journal of Experimental Psychopathology, 2(2), 197–209. https://doi.org/10.5127/jep.008310

