## Supplement 1 for "Systematic Review of Aptamer Sequence Reporting in the Literature Reveals Widespread Unexplained Sequence Alterations"

**Table S1. Search Terms Used for Literature Reviews. Ordered by frequency of aptamer target used in application publications (Dunn et al., 2017).**

| Aptamer Target | "Search within citing articles" Search Term |
|---|---|
| Human Thrombin | "aptamer" "human thrombin" |
| Adenosine Triphosphate (ATP) | "aptamer" "ATP" |
| Vascular Endothelial Growth Factor (VEGF) | "aptamer" "vegf" |
| Platelet Derived Growth Factor (PDGF) BB | "aptamer" "pdgf bb" |
| Cocaine | "aptamer" "cocaine" |
| Theophylline | "aptamer" "theophylline" |
| Lysozyme (Cox et al. 2001 and Tran et al., 2010) | "aptamer" "lysozyme" |
| Nucleolin | "aptamer" "nucleolin" |
| Immunoglobulin E (IgE) | "aptamer" "ige" |
| Ochratoxin A (OTA) | "aptamer" "ochratoxin a" |

**Table S2. Publications Reporting Each Altered Sequence in the Cox et al. 2001 phylogeny.**

| ID | Publications Reporting Sequence |
|---|---|
| 1 | Collett et al., 2004 (12), **Cox et al., 2001** (14), Padlan et al., 2014 (48), Kirby et al., 2004 (30) |
| 2A | Mihai et al., 2014 (45), Mihai et al., 2006 (46), Vasilescu et al., 2013 (61), Vasilescu et al., 2016 (62) |
| 2B | Chen et al., 2013 (7), Chen et al., 2013 (8), Cheng et al., 2007 (9), Giuffrida et al., 2018 (20), Hayashi et al., 2010 (22), Huang et al., 2009 (23), Li et al., 2011 (36), Li et al., 2014 (35), Li et al., 2007 (33), Li et al., 2016 (32), Li et al., 2010 (34), Lin et al., 2017 (38), Peng et al., 2009 (49), Rodriguez et al., 2009 (52), Teller et al., 2009 (57), Truong et al., 2013 (60), Wang et al., 2010 (68), Wang et al., 2011 (67), Yeung et al., 2010 (72), Zhang et al., 2011 (75) |
| 2C | Cho et al., 2006 (11)* |
| 2D | Hybarger et al., 2006 (26) |
| 3A | Bamrungsap et al., 2011 (2), Bi et al., 2017 (4), Cao et al., 2017 (5), Cheglakov et al., 2007 (6), Deore et al., 2019 (16), Fang et al., 2016 (17), Ghosh et al., 2018 (18), Giradot et al., 2011 (19), Huang et al., 2010 (25), Khan et al., 2018 (28), Liao et al., 2013 (37), Liu et al., 2014 (39), Liu et al., 2018 (40), Lu et al., 2013 (42), Ocaña et al., 2015 (46), Ostatná et al., 2017 (47), Song et al., 2011 (54), Subramanian et al., 2013 (),Wang et al., 2010 (64), Wang et al., 2014 (65), Wang et al., 2016 (66), Xia et al., 2013 (71), Xia et al., 2015 (70), Zhao et al., 2014 (76), Zhu et al., 2015 (77) |
| 3B | Shamsipur, 2016 (53) |

| | |
|---|---|
| 3C | Zuo et al., 2019 (79) |
| 4A | Bayramoglu et al., 2015 (3) |
| 0 | Zhang et al., 20121(75), Wood et al., 2012 (69) |

**Table S3. Percentage Publications with Unexplained Sequence Alterations within Each Phylogeny.**

| **Aptamer Phylogeny** | **Percentage Publications with Unexplained Sequence Alterations** |
|---|---|
| Thrombin aptamer | 13% (9 publications) |
| ATP aptamer | 87% (60 publications) |
| VEGF aptamer | 40% (10 publications, all did not provide sequence) |
| PDGF BB aptamer | 92% (65 publications) |
| Cocaine aptamer | 57% (33 publications) |
| Theophylline aptamer | 12% (13 publications) |
| Cox Lysozyme aptamer | 95% (57 publications) |
| Tran Lysozyme aptamer | 68% (15 publications) |
| Nucleolin aptamer | 37% (26 publications) |
| IgE aptamer | 46% (33 publications) |
| OTA aptamer | 23% (39 publications) |

**Table S4. Total Number of Unique Sequences and Sequences with Unexplained Alterations Identified.**

| Aptamer Phylogeny | Number of Unique Sequences (IDs) Identified | Number of Unique Undescribed Sequence Alterations | Notes |
|---|---|---|---|
| Thrombin aptamer | 10 | 9 | One originating sequence (explained ID) |
| ATP aptamer | 13 | 12 | One originating sequence (explained ID) |
| VEGF aptamer | 14 | 0 | All sequences explained or novel selection, 7 originating clones |
| PDGF BB aptamer | 35 | 32 | Three originating sequences |
| Cocaine aptamer | 31 | 21 | 8 described altered sequences and 2 originating clones |
| Theophylline aptamer | 8 | 7 | One originating sequence (explained ID) |
| Cox Lysozyme aptamer | 11 | 9 | One original sequence examined (6 clones originally). |
| Tran Lysozyme aptamer | 4 | 3 | One originating sequence (explained ID) |
| Nucleolin aptamer | 6 | 4 | One originating and one described sequence alteration |
| IgE aptamer | 15 | 13 | Two originating sequences. |
| OTA aptamer | 31 | 27 | Four originating sequences |
| Total Number of Sequences | 178 | 137 (77% of unique sequences) | 41 total described sequences, 23 originating clones total |

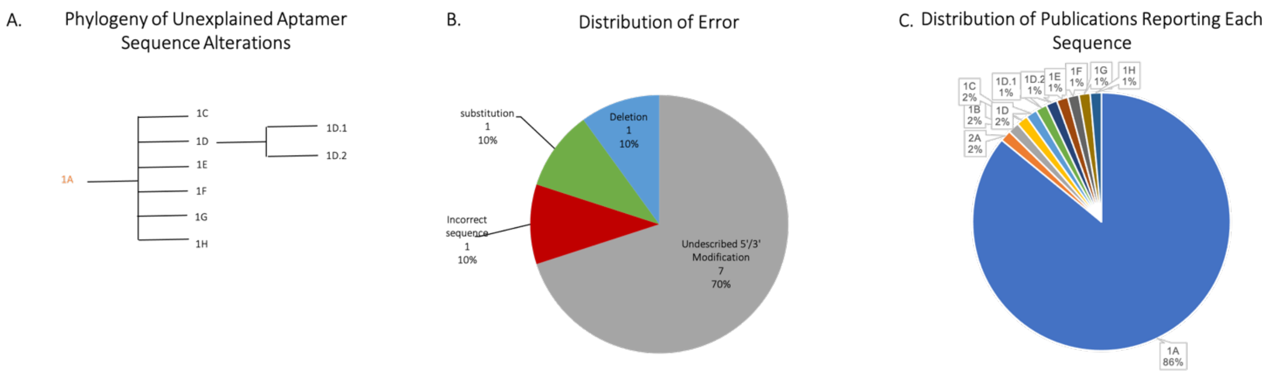


| ID | Publications Reporting Sequence |
|---|---|
| 1A | Avino et al., 2012 (1), Bern Aberem et al., 2006 (3), Bing et al., 2010 (4), **Bock et al., 1992** (5), Bompiani et al., 2012 (6), Boncler et al., 2001 (7), Cai et al., 2006 (8), Centi et al., 2008 (9), Chen et al., 2010 (10), Chen et al., 2010 (11), Cho et al., 2008 (12), Connor et al., 2006 (13), Dougan et al., 2003 (15), Evtugyn et al., 2009 (16), Fialova et al., 2006 (17), Gosai et al., 2016 (18), Gronewold et al., 2005 (19), Hamaguchi et al., 2001 (20), Hasegawa et al., 2008 (21), Holland et al., 2000 (22), Hoon et al., 2011 (23), Huang et al., 2010 (24), Huang & Zhu, 2009 (25), Ikebukuro et al., 2004 (26), Ikebukuro et al., 2006 (27), Jiang et al., 2014 (28), Jung et al., 2010 (29), Krauss et al., 2011 (31), Kretz et al., 2006 (32), Kupakuwana et al., (33), Lin et al., 2011 (38), Martino et al., 2006 (40), Mazurov et al., 2011 (41), Mendelboum et al., 2008 (43), Musumeci et al., 2012 (44), Nagatoishi et al., 2011 (45), Nagatoishi et al., 2007 (46), Najafi-Shoushtari et al., 2007 (47), Nimjee et al., 2009 (48), Pagano et al., 2008 (50), Padmanabhan et al., 1993 (49), Pagba et al., 2010 (51), Park et al., 2013 (52), Pastemak et al., 2011 (53), Pica et al., 2013 (54), Rahman et al., 2009 (55), Rinker et al., 2008 (56), Russo Krauss et al., 2012 (57), Rye et al., 2001 (58), Sosic et al., 2011 (61), Tasset et al., 1997 (63), Tennico et al., 2010 (64), Wang et al., 1993 (65), Wang et al., 2008 (67), Wang et al., 2018 (69), Wang et al., 2011 (70), Yang et al., 2009 (72), Yigit et al., 2008 (73), Zavylova et al., 2013 (75), Zavyalova et al., 2011 (76), Zhang et al., 2011 (77) |
| 1B | Schlensog et al., 2004 (59) |
| 1C | Li et al., 2002 (36) |
| 1D | Bai et al., 2013 (2) |
| 1D.1 | Wang et al., 2015 (66) |
| 1D.2 | Yin et al., 2015 (74) |
| 1E | Wang & Wang, 2013 (68) |
| 1F | Li et al., 2010 (37) |
| 1G | Mccauly et al., 2003 (42) |
| 1H | Kang et al., 2008 (30) |
| 2A | Bock et al., 1992 (5) |

| *Node* | *#* | *Aptamer Sequence Reported* |
|---|---|---|
| *1A* | 46 | GGTTGGTGTGGTTGG |
| *1B* | 1 | GGT~~T~~GGTG**G**TGGT~~T~~G**TT** |
| *1C* | 1 | **T**GGTTGGTGTGGTTGG**T** |
| *1D* | 1 | **TTTTT**GGTTGGTGTGGTTGG |
| *1D.1* | 1 | **TTTTTTTT**GGTTGGTGTGGTTGG |
| *1D.2* | 1 | **TTTTTTTTTT**GGTTGGTGTGGTTGG |
| *1E* | 1 | **ACTGT**GGTTGGTGTGGTTGG |
| *1F* | 1 | **CCATCTCCACTT**GGTTGGTGTGGTTGG |
| *1G* | 1 | **CCACC**GGTTGGTGTGGTTG~~G~~ |
| *1H* | 1 | **GACAGACGATGTGCTGACTACT**GGTTGGTGAGGTTGG**GTAGTCAGCACATCGTCTGTC** |
| *2A* | 1 | GGTTGG |

**S1 | Phylogeny depicting unexplained aptamer sequence alterations introduced to the 15mer DNA thrombin binding aptamer (TBA15, Bock et al., 1992), collected December 2019.** 26 publications reported the same sequence. Unexplained insertions are bolded, unexplained substitutions are bolded and underlined, unexplained deletions are struck out and justified or explained alterations are in light grey. The number (#) column indicates the number of publications found reporting each sequence in our analysis.

Six alternative and unexplained sequences were found in subsequent publications from node 1A. In nodes 1D, 1D.1, 1D.2, with one publication per node, each sequence modification involved a PolyT tail of varying lengths, and in all of the publications, the 5' PolyT modification to the 15-mer original aptamer was not explained. Specifically, in the Bai et al. (2013) publication, there was a Poly5Tail (1D), in the Wang, Zhou et al. (2015) publication it was a Poly8 Tail (1D.1), and in the Yin et al. (2015) publication, it was a Poly10 tail (1D.2).
A Thymine base was added to both the 5' and 3' end of the original 15-mer aptamer sequence, resulting in a 17-mer sequence, in the Li et al. (2002) publication (1C). The purpose of these additions were not explained in the publication.
Finally, in the Schlensog et al. (2004) publication, a series of deletions, insertions, and one substitution was made (1B). This sequence varied the most from the original Bock et al. (1992) sequence and the purposes behind these modifications was nonexistent.

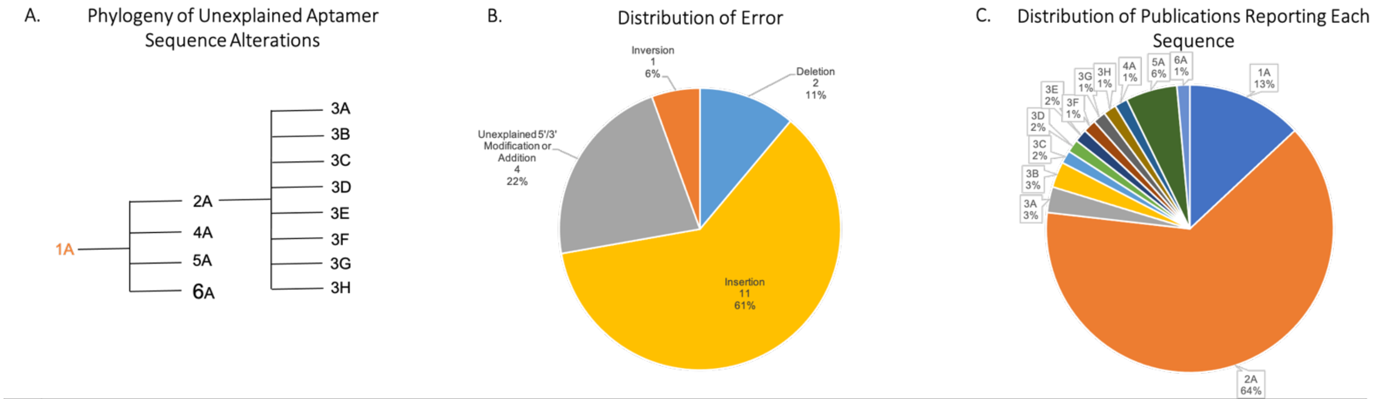


| ID | Publications Reporting Sequence |
|---|---|
| 1A | Arnaut et al., 2013 (1), Baaske et al., 2010 (2), Entzian et al., 2016 (9), **Huizenga et al., 1994** (15), Li et al., 2017 (24), Liao et al., 2016 (26), Urata et al., 2007 (45), Xia et al., 2013 (56), Zhao et al., 2015 (69) |
| 2A | Biniuri et al., 2018 (3), Bu et al., 2013 (4), Cai et al., 2011 (5), Chen et al., 2008 (6), Chen et al., 2008 (7), Cheng et al., 2013 (8), Goda et al., 2011 (11), Goda et al., 2012 (12), He et al., 2010 (13), Huo et al., 2016 (16), Jin et al., 2013 (18), Kashefi-Kheyrabadi et al., 2013 (20), Kashefi-Kheyrabadi et al., 2012 (19), Kong et al., 2013 (21), Li et al., 2008 (23), Liang et al., 2011 (25), Lin et al., 2014 (27), Liu et al., 2011 (28), Liu et al., 2014 (30), Merino and Weeks, 2003 (31), Modh et al., 2017 (32), Nutiu et al., 2003 (35), Nutiu et al., 2005 (36), Park et al., 2015 (40), Qiu et al., 2015 (41), Sitaula et al., 2013 (42), Song et al., 2012 (43), Wang et al., 2007 (47), Wang et al., 2005 (48), Wang et al., 2015 (49), Wang et al., 2012 (50), White et al., 2010 (54), Wu et al., 2013 (55), Xie et al., 2014 (57), Xu et al., 2010 (59), Yamana et al., 2003 (60), Yin et al., 2012 (61), Zhang et al., 2017 (66), Zhao et al., 2015 (67), Zhao et al., 2012 (68), Zhao et al., 2014 (70), Zhou et al., 2010 (71), Zhou et al., 2011 (72), Zuo et al., 2007 (73) |
| 3A | Wang et al., 2008 (51), Wang et al., 2008 (52) |
| 3B | Wang et el., 2014 (53), Xu et al., 2017 (58) |
| 3C | Liu et al., 2013 (29) |
| 3D | Zhang et al., 2010 (64) |
| 3E | Yu et al., 2014 (63) |
| 3F | Ying et al., 2011 (62) |
| 3G | He et al., 2011 (14) |
| 3H | Zhang et al., 2015 (65) |
| 4A | Jiang et al., 2012 (17) |
| 5A | Nielsen et al., 2010 (34), Ozalp et al., 2016 (37), Ozalp et al., 2010 (39), Tang et al., 2008 (44) |
| 6A | Li et al., 2012 (22) |

| *Node* | *#* | *Aptamer Sequence Reported* |
|---|---|---|
| *1A* | 9 | CCTGGGGGAGTATTGCGGAGGAAGG |
| *2A* | 44 | **A**CCTGGGGGAGTATTGCGGAGGAAGG**T** |
| *3A* | 2 | **T**GGAAGGAGGCGTTATGAGGGGGTCC**A** |
| *3B* | 2 | **AA**CCTGGGGGAGTATTGCGGAGGAAGG**T** |
| *3C* | 1 | **CCTCCTA**CCTGGGGGAGTATTGCGGAGGAAGG**TA** |
| *3D* | 1 | **TCTCTCA**CCTGGGGGAGTATTGCGGAGGAAGG**T** |
| *3E* | 1 | **A**CCTGGGGGAGTATTGCGGAGGAAGG**TTTT** |
| *3F* | 1 | **A**CCTGGGGGAGTATTGCGGAGGAAGG**TGTCACA**(A)$_{10}$ |
| *3G* | 1 | **A**CCT**TCCT**GGGGGAGTATTGCGGAGGAAGG**T** |
| *3H* | 1 | **A**CCTGGGGGA~~G~~TA**A**TTGCGGAGGAAGG**T** |
| *4A* | 1 | CCTGGGGGA~~G~~TATTGCGGAGGAAGG |
| *5A* | 4 | **CA**CCTGGGGGAGTATTGCGGAGGAAGG**TT** |
| *6A* | 1 | **GCA**CCTGGGGGAGTATTGCGGAGGAAGG**T** |

**S2 | Phylogeny depicting unexplained aptamer sequence alterations introduced to the 24mer DNA Adenosine Triphosphate (ATP) binding aptamer (DH29.36, Huizenga et al., 1994)**. Unexplained insertions are bolded, unexplained substitutions are bolded and underlined, unexplained deletions are struck out and justified or explained alterations are in light grey. The number (#) column indicates the number of publications found reporting each sequence in our analysis.

**A.** The DNA ATP binding aptamer (ABA) was discovered by Huizenga et al. (1994) and a minimized variant, or conserved sequences, was determined and folding structure was predicted **(ID 1)**. Urata et al. subsequently describe three minimized variants, the 25mer described by Huizenga et al., a 23mer cutting 1 nt at the 5' and 3' ends, and one adding an A to the 5' and T to the 3'. However, several groups **(2A)** use the 27mer without citing Urata's work minimizing the variant or a justification for using this over the 25mew. Xia et al. use the minimized variant described by Urata et al. citing their finding that the 23mer has been shown to have a stronger structural response to ATP and this is well suited for FRET characterization. Several publications include additional sequence information errors in addition to the use of the 27mer. Wang et al., 2008 published the 27mer, but inverted the sequence giving the 3' to 5' sequence labeled as 5' to 3' **(3A)**. Wang et al., 2014 added an additional A to the 5' end **(3B)**. Liu et al. (2013) **(3C)** introduced an additional CCTCCT to the 5' end and an A to the 3' end without explanation of their purpose. The added A and T are presumably for stability, and the additional 5' modification was likely created for the creation of a duplex structure or a hairpin that then changes conformation upon binding to ATP making it vulnerable to Exo III degradation. Though the sequence chose should be thoroughly described. Zhang et al. (2010) **(3D)** include an additional TCTCTC at the 5' end of the sequence. The linker is presumably to maintain attachment to the carbon nanotube matrix when binding ATP, but this is not stated clearly. Yu et al. 2010 include an additional 3 T's at the 3' without justification **(3E)**, and Ying et al. (2011) added a stretch of nucleotides to the 3' end before the stretch of adenosines without justification **(3F)**. He et al. (2011) **(3G)** include a 4nt insertion in the center of the aptamer sequence. Finally, Zhang et al. (2015) provide a sequence with a deletion and insertion.

Importantly, although many of these mutations **(2A, 3B-F)** most likely served a purpose within the application, the modifications made to the original sequence were not described or cited. Modh et al. (2017) give the sequence with added A and T for stability in a split aptamer design and cite the sequence correctly in the text and in most placed in their figure but incorrectly cite the sequence in one of the tertiary structures, deleting a G in the middle of the stem. While this was not cited as an error in the phylogeny due to the presence of correct sequence information in all other places, if this had been the only depiction of the sequence, an error could have been caused. Interestingly, in this phylogeny, no papers did not provide sequence information.

One paper was excluded from the analysis (Mukhurjee et al., 2015) because they used the wild type aptamer sequence cited in the Huizenga paper with the primer sequences included rather than the ATP aptamer they conclude/minimize. **B.** The Distribution of type of errors causing undescribed aptamer mutations are shown, with the predominant cause of sequence error belonging to the sequence information category including insertions and deletions. **C.** The number of uses of each sequence in the reviewed publications is shown, with a predominant use (61% or 31 publications) of node 2A. While Urata et al. (2007) describe a minimized variant and the utility of the 5' A and 3' T addition, several publications fail to make this justification or were published before the description of this aptamer variant.

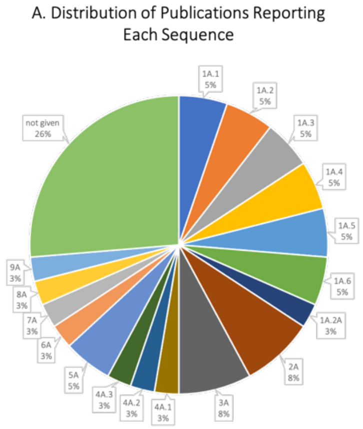


| ID | Nucleic Acid Type | Name | # Uses | Aptamer Sequence Reported (5′-3′) | Publications Reporting Sequence |
|---|---|---|---|---|---|
| 1A.1 | RNA | 100t (family 1) | 2 | NUGAUGVNCAUCCGN | **Jellinek et al., 1994 (34),** Libby, 2010 **(42),** Zhang & Yadavalli, 2010 **(80),** Li et al., 2007 **(41),** Ahirwar et al., 2016 **(1)** |
| 1A.2 | RNA | 44t (family 2) | 2 | AAUGCCGASSSSUUGAUGGGUU | Jellinek et al., 1994 **(34),** Libby, 2010 **(42)** |
| 1A.3 | RNA | 12t (family 3) | 2 | RDGGGAACCUGCGUYUCGGCACCHY | Jellinek et al., 1994 **(34),** Libby, 2010 **(42)** |
| 1A.4 | RNA | 40t (family 4) | 2 | HHGGUUGAGUCUGUCCCDD | Jellinek et al., 1994 **(34),** Libby, 2010 **(42)** |
| 1A.5 | RNA | 84t (family 5) | 2 | NRUAGUUGGNNNCUNSUNNNCGCCGUN | Jellinek et al., 1994 **(34),** Libby, 2010 **(42),** Potty et al., 2008 **(58)** |
| 1A.6 | RNA | 126t (family 6) | 2 | RSGGUUUCRUGUGKRGACUMUGCCGCGGCCSU | Jellinek et al., 1994 **(34),** Libby, 2010 **(42)** |
| 1A.2A | RNA | 44t.mab1 | 1 | GGAAGCUUGAUGGGUGACACACGUCAUGCCGAGCU**GGCUUCC** | Hall et al., 2007 **(27)** |
| 2A | DNA | N/A | 3 | CCGTCTTCCAGACAAGAGTGCAGGG | U.S. Patent No. 6,051,698, 2000 **(30),** Wang and Yadavalli, 2014 **(74),** Hunt., 2017 **(28),** Potty et al., 2008 **(58)** |
| 3A | DNA | V7t1 | 3 | TGTGGGGGTGGACGGGCCGGGTAGA | Eremeeva et al., 2019 **(20),** Lönne et al., 2015 **(45),** Chen, Chernis, et al., 2003 **(13)** |
| 4A.1 | RNA | ARC224 | 1 | CGAUAUGCAGUUUGAGAAGUCGCGCAUUCG | Burmeister et al. 2004 **(11)** |
| 4A.2 | RNA | ARC245 | 1 | AUGCAGUUUGAGAAGUCGCGCAU | Burmeister et al. 2004 **(11)** |
| 4A.3 | RNA | ARC259 | 1 | ACGCAGUUUGAGAAGUCGCGCGU | Burmeister et al. 2004 **(11)** |
| 5A | RNA | NX-107 | 1 | ACCCUGAUGGUAGACGCCGGGGUG | Green et al., 1995 **(26)** |
| 6A.1 | RNA | t22-OMe | 1 | **GCG**G**U**A**GGAAGAAUUGGAAGCGC** | Ruckman et al., 1998 **(62)** |
| 6A.2 | RNA | t2-OMe | 1 | **GCGA**A**CCGAUGGAAUUUUU**G**GACGCUCGC** | Ruckman et al., 1998 **(62)** |
| 6A.3 | RNA | t-44-OMe | 1 | **CGG**AA**UCAGUGAAUGCUUAUACAUCCG** | Ruckman et al., 1998 **(62)** |
| 7A | RNA | ERaptR3 | 1 | GGGGUCAAGGUGACCCC | Ahirwar et al., 2016 **(1)** |
| 8A.1 | RNA | 2-21 | 1 | CUUGUUGUGCUCCACGGUUAACUAACGAGCGUACAUGUGCAUUG**AAGAGACGAGUGUCAUAGCGAG** | Eremeeva et al., 2019 **(20)** |
| 8A.2 | RNA | 4-6 | 1 | CUUGUUGUGCUCCACGGUUAUAUUGCGGCAUUACUGGUACCAUG**AAGAGACGAGUGUCAUAGCGAG** | Eremeeva et al., 2019 **(20)** |
| 8A.3 | RNA | 2-15 | 1 | CUUGUUGUGCUCCACGGUUACACACGUGAUACGGAUAUGGUUAG**AAGAGACGAGUGUCAUAGCGAG** | Eremeeva et al., 2019 **(20)** |
| 9A | RNA | Macugen™ | 1 | CGGAAUCAGUGAAUGCUUAUACAUCCG dT28 | Libby, 2010 **(42)** |
| 0 | - | - | 10 | Not provided | Lee et al., 2005 **(40),** Trujillo et al., 2007 **(71),** Chen, Nakamoto, et al., 2012 **(15),** Bell, Lynam, et al., 1999 **(6),** Meek et al., 2016 **(48),** Chen, Yang-Sung, et al., 2018 **(14),** Shukla et al., 2007 **(64),** Chelyapov, 2006 **(12),** Mori et al., 2010 **(51),** Ruff et al., 2010 **(63)** |

300 search results narrowed to 152 with search terms "aptamer""vegf"; 80 papers reviewed, 26 were usable; 0 were unexplained publications; 10 publications did not report a sequence.

For the Ruckman aptamer:
**red bolded** indicates 5-iodo-U substitutions, black bolded indicates 2' -OMe purines

The modifications to the Macugen sequence can be seen in the above figure

The first 20 nt of the Eremeeva aptamer at the 5′-end correspond to the 2′-OMe-RNA primer sequence **(purple)** followed by a 24 nt random HNA region (green) and 22 nt 3′-HNA primer region **(blue).**

**bolded orange** indicates a described addition, Red highlighted indicates a described substitution.

R= A or G; Y= C or U; K= G or U; M= A or C; S = G or C; D= A, G, or U; H= A, U, or C; V= G, A, or C; N= any base

**S3 | Usage of the RNA vascular endothelial growth factor (VEGF) binding aptamers (family 1 (15mer), family 2 (22), family 3 (25mer), family 4 (19mer), family 5 (27mer), and family 6 (32mer), Jellinek et al., 1994), collected 1/22/20.** Search results up until 78 publications were analyzed however only 26 publications were usable. The remaining 74 publications were not usable in accordance to our protocol since these consisted of reviews, written in languages other than English, or were irrelevant publications. Additionally, towards the 78th search results repeats of previous publications were seen and specifically there were 5 publications that were a repeat of a previous result. Among the 26 publications, 10 publications did not report a sequence.

Among the publications, what sets VEGF apart from other aptamer phylogeny analysis is that most publications reported novel sequences such that very few publications reported the same related sequences. Additionally, among the publications that did report the same sequences, none had any unexplained modifications. Therefore, there is a strong distinction in this VEGF phylogeny such that in comparison to other aptamer analysis, there was much more novel selections than application papers.
The Jellinek et al., 1994 publication multiple clonal sequences (**seen in part C)** of the VEGF aptamer are clearly listed from six different families. Secondary structures of each of the family sequences were clearly reported. These familial sequences are displayed in nodes **1A.1-1A.6.** Additionally, these sequences are also listed in the Libby, 2010 publication with the predicted secondary structures clearly and with no unexplained modifications.
The node **1A.1** was listed in four publications in addition to the Jellinek et al., 1994 publication. These four publications reported the same variations of the sequence which was 5'-CCGGUAGUCGCAUGGCCCAUCGCGCCCGG-3'.
It is interesting to note that in the Potty et al., 2009 publication, it reports the same sequence seen in node **1A.5**; however, it transforms the RNA to a DNA sequence. This transformation is clearly listed and the publications explains that the sequence corresponds to the DNA analog of the complement of the 84t family.
In the node **1A.2A,** the Hall et al., 2007 reported the sequence seen in node **1A.2**, with an explained 3' addition to the 44t aptamer.
In the U.S. Patent No. 6,051,698, 2000 publication, a DNA sequence is clearly reported and patented. Additionally, two consensus sequences are also clearly reported. This sequence is also clearly reported in three subsequent modifications with no unexplained changes **(2A)**.
In the Nonaka et al., 2010 publication, an original aptamer sequence was found, although this publication was not a search result due to the parameters of our protocol, the sequence found in this publication was reported in three publications **(3A).**
Three RNA VEGF aptamer sequences were reported along with their predicted secondary structures in the Burmeister et al., 2005 publication **(4A.1-4A.3).** Sequence **4A.1** is the incomplete minimized variant of sequence **4A.2.** In sequence **4A.3**, there is a substitution of an A-U base pair for a G-C pair in the terminal region from the sequence **4A.2.**
The Green et al., 1995 publication, found a novel 24 nucleotide sequence RNA aptamer, which is seen in node **5A.** This sequence did not have a predicted secondary structure reported, however binding analysis was performed and it was determined that the RNA aptamer binds to VPF/VEGF with high affinity and specificity.
The Ruckman et al., 1998 publication found three truncated ligands, which are clearly listed in nodes **6A.1-6A.3.** These aptamers were chemically synthesized and the substitutions that correspond to each sequence changes are indicated in the table descriptions.
In node **7A,** a novel RNA aptamer was discovered and reported clearly in the Ahirwar et al., 2016 publication. Here, binding conditions are clearly listed however a secondary structure was not predicted. Instead, a 3D analysis of the aptamer complex was predicted in the publication.
In the Eremeeva et al., 2019 publication, three RNA aptamer sequences were discovered **(8A.1-8A.3).** Here, the aptamer sequences are clearly reported with the first 20 nucleotide region on the 5' end corresponding to the 2'-OMe-RNA primer sequence, which was followed by a 24 nucleotide random RNA region, and finally at the 3' end, a 22 nucleotide primer region is listed.
In addition to reporting the six familial aptamer sequences of the Jellinek et al., 1994 publication, the Libby, 2010 publication also reported a novel RNA sequence **(9A).** This RNA aptamer was named Macugen and unique chemical modifications performed during analysis was clearly reported and listed. The chemical modifications and structure of Macugen can be seen in part

26% of publications reviewed did not provide the aptamer sequence, which is particularly problematic due to the large number of original sequences that bind VEGF. Further, this phylogeny illuminates the importance of characterization of the aptamer sequences and selection based on the properties most beneficial for the application.

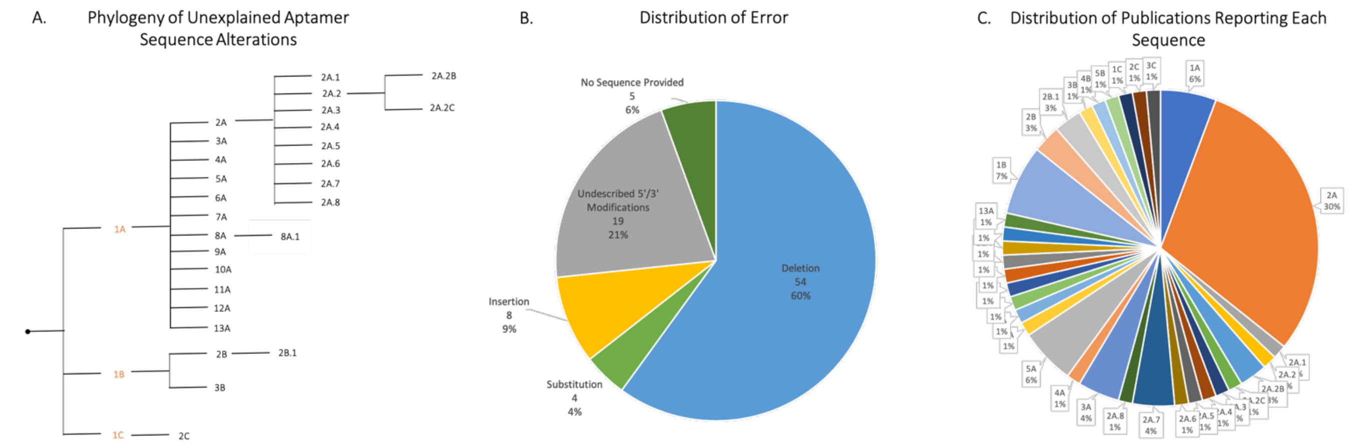


| ID | Publications Reporting Sequence |
|---|---|
| 1A | **Green and Jellinek. 1996 (14)**, Vicens et al., 2005 **(51)**, Vu et al., 2016 (54), Yang et al., 2005 **(59)** |
| 2A | Fang et al., 2001 **(9)**, Fang et al., 2003 **(10)**, Fisher et al., 2012 (11), Huang et al., 2005 (20), Huang et al., 2007 **(19)**, Jiang et al., 2004 (23), Jin et al., 2014 (24), Kim et al., 2009 (25), Lai et al., 2007 **(28)**, Li et al., 2016 (33), Liao et al., 2007 **(35)**, Neumann et al., 2009 (41), Ruslinda et al., 2012 **(45)**, Song et al., 2012 **(47)**, Wang et al., 2009 **(55)**, Wang et al., 2015 (57), Yang et al., 2014 **(60)**, Zhang et al., 2012 (67), Zhang et al., 2015 (65), Zhang et al., 2009 **(64)**, Zhou et al, 2006 **(69)** |
| 2A.1 | Hu et al., 2014 **(18)** |
| 2A.2 | Huang K et al., 2016 **(21)** |
| 2A.2B | Tang et al., 2012 **(50)**, Ma et al., 2014 **(40)** |
| 2A.2C | Huang et al., 2008 **(22)** |
| 2A.3 | Yi et al., 2013 **(62)** |
| 2A.4 | Li et al., 2013 **(31)** |
| 2A.5 | Wang et al., 2013 **(56)** |
| 2A.6 | Ye et al., 2016 **(61)** |
| 2A.7 | Fang et al., 2015 **(8)**, Liu et al., 2014 **(37)**, Zhang et al., 2015 **(66)** |
| 2A.8 | Liu et al., 2012 **(36)** |
| 3A | Csordas et al., 2009 **(7)**, Kim et al., 2011 **(26)**, Gao et al., 2018 **(13)** |
| 4A | Vu et al., 2017 **(53)** |
| 5A | Vu et al., 2018 (52), Battig et al, 2012 **(2)**, Soontornworajit et al., 2010 **(49)**, Zhang et al., 2018 **(68)** |
| 6A | Lai et al., 2017 **(27)** |
| 7A | Li et al., 2013 **(32)** |
| 8A | Battig et al., 2014 **(1)** |
| 8A.1 | Battig et al., 2014 **(1)** |
| 9A | Sennino et al., 2007 **(46)** |
| 10A | Bi et al., 2014 **(3)** |
| 11A | Chhabra et al., 2007 **(5)** |
| 12A | Liang et al., 2013 **(34)** |
| 13A | Chang et al., 2013 **(4)** |
| 14A | Soontornworajit et al., 2010 **(48)** |
| 1B | Green & Jellinek 1996 **(14)**, Guo & Zhao 2016a, 2016b (16) (17), Guo et al., 2016 (15), Song et al., 2012 **(47)** |
| 2B | Fredriksson et al., 2002 **(12)**, **Zhou et al., 2007 (70)** |
| 2B.1 | Zhu et al., 2010 **(71)**, Zhu et al., 2012 **(72)** |
| 3B | Li et al., 2012 **(30)** |
| 4B | Xie et al., 2010 **(58)** |
| 5B | Lv et al., 2015 (39) |
| 1C | Green, Jellinek 1996 **(14)** |
| 2C | Zhang et al., 2012 **(63)** |
| 3C | Zhang et al., 2009 **(64)** |
| 0 | Leppanen et al., 2000 (29), Lu et al., 2010 **(38)**, Ostendorf et al., 2001 (42), Pietras et al., 2002 **(44)** |

| *Node* | *#* | *Aptamer Sequence Reported* |
|---|---|---|
| *1A* | 4 | CACAGGCTACGGCACGTAGAGCATCACCATGATCCTGTG |
| *2A* | 21 | ~~CA~~CAGGCTACGGCACGTAGAGCATCACCATGATCCTG~~TG~~ |

| | | |
|---|---|---|
| *2A.1* | 1 | **AAAAAA**~~CA~~CAGGCTACGGCACGTAGAGCATCACCATGATCCTG~~TG~~ |
| *2A.2* | 1 | ~~CA~~CAGGCTACGGCACGTAGAGCATCACCATGATCCTG~~TG~~**TTTTT** |
| *2A.2B* | 2 | ~~CA~~CAGGCTACGGCACGTAGAGCATCACCATGATCCTG~~TG~~**TTTTTT** |
| *2A.2C* | 1 | ~~CA~~CAGGCTACGGCACGTAGAGCATCACCATGATCCTG~~TG~~**TTTTTTT**CAACACACCTTAACAC |
| *2A.3* | 1 | ~~CA~~CAGGCTACGGCACGTAGAGCATCACCATGATCCTG~~TG~~**CCCAGGTTCTCT($T_{10}$)-HS** |
| *2A.4* | 1 | ~~CA~~CAGGCTACGGCACGTAGAGCATCACCATGATCCTG~~TG~~**AAAAAAAAAAAAAAAAAAAAAAAA** |
| *2A.5* | 1 | **C**~~CA~~CAGGCTACGGCACGTAGAGCATCACCATGATCCTG~~TG~~**G** |
| *2A.6* | 1 | **TCTAGACATTCATCCT**~~CA~~CAGGCTACGGCACGTAGAGCATCACCATGATCCTG~~TG~~ |
| *2A.7* | 3 | **TTTTTTTTTT**~~CA~~CAGGCTACGGCACGTAGAGCATCACCATGATCCTG~~TG~~ |
| *2A.8* | 1 | ~~CA~~CAGGCTACGGCACGTAGAGCATCACCATGATCCTG~~TG~~**A** |
| *3A* | 3 | CACAGGCTACGGCACGTAGAGCATCACCATGATCCTGTG**T** |
| *4A* | 1 | C~~A~~CAGGCTACGGCACGTAGAGCATCACCATGATCCTGTG |
| *5A* | 4 | ~~C~~ACAGGCTACGGCACGTAGAGCATCACCATGATCCTG~~TG~~ |
| *6A* | 1 | ~~C~~ACAGGCTACGGCACGTAGAGCATCACCATGATCCTGTG**ACTAACCTG**($G_4$) |
| *7A* | 1 | ~~C~~ACAGGCTACGGCACGTAGAGCATCACCATGATCCTGT~~G~~ |
| *8A* | 1 | **GCGATACTC**CACAGGCTACGGCACGTAGAGCATCACCATGATCCTG~~TG~~ |
| *8A.1* | 1 | **GCGATACTC**CACAGGCTACGGCACGTAGAGCATCACCATGATCC**CA**~~TG~~ |
| *9A* | 1 | dC**dC**dCdAdGdGdCTdAdCmG~~G~~-HEG-dC~~AC~~dGTdAmGdAmGdCdAm**Um**CmA-HEG~~CCA~~TdGdATmCdCTmGmG**mGdT** |
| *10A* | 1 | C**T**CAGGCTACGGCACGTAGAGCATCACCATGATCCTG**A**G |
| *11A* | 1 | CACAGGCTACGGCACGTAGAGCATCACCATGATCCTGT~~G~~ |
| *12A* | 1 | CACAGGCTACGGCACGTAGAGCATC**ATC**ACCATGATCCTGTG**T** |
| *13A* | 1 | ~~CAC~~AGGCTACGGCACGTAGAGCATCACCATGATCCTG~~TG~~ |
| *14A* | 1 | GCGATACTCCACAGGCTACGGCACGTAGAGCATCACCATGATCC~~TGTG~~ |
| *1B* | 5 | TACTCAGGGCACTGCAAGCAATTGTGGTCCCAATGGGCTGAGTA |
| *2B* | 1 | TACTCAGGGCACTGCAAGCAATTGTGGTCCCAATGGGCTGAGTA**T** |
| *2B.1* | 2 | **AAAAAAAAAA**TACTCAGGGCACTGCAAGCAATTGTGGTCCCAATGGGCTGAGTA**T** |
| *3B* | 1 | ~~T~~ACTCAGGGCACTGCAAGCAATTGTGGTCCCAATGGGCTGAGTA |
| *4B* | 1 | TACTCAGGGCACTGCAAGCAATTGTGGTCCCAATGGGCTGAGTA**TTTTTT** |
| *5B* | 1 | TACTCAGG~~G~~CACTGCAAGCAATTGTGGTCCCAATGGGCTGAGTA |
| *1C* | 1 | TGGGAGGGCGCGTTCTTCGTGGTTACTTTTAGTCCCG |
| *2C* | 1 | TGGG**C**G~~GG~~CG~~CG~~TTCTTCGTGGTTACTTTTAGTCCCG |
| *3C* | 1 | **A**GGGCG~~GG~~CG~~CG~~TTCTTCGTGGTTACTTTTAGTCCCG |
| *0* | 5 | No sequence provided |

**S4 | Phylogeny depicting unexplained aptamer sequence alterations introduced to the three DNA platelet-derived growth factor (PDGF-BB) binding aptamers: 36t a 36mer (node 1A), 41t a 41mer (1B), and 20t a 20mer (1C), Green and Jellinek, 1996.** Unexplained insertions are bolded, unexplained substitutions are bolded and underlined, unexplained deletions are struck out and justified or explained alterations are in light grey. The number (#) column indicates the number of publications found reporting each sequence in our analysis.

The above phylogeny documents the alteration to the PDGF ssDNA aptamer sequences originally identified by Green and Jellinek 1996 using the SELEX methodology. Green and Jellinek identified three different minimized variant sequences that exhibited binding to PDGF BB **(1A, 1B, 1C)**. The most common deviation in the most commonly-reported sequence was cited first by Fang et al., 2001 **(2A);** this group reported a sequence that truncated the original sequence by two nucleotides on either end. Sequences stemming from this node **(2A.1-4)**, employed various unjustified 5' or 3' extensions. Other groups modified the original Green Jellinek sequence in various ways. Most notably, Sennino et al. **(9A)** made several unjustified substitutions in the aptamer sequence in a cancer application. Sequence modifications in **1B** and **1C** largely consisted of unexplained 5' or 3' additions and deletions of nucleotides in the middle of the minimized sequence. Of note, Zhang et al. 2009 and 2012 **(2C and 3C)**, used sequences that included deletions of nucleotides within the region. While this group was able to generate results from this error, target binding characteristics could have changed with the introduction of these errors.

The most commonly cited aptamer in this phylogeny **(1A)** was used in a variety of applications, likely because it was the original aptamer sequence described by Green and Jellinek that exhibited the highest binding affinity. Subsequent papers in this analysis typically cited using "the PDGF BB aptamer", which could refer to any of the three sequences identified by Green and Jellinek. Further clarification could have been made by citing the originally reported dissociation constant or the length of the minimized variant. 2 of the papers in this analysis did not report a sequence for the PDGF aptamer.

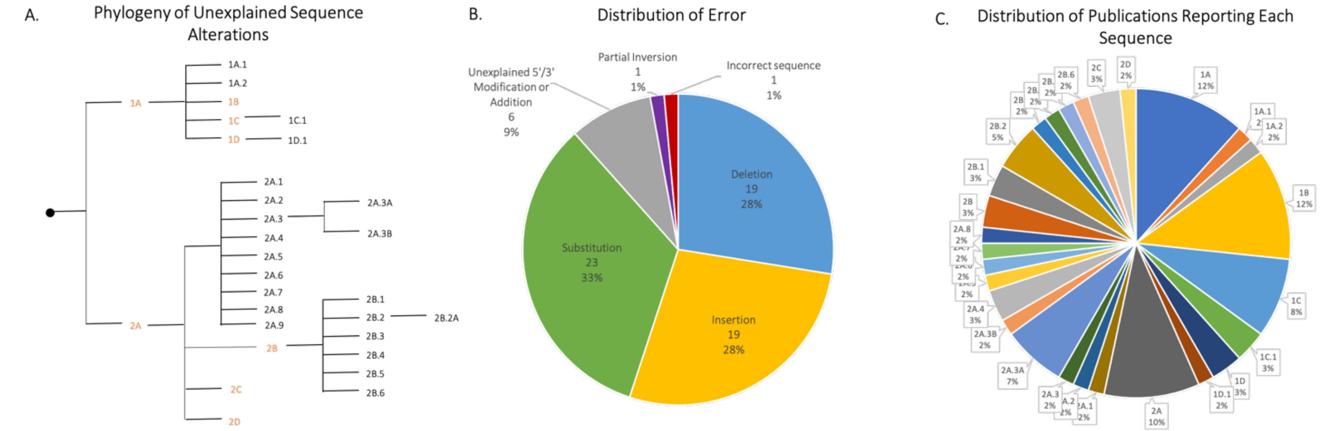


| ID | Publications Reporting Sequence |
|---|---|
| 1A | Baker et al., 2006 (1), Cekan et al., 2009 (2), Chen et al., 2008 (3), Li et al., 2007 (18), Liu et al., 2017 (20), Neves et al., 2010 (26), **Stojanovic et al., 2001** (40), Taghavi et al., 2014 (42), Zhou et al., 2011 (58) |
| 1A.1 | Yang et al., 2016 (48) |
| 1A.2 | Zhou et al., 2012 (57) |
| 1B | Baker at al., 2006 (1), Deng et al., 2012 (5), Li et al., 2008 (17), Roushani et al., 2015 (33), Roushani et al., 2015, (34), Swenson et al., 2009 (41), Zhang et al., 2011 (54) |
| 1C | Neves et al., 2015 (24), Neves et al., 2010 (26), Neves et al., 2017 (28), Reinstein et al., 2013 (31), Slavkovic et al., 2015 (39) |
| 1C.1 | Neves et al., 2016 (25), Neves et al., 2017 (29) |
| 1D | Neves et al., 2010 (26), Reinstein et al., 2013 (31) |
| 1D.1 | Grytz et al., 2016 (9) |
| 2A | Kang et al., 2011 (14), Madru et al., 2009 (22), Neves et al., 2010 (26), Roncancio et al., 2014 (32), Sachan et al., 2016 (35), Stojanovic et al., 2001 (40) |
| 2A.1 | Kawano et al., 2011 (15) |
| 2A.2 | Wang et al., 2018 (44) |
| 2A.3 | Sharon et al., 2009 (37) |
| 2A.3A | Freeman et al., 2009 (6), He et al., 2010 (10), Nie et al., 2013 (30), Shi et al., 2013 (38), Zhang et al., 2012 (53) |
| 2A.3B | Golub et al., 2009 (8) |
| 2A.4 | Li et al., 2013 (16), Wen et al., 2011 (46), Zhang et al., 2008 (55) |
| 2A.5 | Grytz et al., 2016 (9) |
| 2A.6 | Zhao et al., 2015 (56) |
| 2A.7 | Wu et al., 2010 (47) |
| 2A.8 | Zou et al., 2012 (59) |
| 2A.9 | Sassolas et al., 2010 (35) |
| 2B | Roncancio et al., 2014 (32), Zhang et al., 2009 (52) |
| 2B.1 | Li et al., 2011 (19), Ma et al., 2011 (21) |
| 2B.2 | Chen et al., 2016 (4), Hua et al., 2011 (12), Hua et al., 2011 (13) |
| 2B.2A | Yin et al., 2012 (47) |
| 2B.3 | Yan et al., 2010 (48) |
| 2B.4 | Ge et al., 2012 (7) |
| 2B.5 | Tang et al., 2016 (43) |
| 2B.6 | Hilton et al., 2011 (11) |
| 2C | Roncancio et al., 2014 (32), Yu et al., 2016 (51) |
| 2D | Neves et al., 2010 (26) |
| 3 | Wang et al., 2007 (45) |

| *Node* | *#* | *Aptamer Sequence Reported* |
|---|---|---|
| *1A* | 7 | GACAAGGAAAATCCTTCAATGAAGTGGGTC |
| *1A.1* | 1 | GACAAGGA**T**AAATCCTTCAATGAAGTGGGTCACTCATCTGTGA |
| *1A.2* | 1 | **A**GACAAGGAAAATCCTTCAATGAAGTGGGTC**TC** |
| *1B* | 7 | **A**GACAAGGAAAATCCTTCAATGAAGTGGGTC**G** |
| *1C* | 5 | **GGC**GACAAGGAAAATCCTTCAA**C**GAAGTGGGTC**GCC** |
| *1C.1* | 2 | **GGC**GACAAGGAAAATCCTT**G**AA**C**GAAGTGGGTC**GCC** |
| *1D* | 2 | GACAAGGAAAATCCTTCAA**C**GAAGTGGGTC |
| *1D.1* | 1 | GACAAG~~G~~AA**GCG**A**CGTCGC**T**T**CTTCAA**C**GAAGTGGGTC |
| *2A* | 6 | GGGAGACAAGGAAAATCCTTCAATGAAGTGGGTCGACA |
| *2A.1* | 1 | GGGAGACAAGGAAA~~A~~TCCTTCAATGAAGTGGGTCGACA |
| *2A.2* | 1 | GGGAG**T**CAAG~~G~~AA**C,** ACAGC**A**GGGTGAAGTAACTTC**T**T**G** |
| *2A.3* | 1 | GGGAG**T**CAAGGAA**CG**AA**TTCGT**TC~~C~~TTCAATGAAGTGGG**A**CGACA |
| *2A.3A* | 4 | GGGAG**T**CAAG~~G~~AA**CG**AA**TTCGT**TC~~C~~TTCAATGAAGTGGG**A**CGACA |
| *2A.3B* | 1 | GGGAG**T**CAAG~~G~~AA**CG**AA**TTCGT**TCGTT**CTT**CAATGAAGTGGG**A**CGACA |
| *2A.4* | 2 | GGGAG**T**CAAG~~G~~AA**CG**T**T**CTTCAATGAAGTGGG**A**CGACA |
| *2A.5* | 1 | **C**GG**C**GACAAG~~G~~AA**GC**GA**CGTCGC**T**T**CTTCAA**C**GAAGTGGGTCGAC**G** |
| *2A.6* | 1 | GGGAG**T**CAAG~~G~~AA**CTTTG**T**T**CTTCAAT~~GA~~AGTGGG**A**CGACA |
| *2A.7* | 1 | GGGAG**T**CAAG~~G~~AA**CCGG**T**T**CTTCAATGAAGTGG~~TC~~GACA |
| *2A.8* | 1 | GGGAG**T**CAAG~~G~~AA**CA**AA**GT**TC~~C~~TTCAATGAAGT**GT**GGG**A**CGACA |
| *2A.9* | 1 | GGGAGAC**TT**GGATAAATCC**AA**CAATGAAGTGGGTCGAGA |
| *2B* | 2 | GGGAGACAAGGAAAATCCTTCAATGAAGTGGGTC**TCCC** |
| *2B.1* | 2 | **GG**GGGAGACAAGGAAAATCCTTCAATGAAGTGGGTC**TCCC** |
| *2B.2* | 3 | GGGAGACAAGGA**T**AAATCCTTCAATGAAGTGGGTC**TCCC** |
| *2B.2A* | 1 | GGGAGACAAGGA**T**AAATCCTTCAATGAAGTGGGTC~~TCCC~~ |
| *2B.3* | 1 | **ACTCATCTGTGAATCTC**GGGAGACAAGGA**T**AAATCCTTCAATGAAGTGGGTC**TCCC** |

| | | |
|---|---|---|
| *2B.4* | 1 | **CCATA**GGGAGACAAGGA**T**AAATCCTTCAATGAAGTGGGTC**TCCC** |
| *2B.5* | 1 | TTTTTTTACTCATGGGAGACAAGGA**TT**AATCCTTCAATGAAGTGGGTC**TCCC** |
| *2B.6* | 1 | **ATCTC**GGGA**C**GACAAGGA**TTT**TCCTT~~T~~CAATGAAGTGGGTC**GTCCC** |
| *2C* | 2 | GGGAGACAAGGAAAATCCTTCAA**C**GAAGTGGGTC**TCCC** |
| *2D* | 1 | ~~G~~GGAGACAAGGAAAATCCTTCAATGAAGTGGGTCGAC~~A~~ |
| *3* | 1 | AGCAACCTCAAACAGACACCATGG |

**S5 | Phylogeny depicting unexplained aptamer sequence alterations introduced to the two DNA cocaine binding aptamers: MNS4.1 a 30mer (node 2A) and MNS7.9/MSN6 a 38mer (1A), Stojanovic et al., 2001.** Unexplained insertions are bolded, unexplained substitutions are bolded and underlined, unexplained deletions are struck out and justified or explained alterations are in light grey. The number (#) column indicates the number of publications found reporting each sequence in our analysis.

The above phylogeny follows the evolution of the MSN-4.1 **(2A)** and MNS-7.9/MN6 **(1A)** ssDNA cocaine aptamer sequences since their discovery in 2001 by Stojanovic, et al. The MSN-7.9 aptamer was constructed by truncating the MSN-4.1 aptamer and is described "with a shortened $S_1$ stem [and found] that it retained significant, albeit reduced, affinity for cocaine." They further modify MSN-7.9 with a 5'-fluorescein and 3' dabcyl to mitigate effects of decreased binding. A 32-mer **(1B)** with single nucleotide additions to both the 5' and 3' ends of the original MN6 sequence was first described by Baker, et al. Further, Neves, et al. describe two sequence variants that include a single nucleotide substitution in stem-loop 3, shifting a noncanonical G-T base pairing to G-C. In addition to this substitution, the MN4 sequence **(1C)** features three-nucleotide additions to both the 5' and 3' ends, while the MN19 sequence **(1D)** does not share this change. The original 2001 Stojanovic publication included an additional sequence, MNS 4.1 **(2A)**, which included a four-nucleotide addition to both the 5' and 3' ends. Roncancio, et al. describe a 38-mer **(2B)** replaced the final four nucleotides on the 3' end with TCCC for stabilization of the stem, and another 38-mer sequence **(2C)** with a single nucleotide substitution similar to the one described by Neves, et al. Finally, Neves, et al. described the MN1 sequence **(2D)** which featured three-nucleotide additions to both the 5' and 3' ends of the original MN6 sequence. All other changes noted in the phylogeny above made changes to the original MN6 or MNS 4.1 sequences without sufficient explanation of their modifications. One notable error made by Wang et al., 2018 **(2A.2)** used a split aptamer design, inverting one fragment, but not the other in addition to a deletion and five undescribed substitutions.
Upon review of the included publications for this analysis, it was noted that the downstream applications for the cocaine aptamer were almost exclusively sensor-based and diagnostic in nature, primarily using in-vitro methodology. The phylogeny above shows an unusually high rate of error relative to other aptamer targets that our group reviewed, which we speculate to be largely a product of the use of these aptamers as an auxiliary reagent rather than a central component of these groups' research. Further, some groups only reported oligonucleotide sequences within a figure showing the aptamer's secondary structure, which made the manual copying of the sequence tedious and prone to error. The orange ID's were aptamer modifications that were described and fully characterized.

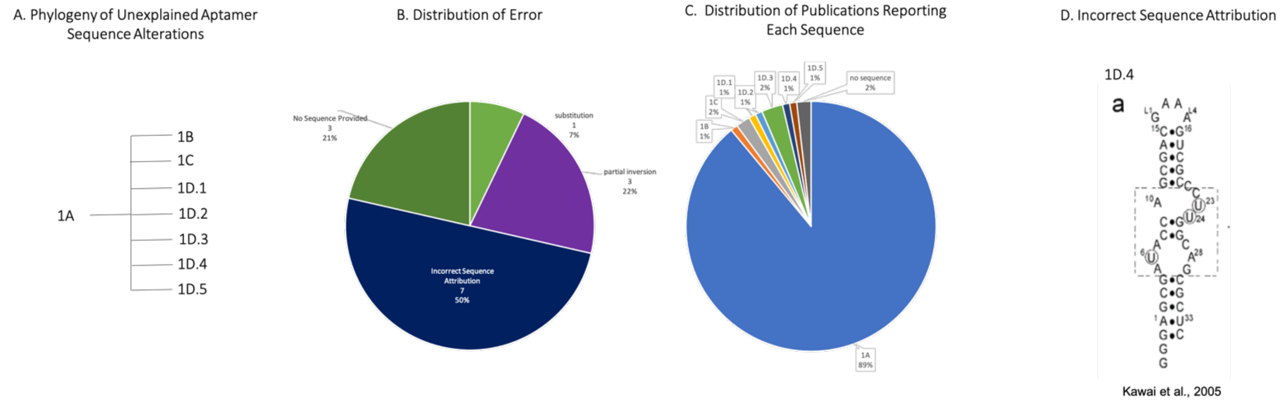


| ID | Publications Reporting Sequence |
|---|---|
| 1A | Amontov & Jäschke, 2006 (1); An et al., 2006 (2); Anderson & Mecozzi, 2005 (3); Ausländer et al., 2011 (4); Autour et al., 2019 (5); Brutscher et al., 1999 (6); Carrasquilla et al., 2012 (8); Chang et al., 2014 (9); Chávez et al., 2010 (10); X. Chen et al., 2018 (12); Chovelon et al., 2016 (13); Chushak et al., 2009 (14), Cui et al., 2013 (15); **De Silva & Walter, 2009 (16),** Dery et al., 2009 (17); Desai & Gallivan, 2004 (18); Domin et al., 2017 (19); Durand et al., 2016 (21); Endo et al., 2004 (25); Endoh & Sugimoto, 2013 (24); Espah Borujeni et al., 2016 (26); Feng et al., 2018 (28); Ferapontova et al., 2008 (31); Ferapontova & Gothelf, 2009 (30); Ferapontova et al., 2009 (29), Fowler et al., 2008 (32); Freedman et al., 2010 (33); Fujita et al., 2012 (34); Gouda et al., 2003 (36); Groher & Suess, 2014 (37); Helm et al., 2005 (42); **Jenison et al., 1994 (45)**; Jiang et al., 2015 (46); Jo et al., 2012 (48); Jose et al., 2001 (49); **Jucker et al., 2003 (50),** Katiyar et al., 2013 (51); Kertsburg & Soukup, 2002 (54); Kim et al., 2005, 2008 (55) (56); Kumar et al., 2009 (57); Lam & Joyce, 2011 (59); Latham et al., 2009 (60); P. S. Lau et al., 2010, 2013 (61) (62); Lee et al., 2010 (64); X. Li et al., 2013 (66); Liao et al., 2018 (67); Ling et al., 2016 (68); **Lou et al., 2019 (69), Luzi et al., 2003 (70),** Lynch et al., 2007 (71); Lynch & Gallivan, 2009 (72); Ma et al., 2017 (73), Michener & Smolke, 2012 (75); Nomura et al., 2013 (77); Ogawa, 2011, 2012 (78) (79); Ogawa et al., 2017, 2018 (81) (82); Ogawa & Maeda, 2007 (80); Ogawa & Tabuchi, 2015 (83); Page et al., 2018 (84); Patel & Suri, 2000 (85); Pei & Stojanovic, 2008 (86) Petermeier & Jäschke, 2008 (87); Qi et al., 2012 (88); Robertson & Ellington, 2000 (90); Sarpong & Datta, 2012 (92); Sekella et al., 2002 (94); Sharma et al., 2008 (95); Sibille et al., 2001 (96), Song et al., 2014 (97); Soukup et al., 2000 (98); Soukup & Breaker, 1999a (99); Tang & Breaker, 1997 (101); Thompson et al., 2002 (102); Topp & Gallivan, 2007, 2008a, 2008b (105) (106) (107); Townshend et al., 2015 (108); Tuleuova et al., 2008 (109); Wachsmuth et al., 2013, 2015 (110) (111); J. Wang et al., 2018 (112); S. Wang et al., 2008 (113); S. Wang & White, 2007 (114); Warfield & Anderson, 2017 (115); Wieland & Hartig, 2008 (116); Xie & Walton, 2009 (118); Zhang et al., 2017 (119); Zhou et al., 2015 (121); Zimmermann et al., 1997, 1998, 2000 (122) (123) (124) |
| 1B | Edwards & Baeumner, 2007 (22) |
| 1C | Dong & Zhao, 2013 (20), Wu et al., 2019 (117), Zhao & Yang, 2010 (119) |
| 1D.1 | Rankin et al., 2006 (89) |
| 1D.2 | Suess et al., 2004 (100) |
| 1D.3 | Endoh et al., 2009 (25); Jo & Shin, 2009 (47); Rudolph et al., 2013 (91) |
| 1D.4 | Kawai et al., 2005 (52) |
| 1D.5 | Lam & Joyce, 2009 (58) |
| 0 | Feng et al., 2016 (27), Harbaugh et al., 2009 (40); Heus, 1997 (43) |

*Node # Aptamer Sequence Reported*

| | | |
|---|---|---|
| *1A* | 99 | (Nx) AUACCA (Nx) CUUGG (C/A) AG (Nx) |
| *1B* | 1 | (AAGUG) **<u>C</u>**UACCA (GCAUCGUCUUGAUGC) CCUUGGCAG (CACUUCA) |
| *1C* | 3 | $(CH_2)_6$ CCUUGGAAG *(CC)* 5′ *(GG)* AUACCA |
| *1D.1* | 1 | (GGC) **G**AUACCA**G** (CCGAAAGG) **C**CCUUGGCAG**C** (GUC) |
| *1D.2* | 1 | (Nx) ~~A~~UACCA (GCCGAAAGGCC) CUUGGCA~~G~~ (Nx) |
| *1D.3* | 3 | *(Nx)* **G**AUACCA *(Nx)* CCUUGGCAG**C** *(Nx)* |
| *1D.4* | 1 | (GGGAGCG) AUACCA (GCGACGAAAGUCGC) **C**CUUGGCAG (CGCUC) |
| *1D.5* | 1 | (UCGG) AUACCA (GCCGAAAGGC) CCUUGGCAG (CUUGA) |
| *0* | 3 | No sequence provided |

**S6 | Phylogeny depicting unexplained aptamer sequence alterations introduced to the RNA theophylline binding aptamer (the conserved theophylline binding region, Jenison et al., 1994).** Unexplained insertions are bolded, unexplained substitutions are bolded and underlined, unexplained deletions are struck out and justified or explained alterations are in light grey. The number (#) column indicates the number of publications found reporting each sequence in our analysis. **A.** Jenison et al. 1994 selected a minimized variant aptamer, or a conserved theophylline binding region, underlined in green (1A). Edwards and Baeumner (2007) made an undescribed substitution to the 5′ A of region 1 (1B). Several publications (1C) use a split aptamer design, anchoring the second region to the surface, effectively inverting the order of the conserved regions. While this will still form the correct stem structure, it inverts the order in which the regions are from 5′ to 3′. Finally, Rankin et al. (2006), Seuss et al. (2004) both contained illustrative errors, shown in part D, that incorrectly box the incorrect "conserved theophylline binding region". While this does not change the effectiveness of the aptamer, it is an undescribed illustrative error. Endoh et al. (2009), Jo and Shin (2009), Rudolph et al. (2013) all contain the same illustrative error (1D.3), adding a G to the 5′ of region 1 and a C to the 3′ of region 2. Kawai et al. (2005) contains a single addition on the 5′ end of region 2, and finally Lam and Joyce (2009) provide the correct sequence (1D.5), but do not clearly indicate the 5′ or 3′ termini, which could result in subsequent inversions. While in the context of their application, the direction could be inferred, but was not made explicitly clear. **D.** Reprinted in part with permission from Kawai, R., Kimoto, M., Ikeda, S., Mitsui, T., Endo, M., Yokoyama, S., & Hirao, I. (2005). Site-specific fluorescent labeling of RNA molecules by specific transcription using unnatural base pairs. *Journal of the American Chemical Society*, *127*(49), 17286-17295. Copyright 2004 American Chemical Society.

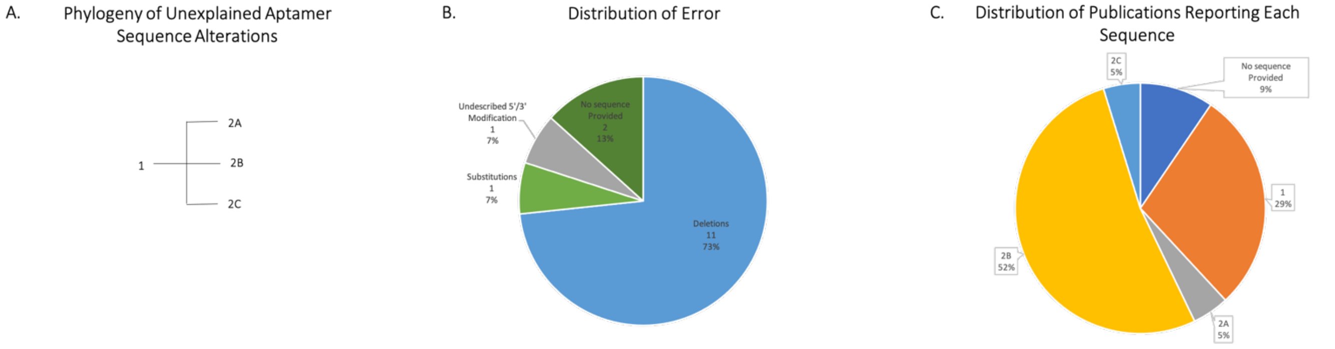


| ID | Publications Reporting Sequence |
|---|---|
| 1 | Khedri et al., 2018 **(15)**, Lam et al., 2018 **(16)**, Meyer, 2014 **(24)**, **Tran et al., 2010 (36)**, Weng et al., 2016 **(46)**, Wood et al., 2014 **(47)** |
| 2A | Zou et al., 2012 **(55)** |
| 2B | Boushell et al., 2017 **(5)**, Han et al., 2012 **(12)**, Liu et al., 2015 **(20)**, Luu et al., 2019 **(21)**, Mishra et al., 2017 **(26)**, Ocaña et al., 2015 **(28)**, Ocaña et al., 2015 **(29)**, Titoiu et al., 2019 **(35)**, Truong et al., 2013 **(38)**, Wang et al., 2011 **(43)**, Xie et al., 2017 **(50)** |
| 2C | Wang et al., 2018 **(42)** |
| 0 | Wood et al., 2012 **(48)**, Zhang et al., 2013 **(51)** |

| *Node* | *#* | *Aptamer Sequence Reported* |
|---|---|---|
| *1* | 6 | AGCAGCACAGAGGTCAGATGGCAGCTAAGCAGGCGGCTCACAAAACCATTCGCATGCGGCCCTATGCGTGCTACCGTGAA |
| *2A* | 1 | AGCAGCACAGAGGTCAGATGGCAG**<u>G</u>**TAAGCAGGCGGCTCACAAAACCATTCGCATGCGGCCCTATGCGTGCTACCGTGAA |
| *2B* | 11 | ~~AGCAGCACAGAGGTCAGATG~~GCAGCTAAGCAGGCGGCTCACAAAACCATTCGCATGCGGC~~CCTATGCGTGCTACCGTGAA~~ |
| *2C* | 1 | **CGAACAACGCTAAAATTC**~~AGCAGCACAGAGGTCAGATG~~GCAGCTAAGCAGGCGGCTCACAAAACCATTCGCATGCGGCCCTATGCGTGCTACCGTGAA**ATAGGAGACGGGCAACAGCTGATCCTGATGG** |
| *0* | 2 | No sequence Provided |

**S7 | Phylogeny depicting unexplained aptamer sequence alterations introduced to the 80mer DNA lysozyme binding aptamer (Apta1, Tran et al., 2010).** Unexplained insertions are bolded, unexplained substitutions are bolded and underlined, unexplained deletions are struck out and justified or explained alterations are in light grey. The number (#) column indicates the number of publications found reporting each sequence in our analysis.

**A.** Tran et al. selected a new ssDNA **(1)** aptamer in 2010 that showed higher binding affinity than the aptamer previously described by Cox et al. Since then it has been used in a variety of aptasensors. In 2011 Zou et al. **(2A)** used the Tran aptamer publishing the sequence with the primers included, indicating that the first and last 22 nt were a part of the primer, with a single nt mutation with no apparent or articulated purpose. Several subsequent publications, Wang et al. 2011, **(2B)** publish the sequence without the primer without citing a minimized variant of the aptamer. Finally, Wang et al. 2008 use the Tran aptamer with novel primers **(2C).** Additionally, 3 publications presumably use the aptamer developed by Tran et al. but do not provide the sequence. This is especially problematic in cases in which several aptamers exist for the same target. I suspect that if we were to review papers that cite the articles that include unintentional errors, we would find a greater percentage of their usage in the literature. **B.** The distribution of the type of error causing unintended changes to sequence information are provided showing that deviations in sequence information is the most common type of error, with deletions causing 50% of error in the 21 publications reviewed. **C.** Illustrates the number of publications using each sequence in the 21 reviewed.

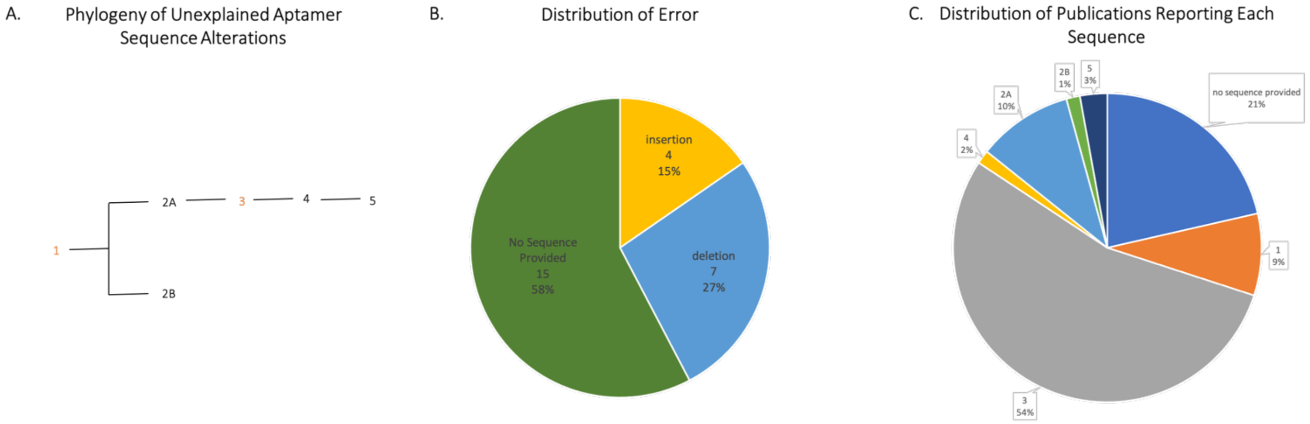


| ID | Publications Reporting Sequence |
|---|---|
| 1 | **Bates et al., 1999 (6)**, Reyes-Reyes et al., 2019 **(66)** |
| 2A | Hong et al., 2018 **(28),** Hwang et al., 2010 **(30),** Ko et al., 2009 **(35)**, Kang et al., **(34)**, Li et al., 2014 **(44)**, Luo et al., 2017 **(49),** Ozalp et al., 2011 **(59)** |
| 2B | Chauhan et al., 2018 **(14)** |
| 3 | Alibolandi et al., 2014 **(2),** Aravind et al., 2012 **(3),** Balinsky et al., 2013 **(4)**, Bates et al., 2010 **(5),** Bates et al., 2017 **(7),** Cao et al., 2009 **(10)**, Carvalho et al., 2019 **(11)**, Charoenphol et al., 2014 **(13),** Cheng et al., 2016 **(15),** Cho et al., 2016 **(16)**, Choi et al., 2009 **(17)**, Dapic et al., 2002 **(20),** Farzin et al., 2018 **(23)**, Feng et al., 2011 **(24)**, Feng et al., 2015 **(25)**, Girvan et al., 2006 **(26)**, Hu et al., 2015 **(29)**, Leaderer et al., 2015 **(38)**, Li et al., 2014 **(42)**, Li et al., 2012 **(43),** Lian et al., 2012 **(45)**, Malik et al., 2015 **(52)**, Maremanda et al., 2015 **(53)**, Metifiot et al., 2015 **(54),** Noaparast et al., 2015 **(57)**, Oh et al., 2014 **(58),** Park et al., 2018 **(60)**, Park et al., 2012 **(61)**, Reyes-Reyes et al., 2010 **(65)**, Reyes-Reyes et al., 2015 **(64)**, Shiao et al., 2014 **(69)**, Shieh et al., 2010 **(71)**, Soundararajan et al., 2008 **(72)**, Soundararajan et al., 2009 **(73),** Taghavia et al., 2017 **(74),** Teng et al., 2007 **(75)**, Trinh et al., 2015 **(77),** Wu et al., 2013 **(81).** Xing et al., 2013 **(82)** |
| 4 | Zhou et al., 2013 **(89)** |
| 5 | Ai et al., 2014, **(1),** Motaghi et al., 2017 **(55)** |
| 0 | Farin et al., 2011 **(22)**, Goldshmit et al., 2014 **(27)**, Ishimaru et al., 2010 **(32)**, Lai et al., 2010 **(37)**, Liao et al., 2015 **(46)**, Litchfield et al., 2012 **(47)**, Lorents et al., 2018 **(48),** Maier and Levy, 2016 **(51),** Perrone et al., 2016 **(63),** Schokoroy et al., 2013 **(67)**, Sharma et al., 2018 **(68), Shieh et al., 2009 (70),** Tosoni et al., 2015 **(76),** Wolfson et al., 2016 **(79)**, Wolfson et al., 2018 **(80)** |

| *Node* | # | *Aptamer Sequence Reported* |
|---|---|---|
| *1* | 6 | TTTGGTGGTGGTGGTTGTGGTGGTGGTGG |
| *2A* | 7 | ~~T~~TTGGTGGTGGTGGTTGTGGTGGTGGTGG |
| *2B* | 1 | TTTTTTGGTGGTGGTGGTTGTGGTGGTGGTGG**TTT** |
| *3* | 38 | ~~TTT~~GGTGGTGGTGGTTGTGGTGGTGGTGG |
| *4* | 1 | ~~TTT~~GGTGGTGGTGGTTGTGGTGGTGGTGG**T** |
| *5* | 2 | ~~TTT~~GGTGGTGGTGGTTGTGGTGGTGGTGG**TTT** |
| *0* | 15 | No sequence provided |

**S8 | Phylogeny depicting unexplained aptamer sequence alterations introduced to the 26mer DNA nucleolin binding aptamer (AS1411/AGRO100, Bates et al., 1999)**. Unexplained insertions are bolded, unexplained substitutions are bolded and underlined, unexplained deletions are struck out and justified or explained alterations are in light grey. The number (#) column indicates the number of publications found reporting each sequence in our analysis.

The phylogeny constructed traces alterations in the AS1411 ssDNA aptamer sequence since its discovery in 1999 by Bates et al. by screening anti-proliferative activity among antisense oligonucleotides rather than the SELEX method typical in aptamer development. One notable early modification made to the GRO29A sequence **(1)** removed the 5' 3T cap and 3'aminoalkyl because they were found to be unnecessary for serum stability and protection from nuclease activity (Girven et al.) resulting in the AS1411/AGRO100 sequence **(3).** A non-intentional mutation occurs **(2)** in which Ko et al. include 2 T's at the 5' end of AS1411. Further, Chauhan et al. **(4A)** report a sequence with explained 5'6T spacer, but do not describe the 3Ts added to the 3' end. Similarly, Zhou et al. **(4B)** state that they use a 26mer but provide a 27nt sequence with an added and unexplained T at the 3' end. There were multiple missteps found in publications within the field that could lead to subsequent unintentional mutations. Lee et al. used GRO29A sequence but called it AS1411, including the TTT. While this does not affect the functionality of the anti-nucleolin aptamer as it was effective in both forms (Bates et al) this is an inconsistency in the naming and sequence that could prove deleterious in other circumstances. Further, 12 of the 50 publications reviewed do not report the aptamer sequence, making it impossible to know from their publication if they used the correct sequence of AS1411 or an unintentionally modified sequence that found its way into the literature. Finally, the final concentrations of aptamer and conditions used varied across the publications. This inconsistency, particularly on dosage, is problematic particularly in the case of an aptamer with therapeutic applications. Lee et al. 2009 used GRO29A sequence but called it AS1411, including the TTT. While this would not have an effect on the functionality of the anti-nucleolin aptamer as it was effective in both forms (Bates et al) this is an inconsistency in the naming and sequence that could prove deleterious in other circumstances.

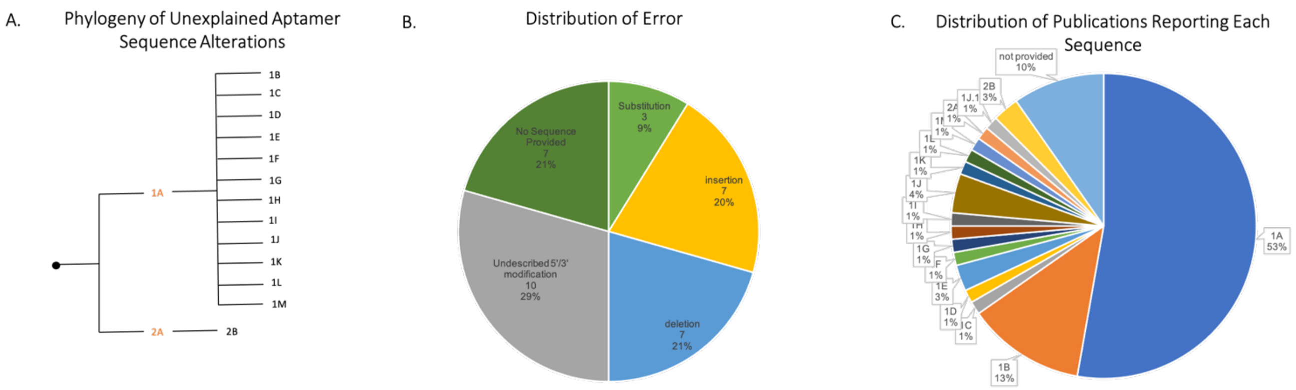


| ID | Publications Reporting Sequence |
|---|---|
| 1A | Adhikari et al., 2015 (1), Buchanan et al., 2003 (6), Cheow & Han, 2011 (7), Cho et al., 2006 (8), Cole et al., 2007 (9), Ding et al., (11), Feng et al., 2008 (13), Feng et al., 2009 (12), German et al., 1998 (17), Gokulrangan et al., 2005 (18), Gong et al., 2007 (19), He et al., 2009 (20), Hu and Easley, 2011 (21), Jiang et al., 2004 (23), Jiang et al., 2003 (24), Katilius et al., 2007 (26), Lautnera et al., N.D. (29), Lee et al., 2013 (30), Liss et al., 2002 (34), Luo et al., 2011 (36), Park et al., 2013 (44), Peng et al., 2011 (46), Pollet et al., 2008 (47), Poongavanam et al., 2016 (49), Tran et al., 2011 (56), Turgeon et al., 2010 (57), Unruh et al., 2005a (58), Unruh et al., 2005b (59), Wang et al., 2008 (62), Wang et al., 2007 (64), **Wiegand et al., 1996** (65), Wu et al., 2009 (65), Xia et al., 2014 (67), Xu et al., 2005 (69), Xu et al., 2006 (70), Yao et al., 2009 (71), Yao et al., 2010 (72), YiXian et al., 2013 (73), Zhang & Yadavalli, 2011 (75) |
| 1B | Kim et al., 2009 (28)*, Lee et al., 2008 (31)*, Maehashi et al., 2007 (37), Ohno et al., 2011 (42)*, Papamichael et al., 2006 (43), Šnejdárková et al., 2008 (53), Stadtherr et al., 2005 (55), Wang et al., 2011 (61) |
| 1C | Lin et al., 2006 (33)* |
| 1D | Wang et el., 2010 (63)* |
| 1E | Fukasawa et al., 2009 (16)*, Yoshida et al., 2008 (74)* |
| 1F | Xiao et al., 2009 (68)* |
| 1G | Liu et al., 2015 (35)* |
| 1H | Jiang et al., 2015 (22)* |
| 1I | Pollet et al., 2012 (48)* |
| 1J | Li et al., 2011 (32)*, Nam et al., 2012 (40)*, Sim et al., 2010 (52)* |
| 1J.1 | Kim et al., 2010 (27)* |
| IK | Bai & Zhao, 2017 (2)* |
| 1L | Schachermeyer et al., 2013 (51)* |
| 1M | Spiridonova et al., 2014 (54)* |
| 2A | Wiegand et al., 1996 (64) |
| 2B | Fischer et al., 2008 (15)*, Fischer & Tarasow, 2011 (14)* |
| 0 | Balmurugan et al., 2008 (4), Bai et al., 2017 (3), Collet et al., 2005 (10), Maehasi et al., 2009 (38), Oh et al., 2014 (41), Pei & Stojanovic, 2008 (45), Vinkenborg et al., 2012 (60) |

| *Node* | *#* | *Aptamer Sequence Reported* |
|---|---|---|
| *1A* | 38 | GGGGCACGTTTATCCGTCCCTCCTAGTGGCGTGCCCC |
| *1B* | 9 | **GCGC**GGGGCACGTTTATCCGTCCCTCCTAGTGGCGTGCCCC**GCGC** |
| *1C* | 1 | GGGGCACGTTTATCCGTCCCT~~CCT~~AGTGGCGTGCCCC |
| *1D* | 1 | **ATCTAC**GGGGCACGTTTATCCGTCCCTCCTAGTGGCGTGCCCC |
| *1E* | 2 | ~~GGGG~~**T**CACGTTTATCCGTCCCTCCTAGTGGCGTG**A**~~CCCC~~ |
| *1F* | 1 | **TTTTT**GGGGCACGTTTATCCGTCCCTCCTAGTGGCGTGCCCC |
| *1G* | 1 | **TTTTTTTTTT**GGGGCACGTTTATCCGTCCCTCCTAGTGGCGTGCCCC |
| *1H* | 1 | **CTAATTGT**GGGGCACGTTTATCCGTCCCTCCTAGTGGCGTGCCCC**TTTTTTTTTTTTTTTTTTAGGAGG** |
| *1I* | 1 | **AAAAA**GGGGCACGTTTATCCGTCCCTCCTAGTGGCGTGCCCC |
| *1J* | 3 | **AAAAAAAAAAAAAAA**GGGGCACGTTTATCCGTCCCTCCTAGTGGCGTGCCCC |
| *1J.1* | 1 | **AAAAAAAAAAAAAAA**GGGGCACGTTTATCCGTCCCTCCTAGTGGCGTGCCC**G** |
| *1K* | 1 | ~~GGG~~GCACGTTTATCCGTCCCTCCTAGTGGCGTGC~~CCC~~ |
| *1L* | 1 | GGGGCACGTTTATCCGTCCCTCCTAGTGGCGTGCCCC**A** |
| *1M* | 1 | GGGGCACGTTTATCCGTCCCT~~CCT~~AGTGGCGTGCCCC**iT** |
| *2A* | 1 | TTTATCCGTTCCTCTTAGTGG |
| *2B* | 2 | TTTATCCGTTCCTC**C**TAGTGG |
| *0* | 7 | No sequence provided |

**S9 | Phylogeny depicting unexplained aptamer sequence alterations introduced to the two DNA Immunoglobulin E (IgE) binding aptamers: D17.4 a 37mer (node 1A) and "consensus sequence" a 21mer (2A) Wiegand et al., 1996**. Unexplained insertions are bolded, unexplained substitutions are bolded and underlined, unexplained deletions are struck out and justified or explained alterations are in light grey. The number (#) column indicates the number of publications found reporting each sequence in our analysis.

The IgE binding aptamer, clone D17.4, was discovered by Wiegand et al. (1996) **(ID 1A)**. Stjovanic et al., also described a DNA IgE consensus sequence was discovered as well **(2A)**. In the Wiegand et al. (1996) publication, a secondary structure was proposed with four added G-C pairs, two to each 5' and 3' end.to promote extra stability of the sequence. This addition of G-C pair nucleotides is present in the Liss et al. (2002) publication in which the aptamer is now referred to as D17.4ext, and this publication clearly reports that the 8 nucleotide expansion was added to promote the stability of the secondary structure **(1A)**. Subsequent publications present in node 1B also reported the same sequence with the G-C pair addition described by Liss et al., 2002; however, two of the publications failed to report the explicit purpose behind this change **(1B).**

In both the Lin et al. (2006)and Spiridonova et al. (2014) publications, 3 nucleotides C-C-T were deleted from the middle of the sequence, but the purpose of this change was not explained nor was a secondary structure prediction made in either publications **(1C)**.

The Wang et al. (2010) publication reported the IgE Aptamer sequence with reference to the Wiegand et al. (1996) publication; however, the nucleotide set ATCTAC was added to the 5' end with no explanation as to this purpose or prediction of how the secondary structure would change as a result of this mutation **(1D).**

The Fukasawa et al. (2009) and Yoshida et al. (2008) publications had both additions and deletions to the 5' and 3' ends of the original sequence **(1E).** The first four GGGG nucleotides were deleted from the 5' end and the last four CCCC nucleotides were deleted from the 3' end, and T nucleotide was added to the beginning of the 5' end and an A nucleotide was added to the end of the 3' end. Interestingly, this aptamer sequence was reported as a 31-mer aptamer, which took note of deletions and insertion changes to the sequence. However, the IgE aptamer cited the Wiegand et al. (1996) publication when reporting the sequence, but did not explain the purpose behind these mutations.

Addition of a polyT tail to the 5' end of the sequence is present in the publications present in node 1F and 1G of varying lengths. Specifically, in the Xiao et al. (2009) publication, the PolyT addition is 5 nucleotides in length **(1F),** and in the Liu et al., 2015 publication this PolyT addition is 10 nucleotides in length **(1G).** In both publications, however, the purpose behind these PolyT 5' modifications is not explained.

The Jiang et al. (2015) publication had large sets of nucleotide additions to both the 5' and 3' ends **(1H).** However, this pseudoknot IgE binding aptamer was designed using Mfold program and the secondary structure of this aptamer is clearly shown. The purposes of this modifications from the original sequence were not clearly explained.

Addition of a polyA tail to the 5' end of the sequence is present in the publications present in nodes 1I and 1J, and once again, these PolyA tails are of varying lengths. In the Pollet et al., 2012 publication, the PolyA tail is 5 nucleotides in length **(1I),** but in the Li et al., 2011 publication, the PolyA addition is 15 nucleotides in length **(1J).** Once again, in both publications, the purpose behind these PolyA 5' modifications were not explained.

In the Bai and Zhao (2017) publication, a three nucleotide deletion was made to both the 5' and 3' end **(1K)**. Specifically, the first three nucleotides of the sequence, GGG, were deleted from the 5' end and the last three nucleotides of the sequence, CCC, were deleted from the 3' end. The purpose behind these deletions were not explained. Interestingly, although 6 nucleotides were deleted from the sequence, the aptamer was still referred to as a 37-mer sequence even though the length was transformed to a 31-mer sequence.

In the Schachermeyer et al. (2013) publication, a single nucleotide A was added to the very end of the sequence at the 3' end **(1L)**. No secondary structure was predicted for the altered sequence, and the purpose behind this lone base modification was not explained.

In the final node, the Xia et al., 2014, although the secondary structure of the DNA aptamer was listed, there was a nucleotide deletion, of 9 bases in length, from the 5' end of the sequence and another nucleotide deletion, of 8 bases in length, from the 3' end of the sequence **(1M).** In addition, a polyT tail was added and modified both the 5' and 3' ends of the sequence **(1M)**. The purposes behind these deletions and modifications were not explained in the publication.

When tracking the changes of the secondary sequence presented in node 2A, in both the Fischer et al. (2008) and Fischer and Tarasow (2011) publications, a substitution was made that converted the thymine base into a cytosine base. The purpose behind this substitution was not explained **(2B)**. Also, it is important to note that both of these publications were from the same group, which explains the continuity of the same substitution error present in both publications.

Additionally, the Luo et al. (2011), Maehashi et al. (2009), Basnar et al. (2006), Collett et al. (2005), Vinkenborg et al. (2012), Oh et al. (2014), Ruff et al. (2010), Jing and Browser (2011), and Pier and Stojanvoic (2008) publications did not report the sequence of the IgE aptamer although they did reference the original Wiegand et al. (1996) publication when referring to the IgE Aptamer. Therefore, these publications were excluded from the analysis.

Mendonsa et al. (2004) did not report the IgE aptamer sequence, but instead did report the primers from the original selection and the subsequent new sequences obtained from their original selection. This was not cited as an error as this reporting style was unique and as they did not report the original sequence from Wiegand et al. (1996), there was not way to digress whether an unexplained modification was made.

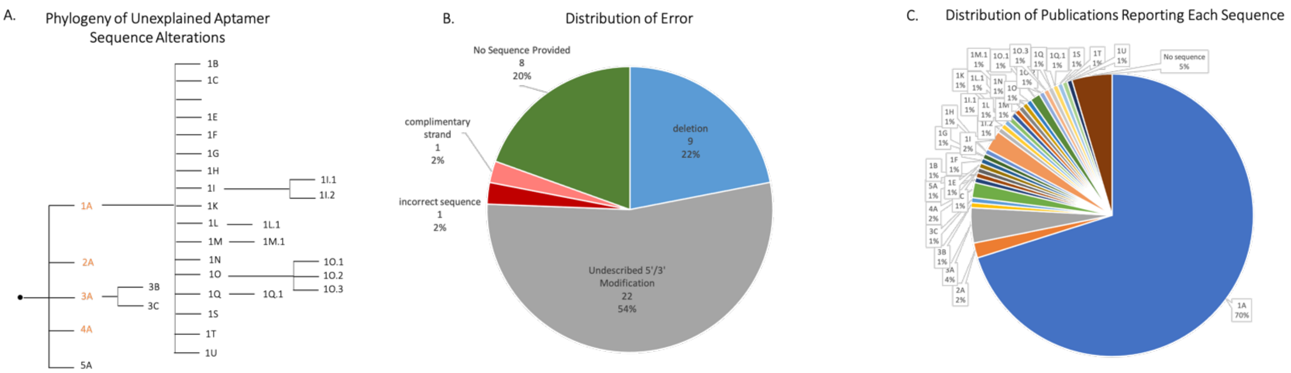


| ID | Publications Reporting Sequence |
|---|---|
| 1A | Ali et al., 2014 (1), Baggiani et al., 2015 (2), Bianco et al., 2017 (5), Brothier et al., 2014 (7), Castillo et al., 2012 (9), Chapuis-Hugon et al., 2011 (10), Chen et al., 2014 (12), Chen et al., 2014 (15), Chen et al., 2019 (14), Chen et al., 2012 (11), Chen et al., 2018 (13), Chrouda et al., 2015 (16), Costantini et al., 2016 (17), **Cruz-Aguado et al., 2008** (18), Dai et al., 2016 (19), Dai et al., 2017 (20), De Girolamo et al., 2011 (22), Dou et al., 2016 (25), Duan et al., 2011 (26), Evtugyn et al., 2013 (28), Evtugyn et al., 2017 (29), Galaretta et al., 2012 (30), Geng et al., 2013 (32), Guo et al., 2011 (34), Hao et al., 2016 (35), Hayat et al., 2013 (36), Hayat et al., 2013 (37), Hayat et al., 2015 (38), Hayat et al., 2013 (39), Huang et al., 2018 (41), Hun et al., 2013 (43), Jiang et al., 2014 (45), Karimi et al., 2017 (49), Kidd et al., 2011 (50), Kuang et a., 2010 (51), Lee et al., 2014 (53), Li et al., 2018 (54), Li et al., 2019 (55), Li et al., 2019 (56), Liu et al., 2016 (62), Liu et al., 2018 (57), Liu et al., 2019 (60), Liu et al., 2015 (61), Liu et al., 2015 (59), Liu et al., 2018 (58), Loo et al., 2015 (63), Lv et al., 2014 (71), Lv et al., 2018 (67), Lv et al., 2016 (66), Lv et al., 2017 (68), Lv et al., 2017 (69), Lv et al., 2017 (70), Ma et al., 2018 (72), Marechal et al., 2015 (74), Mazaafrianto et al., 2018 (75), McKeague et al., 2014 (77), McKeague et al., 2015 (76), Mejri-Omrani et al., 2015 (78), Mishra et al., 2015 (79), Nekrasov et al., 2019 (84), Ni et al., 2018 (85), Perrier et al., 2014 (88), Phanchai et al., 2018 (89), Prabhakar et al., 2011 (90), Prieto-Simón et al., 2014 (91), Qian et al., 2015 (93), Qian et al., 2018 (92), Qing et al., 2017 (94), Rhouati et al., 2013 (95), Samokhvalov et al., 2018 (98), Samokhvalov et al., 2017 (97), Samokhvalov et al., 2018 (99), Sanzani et al., 2014 (100), Schax et al., 2015 (101), Shao et al., 2018 (102), Sharma et al., 2015 (103), Sharma et al., 2016 (104), Shen et al., 2018 (105), Sheng et al., 2011 (107), Simão et al., 2017 (108), Somerson et al., 2018 (109), Song et al., 2018 (111), Sui et al., 2018 (112), Sun et al., 2017 (113), Tan et al., 2015 (114), Tian et al., 2019 (115), Tian et al., 2018 (116), Tong et al., 2012 (118), Tong et al., 2011 (117), Wang et al., 2017 (120), Wang et al., 2011 (126), Wang et al., 2016 (123), Wang et al., 2019 (125), Wang et al., 2010 (130), Wang et al., 2015 (121), Wang et al., 2015 (122), Wei et al., 2015 (132), Wu et al., 2019 (134), Wu et al., 2018 (135), Wu et al., 2012 (133), Wu et al., 2017 (137), Wu et al., 2018 (138), Xiao et al., 2018 (140), Xu et al., 2017 (142), Xu et al., 2016 (144), Yang et al., 2017 (153), Yang et al., 2013 (145), Yang et al., 2011 (147), Yang et al., 2015 (148), Yang et al., 2014 (151), Yang et al., 2014 (152), Yin et al., 2018 (155), Yin et al., 2017 (156), Yu et al., 2018 (157), Yuan et al., 2014 (158), Yue et al., 2014 (159), Zhang et al., 2017 (163), Zhang et al., 2015 (164), Zhang et al., 2013 (162), Zhao et al., 2019 (165), Zhao et al., 2014 (167), Zhao et al., 2013 (166), Zhu et al., 2015 (170) |
| 1B | Yao et al., 2015 (154), Huang et al., 2013 (42) |
| 1C | Barthelmebs et al., 2011 (3) |
| 1F | Song et al., 2018 (110) |
| 1H | Jin et al., 2018 (46) |
| 1I | Jo et al., 2018 (47), Mun et al., 2014 (82), Muthamizh et al., 2017 (83), Park et al., 2014 (87) |
| 1I.1 | Jo et al., 2016 (48) |
| 1I.2 | Jo et al., 2016 (48) |
| 1K | Lee et al., 2018 (52) |
| 1L | Lu et al., 2017 (64) |
| 1L.1 | Luan et al., 2015 (65) |
| 1M | Rivas et al., 2015 (96) |
| 1M.1 | Wang et al., 2011 (127) |
| 1N | Shen et al., 2017 (106) |
| 1O | Yang et al., 2014 (149) |
| 1O.1 | Gu et al., 2016 (33), Wang et al., 2015 (128) |
| 1O.2 | Tian et al., 2019 (115), He et al., 2019 (40) |
| 1O.3 | Zhang et al., 2018 (160) |
| 1Q | Wang et al., 2015 (119) |
| 1Q.1 | Wang et al., 2018 (124) |
| 1S | Wei et al., 2019 (131) |
| 1T | Yang et al., 2011 (146) |
| 1U | Zhang et al., 2012 (161) |
| 2A | Cruz-Aguado et al., 2008 (18), Barthelmebs et al., 2010 (4), Modh et al., 2017 (80), Niazi et al., 2019 (86) |
| 3A | Cruz-Aguado et al., 2008 (18); Duan et al., 2011 (26); Wu et al., 2012 (136), Wu et al., 2011 (137) Ma et al., 2013 (73) |
| 3B | Motyčka et al., 2014 (81) |
| 3C | Zhao et al., 2015 (168) |
| 4A | Deng et al., 2018 (23), Deore et al., 2019 (24), Zhu et al., 2019 (173) |
| 5A | Xu et al., 2019 (141) |
| 0 | Bonel et al., 2010 (6), Cai et al., 2018 (8), De Girolamo et al., 2012 (21), Ganbold et al., 2014 (31), Ji et al., 2019 (44), Rhouati et al., 2013 (95), Xu et al., 2016 (143), Yang et al., 2014 (150) |

| *Node* | *#* | *Aptamer Sequence Reported* |
|---|---|---|
| *1A* | 122 | GATCGGGTGTGGGTGGCGTAAAGGGAGCATCGGACA |
| *1B* | 1 | GATCGGGTGTGGGTGGCGTAAAGGGAGCATCGGACA**GGCGAAGACAGGTGCTTAGT** |
| *1C* | 1 | GGAGGACGAAGCGGAACCGGGTGTGGGTGCCTTGATCCAGGGAGTCTCAGAAGACACGCCCGACA |
| *1F* | 1 | TGTCCGATGCTCCCTTTACGCCACCCACACCCGATC |
| *1H* | 1 | **GCTGAGTCTGAGTC**GATCGGGTGTGGGTGGCGTAAAGGGAGCATCGGACA |
| *1I* | 4 | GCATCTGATCGGGTGTGGGTGGCGTAAAGG |
| *1I.1* | 1 | AAGGGTAGGGCGGGTTGGGTAAATAAAAAGCATCTGATCGGGTGTGGGTGGCGTAAAGG**AAA** |
| *1I.2* | 1 | AAGGGTAGGGCGGGTTGGGTAAATGCATCTGATCGGGTGTGGG~~T~~GGCGTAAAGG**AAA** |
| *1K* | 1 | **TTTTT**GATCGGGTGTGGGTGGCGTAAAGGGAGCATCGGACA |
| *1L* | 1 | CCTGGGAGGGAGGGAGGGATCGGGTGTGGGTGGCGTAAAGGGAGCATCGGACACCCGATCCC |
| *1L.1* | 1 | CTGGGAGGGAGGGAGGGATCGGGTGTGGGTGGCGTAAAGGGAGCATCGGACACCCGATCCC |
| *1M* | 1 | GATCGGGTGTGGGTGGCGTAAAGGGAGCATCGGACA**AAAAAAAAA** |
| *1M.1* | 1 | GATCGGGTGTGGGTGGCGTAAAGGGAGCATCGGACA**AAAAAAAAAAAAAAAAAA** |
| *1N* | 1 | **TGGGTAGGGCGGGTTGGGAAA**GATCGGGTGTGGGTGGCGTAAAGGGAGCATCGGACA |
| *1O* | 1 | **AAA**GATCGGGTGTGGGTGGCGTAAAGGGAGCATCGGACA |
| *1O.1* | 2 | **AAAAAA**GATCGGGTGTGGGTGGCGTAAAGGGAGCATCGGACA |
| *1I.2* | 2 | **AAAAAAAAAAAA**GATCGGGTGTGGGTGGCGTAAAGGGAGCATCGGACA |
| *1O.3* | 1 | **AAAAAAAAAAAAAAAAAA**GATCGGGTGTGGGTGGCGTAAAGGGAGCATCGGACA |
| *1Q* | 1 | GATCGGGTGTGGGTGGCGTAAAGGGAGCATCGGACA**CGCCACCCACAC** |
| *1Q.1* | 1 | GATCGGGTGTGGGTGGCGTAAAGGGAGCATCGGACA**CGCCACCCACACA** |
| *1S* | 1 | **TGAG**GATCGGGTGTGGGTGGCGTAAAGGGAGCATCGGACA |

| | | |
|---|---|---|
| *1T* | 1 | CTGGGAGGGAGGGAGG~~G~~ATCGGGTGTGGGTGGCGTAAAGGGAGCATCGGACACCCGATCCC |
| *1U* | 1 | TTCGATCGGGTGTGGGTGGCGTAAAGGGAGCATCGGA~~CA~~AAAA |
| *2A* | 3 | TGGTGGCTGTAGGTCAGCATCTGATCGGGTGTGGGTGGCGTAAAGGGAGCATCGGACAACG |
| *3A* | 7 | GATCGGGTGTGGGTGGCGTAAAGGGAGCATCGG |
| *3B* | 1 | GATCGGGTGTGGGTGGCGTAAAGG~~G~~AGCATCGG |
| *3C* | 1 | TTTTTTTTTTGATCGGGTGTGGGTGGCGTAAAGGGAGCATCGG |
| *4A* | 3 | GATCGGGTGTGGGTGGCGTAAAGGGAGCATC |
| *5A* | 1 | GGGGTGAAACGGGTCCCG |
| *0* | 8 | No sequence provided |

**S10 | Phylogeny depicting unexplained aptamer sequence alterations introduced to the three DNA Ochratoxin A (OTA) binding aptamers: 1.12 a 61mer (node 2A), 1.12.5 a 36mer (1A), and 1.12.8 33mer (3A), Cruz-Aguado et al., 2008.** Unexplained insertions are bolded, unexplained substitutions are bolded and underlined, unexplained deletions are struck out and justified or explained alterations are in light grey. The number (#) column indicates the number of publications found reporting each sequence in our analysis.

Orange IDs are the clones described by Cruz-Aguado (2008), 1A was 1.12.5 (36mer), 2A was 1.12 (61mer), 3A was 1.12.8 (33mer), 4A was 1.12.12 (31mer), and 5A was the random region of 1.13 without the primers. **A.** Cruz-Aguado et al. selected several novel ssDNA **(1A, 2A, 3A)** aptamers against OTA in 2008. Since then, it has also been used in a variety of aptasensors. **(1U)** A deletion of the last two nucleotides from the variable region was found in a publication (Zhang et al. 2012), an unexplained modification from the any previously published minimized aptamer sequences. **(1C)** Barthelmebs et al., 2011 cite the Cruz-Aguado aptamer but provide the sequence of their own aptamer described in 2010. In addition, Wang et al., 2014 included an unexplained 5′ $A_6$ addition **(1O.1).** Jo et al. (2016) use the 1.12 sequence from Table 2 in the Cruz-Aguado paper without including the primer regions which are stated in the text to be excluded from the sequences in the table. In addition, in the same table Jo et al provide a sequence with a deletion of a thymine nucleotide from the variable region in addition to an undescribed 3′ poly A addition found in both **(1I.1 and 1I.2 respectively)**. Park et al., 2014 included 5′ and 3′ poly A additions without describing these modifications in addition to using the 1.12 sequence from Table 2 in the Cruz-Aguado paper without including the primer regions which are stated in the text to be excluded from the sequences in the table **(1I).** Luan et al. (2015 **1L.1**) and Shen et al. (2017, **1N)** both include undescribed 5′/3′ additions, and Yang et al. (2011, **1T**), provide the correct sequence and fully described their additions, but in their figure they include the 5′ G in the added region rather than the aptamer region. **B.** The distribution of the type of error causing unintended changes to sequence information are provided, showing that deviations in sequence information are the most common type of error, notwithstanding publications that did not report a sequence. Deletions caused 33% of error in the 50 publications reviewed. **C.** Illustrates the number of publications using each sequence in the 50 publications reviewed. ID's depicted with orange text were described variants that were fully characterized.

### Works Cited

The red text are references that were excluded from the analysis (i.e., because they did not use the sequence experimentally, were reviews, etc.).

***Figure 1 80mer RNA anti- lysozyme aptamer (clone 1, Cox et al., 2001) References***


1. Baldrich, E., Restrepo, A., & O'Sullivan, C. K. (2004). Aptasensor development: elucidation of critical parameters for optimal aptamer performance. *Analytical chemistry*, *76*(23), 7053-7063.
2. Bamrungsap, S., Shukoor, M. I., Chen, T., Sefah, K., & Tan, W. (2011). Detection of Lysozyme Magnetic Relaxation Switches Based on. *Analytical Chemistry*, *83*(20), 7795–7799. https://doi.org/10.1021/ac201442a
3. Bayramoglu, G., Ozalp, V. C., Yilmaz, M., Guler, U., Salih, B., & Arica, M. Y. (2015). Lysozyme specific aptamer immobilized MCM-41 silicate for single-step purification and quartz crystal microbalance (QCM)-based determination of lysozyme from chicken egg white. *Microporous and Mesoporous Materials*, *207*, 95–104. https://doi.org/10.1016/j.micromeso.2015.01.009
4. Bi, W., Bai, X., Gao, F., Lu, C., Wang, Y., Zhai, G., … Zhang, K. (2017). DNA-Templated Aptamer Probe for Identification of Target Proteins. *Analytical Chemistry*, *89*(7), 4071–4076. https://doi.org/10.1021/acs.analchem.6b04895
5. Cao, X., Xia, J., Liu, H., Zhang, F., Wang, Z., & Lu, L. (2017). A new dual-signalling electrochemical aptasensor with the integration of "signal on/off" and "labeling/label-free" strategies. *Sensors and Actuators, B: Chemical*, *239*, 166–171. https://doi.org/10.1016/j.snb.2016.08.009
6. Cheglakov, Z., Weizmann, Y., Braunschweig, A. B., Wilner, O. I., & Willner, I. (2008). Increasing the complexity of periodic protein nanostructures by the rolling-circle-amplified synthesis of aptamers. *Angewandte Chemie - International Edition*, *47*(1), 126–130. https://doi.org/10.1002/anie.200703688
7. Chen, F., Gülbakan, B., & Zenobi, R. (2013). Direct access to aptamer–protein complexes via MALDI-MS. *Chemical Science*, *4*(10), 4071–4078. https://doi.org/10.1039/c3sc51410b
8. Chen, Z., Guo, J., Li, J., & Guo, L. (2013). Tetrahexahedral Au nanocrystals/aptamer based ultrasensitive electrochemical biosensor. *RSC Advances*, *3*(34), 14385–14389. https://doi.org/10.1039/c3ra41065j
9. Cheng, A. K. H., Ge, B., & Yu, H. Z. (2007). Aptamer-based biosensors for label-free voltammetric detection of lysozyme. *Analytical Chemistry*, *79*(14), 5158–5164. https://doi.org/10.1021/ac062214q
10. Cheng, A. K., Su, H., Wang, Y. A., & Yu, H. Z. (2009). Aptamer-based detection of epithelial tumor marker mucin 1 with quantum dot-based fluorescence readout. *Analytical chemistry*, *81*(15), 6130-6139.
11. Cho, E. J., Collett, J. R., Szafranska, A. E., & Ellington, A. D. (2006). Optimization of aptamer microarray technology for multiple protein targets. *Analytica Chimica Acta*, *564*(1), 82–90. https://doi.org/10.1016/j.aca.2005.12.038
12. Collett, J. R., Cho, E. J., Lee, J. F., Levy, M., Hood, A. J., Wan, C., & Ellington, A. D. (2005). Functional RNA microarrays for high-throughput screening of antiprotein aptamers. *Analytical Biochemistry*, *338*(1), 113–123. https://doi.org/10.1016/j.ab.2004.11.027
13. Collett, J. R., Eun, J. C., & Ellington, A. D. (2005). Production and processing of aptamer microarrays. *Methods*, *37*(1), 4–15. https://doi.org/10.1016/j.ymeth.2005.05.009

14. Cox, J. C., & Ellington, A. D. (2001). Automated Selection of Anti-Protein Aptamers. *Bioorganic and Medicinal Chemistry*, *9*, 2525–2531. https://doi.org/10.1016/s0968-0896(01)00028-1
15. Cox, J. C., Hayhurst, A., Hesselberth, J., Bayer, T. S., Georgiou, G., & Ellington, A. D. (2002). Automated selection of aptamers against protein targets translated in vitro: from gene to aptamer. *Nucleic Acids Research*, *30*(20), e108-e108.
16. Deore, P. S., & Manderville, R. A. (2019). Aptamer-induced thermofluorimetric protein stabilization and G-quadruplex nucleic acid staining by SYPRO orange dye. *New Journal of Chemistry*, *43*(13), 4994–4997. https://doi.org/10.1039/C9NJ00188C
17. Fang, S., Dong, X., Ji, H., Liu, S., Yan, F., Peng, D., … Zhang, Z. (2016). Electrochemical aptasensor for lysozyme based on a gold electrode modified with a nanocomposite consisting of reduced graphene oxide, cuprous oxide, and plasma-polymerized propargylamine. *Microchimica Acta*, *183*(2), 633–642. https://doi.org/10.1007/s00604-015-1675-5
18. Ghosh, S., Khan, N. I., Tsavalas, J. G., & Song, E. (2018). Selective Detection of Lysozyme Biomarker Utilizing Large Area Chemical Vapor Deposition-Grown Graphene-Based Field-Effect Transistor. *Frontiers in Bioengineering and Biotechnology*, *6*(March), 1–7. https://doi.org/10.3389/fbioe.2018.00029
19. Girardot, M., Li, H. Y., Descroix, S., & Varenne, A. (2011). Determination of binding parameters between lysozyme and its aptamer by frontal analysis continuous microchip electrophoresis (FACMCE). *Journal of Chromatography A*, *1218*(26), 4052–4058. https://doi.org/10.1016/j.chroma.2011.04.077
20. Giuffrida, M. C., Cigliana, G., & Spoto, G. (2018). Ultrasensitive detection of lysozyme in droplet-based microfluidic devices. *Biosensors and Bioelectronics*, *104*, 8–14. https://doi.org/10.1016/j.bios.2017.12.042
21. Gokulrangan, G., Unruh, J. R., Holub, D. F., Ingram, B., Johnson, C. K., & Wilson, G. S. (2005). DNA aptamer-based bioanalysis of IgE by fluorescence anisotropy. *Analytical chemistry*, *77*(7), 1963-1970.
22. Hayashi, E., Takada, T., Nakamura, M., & Yamana, K. (2010). Electronic aptamer-based biosensor for multiprotein analytes on a single paltform. *Chemistry Letters*, *39*(5), 454–455. https://doi.org/10.1246/cl.2010.454
23. Huang, H., Jie, G., Cui, R., & Zhu, J. J. (2009). DNA aptamer-based detection of lysozyme by an electrochemiluminescence assay coupled to quantum dots. *Electrochemistry Communications*, *11*(4), 816–818. https://doi.org/10.1016/j.elecom.2009.01.009
24. Huang, H., Zhang, Q., Luo, J., & Zhao, Y. (2012). Sensitive colorimetric detection of lysozyme in human serum using peptide-capped gold nanoparticles. *Analytical Methods*, *4*(11), 3874-3878.
25. Huang, J., Zhu, Z., Bamrungsap, S., Zhu, G., You, M., He, X., ... & Tan, W. (2010). Competition-mediated pyrene-switching aptasensor: probing lysozyme in human serum with a monomer-excimer fluorescence switch. *Analytical chemistry*, *82*(24), 10158-10163.
26. Hybarger, G., Bynum, J., Williams, R. F., Valdes, J. J., & Chambers, J. P. (2006). A microfluidic SELEX prototype. *Analytical and Bioanalytical Chemistry*, *384*(1), 191–198. https://doi.org/10.1007/s00216-005-0089-3
27. Kawde, A. N., Rodriguez, M. C., Lee, T. M. H., & Wang, J. (2005). Label-free bioelectronic detection of aptamer-protein interactions. *Electrochemistry Communications*, *7*(5), 537–540. https://doi.org/10.1016/j.elecom.2005.03.008
28. Khan, N. I., Maddaus, A. G., & Song, E. (2018). A low-cost inkjet-printed aptamer-based electrochemical biosensor for the selective detection of lysozyme. *Biosensors*, *8*(1). https://doi.org/10.3390/bios8010007
29. Khedri, M., Ramezani, M., Rafatpanah, H., & Abnous, K. (2018). Detection of food-born allergens with aptamer-based biosensors. *TrAC Trends in Analytical Chemistry*, *103*, 126-136.
30. Kirby, R., Cho, E. J., Gehrke, B., Bayer, T., Park, Y. S., Neikirk, D. P., … Ellington, A. D. (2004). Aptamer-based sensor arrays for the detection and quantitation of proteins. *Analytical Chemistry*, *76*(14), 4066–4075. https://doi.org/10.1021/ac049858n
31. Lee, J. F., Cox, J. C., Collett, J. R., & Ellington, A. D. (2005). Exploring sequence space through automated aptamer selection. *JALA: Journal of the Association for Laboratory Automation*, *10*(4), 213-218.
32. Li, C. M., Zhan, L., Zheng, L. L., Li, Y. F., & Huang, C. Z. (2016). A magnetic nanoparticle-based aptasensor for selective and sensitive determination of lysozyme with strongly scattering silver nanoparticles. *Analyst*, *141*(10), 3020–3026. https://doi.org/10.1039/c6an00489j
33. Li, D., Shlyahovsky, B., Elbaz, J., & Willner, I. (2007). Amplified analysis of low-molecular-weight substrates or proteins by the self-assembly of DNAzyme-aptamer conjugates. *Journal of the American Chemical Society*, *129*(18), 5804–5805. https://doi.org/10.1021/ja070180d
34. Li, L. D., Chen, Z. B., Zhao, H. T., Guo, L., & Mu, X. (2010). An aptamer-based biosensor for the detection of lysozyme with gold nanoparticles amplification. *Sensors and Actuators, B: Chemical*, *149*(1), 110–115. https://doi.org/10.1016/j.snb.2010.06.015
35. Li, S., Gao, Z., & Shao, N. (2014). Non-covalent conjugation of CdTe QDs with lysozyme binding DNA for fluorescent sensing of lysozyme in complex biological sample. *Talanta*, *129*, 86–92. https://doi.org/10.1016/j.talanta.2014.04.062
36. Li, Y., Qi, H., Gao, Q., & Zhang, C. (2011). Label-free and sensitive electrogenerated chemiluminescence aptasensor for the determination of lysozyme. *Biosensors and Bioelectronics*, *26*(5), 2733–2736. https://doi.org/10.1016/j.bios.2010.09.048
37. Liao, D., Chen, J., Li, W., Zhang, Q., Wang, F., Li, Y., & Yu, C. (2013). Fluorescence turn-on detection of a protein using cytochrome c as a quencher. *Chemical Communications*, *49*(82), 9458–9460. https://doi.org/10.1039/c3cc43985b
38. Lin, X., Ivanov, A. P., & Edel, J. B. (2017). Selective single molecule nanopore sensing of proteins using DNA aptamer-functionalised gold nanoparticles. *Chemical Science*, *8*(5), 3905–3912. https://doi.org/10.1039/c7sc00415j
39. Liu, S., Na, W., Pang, S., Shi, F., & Su, X. (2014). A label-free fluorescence detection strategy for lysozyme assay using CuInS2quantum dots. *Analyst*, *139*(12), 3048–3054. https://doi.org/10.1039/c4an00160e
40. Liu, X., Li, X., Lu, Y., Cao, J., & Li, F. (2018). A split aptamer-based imaging solution for the visualization of latent fingerprints. *Analytical Methods*, *10*(19), 2281–2286. https://doi.org/10.1039/c8ay00538a
41. Liu, Y., Lin, C., Li, H., & Yan, H. (2005). Aptamer-directed self-assembly of protein arrays on a DNA nanostructure. *Angewandte Chemie International Edition*, *44*(28), 4333-4338.
42. Lu, Y., Liu, Y., Zhang, S., Wang, S., & Zhang, X. (2013). Aptamer-based plasmonic sensor array for discrimination of proteins and cells with the naked eye. *Analytical Chemistry*, *85*(14), 6571–6574. https://doi.org/10.1021/ac4014594.
43. Lv, Z., Liu, J., Bai, W., Yang, S., & Chen, A. (2015). A simple and sensitive label-free fluorescent approach for protein detection based on a Perylene probe and aptamer. *Biosensors and Bioelectronics*, *64*, 530-534.
44. Mihai, I., Vezeanu, A., Polonschii, C., Albu, C., Radu, G. L., & Vasilescu, A. (2015). Label-free detection of lysozyme in wines using an aptamer based biosensor and SPR detection. *Sensors and Actuators, B: Chemical*, *206*, 198–204. https://doi.org/10.1016/j.snb.2014.09.050
45. Mihai, I., Vezeanu, A., Polonschii, C., David, S., Gáspár, S., Bucur, B., … Vasilescu, A. (2014). Low-fouling SPR detection of lysozyme and its aggregates. *Analytical Methods*, *6*(19), 7646–7654. https://doi.org/10.1039/c4ay01237b
46. Ocaña, C., Hayat, A., Mishra, R. K., Vasilescu, A., del Valle, M., & Marty, J. L. (2015). Label free aptasensor for Lysozyme detection: A comparison of the analytical performance of two aptamers. *Bioelectrochemistry*, *105*(October), 72–77. https://doi.org/10.1016/j.bioelechem.2015.05.009
47. Ostatná, V., Kasalová-Vargová, V., Kékedy-Nagy, L., Černocká, H., & Ferapontova, E. E. (2017). Chronopotentiometric sensing of specific interactions between lysozyme and the DNA aptamer. *Bioelectrochemistry*, *114*, 42–47. https://doi.org/10.1016/j.bioelechem.2016.12.003
48. Padlan, C. S., Malashkevich, V. N., Almo, S. C., Levy, M., Brenowitz, M., & Girvin, M. E. (2014). An RNA aptamer possessing a novel monovalent cation-mediated fold inhibits lysozyme catalysis by inhibiting the binding of long natural substrates. *Rna*, *20*(4), 447–461. https://doi.org/10.1261/rna.043034.113
49. Peng, Y., Zhang, D., Li, Y., Qi, H., Gao, Q., & Zhang, C. (2009). Label-free and sensitive faradic impedance aptasensor for the determination of lysozyme based on target-induced aptamer

displacement. *Biosensors and Bioelectronics*, *25*(1), 94–99. https://doi.org/10.1016/j.bios.2009.06.001
50. Potty, A. S. R., Kourentzi, K., Fang, H., Schuck, P., & Willson, R. C. (2011). Biophysical characterization of DNA and RNA aptamer interactions with hen egg lysozyme. *International Journal of Biological Macromolecules*, *48*(3), 392–397. https://doi.org/10.1016/j.ijbiomac.2010.12.007
51. Robertson, M. P., & Ellington, A. D. (2001). In vitro selection of nucleoprotein enzymes. *nature biotechnology*, *19*(7), 650-655.
52. Rodríguez, M. C., & Rivas, G. A. (2009). Label-free electrochemical aptasensor for the detection of lysozyme. *Talanta*, *78*(1), 212–216. https://doi.org/10.1016/j.talanta.2008.11.002
53. Shamsipur, M., Farzin, L., & Tabrizi, M. A. (2016). Ultrasensitive aptamer-based on-off assay for lysozyme using a glassy carbon electrode modified with gold nanoparticles and electrochemically reduced graphene oxide. *Microchimica Acta*, *183*(10), 2733–2743. https://doi.org/10.1007/s00604-016-1920-6
54. Song, Y., Xu, C., Wei, W., Ren, J., & Qu, X. (2011). Light regulation of peroxidase activity by spiropyran functionalized carbon nanotubes used for label-free colorimetric detection of lysozyme. *Chemical Communications*, *47*(32), 9083–9085. https://doi.org/10.1039/c1cc13279b
55. Stovall, G. M., Cox, J. C., & Ellington, A. D. (2004). Automated optimization of aptamer selection buffer conditions. *JALA: Journal of the Association for Laboratory Automation*, *9*(3), 117-122.
56. Subramanian, P., Lesniewski, A., Kaminska, I., Vlandas, A., Vasilescu, A., Niedziolka-Jonsson, J., … Szunerits, S. (2013). Lysozyme detection on aptamer functionalized graphene-coated SPR interfaces. *Biosensors and Bioelectronics*, *50*, 239–243. https://doi.org/10.1016/j.bios.2013.06.026
57. Teller, C., Shimron, S., & Willner, I. (2009). Aptamer-DNAzyme hairpins for amplified biosensing. *Analytical Chemistry*, *81*(21), 9114–9119. https://doi.org/10.1021/ac901773b
58. Titoiu, A. M., Porumb, R., Fanjul-Bolado, P., Epure, P., Zamfir, M., & Vasilescu, A. (2019). Detection of allergenic lysozyme during winemaking with an electrochemical aptasensor. *Electroanalysis*, *31*(11), 2262-2273.
59. Tran, D. T., Janssen, K. P., Pollet, J., Lammertyn, E., Anné, J., Van Schepdael, A., & Lammertyn, J. (2010). Selection and characterization of DNA aptamers for egg white lysozyme. *Molecules*, *15*(3), 1127-1140.
60. Truong, P. L., Choi, S. P., & Sim, S. J. (2013). Amplification of resonant rayleigh light scattering response using immunogold colloids for detection of lysozyme. *Small*, *9*(20), 3485–3492. https://doi.org/10.1002/smll.201202638
61. Vasilescu, A., Gaspar, S., Mihai, I., Tache, A., & Litescu, S. C. (2013). Development of a label-free aptasensor for monitoring the self-association of lysozyme. *Analyst*, *138*(12), 3530–3537. https://doi.org/10.1039/c3an00229b
62. Vasilescu, A., Purcarea, C., Popa, E., Zamfir, M., Mihai, I., Litescu, S., … Marty, J. L. (2016). Versatile SPR aptasensor for detection of lysozyme dimer in oligomeric and aggregated mixtures. *Biosensors and Bioelectronics*, *83*, 353–360. https://doi.org/10.1016/j.bios.2016.04.080
63. Vasilescu, A., Wang, Q., Li, M., Boukherroub, R., & Szunerits, S. (2016). Aptamer-based electrochemical sensing of lysozyme. *Chemosensors*, *4*(2), 10.
64. Wang, B., & Yu, C. (2010). Fluorescence Turn-On Detection of a Protein through the Reduced Aggregation of a Perylene Probe. *Angewandte Chemie*, *122*(8), 1527–1530. https://doi.org/10.1002/ange.200905237
65. Wang, J., Wei, T., Li, X., Zhang, B., Wang, J., Huang, C., & Yuan, Q. (2014). Near-infrared-light-mediated imaging of latent fingerprints based on molecular recognition. *Angewandte Chemie - International Edition*, *53*(6), 1616–1620. https://doi.org/10.1002/anie.201308843
66. Wang, Q., Subramanian, P., Schechter, A., Teblum, E., Yemini, R., Nessim, G. D., … Szunerits, S. (2016). Vertically Aligned Nitrogen-Doped Carbon Nanotube Carpet Electrodes: Highly Sensitive Interfaces for the Analysis of Serum from Patients with Inflammatory Bowel Disease. *ACS Applied Materials and Interfaces*, *8*(15), 9600–9609. https://doi.org/10.1021/acsami.6b00663
67. Wang, X., Xu, Y., Chen, Y., Li, L., Liu, F., & Li, N. (2011). The gold-nanoparticle-based surface plasmon resonance light scattering and visual DNA aptasensor for lysozyme. *Analytical and Bioanalytical Chemistry*, *400*(7), 2085–2091. https://doi.org/10.1007/s00216-011-4943-1
68. Wang, Y., Pu, K. Y., & Liu, B. (2010). Anionic conjugated polymer with aptamer-functionalized silica nanoparticle for label-free naked-eye detection of lysozyme in protein mixtures. *Langmuir*, *26*(12), 10025–10030. https://doi.org/10.1021/la100139p
69. Wood, M., Maynard, P., Spindler, X., Lennard, C., & Roux, C. (2012). Visualization of latent fingermarks using an aptamer-based reagent. *Angewandte Chemie - International Edition*, *51*(49), 12272–12274. https://doi.org/10.1002/anie.201207394
70. Xia, J., Song, D., Wang, Z., Zhang, F., Yang, M., Gui, R., … Xia, L. (2015). Single electrode biosensor for simultaneous determination of interferon gamma and lysozyme. *Biosensors and Bioelectronics*, *68*, 55–61. https://doi.org/10.1016/j.bios.2014.12.045
71. Xia, Y., Gan, S., Xu, Q., Qiu, X., Gao, P., & Huang, S. (2013). A three-way junction aptasensor for lysozyme detection. *Biosensors and Bioelectronics*, *39*(1), 250–254. https://doi.org/10.1016/j.bios.2012.07.053
72. Yeung, M. C. L., Wong, K. M. C., Tsang, Y. K. T., & Yam, V. W. W. (2010). Aptamer-induced self-assembly of a NIR-emissive platinum(ii) terpyridyl complex for label- and immobilization-free detection of lysozyme and thrombin. *Chemical Communications*, *46*(41), 7709–7711. https://doi.org/10.1039/c0cc02631j
73. Yüce, M., Ullah, N., & Budak, H. (2015). Trends in aptamer selection methods and applications. *Analyst*, *140*(16), 5379-5399.
74. Zhang, F., Zhao, Y. Y., Chen, H., Wang, X. H., Chen, Q., & He, P. G. (2015). Sensitive fluorescence detection of lysozyme using a tris(bipyridine)ruthenium(ii) complex containing multiple cyclodextrins. *Chemical Communications*, *51*(30), 6613–6616. https://doi.org/10.1039/c5cc00428d
75. Zhang, H., Fang, C., & Zhang, S. (2011). Ultrasensitive electrochemical analysis of two analytes by using an autonomous DNA machine that works in a two-cycle mode. *Chemistry - A European Journal*, *17*(27), 7531–7537. https://doi.org/10.1002/chem.201002767
76. Zhao, Y., Chen, H., Chen, Q., Qi, Y., Yang, F., Tang, J., … Zhang, F. (2014). Ultrasensitive and signal-on electrochemiluminescence aptasensor using the multi-tris(bipyridine)ruthenium(II)-β-cyclodextrin complexes. *Chinese Journal of Chemistry*, *32*(11), 1161–1168. https://doi.org/10.1002/cjoc.201400511
77. Zhu, H., Ding, Y., Wang, A., Sun, X., Wu, X. C., & Zhu, J. J. (2015). A simple strategy based on upconversion nanoparticles for a fluorescent resonant energy transfer biosensor. *Journal of Materials Chemistry B*, *3*(3), 458–464. https://doi.org/10.1039/c4tb01320d
78. Zou, M., Chen, Y., Xu, X., Huang, H., Liu, F., & Li, N. (2012). The homogeneous fluorescence anisotropic sensing of salivary lysozyme using the 6-carboxyfluorescein-labeled DNA aptamer. *Biosensors and Bioelectronics*, *32*(1), 148-154.
79. Zuo, L., Qin, G., Lan, Y., Wei, Y., & Dong, C. (2019). A turn-on phosphorescence aptasensor for ultrasensitive detection of lysozyme in humoral samples. *Sensors and Actuators, B: Chemical*, *289*(December 2018), 100–105. https://doi.org/10.1016/j.snb.2019.03.08

***S1: 15mer DNA anti-thrombin aptamer (TBA15, Bock et al., 1992) References***

1. Aviñó, A., Fabrega, C., Tintoré, M., & Eritja, R. (2012). Thrombin Binding Aptamer, more than a simple aptamer: Chemically modified derivatives and biomedical applications. *Current Pharmaceutical Design*, *18*(14), 2036–2047.
2. Bai, Y., Feng, F., Zhao, L., Wang, C., Wang, H., Tian, M., … He, X. (2013). Aptamer/thrombin/aptamer-AuNPs sandwich enhanced surface plasmon resonance sensor for the detection of

subnanomolar thrombin. *Biosensors and Bioelectronics*, *47*, 265–270. https://doi.org/10.1016/j.bios.2013.02.004
3. Bérn Abérem, M., Najari, A., Ho, A. A., Gravel, J. F., Nobert, P., Boudreau, D., & Leclerc, M. (2006). Protein detecting arrays based on cationic polythiophene-DNA-aptamer complexes. *Advanced Materials*, *18*(20), 2703–2707. https://doi.org/10.1002/adma.200601651
4. Bing, T., Liu, X., Cheng, X., Cao, Z., & Shangguan, D. (2010). Bifunctional combined aptamer for simultaneous separation and detection of thrombin. *Biosensors and Bioelectronics*, *25*(6), 1487–1492. https://doi.org/10.1016/j.bios.2009.11.003
5. Bock, L. C., Griffin, L. C., Latham, J. A., Vermaas, E. H., & Toole, J. J. (1992). Selection of single-stranded DNA molecules that bind and inhibit human thrombin. *Nature*, *355*(6), 564–566. https://doi.org/10.1038/255242a0
6. Bompiani, K. M., Monroe, D. M., Church, F. C., & Sullenger, B. A. (2012). A high affinity, antidote-controllable prothrombin and thrombin-binding RNA aptamer inhibits thrombin generation and thrombin activity. *Journal of Thrombosis and Haemostasis*, *10*(5), 870–880. https://doi.org/10.1111/j.1538-7836.2012.04679.x
7. Boncler, M. A., Koziolkiewicz, M., & Watala, C. (2001). Aptamer inhibits degradation of platelet proteolytically activatable receptor, PAR-1, by thrombin. *Thrombosis Research*, *104*(3), 215–222. https://doi.org/10.1016/S0049-3848(01)00357-7
8. Cai, H., Lee, T. M. H., & Hsing, I. M. (2006). Label-free protein recognition using an aptamer-based impedance measurement assay. *Sensors and Actuators, B: Chemical*, *114*(1), 433–437. https://doi.org/10.1016/j.snb.2005.06.017
9. Centi, S., Messina, G., Tombelli, S., Palchetti, I., & Mascini, M. (2008). Different approaches for the detection of thrombin by an electrochemical aptamer-based assay coupled to magnetic beads. *Biosensors and Bioelectronics*, *23*(11), 1602–1609. https://doi.org/10.1016/j.bios.2008.01.020
10. Chen, J., Zhang, J., Li, J., Yang, H. H., Fu, F., & Chen, G. (2010). An ultrasensitive signal-on electrochemical aptasensor via target-induced conjunction of split aptamer fragments. *Biosensors and Bioelectronics*, *25*(5), 996–1000. https://doi.org/10.1016/j.bios.2009.09.015
11. Chen, Q., Tang, W., Wang, D., Wu, X., Li, N., & Liu, F. (2010). Amplified QCM-D biosensor for protein based on aptamer-functionalized gold nanoparticles. *Biosensors and Bioelectronics*, *26*(2), 575–579. https://doi.org/10.1016/j.bios.2010.07.034
12. Cho, H., Baker, B. R., Wachsmann-Hogiu, S., Pagba, C. V., Laurence, T. A., Lane, S. M., … Tok, J. B. H. (2008). Aptamer-based SERRS sensor for thrombin detection. *Nano Letters*, *8*(12), 4386–4390. https://doi.org/10.1021/nl802245w
13. Connor, A. C., & McGown, L. B. (2006). Aptamer stationary phase for protein capture in affinity capillary chromatography. *Journal of Chromatography A*, *1111*(2), 115–119. https://doi.org/10.1016/j.chroma.2005.05.012
14. Deng, B., Lin, Y., Wang, C., Li, F., Wang, Z., Zhang, H., … Le, X. C. (2014). Aptamer binding assays for proteins: The thrombin example-A review. *Analytica Chimica Acta*, *837*, 1–15. https://doi.org/10.1016/j.aca.2014.04.055
15. Dougan, H., Weitz, J. I., Stafford, A. R., Gillespie, K. D., Klement, P., Hobbs, J. B., & Lyster, D. M. (2003). Evaluation of DNA aptamers directed to thrombin as potential thrombus imaging agents. *Nuclear Medicine and Biology*, *30*(1), 61–72. https://doi.org/10.1016/S0969-8051(02)00378-5
16. Evtugyn, G., Porfireva, A., Ivanov, A., Konovalov, O., & Hianik, T. (2009). Molecularly imprinted polymerized methylene green as a platform for electrochemical sensing of aptamer- thrombin interactions. *Electroanalysis*, *21*(11), 1272–1277. https://doi.org/10.1002/elan.200804556
17. Fialová, M., Kypr, J., & Vorlíčková, M. (2006). The thrombin binding aptamer GGTTGGTGTGGTTGG forms a bimolecular guanine tetraplex. *Biochemical and Biophysical Research Communications*, *344*(1), 50–54. https://doi.org/10.1016/j.bbrc.2006.03.144
18. Gosai, A., Ma, X., Balasubramanian, G., & Shrotriya, P. (2016). Electrical Stimulus Controlled Binding/Unbinding of Human Thrombin-Aptamer Complex. *Scientific Reports*, *6*(November), 1–12. https://doi.org/10.1038/srep37449
19. Gronewold, T. M. A., Glass, S., Quandt, E., & Famulok, M. (2005). Monitoring complex formation in the blood-coagulation cascade using aptamer-coated SAW sensors. *Biosensors and Bioelectronics*, *20*(10 SPEC. ISS.), 2044–2052. https://doi.org/10.1016/j.bios.2004.09.007
20. Hamaguchi, N., Ellington, A., & Stanton, M. (2001). Aptamer beacons for the direct detection of proteins. *Analytical Biochemistry*, *294*(2), 126–131. https://doi.org/10.1006/abio.2001.5169
21. Hasegawa, H., Taira, K. I., Sode, K., & Ikebukuro, K. (2008). Improvement of aptamer affinity by dimerization. *Sensors*, *8*(2), 1090–1098. https://doi.org/10.3390/s8021090
22. Holland, C. A., Henry, A. T., Whinna, H. C., & Church, F. C. (2000). Effect of oligodeoxynucleotide thrombin aptamer on thrombin inhibition by heparin cofactor II and antithrombin. *FEBS Letters*, *484*(2), 87–91. https://doi.org/10.1016/S0014-5793(00)02131-1
23. Hoon, S., Zhou, B., Janda, K. D., Brenner, S., & Scolnick, J. (2011). Aptamer selection by high-throughput sequencing and informatic analysis. *BioTechniques*, *51*(6), 413–416. https://doi.org/10.2144/000113786
24. Huang, D. W., Niu, C. G., Qin, P. Z., Ruan, M., & Zeng, G. M. (2010). Time-resolved fluorescence aptamer-based sandwich assay for thrombin detection. *Talanta*, *83*(1), 185–189. https://doi.org/10.1016/j.talanta.2010.09.004
25. Huang, H., & Zhu, J. J. (2009). DNA aptamer-based QDs electrochemiluminescence biosensor for the detection of thrombin. *Biosensors and Bioelectronics*, *25*(4), 927–930. https://doi.org/10.1016/j.bios.2009.08.008
26. Ikebukuro, K., Kiyohara, C., & Sode, K. (2004). Electrochemical detection of protein using a double aptamer sandwich. *Analytical Letters*, *37*(14), 2901–2909. https://doi.org/10.1081/AL-200035778
27. Ikebukuro, K., Yoshida, W., Noma, T., & Sode, K. (2006). Analysis of the evolution of the thrombin-inhibiting DNA aptamers using a genetic algorithm. *Biotechnology Letters*, *28*(23), 1933–1937. https://doi.org/10.1007/s10529-006-9174-8
28. Jiang, Z., Yang, T., Liu, M., Hu, Y., & Wang, J. (2014). An aptamer-based biosensor for sensitive thrombin detection with phthalocyanine@SiO2 mesoporous nanoparticles. *Biosensors and Bioelectronics*, *53*, 340–345. https://doi.org/10.1016/j.bios.2013.10.005
29. Jung, Y. K., Kim, T. W., Park, H. G., & Soh, H. T. (2010). Specific colorimetric detection of proteins using bidentate aptamer-conjugated polydiacetylene (PDA) liposomes. *Advanced Functional Materials*, *20*(18), 3092–3097. https://doi.org/10.1002/adfm.201001008
30. Kang, Y., Feng, K. J., Chen, J. W., Jiang, J. H., Shen, G. L., & Yu, R. Q. (2008). Electrochemical detection of thrombin by sandwich approach using antibody and aptamer. *Bioelectrochemistry*, *73*(1), 76–81. https://doi.org/10.1016/j.bioelechem.2008.04.024
31. Krauss, I. R., Merlino, A., Giancola, C., Randazzo, A., Mazzarella, L., & Sica, F. (2011). Thrombin-aptamer recognition: A revealed ambiguity. *Nucleic Acids Research*, *39*(17), 7858–7867. https://doi.org/10.1093/nar/gkr522
32. Kretz, C. A., Stafford, A. R., Fredenburgh, J. C., & Weitz, J. I. (2006). HD1, a thrombin-directed aptamer, binds exosite 1 on prothrombin with high affinity and inhibits its activation by prothrombinase. *Journal of Biological Chemistry*, *281*(49), 37477–37485. https://doi.org/10.1074/jbc.M607359200
33. Kupakuwana, G. V., Crill, J. E., McPike, M. P., & Borer, P. N. (2011). Acyclic identification of aptamers for human alpha-thrombin using over-represented libraries and deep sequencing. *PLoS ONE*, *6*(5). https://doi.org/10.1371/journal.pone.0019395

34. Latham, J. A., Johnson, R., & Toole, J. J. (1994). The application of a modified nucleotide in aptamer selection: novel thrombin aptamers containing-(1-pentynyl)-2′-deoxyuridine. *Nucleic acids research*, *22*(14), 2817-2822.
35. Lee, J. F., Stovall, G. M., & Ellington, A. D. (2006). Aptamer therapeutics advance. *Current Opinion in Chemical Biology*, *10*(3), 282–289. https://doi.org/10.1016/j.cbpa.2006.03.015
36. Li, J. J., Fang, X., & Tan, W. (2002). Molecular aptamer beacons for real-time protein recognition. *Biochemical and Biophysical Research Communications*, *292*(1), 31–40. https://doi.org/10.1006/bbrc.2002.6581
37. Li, L., Zhao, H., Chen, Z., Mu, X., & Guo, L. (2010). Aptamer-based electrochemical approach to the detection of thrombin by modification of gold nanoparticles. *Analytical and Bioanalytical Chemistry*, *398*(1), 563–570. https://doi.org/10.1007/s00216-010-3922-2
38. Lin, P. H., Chen, R. H., Lee, C. H., Chang, Y., Chen, C. S., & Chen, W. Y. (2011). Studies of the binding mechanism between aptamers and thrombin by circular dichroism, surface plasmon resonance and isothermal titration calorimetry. *Colloids and Surfaces B: Biointerfaces*, *88*(2), 552–558. https://doi.org/10.1016/j.colsurfb.2011.07.032
39. Long, S. B., Long, M. B., White, R. R., & Sullenger, B. A. (2008). Crystal structure of an RNA aptamer bound to thrombin. *Rna*, *14*(12), 2504–2512. https://doi.org/10.1261/rna.1239308
40. Martino, L., Virno, A., Randazzo, A., Virgilio, A., Esposito, V., Giancola, C., … Mayol, L. (2006). A new modified thrombin binding aptamer containing a 5′-5′ inversion of polarity site. *Nucleic Acids Research*, *34*(22), 6653–6662. https://doi.org/10.1093/nar/gkl915
41. Mazurov, A. V., Titaeva, E. V., Khaspekova, S. G., Storojilova, A. N., Spiridonova, V. A., Kopylov, A. M., & Dobrovolsky, A. B. (2011). Characteristics of a new DNA aptamer, direct inhibitor of thrombin. *Bulletin of Experimental Biology and Medicine*, *150*(4), 422–425. https://doi.org/10.1007/s10517-011-1158-6
42. McCauley, T. G., Hamaguchi, N., & Stanton, M. (2003). Aptamer-based biosensor arrays for detection and quantification of biological macromolecules. *Analytical Biochemistry*, *319*(2), 244–250. https://doi.org/10.1016/S0003-2697(03)00297-5
43. Mendelboum Raviv, S., Horváth, A., Aradi, J., Bagoly, Z., Fazakas, F., Batta, Z., … Hársfalvi, J. (2008). 4-Thio-deoxyuridylate-modified thrombin aptamer and its inhibitory effect on fibrin clot formation, platelet aggregation and thrombus growth on subendothelial matrix. *Journal of Thrombosis and Haemostasis*, *6*(10), 1764–1771. https://doi.org/10.1111/j.1538-7836.2008.03106.x
44. Musumeci, D., & Montesarchio, D. (2012). Polyvalent nucleic acid aptamers and modulation of their activity: A focus on the thrombin binding aptamer. *Pharmacology and Therapeutics*, *136*(2), 202–215. https://doi.org/10.1016/j.pharmthera.2012.07.011
45. Nagatoishi, S., Isono, N., Tsumoto, K., & Sugimoto, N. (2011). Loop residues of thrombin-binding DNA aptamer impact G-quadruplex stability and thrombin binding. *Biochimie*, *93*(8), 1231–1238. https://doi.org/10.1016/j.biochi.2011.03.013
46. Nagatoishi, S., Tanaka, Y., & Tsumoto, K. (2007). Circular dichroism spectra demonstrate formation of the thrombin-binding DNA aptamer G-quadruplex under stabilizing-cation-deficient conditions. *Biochemical and Biophysical Research Communications*, *352*(3), 812–817. https://doi.org/10.1016/j.bbrc.2006.11.088
47. Najafi-Shoushtari, S. H., & Famulok, M. (2007). DNA aptamer-mediated regulation of the hairpin ribozyme by human α-thrombin. *Blood Cells, Molecules, and Diseases*, *38*(1), 19–24. https://doi.org/10.1016/j.bcmd.2006.10.007
48. Nimjee, S. M., Oney, S., Volovyk, Z., Bompiani, K. M., Long, S. B., Hoffman, M., & Sullenger, B. A. (2009). Synergistic effect of aptamers that inhibit exosites 1 and 2 on thrombin. *Rna*, *15*(12), 2105–2111. https://doi.org/10.1261/rna.1240109
49. Padmanabhan, K., Padmanabhan, K. P., Ferrara, J. D., Sadler, J. E., & Tulinsky, A. (1993). The structure of α-thrombin inhibited by a 15-mer single-stranded DNA aptamer. *Journal of Biological Chemistry*, *268*(24), 17651–17654. https://doi.org/10.2210/pdb1hut/pdb
50. Pagano, B., Martino, L., Randazzo, A., & Giancola, C. (2008). Stability and binding properties of a modified thrombin binding aptamer. *Biophysical Journal*, *94*(2), 562–569. https://doi.org/10.1529/biophysj.107.117382
51. Pagba, C. V., Lane, S. M., Cho, H., & Wachsmann-Hogiu, S. (2010). Direct detection of aptamer-thrombin binding via surface-enhanced Raman spectroscopy. *Journal of Biomedical Optics*, *15*(4), 047006. https://doi.org/10.1117/1.3465594
52. Park, M. K., Kee, J. S., Quah, J. Y., Netto, V., Song, J., Fang, Q., … Lo, G. Q. (2013). Label-free aptamer sensor based on silicon microring resonators. *Sensors and Actuators, B: Chemical*, *176*, 552–559. https://doi.org/10.1016/j.snb.2012.08.078
53. Pasternak, A., Hernandez, F. J., Rasmussen, L. M., Vester, B., & Wengel, J. (2011). Improved thrombin binding aptamer by incorporation of a single unlocked nucleic acid monomer. *Nucleic Acids Research*, *39*(3), 1155–1164. https://doi.org/10.1093/nar/gkq823
54. Pica, A., Russo Krauss, I., Merlino, A., Nagatoishi, S., Sugimoto, N., & Sica, F. (2013). Dissecting the contribution of thrombin exosite i in the recognition of thrombin binding aptamer. *FEBS Journal*, *280*(24), 6581–6588. https://doi.org/10.1111/febs.12561
55. Rahman, M. A., Jung, I. S., Won, M. S., & Shim, Y. B. (2009). Gold nanoparticles doped conducting polymer nanorod electrodes: Ferrocene catalyzed aptamer-based thrombin immunosensor. *Analytical Chemistry*, *81*(16), 6604–6611. https://doi.org/10.1021/ac900285v
56. Rinker, S., Ke, Y., Liu, Y., Chhabra, R., & Yan, H. (2008). Self-assembled DNA nanostructures for distance-dependent multivalent ligand-protein binding. *Nature Nanotechnology*, *3*(7), 418–422. https://doi.org/10.1038/nnano.2008.164
57. Russo Krauss, I., Merlino, A., Randazzo, A., Novellino, E., Mazzarella, L., & Sica, F. (2012). High-resolution structures of two complexes between thrombin and thrombin-binding aptamer shed light on the role of cations in the aptamer inhibitory activity. *Nucleic Acids Research*, *40*(16), 8119–8128. https://doi.org/10.1093/nar/gks512
58. Rye, P. D., & Nustad, K. (2001). Immunomagnetic DNA aptamer assay. *BioTechniques*, *30*(2), 290–295. https://doi.org/10.2144/01302st01
59. Schlensog, M. D., Gronewold, T. M. A., Tewes, M., Famulok, M., & Quandt, E. (2004). A Love-wave biosensor using nucleic acids as ligands. *Sensors and Actuators, B: Chemical*, *101*(3), 308–315. https://doi.org/10.1016/j.snb.2004.03.015
60. Song, M., Zhang, Y., Li, T., Wang, Z., Yin, J., & Wang, H. (2009). Highly sensitive detection of human thrombin in serum by affinity capillary electrophoresis/laser-induced fluorescence polarization using aptamers as probes. *Journal of Chromatography A*, *1216*(5), 873–878. https://doi.org/10.1016/j.chroma.2008.11.085
61. Sosic, A., Meneghello, A., Cretaio, E., & Gatto, B. (2011). Human thrombin detection through a sandwich aptamer microarray: Interaction analysis in solution and in solid phase. *Sensors*, *11*(10), 9426–9441. https://doi.org/10.3390/s111009426
62. Strehlitz, B., Nikolaus, N., & Stoltenburg, R. (2008). Protein detection with aptamer biosensors. *Sensors*, *8*(7), 4296-4307.
63. Tasset, D. M., Kubik, M. F., & Steiner, W. (1997). Oligonucleotide inhibitors of human thrombin that bind distinct epitopes. *Journal of Molecular Biology*, *272*(5), 688–698. https://doi.org/10.1006/jmbi.1997.1275
64. Tennico, Y. H., Hutanu, D., Koesdjojo, M. T., Bartel, C. M., & Remcho, V. T. (2010). On-chip aptamer-based sandwich assay for thrombin detection employing magnetic beads and quantum dots. *Analytical Chemistry*, *82*(13), 5591–5597. https://doi.org/10.1021/ac101269u

65. Wang, K. Y., Bolton, P. H., Krawczyk, S. H., Bischofberger, N., & Swaminathan, S. (1993). The Tertiary Structure of a DNA Aptamer Which Binds to and Inhibits Thrombin Determines Activity. *Biochemistry*, *32*(42), 11285–11292. https://doi.org/10.1021/bi00093a004
66. Wang, Q., Zhou, Z., Zhai, Y., Zhang, L., Hong, W., Zhang, Z., & Dong, S. (2015). Label-free aptamer biosensor for thrombin detection based on functionalized graphene nanocomposites. *Talanta*, *141*, 247–252. https://doi.org/10.1016/j.talanta.2015.04.012
67. Wang, W., Chen, C., Qian, M., & Zhao, X. S. (2008). Aptamer biosensor for protein detection using gold nanoparticles. *Analytical Biochemistry*, *373*(2), 213–219. https://doi.org/10.1016/j.ab.2007.11.013
68. Wang, X., & Wang, X. (2013). Aptamer-functionalized hydrogel diffraction gratings for the human thrombin detection. *Chemical Communications*, *49*(53), 5957–5959. https://doi.org/10.1039/c3cc41827h
69. Wang, Y. H., Xia, H., Huang, K. J., Wu, X., Ma, Y. Y., Deng, R., … Han, Z. W. (2018). Ultrasensitive determination of thrombin by using an electrode modified with WSe 2 and gold nanoparticles, aptamer-thrombin-aptamer sandwiching, redox cycling, and signal enhancement by alkaline phosphatase. *Microchimica Acta*, *185*(11), 1–7. https://doi.org/10.1007/s00604-018-3028-7
70. Wang, Y., He, X., Wang, K., Ni, X., Su, J., & Chen, Z. (2011). Electrochemical detection of thrombin based on aptamer and ferrocenylhexanethiol loaded silica nanocapsules. *Biosensors and Bioelectronics*, *26*(8), 3536–3541. https://doi.org/10.1016/j.bios.2011.01.041
71. Wang, Z., Zhao, J. C., Lian, H. Z., & Chen, H. Y. (2015). Aptamer-based organic-silica hybrid affinity monolith prepared via "thiol-ene" click reaction for extraction of thrombin. *Talanta*, *138*, 52–58. https://doi.org/10.1016/j.talanta.2015.02.009
72. Yang, H., Ji, J., Liu, Y., Kong, J., & Liu, B. (2009). An aptamer-based biosensor for sensitive thrombin detection. *Electrochemistry Communications*, *11*(1), 38–40. https://doi.org/10.1016/j.elecom.2008.10.024
73. Yigit, M. V., Mazumdar, D., & Lu, Y. (2008). MRI detection of thrombin with aptamer functionalized superparamagnetic iron oxide nanoparticles. *Bioconjugate Chemistry*, *19*(2), 412–417. https://doi.org/10.1021/bc7003928
74. Yin, J., Zhang, A., Dong, C., & Ren, J. (2015). An aptamer-based single particle method for sensitive detection of thrombin using fluorescent quantum dots as labeling probes. *Talanta*, *144*, 13–19. https://doi.org/10.1016/j.talanta.2015.05.034
75. Zavyalova, E., Golovin, A., Pavlova, G., & Kopylov, A. (2013). Module-Activity Relationship of G-quadruplex Based DNA Aptamers for Human Thrombin. *Current Medicinal Chemistry*. https://doi.org/10.2174/09298673113206660283
76. Zavyalova, E., Golovin, A., Reshetnikov, R., Mudrik, N., Panteleyev, D., Pavlova, G., & Kopylov, A. (2011). Novel modular DNA aptamer for human thrombin with high anticoagulant activity. *Current medicinal chemistry*, *18*(22), 3343-3350.
77. Zhang, J., Chen, P., Wu, X. Y., Chen, J. H., Xu, L. J., Chen, G. N., & Fu, F. F. (2011). A signal-on electrochemiluminescence aptamer biosensor for the detection of ultratrace thrombin based on junction-probe. *Biosensors and Bioelectronics*, *26*(5), 2645–2650. https://doi.org/10.1016/j.bios.2010.11.028

***S2: 24mer DNA anti- Adenosine Triphosphate (ATP) aptamer (DH29.36, Huizenga et al., 1994) References***

1. Arnaut, V., Langecker, M., & Simmel, F. C. (2013). Nanopore force spectroscopy of aptamer-ligand complexes. *Biophysical Journal*, *105*(5), 1199–1207. https://doi.org/10.1016/j.bpj.2013.07.047
2. Baaske, P., Wienken, C. J., Reineck, P., Duhr, S., & Braun, D. (2010). Optical thermophoresis for quantifying the buffer dependence of aptamer binding. *Angewandte Chemie - International Edition*, *49*(12), 2238–2241. https://doi.org/10.1002/anie.200903998
3. Biniuri, Y., Albada, B., & Willner, I. (2018). Probing ATP/ATP-Aptamer or ATP-Aptamer Mutant Complexes by Microscale Thermophoresis and Molecular Dynamics Simulations: Discovery of an ATP-Aptamer Sequence of Superior Binding Properties. *Journal of Physical Chemistry B*, *122*(39), 9102–9109. https://doi.org/10.1021/acs.jpcb.8b06802
4. Bu, N. N., Gao, A., He, X. W., & Yin, X. B. (2013). Electrochemiluminescent biosensor of ATP using tetrahedron structured DNA and a functional oligonucleotide for Ru(phen)32+ intercalation and target identification. *Biosensors and Bioelectronics*, *43*(1), 200–204. https://doi.org/10.1016/j.bios.2012.11.027
5. Cai, L., Chen, Z. Z., Dong, X. M., Tang, H. W., & Pang, D. W. (2011). Silica nanoparticles based label-free aptamer hybridization for ATP detection using hoechst33258 as the signal reporter. *Biosensors and Bioelectronics*, *29*(1), 46–52. https://doi.org/10.1016/j.bios.2011.07.064
6. Chen, S. J., Huang, Y. F., Huang, C. C., Lee, K. H., Lin, Z. H., & Chang, H. T. (2008). Colorimetric determination of urinary adenosine using aptamer-modified gold nanoparticles. *Biosensors and Bioelectronics*, *23*(11), 1749–1753. https://doi.org/10.1016/j.bios.2008.02.008
7. Chen, Z., Li, G., Zhang, L., Jiang, J., Li, Z., Peng, Z., & Deng, L. (2008). A new method for the detection of ATP using a quantum-dot-tagged aptamer. *Analytical and Bioanalytical Chemistry*, *392*(6), 1185–1188. https://doi.org/10.1007/s00216-008-2342-z
8. Cheng, S., Zheng, B., Wang, M., Lam, M. H. W., & Ge, X. (2013). Double-functionalized gold nanoparticles with split aptamer for the detection of adenosine triphosphate. *Talanta*, *115*, 506–511. https://doi.org/10.1016/j.talanta.2013.05.065
9. Entzian, C., & Schubert, T. (2016). Studying small molecule-aptamer interactions using MicroScale Thermophoresis (MST). *Methods*, *97*, 27–34. https://doi.org/10.1016/j.ymeth.2015.08.023
10. Feigon, J., Dieckmann, T., & Smith, F. W. (1996). Aptamer structures from A to ζ. *Chemistry and Biology*, *3*(8), 611–617. https://doi.org/10.1016/S1074-5521(96)90127-1
11. Goda, T., & Miyahara, Y. (2011). Thermo-responsive molecular switches for ATP using hairpin DNA aptamers. *Biosensors and Bioelectronics*, *26*(9), 3949–3952. https://doi.org/10.1016/j.bios.2011.02.041
12. Goda, T., & Miyahara, Y. (2012). A hairpin DNA aptamer coupled with groove binders as a smart switch for a field-effect transistor biosensor. *Biosensors and Bioelectronics*, *32*(1), 244–249. https://doi.org/10.1016/j.bios.2011.12.022
13. He, X., Wei, B., & Mi, Y. (2010). Aptamer based reversible DNA induced hydrogel system for molecular recognition and separation. *Chemical Communications*, *46*(34), 6308–6310. https://doi.org/10.1039/c0cc01392g
14. He, Y., Wang, Z. G., Tang, H. W., & Pang, D. W. (2011). Low background signal platform for the detection of ATP: When a molecular aptamer beacon meets graphene oxide. *Biosensors and Bioelectronics*, *29*(1), 76–81. https://doi.org/10.1016/j.bios.2011.07.069

15. Huizenga, D. E., & Szostak, J. W. (1995). A DNA Aptamer That Binds Adenosine and ATP. *Biochemistry*, *34*(2), 656–665. https://doi.org/10.1021/bi00002a033
16. Huo, Y., Qi, L., Lv, X. J., Lai, T., Zhang, J., & Zhang, Z. Q. (2016). A sensitive aptasensor for colorimetric detection of adenosine triphosphate based on the protective effect of ATP-aptamer complexes on unmodified gold nanoparticles. *Biosensors and Bioelectronics*, *78*, 315–320. https://doi.org/10.1016/j.bios.2015.11.043
17. Jiang, Y., Liu, N., Guo, W., Xia, F., & Jiang, L. (2012). Highly-Efficient Gating of Solid-State Nanochannels by DNA Supersandwhich Structure Containing ATP Aptamers: A Nanofluidic IMPLICATION Logic Device. *Journal of the American Chemical Society*, *134*, 15395−15401. https://doi.org/10.1021/ja3053333
18. Jin, F., Lian, Y., Li, J., Zheng, J., Hu, Y., Liu, J., … Yang, R. (2013). Molecule-binding dependent assembly of split aptamer and γ-cyclodextrin: A sensitive excimer signaling approach for aptamer biosensors. *Analytica Chimica Acta*, *799*, 44–50. https://doi.org/10.1016/j.aca.2013.08.012
19. Kashefi-Kheyrabadi, L., & Mehrgardi, M. A. (2012). Aptamer-conjugated silver nanoparticles for electrochemical detection of adenosine triphosphate. *Biosensors and Bioelectronics*, *37*(1), 94–98. https://doi.org/10.1016/j.bios.2012.04.045
20. Kashefi-Kheyrabadi, L., & Mehrgardi, M. A. (2013). Aptamer-based electrochemical biosensor for detection of adenosine triphosphate using a nanoporous gold platform. *Bioelectrochemistry*, *94*, 47–52. https://doi.org/10.1016/j.bioelechem.2013.05.005
21. Kong, L., Xu, J., Xu, Y., Xiang, Y., Yuan, R., & Chai, Y. (2013). A universal and label-free aptasensor for fluorescent detection of ATP and thrombin based on SYBR Green I dye. *Biosensors and Bioelectronics*, *42*(1), 193–197. https://doi.org/10.1016/j.bios.2012.10.064
22. Li, M., Zhang, J., Suri, S., Sooter, L. J., Ma, D., & Wu, N. (2012). Detection of adenosine triphosphate with an aptamer biosensor based on surface-enhanced Raman scattering. *Analytical Chemistry*, *84*(6), 2837–2842. https://doi.org/10.1021/ac203325z
23. Li, N., & Ho, C. M. (2008). Aptamer-based optical probes with separated molecular recognition and signal transduction modules. *Journal of the American Chemical Society*, *130*(8), 2380–2381. https://doi.org/10.1021/ja076787b
24. Li, Y., Sun, L., & Zhao, Q. (2017). Competitive fluorescence anisotropy/polarization assay for ATP using aptamer as affinity ligand and dye-labeled ATP as fluorescence tracer. *Talanta*, *174*(May), 7–13. https://doi.org/10.1016/j.talanta.2017.05.077
25. Liang, A., Ouyang, H., & Jiang, Z. (2011). Resonance scattering spectral detection of trace ATP based on label-free aptamer reaction and nanogold catalysis. *Analyst*, *136*(21), 4514–4519. https://doi.org/10.1039/c1an15542c
26. Liao, W. C., Sohn, Y. S., Riutin, M., Cecconello, A., Parak, W. J., Nechushtai, R., & Willner, I. (2016). The Application of Stimuli-Responsive VEGF- and ATP-Aptamer-Based Microcapsules for the Controlled Release of an Anticancer Drug, and the Selective Targeted Cytotoxicity toward Cancer Cells. *Advanced Functional Materials*, *26*(24), 4262–4273. https://doi.org/10.1002/adfm.201600069
27. Lin, X., Cui, L., Huang, Y., Lin, Y., Xie, Y., Zhu, Z., … Yang, C. J. (2014). Carbon nanoparticle-protected aptamers for highly sensitive and selective detection of biomolecules based on nuclease-assisted target recycling signal amplification. *Chemical Communications*, *50*(57), 7646–7648. https://doi.org/10.1039/c4cc02184c
28. Liu, F., Zhang, J., Chen, R., Chen, L., & Deng, L. (2011). Highly effective colorimetric and visual detection of ATP by a DNAzyme-aptamer sensor. *Chemistry and Biodiversity*, *8*(2), 311–316. https://doi.org/10.1002/cbdv.201000130
29. Liu, S., Wang, Y., Zhang, C., Lin, Y., & Li, F. (2013). Homogeneous electrochemical aptamer-based ATP assay with signal amplification by exonuclease III assisted target recycling. *Chemical Communications*, *49*(23), 2335–2337. https://doi.org/10.1039/c3cc39082a
30. Liu, Z., Chen, S., Liu, B., Wu, J., Zhou, Y., He, L., … Liu, J. (2014). Intracellular detection of ATP using an aptamer beacon covalently linked to graphene oxide resisting nonspecific probe displacement. *Analytical Chemistry*, *86*(24), 12229–12235. https://doi.org/10.1021/ac503358m
31. Merino, E. J., & Weeks, K. M. (2003). Fluorogenic resolution of ligand binding by a nucleic acid aptamer. *Journal of the American Chemical Society*, *125*(41), 12370–12371. https://doi.org/10.1021/ja035299a
32. Modh, H., Witt, M., Urmann, K., Lavrentieva, A., Segal, E., Scheper, T., & Walter, J. G. (2017). Aptamer-based detection of adenosine triphosphate via qPCR. *Talanta*, *172*(January), 199–205. https://doi.org/10.1016/j.talanta.2017.05.037
33. Mukherjee, S., Meshik, X., Choi, M., Farid, S., Datta, D., Lan, Y., … Stroscio, M. A. (2015). A Graphene and Aptamer Based Liquid Gated FET-Like Electrochemical Biosensor to Detect Adenosine Triphosphate. *IEEE Transactions on Nanobioscience*, *14*(8), 967–972. https://doi.org/10.1109/TNB.2015.2501364
34. Nielsen, L. J., Olsen, L. F., & Ozalp, V. C. (2010). Aptamers embedded in polyacrylamide nanoparticles: A tool for in vivo metabolite sensing. *ACS Nano*, *4*(8), 4361–4370. https://doi.org/10.1021/nn100635j
35. Nutiu, R., & Li, Y. (2003). Structure-switching signaling aptamers. *Journal of the American Chemical Society*, *125*(16), 4771–4778. https://doi.org/10.1021/ja028962o
36. Nutiu, R., & Li, Y. (2005). Aptamers with fluorescence-signaling properties. *Methods*, *37*(1), 16–25. https://doi.org/10.1016/j.ymeth.2005.07.001
37. Özalp, V. C., Çam, D., Hernandez, F. J., Hernandez, L. I., Schäfer, T., & Öktem, H. A. (2016). Small molecule detection by lateral flow strips via aptamer-gated silica nanoprobes. *Analyst*, *141*(8), 2595–2599. https://doi.org/10.1039/c6an00273k
38. Özalp, V. C., & Schäfer, T. (2011). Aptamer-based switchable nanovalves for stimuli-responsive drug delivery. *Chemistry - A European Journal*, *17*(36), 9893–9896. https://doi.org/10.1002/chem.201101403
39. Ozalp, V. C., Nielsen, L. J., & Olsen, L. F. (2010). An Aptamer-Based Nanobiosensor for Real-Time Measurements of ATP Dynamics. *ChemBioChem*, *11*, 2538–2541.
40. Park, J. W., Park, Y., & Kim, B. H. (2015). Quencher-free molecular aptamer beacons (QF-MABs) for detection of ATP. *Bioorganic and Medicinal Chemistry Letters*, *25*(20), 4597–4600. https://doi.org/10.1016/j.bmcl.2015.08.052
41. Qiu, H., Liu, Z., Huang, Z., Chen, M., Cai, X., Weng, S., & Lin, X. (2015). Aptamer based turn-off fluorescent ATP assay using DNA concatamers. *Microchimica Acta*, *182*(15–16), 2387–2393. https://doi.org/10.1007/s00604-015-1578-5
42. Sitaula, S., Branch, S. D., & Ali, M. F. (2012). GOx signaling triggered by aptamer-based ATP detection. *Chemical Communications*, *48*(74), 9284–9286. https://doi.org/10.1039/c2cc34279k
43. Song, K., Kong, X., Liu, X., Zhang, Y., Zeng, Q., Tu, L., … Zhang, H. (2012). Aptamer optical biosensor without bio-breakage using upconversion nanoparticles as donors. *Chemical Communications*, *48*(8), 1156–1158. https://doi.org/10.1039/c2cc16817k
44. Tang, Z., Mallikaratchy, P., Yang, R., Kim, Y., Zhu, Z., Wang, H., & Tan, W. (2008). Aptamer switch probe based on intramolecular displacement. *Journal of the American Chemical Society*, *130*(34), 11268–11269. https://doi.org/10.1021/ja804119s
45. Urata, H., Nomura, K., Wada, S. ichi, & Akagi, M. (2007). Fluorescent-labeled single-strand ATP aptamer DNA: Chemo- and enantio-selectivity in sensing adenosine. *Biochemical and Biophysical Research Communications*, *360*(2), 459–463. https://doi.org/10.1016/j.bbrc.2007.06.075
46. Vaish, N. K., Larralde, R., Fraley, A. W., Szostak, J. W., & McLaughlin, L. W. (2003). A novel, modification-dependent ATP-binding aptamer selected from an RNA library incorporating a

cationic functionality. *Biochemistry*, *42*(29), 8842–8851. https://doi.org/10.1021/bi027354i
47. Wang, J., Wang, L., Liu, X., Liang, Z., Song, S., Li, W., … Fan, C. (2007). A gold nanoparticle-based aptamer target binding readout for ATP assay. *Advanced Materials*, *19*(22), 3943–3946. https://doi.org/10.1002/adma.200602256
48. Wang, J., Jiang, Y., Zhou, C., & Fang, X. (2005). Aptamer-based ATP assay using a luminescent light switching complex. *Analytical Chemistry*, *77*(11), 3542–3546. https://doi.org/10.1021/ac050165w
49. Wang, K., Liao, J., Yang, X., Zhao, M., Chen, M., Yao, W., … Lan, X. (2015). A label-free aptasensor for highly sensitive detection of ATP and thrombin based on metal-enhanced PicoGreen fluorescence. *Biosensors and Bioelectronics*, *63*, 172–177. https://doi.org/10.1016/j.bios.2014.07.022
50. Wang, L., Zhu, J., Han, L., Jin, L., Zhu, C., Wang, E., & Dong, S. (2012). Graphene-based aptamer logic gates and their application to multiplex detection. *ACS Nano*, *6*(8), 6659–6666. https://doi.org/10.1021/nn300997f
51. Wang, Y., & Liu, B. (2008). ATP detection using a label-free DNA aptamer and a cationic tetrahedralfluorene. *Analyst*, *133*(11), 1593–1598. https://doi.org/10.1039/b806908e
52. Wang, Y., Wang, Y., & Liu, B. (2008). Fluorescent detection of ATP based on signaling DNA aptamer attached silica nanoparticles. *Nanotechnology*, *19*(41). https://doi.org/10.1088/0957-4484/19/41/415605
53. Wang, Y., Tang, L., Li, Z., Lin, Y., & Li, J. (2014). In situ simultaneous monitoring of ATP and GTP using a graphene oxide nanosheet-based sensing platform in living cells. *Nature Protocols*, *9*(8), 1944–1955. https://doi.org/10.1038/nprot.2014.126
54. White, R. J., Rowe, A. A., & Plaxco, K. W. (2010). Re-engineering aptamers to support reagentless, self-reporting electrochemical sensors. *Analyst*, *135*(3), 589–594. https://doi.org/10.1039/b921253a
55. Wu, J., Coradin, T., & Aimé, C. (2013). Reversible bioresponsive aptamer-based nanocomposites: ATP binding and removal from DNA-grafted silica nanoparticles. *Journal of Materials Chemistry B*, *1*(39), 5353–5359. https://doi.org/10.1039/c3tb20499e
56. Xia, T., Yuan, J., & Fang, X. (2013). Conformational Dynamics of an ATP-Binding DNA Aptamer: A Single-Molecule Study. *Journal of Physical Chemistry B*, *117*(48), 14994–15003. https://doi.org/10.1021/jp4099667
57. Xie, L., Yan, X., & Du, Y. (2014). An aptamer based wall-less LSPR array chip for label-free and high throughput detection of biomolecules. *Biosensors and Bioelectronics*, *53*, 58–64. https://doi.org/10.1016/j.bios.2013.09.031
58. Xu, J., & Wei, C. (2017). The aptamer DNA-templated fluorescence silver nanoclusters: ATP detection and preliminary mechanism investigation. *Biosensors and Bioelectronics*, *87*, 422–427. https://doi.org/10.1016/j.bios.2016.08.079
59. Xu, W., & Lu, Y. (2010). Label-free fluorescent aptamer sensor based on regulation of malachite green fluorescence. *Analytical Chemistry*, *82*(2), 574–578. https://doi.org/10.1021/ac9018473
60. Yamana, K., Ohtani, Y., Nakano, H., & Saito, I. (2003). Bis-pyrene labeled DNA aptamer as an intelligent fluorescent biosensor. *Bioorganic and Medicinal Chemistry Letters*, *13*(20), 3429–3431. https://doi.org/10.1016/S0960-894X(03)00799-6
61. Yin, B. C., Ye, B. C., Wang, H., Zhu, Z., & Tan, W. (2012). Colorimetric logic gates based on aptamer-crosslinked hydrogels. *Chemical Communications*, *48*(9), 1248–1250. https://doi.org/10.1039/c1cc15639j
62. Ying, Y. L., Wang, H. Y., Sutherland, T. C., & Long, Y. T. (2011). Monitoring of an ATP-binding aptamer and its conformational changes using an α-hemolysin nanopore. *Small*, *7*(1), 87–94. https://doi.org/10.1002/smll.201001428
63. Yu, P., He, X., Zhang, L., & Mao, L. (2015). Dual recognition unit strategy improves the specificity of the adenosine triphosphate (ATP) aptamer biosensor for cerebral ATP assay. *Analytical Chemistry*, *87*(2), 1373–1380. https://doi.org/10.1021/ac504249k
64. Zhang, L., Wei, H., Li, J., Li, T., Li, D., Li, Y., & Wang, E. (2010). A carbon nanotubes based ATP apta-sensing platform and its application in cellular assay. *Biosensors and Bioelectronics*, *25*(8), 1897–1901. https://doi.org/10.1016/j.bios.2010.01.002
65. Zhang, S., Wang, K., Li, J., Li, Z., & Sun, T. (2015). Highly efficient colorimetric detection of ATP utilizing a split aptamer target binding strategy and superior catalytic activity of graphene oxide-platinum/gold nanoparticles. *RSC Advances*, *5*(92), 75746–75752. https://doi.org/10.1039/c5ra13550h
66. Zhang, Z., Oni, O., & Liu, J. (2017). New insights into a classic aptamer: Binding sites, cooperativity and more sensitive adenosine detection. *Nucleic Acids Research*, *45*(13), 7593–7601. https://doi.org/10.1093/nar/gkx517
67. Zhao, J., Katsube, S., Yamamoto, J., Yamasaki, K., Miyagishi, M., & Iwai, S. (2015). Analysis of ATP and AMP binding to a DNA aptamer and its imidazole-tethered derivatives by surface plasmon resonance. *Analyst*, *140*(17), 5881–5884. https://doi.org/10.1039/c5an01347j
68. Zhao, M., Liao, L., Wu, M., Lin, Y., Xiao, X., & Nie, C. (2012). Double-receptor sandwich supramolecule sensing method for the determination of ATP based on uranyl-salophen complex and aptamer. *Biosensors and Bioelectronics*, *34*(1), 106–111. https://doi.org/10.1016/j.bios.2012.01.025
69. Zhao, Q., Lv, Q., & Wang, H. (2015). Aptamer fluorescence anisotropy sensors for adenosine triphosphate by comprehensive screening tetramethylrhodamine labeled nucleotides. *Biosensors and Bioelectronics*, *70*, 188–193. https://doi.org/10.1016/j.bios.2015.03.031
70. Zhao, Q., Zhang, Z., Xu, L., Xia, T., Li, N., Liu, J., & Fang, X. (2014). Exonuclease I aided enzyme-linked aptamer assay for small-molecule detection. *Analytical and Bioanalytical Chemistry*, *406*(12), 2949–2955. https://doi.org/10.1007/s00216-014-7705-z
71. Zhou, J., Huang, H., Xuan, J., Zhang, J., & Zhu, J. J. (2010). Quantum dots electrochemical aptasensor based on three-dimensionally ordered macroporous gold film for the detection of ATP. *Biosensors and Bioelectronics*, *26*(2), 834–840. https://doi.org/10.1016/j.bios.2010.05.021
72. Zhou, Z., Du, Y., & Dong, S. (2011). Double-strand DNA-templated formation of copper nanoparticles as fluorescent probe for label-free aptamer sensor. *Analytical Chemistry*, *83*(13), 5122–5127. https://doi.org/10.1021/ac200120g
73. Zuo, X., Song, S., Zhang, J., Pan, D., Wang, L., & Fan, C. (2007). A target-responsive electrochemical aptamer switch (TREAS) for reagentless detection of nanomolar ATP. *Journal of the American Chemical Society*, *129*(5), 1042–1043. https://doi.org/10.1021/ja067024

***S3 The RNA anti-vascular endothelial growth factor (VEGF) binding aptamers (family 1 (15mer), family 2 (22), family 3 (25mer), famiy 4 (19mer), family 5 (27mer), and family 6 (32mer), Jellinek et al., 1994) References***

1. Ahirwar, R., Nahar, S., Aggarwal, S., Ramachandan, S., Maiti, S., & Nahar, P. (2016). In silico selection of an aptamer to estrogen receptor alpha using computational docking employing estrogen response elements as aptamer-alike molecules. *Scientific Reports*, *6*, 21285. https://doi.org/10.1038/srep21285
2. Amadio, M., Govoni, S., & Pascale, A. (2016). Targeting VEGF in eye neovascularization: What's new?: A comprehensive review on

current therapies and oligonucleotide-based interventions under development. *Pharmacol Res, 103*, 253-69. doi: 10.1016/j.phrs.2015.11.027
3. Bacher, J. M., & Ellington, A. D. (1998). Nucleic acid selection as a tool for drug discovery. *Drug Discovery Today, 3*(6), 265-273. https://doi.org/10.1016/S1359-6446(97)01166-5
4. Bayat, P., Nosrati, R., Alibolandi, M., Rafatpanah, H., Abnous, K., Khedri, M., & Ramezani, M. SELEX methods on the road to protein targeting with nucleic acid aptamers. *Biochimie, 154,* 132-155. doi: 10.1016/j.biochi.2018.09.001
5. Bell, C., Lynam, E., Landfair, D. J., Janjic N., & Wiles, M. E. (1999). Oligonucleotide NX1838 inhibits VEGF165-mediated cellular responses in vitro. *In Vitro Cell Dev Biol Anim, 35*(9), 533-42. doi: 10.1007/s11626-999-0064-y
6. Bell, S. D., Denu, J. M., Dixon, J. E., & Ellington, A. D. (1998). RNA molecules that bind to and inhibit the active site of a tyrosine phosphatase. *J Biol Chem, 273*(23), 14309-14. doi: 10.1074/jbc.273.23.14309
7. Biesecker, G., Dihel, L., Enney, K., & Bendele, R. A. (1999). Derivation of RNA aptamer inhibitors of human complement C5. *Immunopharmacology, 42*(1-3), 219-30. doi: 10.1016/s0162-3109(99)00020-x
8. Bridonneau, P., Bunch, S., Tengler, R., Hill, K., Carter, J., Pieken, W., Tinnermeier, D., Lehrman, R., & Drolet, D. W. (1999). Purification of a highly modified RNA-aptamer: Effect of complete denaturation during chromatography on product recovery and specific activity. *Journal of Chromatography B: Biomedical Sciences and Applications, 726*(1-2), 237-247. https://doi.org/10.1016/S0378-4347(99)00037-7
9. Bunka, D. H. J., Stockley, P. G. (2006). Aptamers come of age – at last. *Nature Reviews Microbiology, 4,* 588-596. https://doi.org/10.1038/nrmicro1458
10. Bunka, D. H., Platonova, O., & Stockley, P. G. (2010). Development of aptamer therapeutics. *Curr Opin Pharmacol, 10*(5), 557-62. doi: 10.1016/j.coph.2010.06.009
11. Burmeister, P. E., Lewis, S. D., Silva, R. F., Preiss, J. R., Horwitz, L. R., Pendergrast, P. S., McCauley, T. G., Kurz, J. C., Epstein, D. M., Wilson, C., & Keefe, A. D. (2005). Direct in vitro selection of a 2-O-Methyl aptamer to VEGF. *Chemistry & Biology, 12,* 25-33. doi: 10.1016/j.chembiol.2004.10.017
12. Chelyapov, N. (2006). Allosteric aptamers controlling a signal amplification cascade allow visual detection of molecules at picomolar concentrations. *Biochemistry, 45*(7), 2461-6. doi: 10.1021/bi052106i
13. Chen, C. B., Chernis, G. A., Hoang, V. Q., & Landgraf, R. (2003). Inhibition of heregulin signaling by an aptamer that preferentially binds to the oligomeric form of human epidermal growth factor receptor-3. *PNAS, 100*(16), 9226-9231. https://doi.org/10.1073/pnas.1332660100
14. Chen, W. H., Yang-Sung, S., Fadeev, M., Cecconello, A., Nechushtai, R., & Willner, I. (2018). Targeted VEGF-triggered release of an anti-cancer drug from aptamer-functionalized metal-organic framework nanoparticles. *Nanoscale, 10*(10), 4650-4657. doi: 10.1039/c8nr00193f
15. Chen, Y., Nakamoto, K., Niwa, O., & Corn, R. M. (2012). On-chip synthesis of RNA aptamer microarrays for multiplexed protein biosensing with SPR imaging measurements. *Langmuir : the ACS journal of surfaces and colloids, 28*(22), 8281–8285. https://doi.org/10.1021/la300656c
16. Cockrum, S. E. (2006). Aptamer selections against bacterial toxins and cells. http://hdl.handle.net/2152/21911
17. Ditzler, M. A., Bose, D., Shkriabai, N., Marchand, B., Sarafianos, S. G., Kvaratskhelia, M., & Burke, D. H. (2011). Broad-spectrum aptamer inhibitors of HIV reverse transcriptase closely mimic natural substrates. *Nucleic acids research, 39*(18), 8237–8247. https://doi.org/10.1093/nar/gkr381
18. Drolet, D. W., Green, L. S., Gold, L., & Janjic, N. (2016). Fit for the Eye: Aptamers in Ocular Disorders. *Nucleic acid therapeutics, 26*(3), 127–146. https://doi.org/10.1089/nat.2015.0573
19. Eaton, B. E., Gold, L., & Zichi, D. A. (1995). Let's get specific: the relationship between specificity and affinity. *Chemistry & Biology, 2*(10), 633-638. https://doi.org/10.1016/1074-5521(95)90023-3
20. Eremeeva, E., Fikatas, A., Margamuljana, L., Abramov, M., Schols, D., Groaz, E., & Herdewijn, P. (2019). Highly stable hexitol based XNA aptamers targeting the vascular endothelial growth factor. *Nucleic Acids Research, 47*(10), 4927-4939. https://doi.org/10.1093/nar/gkz252
21. Famulok, M., Hartig, J. S., & Mayer, G. (2007). Functional Aptamers and Aptazymes in Biotechnology, Diagnostics, and Therapy. *Chem. Rev., 107*(9), 3715-3743. https://doi.org/10.1021/cr0306743
22. Gatto, B., & Cavali, M. (2006). From proteins to nucleic acid-based drugs: the role of biotech in anti-VEGF therapy. *Anticancer Agents Med Chem, 6*(4), 287-301. doi: 10.2174/187152006777698178
23. Germer, K., Leonard, M., & Zhang, X. (2013). RNA aptamers and their therapeutic and diagnostic applications. *International journal of biochemistry and molecular biology, 4*(1), 27–40.
24. Gopinath, S. C. B. (2009). Mapping of RNA-protein interactions. *Analytica Chimica Acta, 636*(2), 117-128. https://doi.org/10.1016/j.aca.2009.01.052
25. Gopinath, S. C., Sakamaki, Y., Kawasaki, K., & Kumar, P. K. (2006). An efficient RNA aptamer against human influenza B virus hemagglutinin. *J Biochem, 139*(5), 837-46. doi: 10.1093/jb/mvj095
26. Green, L. S., Jellinek, D., Bell, C., Beebe, L. A., Feistner, B. D., Gill, S. C., Jucker, F. M., & Janjic, N. (1995). Nuclease-resistant nucleic acid ligands to vascular permeability factor/vascular endothelial growth factor. *Chem Biol, 2*(10), 683-95. doi: 10.1016/1074-5521(95)90032-2
27. Hall, B., Hesselberth, J. R., & Ellington, A. D. (2007). Computational selection of nucleic acid biosensors via a slip structure model. *Biosens Bioelectron, 22*(9-10), 1939-47. doi: 10.1016/j.bios.2006.08.019
28. Hunt, S. Structure Determination of Vascular Endothelial Growth Factor Heparin-Binding Domain in Complex with a Dna Aptamer.https://scholar.colorado.edu/concern/graduate_thesis_or_dissertations/1g05fb71z
29. James, W. (2007). Aptamers in the virologists' toolkit. *J Gen Virol, 88*(Pt. 2), 351-64. doi: 10.1099/vir.0.82442-0
30. Janjic, N., Gold, L., Schmidt, P., & Vargeese, C. (2000). *U.S. Patent No. 6,051,698.* Washington, DC: U.S. Patent and Trademark Office.
31. Jellinek, D., Green, L. S., Bell, C., & Janjic, N. (1994). Inhibition of receptor binding by high-affinity RNA ligands to vascular endothelial growth factor. *Biochemistry, 33*(34), 10450-10456. https://doi.org/10.1021/bi00200a028
32. Joshi, M., Sodhi, K. S., Pandey, R., Singh, J., & Goyal, S. (2014). RNA Aptamer Technology. *Indo American Journal of Pharmaceutical Research, 4*(9), 3676-3682.
33. Kanwar, J. R., Shankaranarayanan, J. S., Gurudevan, S., & Kanwar, R. K. (2014). Aptamer-based therapeutics of the past, present and future: from the perspective of eye-related diseases. *Drug Discov Today, 19*(9), 1309-21. doi: 10.1016/j.drudis.2014.02.009
34. Kasahara, Y., & Kuwahara, M. (2012). Artificial specific binders directly recovered from chemically modified nucleic acid libraries. *Journal of nucleic acids, 2012*, 156482. https://doi.org/10.1155/2012/156482
35. Kaur, G., & Roy, I. (2008). Therapeutic applications of aptamers. *Expert Opin Investig Drugs, 17*(1), 43-60. doi: 10.1517/13543784.17.1.43
36. Kayhan, B., & Kayabas, U. (2013). Aptamers: An in vitro Evolution of Therapeutic and Diagnostic Applications in Medicine. *Disease and Molecular Medicine, 1,* 54-60. doi: 10.5455/dmm.20131104040658
37. Kourlas, H., & Schiller, D. S. (2006). Pegaptanib sodium for the treatment of age-related macular degeneration. *Clin Ther, 28*(1), 36-44. doi: 10.1016/j.clinthera.2006.01.009

38. Kujau, M. J., Siebert, A., & Wolfl, S. (1997). Design of leader sequences that improve the efficiency of the enzymatic synthesis of 2′-amino-pyrimidine RNA for in vitro selection. *J Biochem Biophys Methods, 35*(3), 141-51. doi: 10.1016/s0165-022x(97)00039-0
39. Kuwahara, M. (2014). Progress in chemically modified nucleic acid aptamers. *Chemical Biology of Nucleic Acids,* 243-270. https://doi.org/10.1007/978-3-642-54452-1_14
40. Lee, J. H., Canny, M. D., De Erkenez, A., Krilleke, D., Ng, Y. S., Shima, D. T., Pardi, A., & Jucker, F. (2005). A therapeutic aptamer inhibits angiogenesis by specifically targeting the heparin binding domain of VEGF165. *Proc Natl Acad Sci USA, 102*(52), 18902-7. doi: 10.1073/pnas.0509069102
41. Li, Y., Lee, H. J., & Corn, R. M. (2007). Detection of Protein Biomarkers Using RNA Aptamer Microarrays and Enzymatically Amplified Surface Plasmon Resonance Imaging. *Anal. Chem., 79*(3), 1082-1088. https://doi.org/10.1021/ac061849m
42. Libby, A. (2010). NMR Resonance Assignment of the Therapeutic RNA Aptamer Macugen in Complex with its in Vivo Target, the Heparin Binding Domain of VEGF165. https://scholar.colorado.edu/concern/graduate_thesis_or_dissertations/n009w266w
43. Lin, Y., & Jayasena, S. D. (1997). Inhibition of multiple thermostable DNA polymerases by a heterodimeric aptamer. *J Mol Biol, 271*(1), 100-11. doi: 10.1006/jmbi.1997.1165
44. Lollo, B., Steele, F., & Gold, L. (2014). Beyond antibodies: New affinity reagents to unlock the proteome. *Proteomics, 14*(6), 638-44. doi: 10.1002/pmic.201300187
45. Lonne, M., Bolten, S., Lavrentieva A., Stahl, F., Scheper, T., & Walter, J. G. (2015). Development of an aptamer-based affinity purification method for vascular endothelial growth factor. *Biotechnol Rep (Amst), 8*, 16-23. doi: 10.1016/j.btre.2015.08.006
46. Ma, Y., Li, W., Zhou, Z., Qin, X., Wang, D., Gao, Y., Yu, Z., Yin, F., & Li, Z. (2019). Peptide-Aptamer Coassembly Nanocarrier for Cancer Therapy. *Bioconjug Chem, 30*(3), 536-540. doi: 10.1021/acs.bioconjchem.8b00903
47. Majumder, P., Gomes, K. N., & Ulrich, H. (2009). Aptamers: from bench side research towards patented molecules with therapeutic applications. *Expert Opin Ther Pat, 19*(11), 1603-13. doi: 10.1517/13543770903313746
48. Meek, K. N., Rangel, A. E., & Heemstra, J. M. (2016). Enhancing aptamer function and stability via in vitro selection using modified nucleic acids. *Methods, 106*, 29-36. doi: 10.1016/j.ymeth.2016.03.008
49. Mencin, N., Šmuc, T., Vraničar, M., Mavri, J., Hren, M., Galeša, K., Krkoč, P., Ulrich, H., & Šolar, B. (2014). Optimization of SELEX: Comparison of different methods for monitoring the progress of in vitro selection of aptamers. *Journal of Pharmaceutical and Biomedical Analysis, 91,* 151-159. https://doi.org/10.1016/j.jpba.2013.12.031
50. Mie, M., Kai, T., Le, T., Cass, A. E., & Kobatake, E. (2013). Selection of DNA Aptamers with Affinity for Pro-gastrin-Releasing Peptide (proGRP), a Tumor Marker for Small Cell Lung Cancer. *Appl Biochem Biotechnol, 169*(1), 250-5. doi: 10.1007/s12010-012-9956-5
51. Mori, T., Sasaki, J., Aoyama, Y., & Sera, T. (2010). Hypoxia-specific downregulation of endogenous human VEGF-A gene by hypoxia-driven expression of artificial transcription factor. *Mol Biotechnol, 46*(2), 134-9. doi: 10.1007/s12033-010-9288-z
52. Morita, Y., Leslie, M., Kameyama, H., Volk, D. E., & Tanaka, T. (2018). Aptamer Therapeutics in Cancer: Current and Future. *Cancers, 10*(3), 80. https://doi.org/10.3390/cancers10030080
53. Ng, E. W., & Adamis, A. P. (2006). Anti-VEGF aptamer (pegaptanib) therapy for ocular vascular diseases. *Ann N Y Acad Sci, 1082*, 151-71. doi: 10.1196/annals.1348.062
54. Ng, E. W., Shima, D. T., Calias, P., Cunnigham, E. T., Guyer, D. R., & Adamis, A. P. (2006). Pegaptanib, a targeted anti-VEGF aptamer for ocular vascular disease. *Nat Rev Drug Discov, 5*(2), 123-32.
55. Niu, G., & Chen, X. (2010). Vascular endothelial growth factor as an anti-angiogenic target for cancer therapy. *Current drug targets, 11*(8), 1000–1017. https://doi.org/10.2174/138945010791591395
56. Orava, E. W., Jarvik, N., Shek, Y. L., Sidhu, S. S., & Gariepy, J. (2013). A short DNA aptamer that recognizes TNFα and blocks its activity in vitro. *ACS Chem Biol, 8*(1), 170-8. doi: 10.1021/cb3003557
57. Osborne, S. E., Matsumura, I., & Ellington, A. D. (1997). Aptamers as therapeutic and diagnostic reagents: problems and prospects. *Curr Opin Chem Biol, 1*(1), 5-9. doi: 10.1016/s1367-5931(97)80102-0
58. Potty, A. S., Kourentzi, K., Fang, H., Jackson, G. W., Zhang, X., Legge, G. B., & Wilson, R. C. (2009). Biophysical characterization of DNA aptamer interactions with vascular endothelial growth factor. *Biopolymers, 91*(2), 145-56. doi: 10.1002/bip.21097
59. Romero-Lopez, C., Guadix-Arroyo, C., & Berzal-Herranz, A. (2012). Aptamers: Future pharmaceutical drugs. *International Research Journal of Pharmacy and Pharmacology, 2*(13), 311-322.
60. Rosch, J. C., Hollmann, E. K., & Lippmann, E. S. (2016). In vitro selection technologies to enhance biomaterial functionality. *Experimental biology and medicine (Maywood, N.J.), 241*(9), 962–971. https://doi.org/10.1177/1535370216647182
61. Rozenblum, G. T., Kaufman, T., & Vitullo, A. D. (2014). Myelin Basic Protein and a Multiple Sclerosis-related MBP-peptide Bind to Oligonucleotides. Molecular therapy. *Nucleic acids, 3*(9), e192. https://doi.org/10.1038/mtna.2014.43
62. Ruckman, J., Green, L. S., Beeson, J., Waugh, S., Gillette, W. L., Henninger, D. D., Claesson-Welsh, L., & Janjic, N. (1998). 2'-Fluoropyrimidine RNA-based aptamers to the 165-amino acid form of vascular endothelial growth factor (VEGF165). Inhibition of receptor binding and VEGF-induced vascular permeability through interactions requiring the exon 7-encoded domain. *J Biol Chem, 273*(32), 20556-67. doi: 10.1074/jbc.273.32.20556
63. Ruff, K. M., Snyder, T. M., & Liu, D. R. (2010). Enhanced functional potential of nucleic acid aptamer libraries patterned to increase secondary structure. *J. Am. Chem. Soc, 132*(27), 9453-9464. https://doi.org/10.1021/ja103023m
64. Shukla, D., Namperumalsamy, P., Goldbaum, M., & Cunningham, E. T., Jr (2007). Pegaptanib sodium for ocular vascular disease. *Indian journal of ophthalmology, 55*(6), 427–430. https://doi.org/10.4103/0301-4738.36476
65. Silverman, S. K. (2009). Artificial Functional Nucleic Acids: Aptamers, Ribozymes, and Deoxyribozymes Identified by In Vitro Selection. *Functional Nucleic Acids for Analytical Applications,* 47-108. doi: 10.1007/978-0-387-73711-9_3
66. Stein, C. A., & Castanotto, D. (2017). FDA-Approved Oligonucleotide Therapies in 2017. *Molecular therapy : the journal of the American Society of Gene Therapy, 25*(5), 1069–1075. https://doi.org/10.1016/j.ymthe.2017.03.023
67. Stoltenburg, R., Reinemann, C., & Strehlitz, B. (2007). SELEX—A (r)evolutionary method to generate high-affinity nucleic acid ligands. *Biomol Eng, 24*(4), 381-403. doi: 10.1016/j.bioeng.2007.06.001
68. Strehlitz, B., & Stoltenburg, R. (2008). SELEX and its recent optimizations. Aptamers in Bioanalysis, 31-59. doi: 10.1002/9780470380772.ch2
69. Sullivan, R. S. (2017). Expanding quantitative tools for secondary structure analysis of DNA aptamer candidates selected via CompELS. http://hdl.handle.net/1853/60662
70. Tian, Y., Adya, N., Wagner, S., Giam, C. Z., Green, M. R., & Ellington, A. D. (1995). Dissecting protein:protein interactions between transcription factors with an RNA aptamer. *RNA (New York, N.Y.), 1*(3), 317–326.
71. Trujillo, C. A., Nery, A. A., Alves, J. M., Martins, A. H., & Ulrich, H. (2007). Development of the anti-VEGF aptamer to a therapeutic agent for clinical ophthalmology. *Clin Ophthalmol, 1*(4), 393-402.
72. Urvil, P. T., Kakiuchi, N., Zhou, D. M., Shimotohno, K., Kumar, P. K., & Nishikawa, S. (1997). Selection of RNA aptamers that bind specifically to the NS3 protease of hepatitis C virus. *Eur J*

*Biochem, 248*(1), 130-8. doi: 10.1111/j.1432-1033.1997.t01-1-00130.x
73. Waltenberger, J. (1997). Modulation of growth factor action: implications for the treatment of cardiovascular diseases. *Circulation, 96*(11), 4083-94. doi: 10.1161/01.cir.96.11.4083
74. Wang, C., & Yadavalli, V. K. (2014). Spatial recognition and mapping of proteins using DNA aptamers. *Nanotechnology, 25*(45), 455101. doi: 10.1088/0957-4484/25/45/455101
75. Wolfl, S., Kujau, M., Siebert, A., & Wolters, M. (1997). Oligonucleotide Aptamers as Specific Targeting Devices in Diagnostics and Therapy. *Impact of Molecular Biology and New Technical Developments in Diagnostic Imaging,* 135-159. https://doi.org/10.1007/978-3-642-60844-5_10
76. Xi, Z., Huang, R., Deng, Y., & He, N. (2014). Progress in selection and biomedical applications of aptamers. *J Biomed Nanotechnol, 10*(10), 3043-62. doi: 10.1166/jbn.2014.1979
77. Yan, A. C., Bell, K. M., Breeden, M. M., & Ellington, A. D. (2005). Aptamers: prospects in therapeutics and biomedicine. *Front Biosci, 10*, 1802-27. doi: 10.2741/1663
78. Yu, Y., Liang, C., Lv, Q., Li, D., Xu, X., Liu, B., Lu, A., & Zhang, G. (2016). Molecular Selection, Modification and Development of Therapeutic Oligonucleotide Aptamers. *International journal of molecular sciences, 17*(3), 358. https://doi.org/10.3390/ijms17030358
79. Zhang, H., Zhou, L., Zhu, Z., & Yang, C. (2016). Recent progress in aptamer-based functional probes for bioanalysis and biomedicine. *Chemistry, 22*(29), 9886-900. doi: 10.1002/chem.201503543
80. Zhang, X., & Yadavalli, V. K. (2010). Molecular interaction studies of vascular endothelial growth factor with RNA aptamers. *Analyst, 135*(8), 2014-2021. https://doi.org/10.1039/C0AN00200C

***S4: DNA anti platelet-derived growth factor (PDGF-BB) aptamers (36t (1A, 36mer), 41t (1B, 41mer) and 20t (1C, 20mer), Green and Jellinek, 1996) References***

1. Battig, M. R., Huang, Y., Chen, N., & Wang, Y. (2014). Aptamer-functionalized superporous hydrogels for sequestration and release of growth factors regulated via molecular recognition. *Biomaterials*, *35*(27), 8040–8048. https://doi.org/10.1016/j.biomaterials.2014.06.001
2. Battig, M. R., Soontornworajit, B., & Wang, Y. (2012). Programmable release of multiple protein drugs from aptamer-functionalized hydrogels via nucleic acid hybridization. *Journal of the American Chemical Society*, *134*(30), 12410–12413. https://doi.org/10.1021/ja305238a
3. Bi, S., Luo, B., Ye, J., & Wang, Z. (2014). Label-free chemiluminescent aptasensor for platelet-derived growth factor detection based on exonuclease-assisted cascade autocatalytic recycling amplification. *Biosensors and Bioelectronics*, *62*, 208–213. https://doi.org/10.1016/j.bios.2014.06.057
4. Chang, C. C., Wei, S. C., Wu, T. H., Lee, C. H., & Lin, C. W. (2013). Aptamer-based colorimetric detection of platelet-derived growth factor using unmodified gold nanoparticles. *Biosensors and Bioelectronics*, *42*(1), 119–123. https://doi.org/10.1016/j.bios.2012.10.072
5. Chhabra, R., Sharma, J., Ke, Y., Liu, Y., Rinker, S., Lindsay, S., & Yan, H. (2007). Spatially addressable multiprotein nanoarrays templated by aptamer-tagged DNA nanoarchitectures. *Journal of the American Chemical Society*, *129*(34), 10304–10305. https://doi.org/10.1021/ja072410u
6. Chiu, T. C., & Huang, C. C. (2009). *Aptamer-functionalized nano-biosensors*. *Sensors* (Vol. 9). https://doi.org/10.3390/s91210356
7. Csordas, A., Gerdon, A. E., Adams, J. D., Qian, J., Oh, S. S., Xiao, Y., & Soh, H. T. (2010). Detection of proteins in serum by micromagnetic aptamer PCR (MAP) technology. *Angewandte Chemie - International Edition*, *49*(2), 355–358. https://doi.org/10.1002/anie.200904846
8. Fang, L. X., Huang, K. J., & Liu, Y. (2015). Novel electrochemical dual-aptamer-based sandwich biosensor using molybdenum disulfide/carbon aerogel composites and Au nanoparticles for signal amplification. *Biosensors and Bioelectronics*, *71*, 171–178. https://doi.org/10.1016/j.bios.2015.04.031
9. Fang, X., Cao, Z., Beck, T., & Tan, W. (2001). Molecular aptamer for real-time oncoprotein platelet-derived growth factor monitoring by fluorescence anisotropy. *Analytical Chemistry*, *73*(23), 5752–5757. https://doi.org/10.1021/ac010703e
10. Fang, X., Sen, A., Vicens, M., & Tan, W. (2003). Synthetic DNA aptamers to detect protein molecular variants in a high-throughput fluorescence quenching assay. *ChemBioChem*, *4*(9), 829–834. https://doi.org/10.1002/cbic.200300615
11. Fisher, J. D., DiLeo, M. V., & Federspiel, W. J. (2012). Investigating Cytokine Binding Using a Previously Reported TNF-Specific Aptamer. *Open Journal of Applied Sciences*, *02*(03), 135–138. https://doi.org/10.4236/ojapps.2012.23019
12. Fredriksson, S., Gullberg, M., Jarvius, J., Olsson, C., Pietras, K., … Landegren, U. (2014). Protein detection using proximity-dependent DNA Ligation Assays. *Nature Biotechnology*, *20*(May), 473–477. Retrieved from https://www.wakopyrostar.com/blog/post/the-detection-of-endotoxins-via-the-lal-test-the-chromogenic-method/
13. Gao, S., Zheng, X., & Wu, J. (2018). A biolayer interferometry-based enzyme-linked aptamer sorbent assay for real-time and highly sensitive detection of PDGF-BB. *Biosensors and Bioelectronics*, *102*(September 2017), 57–62. https://doi.org/10.1016/j.bios.2017.11.017
14. Green, L. S., Jellinek, D., Jenison, R., Östman, A., Heldin, C. H., & Janjic, N. (1996). Inhibitory DNA ligands to platelet-derived growth factor B-chain. *Biochemistry*, *35*(45), 14413–14424. https://doi.org/10.1021/bi961544+
15. Guo, L., Hao, L., & Zhao, Q. (2016). An aptamer assay using rolling circle amplification coupled with thrombin catalysis for protein detection. *Analytical and Bioanalytical Chemistry*, *408*(17), 4715–4722. https://doi.org/10.1007/s00216-016-9558-0
16. Guo, L., & Zhao, Q. (2016). Thrombin-linked aptamer assay for detection of platelet derived growth factor BB on magnetic beads in a sandwich format. *Talanta*, *158*, 159–164. https://doi.org/10.1016/j.talanta.2016.05.037
17. Guo, L., & Zhao, Q. (2016). Determination of the platelet-derived growth factor BB by a competitive thrombin-linked aptamer-based Fluorometric assay. *Microchimica Acta*, *183*(12), 3229–3235. https://doi.org/10.1007/s00604-016-1978-1
18. Hu, H., Li, H., Zhao, Y., Dong, S., Li, W., Qiang, W., & Xu, D. (2014). Aptamer-functionalized silver nanoparticles for scanometric detection of platelet-derived growth factor-BB. *Analytica Chimica Acta*, *812*, 152–160. https://doi.org/10.1016/j.aca.2013.12.026
19. Huang, C. C., Chiu, S. H., Huang, Y. F., & Chang, H. T. (2007). Aptamer-functionalized gold nanoparticles for turn-on light switch detection of platelet-derived growth factor. *Analytical Chemistry*, *79*(13), 4798–4804. https://doi.org/10.1021/ac0707075
20. Huang, C. C., Huang, Y. F., Cao, Z., Tan, W., & Chang, H. T. (2005). Aptamer-modified gold nanoparticles for colorimetric determination of platelet-derived growth factors and their receptors. *Analytical Chemistry*, *77*(17), 5735–5741. https://doi.org/10.1021/ac050957q

21. Huang, K. J., Shuai, H. L., & Zhang, J. Z. (2016). Ultrasensitive sensing platform for platelet-derived growth factor BB detection based on layered molybdenum selenide-graphene composites and Exonuclease III assisted signal amplification. *Biosensors and Bioelectronics*, *77*, 69–75. https://doi.org/10.1016/j.bios.2015.09.026
22. Huang, Y., Nie, X. M., Gan, S. L., Jiang, J. H., Shen, G. L., & Yu, R. Q. (2008). Electrochemical immunosensor of platelet-derived growth factor with aptamer-primed polymerase amplification. *Analytical Biochemistry*, *382*(1), 16–22. https://doi.org/10.1016/j.ab.2008.07.008
23. Jiang, Y., Fang, X., & Bai, C. (2004). Signaling aptamer/protein binding by a molecular light switch complex. *Analytical Chemistry*, *76*(17), 5230–5235. https://doi.org/10.1021/ac049565u
24. Jin, X., Zhao, J., Zhang, L., Huang, Y., & Zhao, S. (2014). An enhanced fluorescence polarization strategy based on multiple protein-DNA-protein structures for sensitive detection of PDGF-BB. *RSC Advances*, *4*(13), 6850–6853. https://doi.org/10.1039/c3ra44092c
25. Kim, G. Il, Kim, K. W., Oh, M. K., & Sung, Y. M. (2009). The detection of platelet derived growth factor using decoupling of quencher-oligonucleotide from aptamer/quantum dot bioconjugates. *Nanotechnology*, *20*(17). https://doi.org/10.1088/0957-4484/20/17/175503
26. Kim, S. E., Ahn, K. Y., Park, J. S., Kim, K. R., Lee, K. E., Han, S. S., & Lee, J. (2011). Fluorescent ferritin nanoparticles and application to the aptamer sensor. *Analytical Chemistry*, *83*(15), 5834–5843. https://doi.org/10.1021/ac200657s
27. Lai, J., Li, S., Shi, X., Coyne, J., Zhao, N., Dong, F., … Wang, Y. (2017). Displacement and hybridization reactions in aptamer-functionalized hydrogels for biomimetic protein release and signal transduction. *Chemical Science*, *8*(11), 7306–7311. https://doi.org/10.1039/c7sc03023a
28. Lai, R. Y., Plaxco, K. W., & Heeger, A. J. (2007). Aptamer-based electrochemical detection of picomolar platelet-derived growth factor directly in blood serum. *Analytical Chemistry*, *79*(1), 229–233. https://doi.org/10.1021/ac061592s
29. Leppänen, O., Janjic, N., Carlsson, M. A., Pietras, K., Levin, M., Vargeese, C., … Heldin, C. H. (2000). Intimal hyperplasia recurs after removal of PDGF-AB and -BB inhibition in the rat carotid artery injury model. *Arteriosclerosis, Thrombosis, and Vascular Biology*, *20*(11). https://doi.org/10.1161/01.atv.20.11.e89
30. Li, F., Zhang, H., Lai, C., Li, X. F., & Le, X. C. (2012). A molecular translator that acts by binding-induced DNA strand displacement for a homogeneous protein assay. *Angewandte Chemie - International Edition*, *51*(37), 9317–9320. https://doi.org/10.1002/anie.201202677
31. Li, H., Wang, M., Wang, C., Li, W., Qiang, W., & Xu, D. (2013). Silver nanoparticle-enhanced fluorescence resonance energy transfer sensor for human platelet-derived growth factor-BB detection. *Analytical Chemistry*, *85*(9), 4492–4499. https://doi.org/10.1021/ac400047d
32. Li, J., Jia, Y., Zheng, J., Zhong, W., Shen, G., Yang, R., & Tan, W. (2013). Aptamer degradation inhibition combined with DNAzyme cascade-based signal amplification for colorimetric detection of proteins. *Chemical Communications*, *49*(55), 6137–6139. https://doi.org/10.1039/c3cc42148a
33. Li, W., Jiang, W., & Wang, L. (2016). Self-locked aptamer probe mediated cascade amplification strategy for highly sensitive and selective detection of protein and small molecule. *Analytica Chimica Acta*, *940*, 1–7. https://doi.org/10.1016/j.aca.2016.08.017
34. Liang, J., Wei, R., He, S., Liu, Y., Guo, L., & Li, L. (2013). A highly sensitive and selective aptasensor based on graphene oxide fluorescence resonance energy transfer for the rapid determination of oncoprotein PDGF-BB. *Analyst*, *138*(6), 1726–1732. https://doi.org/10.1039/c2an36529d
35. Liao, W., & Cui, X. T. (2007). Reagentless aptamer based impedance biosensor for monitoring a neuro-inflammatory cytokine PDGF. *Biosensors and Bioelectronics*, *23*(2), 218–224. https://doi.org/10.1016/j.bios.2007.04.004
36. Liu, J. J., Song, X. R., Wang, Y. W., Zheng, A. X., Chen, G. N., & Yang, H. H. (2012). Label-free and fluorescence turn-on aptasensor for protein detection via target-induced silver nanoclusters formation. *Analytica Chimica Acta*, *749*, 70–74. https://doi.org/10.1016/j.aca.2012.09.002
37. Liu, Y. M., Zhou, M., Liu, Y. Y., Shi, G. F., Zhang, J. J., Cao, J. T., … Chen, Y. H. (2014). Fabrication of electrochemiluminescence aptasensor based on in situ growth of gold nanoparticles on layered molybdenum disulfide for sensitive detection of platelet-derived growth factor-BB. *RSC Advances*, *4*(44), 22888–22893. https://doi.org/10.1039/c4ra02162b
38. Lu, C., Shahzad, M. M., Moreno-Smith, M., Lin, Y., Jennings, N. B., Allen, J. K., ... & Nick, A. M. (2010). Targeting pericytes with a PDGF-B aptamer in human ovarian carcinoma models. *Cancer biology & therapy*, *9*(3), 176-182.
39. Lv, L., Guo, L., & Zhao, Q. (2015). Aptamer and rolling circle amplification-involved sandwich assay for platelet-derived growth factor-BB with absorbance analysis. *Analytical Methods*, *7*(5), 1855–1859. https://doi.org/10.1039/c4ay02921f
40. Ma, X., Chen, Z., Zhou, J., Weng, W., Zheng, O., Lin, Z., … Chen, G. (2014). Aptamer-based portable biosensor for platelet-derived growth factor-BB (PDGF-BB) with personal glucose meter readout. *Biosensors and Bioelectronics*, *55*, 412–416. https://doi.org/10.1016/j.bios.2013.12.041
41. Neumann, O., Zhang, D., Tam, F., Lal, S., Wittung-Stafshede, P., & Halas, N. J. (2009). Direct optical detection of aptamer conformational changes induced by target molecules. *Analytical Chemistry*, *81*(24), 10002–10006. https://doi.org/10.1021/ac901849k
42. Ostendorf, T., Kunter, U., Gröne, H. J., Bahlmann, F., Kawachi, H., Shimizu, F., … Floege, J. (2001). Specific antagonism of PDGF prevents renal scarring in experimental glomerulonephritis. *Journal of the American Society of Nephrology*, *12*(5), 909–918.
43. Pei, R., & Stojanovic, M. N. (2008). Study of thiazole orange in aptamer-based dye-displacement assays. *Analytical and Bioanalytical Chemistry*, *390*(4), 1093–1099. https://doi.org/10.1007/s00216-007-1773-2
44. Pietras, K., Rubin, K., Sjöblom, T., Buchdunger, E., Sjöquist, M., Heldin, C. H., & Östman, A. (2002). Inhibition of PDGF receptor signaling in tumor stroma enhances antitumor effect of chemotherapy. *Cancer Research*, *62*(19), 5476–5484.
45. Ruslinda, A. R., Penmatsa, V., Ishii, Y., Tajima, S., & Kawarada, H. (2012). Highly sensitive detection of platelet-derived growth factor on a functionalized diamond surface using aptamer sandwich design. *Analyst*, *137*(7), 1692–1697. https://doi.org/10.1039/c2an15933c
46. Sennino, B., Falcón, B. L., McCauley, D., Le, T., McCauley, T., Kurz, J. C., … McDonald, D. M. (2007). Sequential loss of tumor vessel pericytes and endothelial cells after inhibition of platelet-derived growth factor B by selective aptamer AX102. *Cancer Research*, *67*(15), 7358–7367. https://doi.org/10.1158/0008-5472.CAN-07-0293
47. Song, W., Zhu, K., Cao, Z., Lau, C., & Lu, J. (2012). Hybridization chain reaction-based aptameric system for the highly selective and sensitive detection of protein. *Analyst*, *137*(6), 1396–1401. https://doi.org/10.1039/c2an16232f
48. Soontornworajit, B., Zhou, J., Shaw, M. T., Fan, T. H., & Wang, Y. (2010). Hydrogel functionalization with DNA aptamers for sustained PDGF-BB release. *Chemical Communications*, *46*(11), 1857–1859. https://doi.org/10.1039/b924909e
49. Soontornworajit, B., Zhou, J., Zhang, Z., & Wang, Y. (2010). Aptamer-functionalized in situ injectable hydrogel for controlled protein release. *Biomacromolecules*, *11*(10), 2724–2730. https://doi.org/10.1021/bm100774t
50. Tang, L., Liu, Y., Ali, M. M., Kang, D. K., Zhao, W., & Li, J. (2012). Colorimetric and ultrasensitive bioassay based on a dual-amplification system using aptamer and DNAzyme. *Analytical Chemistry*, *84*(11), 4711–4717. https://doi.org/10.1021/ac203274k
51. Vicens, M. C., Sen, A., Vanderlaan, A., Drake, T. J., & Tan, W. (2005). Investigation of molecular beacon aptamer-based bioassay

for platelet-derived growth factor detection. *ChemBioChem*, *6*(5), 900–907. https://doi.org/10.1002/cbic.200400308
52. Vu, C. Q., Rotkrua, P., Soontornworajit, B., & Tantirungrotechai, Y. (2018). Effect of PDGF-B aptamer on PDGFRβ/PDGF-B interaction: Molecular dynamics study. *Journal of Molecular Graphics and Modelling*, *82*, 145–156. https://doi.org/10.1016/j.jmgm.2018.04.012
53. Vu, C. Q., Rotkrua, P., Tantirungrotechai, Y., & Soontornworajit, B. (2017). Oligonucleotide Hybridization Combined with Competitive Antibody Binding for the Truncation of a High-Affinity Aptamer. *ACS Combinatorial Science*, *19*(10), 609–617. https://doi.org/10.1021/acscombsci.6b00163
54. Vu, C. Q., Tantirungrotechai, Y., & Soontornworajit, B. (2016). Truncation of PDGF-BB Aptamer by Secondary Structural Analysis and Immunoassay. *International Journal of Pharma Medicine and Biological Sciences*, *5*(1), 86–90. https://doi.org/10.18178/ijpmbs.5.1.86-90
55. Wang, J., Meng, W., Zheng, X., Liu, S., & Li, G. (2009). Combination of aptamer with gold nanoparticles for electrochemical signal amplification: Application to sensitive detection of platelet-derived growth factor. *Biosensors and Bioelectronics*, *24*(6), 1598–1602. https://doi.org/10.1016/j.bios.2008.08.030
56. Wang, Q., Zheng, H., Gao, X., Lin, Z., & Chen, G. (2013). A label-free ultrasensitive electrochemical aptameric recognition system for protein assay based on hyperbranched rolling circle amplification. *Chemical Communications*, *49*(97), 11418–11420. https://doi.org/10.1039/c3cc46274a
57. Wang, R., Lu, D., Bai, H., Jin, C., Yan, G., Ye, M., … Tan, W. (2016). Using modified aptamers for site specific protein-aptamer conjugations. *Chemical Science*, *7*(3), 2157–2161. https://doi.org/10.1039/c5sc02631h
58. Xie, S., & Walton, S. P. (2010). Development of a dual-aptamer-based multiplex protein biosensor. *Biosensors and Bioelectronics*, *25*(12), 2663–2668. https://doi.org/10.1016/j.bios.2010.04.034
59. Yang, C. J., Jockusch, S., Vicens, M., Turro, N. J., & Tan, W. (2005). Light-switching excimer probes for rapid protein monitoring in complex biological fluids. *Proceedings of the National Academy of Sciences of the United States of America*, *102*(48), 17278–17283. https://doi.org/10.1073/pnas.0508821102
60. Yang, X. H., Sun, S., Liu, P., Wang, K. M., Wang, Q., Liu, J. B., … He, L. L. (2014). A novel fluorescent detection for PDGF-BB based on dsDNA-templated copper nanoparticles. *Chinese Chemical Letters*, *25*(1), 9–14. https://doi.org/10.1016/j.cclet.2013.10.032
61. Ye, S. J., Zhai, X. M., Wu, Y. Y., & Kuang, S. P. (2016). Dual-primer self-generation SERS signal amplification assay for PDGF-BB using label-free aptamer. *Biosensors and Bioelectronics*, *79*, 130–135. https://doi.org/10.1016/j.bios.2015.11.090
62. Yi, Y., Huang, Y., Zhu, G., Lin, F., Zhang, L., Li, H., … Yao, S. (2013). A colorimetric and fluorescence sensing platform for two analytes in homogenous solution based on aptamer-modified gold nanoparticles. *Analytical Methods*, *5*(10), 2477–2484. https://doi.org/10.1039/c3ay40087e
63. Zhang, D., Zhao, Q., Zhao, B., & Wang, H. (2012). Fluorescence anisotropy reduction of allosteric aptamer for sensitive and specific protein signaling. *Analytical Chemistry*, *84*(7), 3070–3074. https://doi.org/10.1021/ac3004133
64. Zhang, H., Li, X. F., & Le, X. C. (2009). Differentiation and detection of PDGF isomers and their receptors by tunable aptamer capillary electrophoresis. *Analytical Chemistry*, *81*(18), 7795–7800. https://doi.org/10.1021/ac901471w
65. Zhang, H., Li, F., Chen, H., Ma, Y., Qi, S., Chen, X., & Zhou, L. (2015). AuNPs colorimetric sensor for detecting platelet-derived growth factor-BB based on isothermal target-triggering strand displacement amplification. *Sensors and Actuators, B: Chemical*, *207*(PartA), 748–755. https://doi.org/10.1016/j.snb.2014.11.007
66. Zhang, J. J., Cao, J. T., Shi, G. F., Huang, K. J., Liu, Y. M., & Ren, S. W. (2015). A luminol electrochemiluminescence aptasensor based on glucose oxidase modified gold nanoparticles for measurement of platelet-derived growth factor BB. *Talanta*, *132*, 65–71. https://doi.org/10.1016/j.talanta.2014.08.058
67. Zhang, Z. Z., & Zhang, C. Y. (2012). Highly sensitive detection of protein with aptamer-based target-triggering two-stage amplification. *Analytical Chemistry*, *84*(3), 1623–1629. https://doi.org/10.1021/ac2029002
68. Zhang, Z., Liu, C., Yang, C., Wu, Y., Yu, F., Chen, Y., & Du, J. (2018). Aptamer-Patterned Hydrogel Films for Spatiotemporally Programmable Capture and Release of Multiple Proteins. *ACS Applied Materials and Interfaces*, *10*(10), 8546–8554. https://doi.org/10.1021/acsami.8b00191
69. Zhou, C., Jiang, Y., Hou, S., Ma, B., Fang, X., & Li, M. (2006). Detection of oncoprotein platelet-derived growth factor using a fluorescent signaling complex of an aptamer and TOTO. *Analytical and Bioanalytical Chemistry*, *384*(5), 1175–1180. https://doi.org/10.1007/s00216-005-0276-2
70. Zhou, L., Ou, L. J., Chu, X., Shen, G. L., & Yu, R. Q. (2007). Aptamer-based rolling circle amplification: A platform for electrochemical detection of protein. *Analytical Chemistry*, *79*(19), 7492–7500. https://doi.org/10.1021/ac071059s
71. Zhu, D., Zhou, X., & Xing, D. (2010). A new kind of aptamer-based immunomagnetic electrochemiluminescence assay for quantitative detection of protein. *Biosensors and Bioelectronics*, *26*(1), 285–288. https://doi.org/10.1016/j.bios.2010.06.028
72. Zhu, D., Zhou, X., & Xing, D. (2012). Ultrasensitive aptamer-based bio bar code immunomagnetic separation and electrochemiluminescence method for the detection of protein. *Analytica Chimica Acta*, *725*, 39–43. https://doi.org/10.1016/j.aca.2012.03.006

***S5: DNA anti-cocaine aptamers (MNS4.1 (2A 30mer) and MNS7.9/MSN6 (1A, 38mer), Stojanovic et al., 2001) References***

1. Baker, B. R., Lai, R. Y., Wood, M. S., Doctor, E. H., Heeger, A. J., & Plaxco, K. W. (2006). An electronic, aptamer-based small-molecule sensor for the rapid, label-free detection of cocaine in adulterated samples and biological fluids. *Journal of the American Chemical Society*, *128*(10), 3138–3139. https://doi.org/10.1021/ja056957p
2. Cekan, P., Jonsson, E. Ö., & Sigurdsson, S. T. (2009). Folding of the cocaine aptamer studied by EPR and fluorescence spectroscopies using the bifunctional spectroscopic probe Ç. *Nucleic Acids Research*, *37*(12), 3990–3995. https://doi.org/10.1093/nar/gkp277
3. Chen, J., Jiang, J., Gao, X., Liu, G., Shen, G., & Yu, R. (2008). A new aptameric biosensor for cocaine based on surface-enhanced raman scattering spectroscopy. *Chemistry - A European Journal*, *14*(27), 8374–8382. https://doi.org/10.1002/chem.200701307

4. Chen, Z., Tan, Y., Xu, K., Zhang, L., Qiu, B., Guo, L., … Chen, G. (2016). Stimulus-response mesoporous silica nanoparticle-based chemiluminescence biosensor for cocaine determination. *Biosensors and Bioelectronics*, *75*, 8–14. https://doi.org/10.1016/j.bios.2015.08.006
5. Deng, Q. P., Tie, C., Zhou, Y. L., & Zhang, X. X. (2012). Cocaine detection by structure-switch aptamer-based capillary zone electrophoresis. *Electrophoresis*, *33*(9–10), 1465–1470. https://doi.org/10.1002/elps.201100680
6. Freeman, R., Li, Y., Tel-Vered, R., Sharon, E., Elbaz, J., & Willner, I. (2009). Self-assembly of supramolecular aptamer structures for optical or electrochemical sensing. *Analyst*, *134*(4), 653–656. https://doi.org/10.1039/b822836c
7. Ge, J., Liu, Z., & Zhao, X. S. (2012). Cocaine detection in blood serum using aptamer biosensor on gold nanoparticles and progressive dilution. *Chinese Journal of Chemistry*, *30*(9), 2023–2028. https://doi.org/10.1002/cjoc.201200256
8. Golub, E., Pelossof, G., Freeman, R., Zhang, H., & Willner, I. (2009). Electrochemical, photoelectrochemical, and surface plasmon resonance detection of cocaine using supramolecular aptamer complexes and metallic or semiconductor nanoparticles. *Analytical Chemistry*, *81*(22), 9291–9298. https://doi.org/10.1021/ac901551q
9. Grytz, C. M., Marko, A., Cekan, P., Sigurdsson, S. T., & Prisner, T. F. (2016). Flexibility and conformation of the cocaine aptamer studied by PELDOR. *Physical Chemistry Chemical Physics*, *18*(4), 2993–3002. https://doi.org/10.1039/c5cp06158j
10. He, J. L., Wu, Z. S., Zhou, H., Wang, H. Q., Jiang, J. H., Shen, G. L., & Yu, R. Q. (2010). Fluorescence aptameric sensor for strand displacement amplification detection of cocaine. *Analytical Chemistry*, *82*(4), 1358–1364. https://doi.org/10.1021/ac902416u
11. Hilton, J. P., Nguyen, T. H., Pei, R., Stojanovic, M., & Lin, Q. (2011). A microfluidic affinity sensor for the detection of cocaine. *Sensors and Actuators, A: Physical*, *166*(2), 241–246. https://doi.org/10.1016/j.sna.2009.12.006
12. Hua, M., Li, P., Li, L., Huang, L., Zhao, X., Feng, Y., & Yang, Y. (2011). Quantum dots as immobilized substrate for electrochemical detection of cocaine based on conformational switching of aptamer. *Journal of Electroanalytical Chemistry*, *662*(2), 306–311. https://doi.org/10.1016/j.jelechem.2011.08.017
13. Hua, M., Tao, M., Wang, P., Zhang, Y., Wu, Z., Chang, Y., & Yang, Y. (2010). Label-free electrochemical cocaine aptasensor based on a target-inducing aptamer switching conformation. *Analytical Sciences*, *26*(12), 1265–1270. https://doi.org/10.2116/analsci.26.1265
14. Kang, K., Sachan, A., Nilsen-Hamilton, M., & Shrotriya, P. (2011). Aptamer functionalized microcantilever sensors for cocaine detection. *Langmuir*, *27*(23), 14696–14702. https://doi.org/10.1021/la202067y
15. Kawano, R., Osaki, T., Sasaki, H., Takinoue, M., Yoshizawa, S., & Takeuchi, S. (2011). Rapid detection of a cocaine-binding aptamer using biological nanopores on a chip. *Journal of the American Chemical Society*, *133*(22), 8474–8477. https://doi.org/10.1021/ja2026085
16. Li, K., Qin, W., Li, F., Zhao, X., Jiang, B., Wang, K., … Li, D. (2013). Nanoplasmonic imaging of latent fingerprints and identification of cocaine. *Angewandte Chemie - International Edition*, *52*(44), 11542–11545. https://doi.org/10.1002/anie.201305980
17. Li, X., Qi, H., Shen, L., Gao, Q., & Zhang, C. (2008). Electrochemical aptasensor for the determination of cocaine incorporating gold nanoparticles modification. *Electroanalysis*, *20*(13), 1475–1482. https://doi.org/10.1002/elan.200704193
18. Li, Y., Qi, H., Peng, Y., Yang, J., & Zhang, C. (2007). Electrogenerated chemiluminescence aptamer-based biosensor for the determination of cocaine. *Electrochemistry Communications*, *9*(10), 2571–2575. https://doi.org/10.1016/j.elecom.2007.07.038
19. Li, Y., Ji, X., & Liu, B. (2011). Chemiluminescence aptasensor for cocaine based on double-functionalized gold nanoprobes and functionalized magnetic microbeads. *Analytical and Bioanalytical Chemistry*, *401*(1), 213–219. https://doi.org/10.1007/s00216-011-5064-6
20. Liu, Y., & Zhao, Q. (2017). Direct fluorescence anisotropy assay for cocaine using tetramethylrhodamine-labeled aptamer. *Analytical and Bioanalytical Chemistry*, *409*(16), 3993–4000. https://doi.org/10.1007/s00216-017-0349-z
21. Ma, C., Wang, W., Yang, Q., Shi, C., & Cao, L. (2011). Cocaine detection via rolling circle amplification of short DNA strand separated by magnetic beads. *Biosensors and Bioelectronics*, *26*(7), 3309–3312. https://doi.org/10.1016/j.bios.2011.01.003
22. Madru, B., Chapuis-Hugon, F., Peyrin, E., & Pichon, V. (2009). Determination of cocaine in human plasma by selective solid-phase extraction using an aptamer-based sorbent. *Analytical Chemistry*, *81*(16), 7081–7086. https://doi.org/10.1021/ac9006667
23. Mokhtarzadeh, A., Ezzati Nazhad Dolatabadi, J., Abnous, K., de la Guardia, M., & Ramezani, M. (2015). Nanomaterial-based cocaine aptasensors. *Biosensors and Bioelectronics*, *68*, 95–106. https://doi.org/10.1016/j.bios.2014.12.052
24. Neves, M. A. D., Blaszykowski, C., Bokhari, S., & Thompson, M. (2015). Ultra-high frequency piezoelectric aptasensor for the label-free detection of cocaine. *Biosensors and Bioelectronics*, *72*, 383–392. https://doi.org/10.1016/j.bios.2015.05.038
25. Neves, M. A. D., Blaszykowski, C., & Thompson, M. (2016). Utilizing a Key Aptamer Structure-Switching Mechanism for the Ultrahigh Frequency Detection of Cocaine. *Analytical Chemistry*, *88*(6), 3098–3106. https://doi.org/10.1021/acs.analchem.5b04010
26. Neves, M. A. D., Reinstein, O., & Johnson, P. E. (2010). Defining a stem length-dependent binding mechanism for the cocaine-binding aptamer. A combined NMR and calorimetry study. *Biochemistry*, *49*(39), 8478–8487. https://doi.org/10.1021/bi100952k
27. Neves, M. A. D., Reinstein, O., Saad, M., & Johnson, P. E. (2010). Defining the secondary structural requirements of a cocaine-binding aptamer by a thermodynamic and mutation study. *Biophysical Chemistry*, *153*(1), 9–16. https://doi.org/10.1016/j.bpc.2010.09.009
28. Neves, M. A. D., Shoara, A. A., Reinstein, O., Abbasi Borhani, O., Martin, T. R., & Johnson, P. E. (2017). Optimizing Stem Length to Improve Ligand Selectivity in a Structure-Switching Cocaine-Binding Aptamer. *ACS Sensors*, *2*(10), 1539–1545. https://doi.org/10.1021/acssensors.7b00619
29. Neves, M. A. D., Slavkovic, S., Churcher, Z. R., & Johnson, P. E. (2017). Salt-mediated two-site ligand binding by the cocaine-binding aptamer. *Nucleic Acids Research*, *45*(3), 1041–1048. https://doi.org/10.1093/nar/gkw1294
30. Nie, J., Deng, Y., Deng, Q. P., Zhang, D. W., Zhou, Y. L., & Zhang, X. X. (2013). A self-assemble aptamer fragment/target complex based high-throughput colorimetric aptasensor using enzyme linked aptamer assay. *Talanta*, *106*, 309–314. https://doi.org/10.1016/j.talanta.2012.11.018
31. Reinstein, O., Yoo, M., Han, C., Palmo, T., Beckham, S. A., Wilce, M. C. J., & Johnson, P. E. (2013). Quinine binding by the cocaine-binding aptamer. thermodynamic and hydrodynamic analysis of high-affinity binding of an off-target ligand. *Biochemistry*, *52*(48), 8652–8662. https://doi.org/10.1021/bi4010039
32. Roncancio, D., Yu, H., Xu, X., Wu, S., Liu, R., Debord, J., … Xiao, Y. (2014). A label-free aptamer-fluorophore assembly for rapid and specific detection of cocaine in biofluids. *Analytical Chemistry*, *86*(22), 11100–11106. https://doi.org/10.1021/ac503360n
33. Roushani, M., & Shahdost-Fard, F. (2015). A highly selective and sensitive cocaine aptasensor based on covalent attachment of the aptamer-functionalized AuNPs onto nanocomposite as the support platform. *Analytica Chimica Acta*, *853*(1), 214–221. https://doi.org/10.1016/j.aca.2014.09.031
34. Roushani, M., & Shahdost-Fard, F. (2015). A novel ultrasensitive aptasensor based on silver nanoparticles measured via enhanced voltammetric response of electrochemical reduction of riboflavin as redox probe for cocaine detection. *Sensors and Actuators, B:*

Chemical, 207(PartA), 764–771. https://doi.org/10.1016/j.snb.2014.10.131
35. Sachan, A., Ilgu, M., Kempema, A., Kraus, G. A., & Nilsen-Hamilton, M. (2016). Specificity and Ligand Affinities of the Cocaine Aptamer: Impact of Structural Features and Physiological NaCl. *Analytical Chemistry*, *88*(15), 7715–7723. https://doi.org/10.1021/acs.analchem.6b01633
36. Sassolas, A., Blum, L. J., & Leca-Bouvier, B. D. (2011). Homogeneous assays using aptamers. *Analyst*, *136*(2), 257–274. https://doi.org/10.1039/c0an00281j
37. Sharon, E., Freeman, R., Ran, T. V., & Willner, I. (2009). Impedimetric or ion-sensitive field-effect transistor (ISFET) aptasensors based on the self-assembly of au nanoparticle-functionalized supramolecular aptamer nanostructures. *Electroanalysis*, *21*(11), 1291–1296. https://doi.org/10.1002/elan.200804565
38. Shi, Y., Dai, H., Sun, Y., Hu, J., Ni, P., & Li, Z. (2013). Fluorescent sensing of cocaine based on a structure switching aptamer, gold nanoparticles and graphene oxide. *Analyst*, *138*(23), 7152–7156. https://doi.org/10.1039/c3an00897e
39. Slavkovic, S., Altunisik, M., Reinstein, O., & Johnson, P. E. (2015). Structure-affinity relationship of the cocaine-binding aptamer with quinine derivatives. *Bioorganic and Medicinal Chemistry*, *23*(10), 2593–2597. https://doi.org/10.1016/j.bmc.2015.02.052
40. Stojanovic, M. N., de Prada, P., & Landry, D. W. (2001). Aptamer-based folding fluorescent sensor for cocaine. *Journal of the American Chemical Society*, *123*(21), 4928–4931. https://doi.org/10.1021/ja0038171
41. Swensen, J. S., Xiao, Y., Ferguson, B. S., Lubin, A. A., Lai, R. Y., Heeger, A. J., … Soh, H. T. (2009). Continuous, real-time monitoring of cocaine in undiluted blood serum via a microfluidic, electrochemical aptamer-based sensor. *Journal of the American Chemical Society*, *131*(12), 4262–4266. https://doi.org/10.1021/ja806531z
42. Taghavi, S., Ayatollahi, S., Alibolandi, M., Lavaee, P., Ramezani, M., & Abnous, K. (2014). A novel label-free cocaine assay based on aptamer-wrapped single-walled carbon nanotubes Cocaine assay by aptamer–SWNTs conjugate. *Nanomed J*, *1*(2), 100–106. Retrieved from http://nmj.mums.ac.ir
43. Tang, Y., Long, F., Gu, C., Wang, C., Han, S., & He, M. (2016). Reusable split-aptamer-based biosensor for rapid detection of cocaine in serum by using an all-fiber evanescent wave optical biosensing platform. *Analytica Chimica Acta*, *933*, 182–188. https://doi.org/10.1016/j.aca.2016.05.021
44. Wang, L., Musile, G., & McCord, B. R. (2018). An aptamer-based paper microfluidic device for the colorimetric determination of cocaine. *Electrophoresis*, *39*(3), 470–475. https://doi.org/10.1002/elps.201700254
45. Wang, J., Wang, L., Liu, X., Liang, Z., Song, S., Li, W., ... & Fan, C. (2007). A gold nanoparticle-based aptamer target binding readout for ATP assay. *Advanced Materials*, *19*(22), 3943-3946.
46. Wen, Y., Pei, H., Wan, Y., Su, Y., Huang, Q., Song, S., & Fan, C. (2011). DNA nanostructure-decorated surfaces for enhanced aptamer-target binding and electrochemical cocaine sensors. *Analytical Chemistry*, *83*(19), 7418–7423. https://doi.org/10.1021/ac201491p
47. Wu, C., Yan, L., Wang, C., Lin, H., Wang, C., Chen, X., & Yang, C. J. (2010). A general excimer signaling approach for aptamer sensors. *Biosensors and Bioelectronics*, *25*(10), 2232–2237. https://doi.org/10.1016/j.bios.2010.02.030
48. Yan, X., Cao, Z., Lau, C., & Lu, J. (2010). DNA aptamer folding on magnetic beads for sequential detection of adenosine and cocaine by substrate-resolved chemiluminescence technology. *Analyst*, *135*(9), 2400–2407. https://doi.org/10.1039/c0an00163e
49. Yang, Z., Castrignanò, E., Estrela, P., Frost, C. G., & Kasprzyk-Hordern, B. (2016). Community Sewage Sensors towards Evaluation of Drug Use Trends: Detection of Cocaine in Wastewater with DNA-Directed Immobilization Aptamer Sensors. *Scientific Reports*, *6*(February), 1–10. https://doi.org/10.1038/srep21024
50. Yin, B. C., Ye, B. C., Wang, H., Zhu, Z., & Tan, W. (2012). Colorimetric logic gates based on aptamer-crosslinked hydrogels. *Chemical Communications*, *48*(9), 1248–1250. https://doi.org/10.1039/c1cc15639j
51. Yu, H., Canoura, J., Guntupalli, B., Lou, X., & Xiao, Y. (2016). A cooperative-binding split aptamer assay for rapid, specific and ultra-sensitive fluorescence detection of cocaine in saliva. *Chemical Science*, *8*(1), 131–141. https://doi.org/10.1039/C6SC01833E
52. Zhang, C. Y., & Johnson, L. W. (2009). Single quantum-dot-based aptameric nanosensor for cocaine. *Analytical Chemistry*, *81*(8), 3051–3055. https://doi.org/10.1021/ac802737b
53. Zhang, D. W., Zhang, F. T., Cui, Y. R., Deng, Q. P., Krause, S., Zhou, Y. L., & Zhang, X. X. (2012). A label-free aptasensor for the sensitive and specific detection of cocaine using supramolecular aptamer fragments/target complex by electrochemical impedance spectroscopy. *Talanta*, *92*, 65–71. https://doi.org/10.1016/j.talanta.2012.01.049
54. Zhang, H., Jiang, B., Xiang, Y., Zhang, Y., Chai, Y., & Yuan, R. (2011). Aptamer/quantum dot-based simultaneous electrochemical detection of multiple small molecules. *Analytica Chimica Acta*, *688*(2), 99–103. https://doi.org/10.1016/j.aca.2010.12.017
55. Zhang, J., Wang, L., Pan, D., Song, S., Boey, F. Y. C., Zhang, H., & Fan, C. (2008). Visual cocaine detection with gold nanoparticles and rationally engineered aptamer structures. *Small*, *4*(8), 1196–1200. https://doi.org/10.1002/smll.200800057
56. Zhao, T., Liu, R., Ding, X., Zhao, J., Yu, H., Wang, L., … Xiao, Y. (2015). Nanoprobe-Enhanced, Split Aptamer-Based Electrochemical Sandwich Assay for Ultrasensitive Detection of Small Molecules. *Analytical Chemistry*, *87*(15), 7712–7719. https://doi.org/10.1021/acs.analchem.5b01178
57. Zhou, J., Ellis, A. V., Kobus, H., & Voelcker, N. H. (2012). Aptamer sensor for cocaine using minor groove binder based energy transfer. *Analytica Chimica Acta*, *719*, 76–81. https://doi.org/10.1016/j.aca.2012.01.011
58. Zhou, Z., Du, Y., & Dong, S. (2011). Double-strand DNA-templated formation of copper nanoparticles as fluorescent probe for label-free aptamer sensor. *Analytical Chemistry*, *83*(13), 5122–5127. https://doi.org/10.1021/ac200120g
59. Zou, R., Lou, X., Ou, H., Zhang, Y., Wang, W., Yuan, M., … Liu, Y. (2012). Highly specific triple-fragment aptamer for optical detection of cocaine. *RSC Advances*, *2*(11), 4636–4638. https://doi.org/10.1039/c2ra20307c

***S6 RNA anti-theophylline aptamer (the conserved theophylline binding region, Jenison et al., 1994) References***

1. Amontov, S., & Jäschke, A. (2006). Controlling the rate of organic reactions: rational design of allosteric Diels-Alderase ribozymes. *Nucleic acids research*, *34*(18), 5032-5038.
2. An, C. Il, Trinh, V. B., & Yokobayashi, Y. (2006). Artificial control of gene expression in mammalian cells by modulating RNA interference through aptamer-small molecule interaction. *Rna*, *12*(5), 710–716. https://doi.org/10.1261/rna.2299306
3. Anderson, P. C., & Mecozzi, S. (2005). Unusually Short RNA Sequences: Design of a 13-mer RNA that Selectively Binds and Recognizes Theophylline. *Journal of the American Chemical Society*, *127*(15), 5290–5291. https://doi.org/10.1021/ja0432463
4. Ausländer, D., Wieland, M., Ausländer, S., Tigges, M., & Fussenegger, M. (2011). Rational design of a small molecule-responsive intramer controlling transgene expression in mammalian cells. *Nucleic acids research*, *39*(22), e155-e155.

5. Autour, A., Bouhedda, F., Cubi, R., & Ryckelynck, M. (2019). Optimization of fluorogenic RNA-based biosensors using droplet-based microfluidic ultrahigh-throughput screening. *Methods*, *161*, 46-53.
6. Brutscher, B., Simorre, J. P., & Marion, D. (1999). 13C spin relaxation measurements in RNA: sensitivity and resolution improvement using spin-state selective correlation experiments. *Journal of Biomolecular NMR*, *14*(3), 241-252.
7. Bunka, D. H. J., & Stockley, P. G. (2006). Aptamers come of age – at last. *Nature Reviews Microbiology*, *4*(8), 588–596. https://doi.org/10.1038/nrmicro1458
8. Carrasquilla, C., Lau, P. S., Li, Y., & Brennan, J. D. (2012). Stabilizing structure-switching signaling rna aptamers by entrapment in sol–gel derived materials for solid-phase assays. *Journal of the American Chemical Society*, *134*(26), 10998-11005.
9. Chang, A. L., McKeague, M., Liang, J. C., & Smolke, C. D. (2014). Kinetic and equilibrium binding characterization of aptamers to small molecules using a label-free, sensitive, and scalable platform. *Analytical chemistry*, *86*(7), 3273-3278.
10. Chávez, J. L., Lyon, W., Kelley-Loughnane, N., & Stone, M. O. (2010). Theophylline detection using an aptamer and DNA-gold nanoparticle conjugates. *Biosensors and Bioelectronics*, *26*(1), 23–28. https://doi.org/10.1016/j.bios.2010.04.049
11. Chen, X., Guo, Z., Tang, Y., Shen, Y., & Miao, P. (2018). A highly sensitive gold nanoparticle-based electrochemical aptasensor for theophylline detection. *Analytica Chimica Acta*, *999*, 54–59. https://doi.org/10.1016/j.aca.2017.10.039
12. Chen, Z., Liu, Y., He, A., Li, J., Chen, M., Zhan, Y., … Cai, Z. (2016). Theophylline controllable RNAi-based genetic switches regulate expression of lncRNA TINCR and malignant phenotypes in bladder cancer cells. *Scientific Reports*, *6*(1), 1–12. https://doi.org/10.1038/srep30798
13. Chovelon, B., Durand, G., Dausse, E., Toulmé, J. J., Faure, P., Peyrin, E., & Ravelet, C. (2016). ELAKCA: enzyme-linked aptamer kissing complex assay as a small molecule sensing platform. *Analytical chemistry*, *88*(5), 2570-2575.
14. Chushak, Y., & Stone, M. O. (2009). In silico selection of RNA aptamers. *Nucleic Acids Research*, *37*(12), e87–e87. https://doi.org/10.1093/nar/gkp408
15. Cui, L., Chen, Z., Zhu, Z., Lin, X., Chen, X., & Yang, C. J. (2013). Stabilization of ssRNA on graphene oxide surface: an effective way to design highly robust RNA probes. *Analytical chemistry*, *85*(4), 2269-2275.
16. De Silva, C., & Walter, N. G. (2009). Leakage and slow allostery limit performance of single drug-sensing aptazyme molecules based on the hammerhead ribozyme. *Rna*, *15*(1), 76–84. https://doi.org/10.1261/rna.1346609
17. Dery, K. J., Gusti, V., Gaur, S., Shively, J. E., Yen, Y., & Gaur, R. K. (2009). Alternative splicing as a therapeutic target for human diseases. In *Therapeutic Applications of RNAi* (pp. 127-144). Humana Press.
18. Desai, S. K., & Gallivan, J. P. (2004). Genetic screens and selections for small molecules based on a synthetic riboswitch that activates protein translation. *Journal of the American Chemical Society*, *126*(41), 13247-13254.
19. Domin, G., Findeiß, S., Wachsmuth, M., Will, S., Stadler, P. F., & Mörl, M. (2017). Applicability of a computational design approach for synthetic riboswitches. *Nucleic acids research*, *45*(7), 4108-4119.
20. Dong, Z. M., & Zhao, G. C. (2013). A theophylline quartz crystal microbalance biosensor based on recognition of RNA aptamer and amplification of signal. *Analyst*, *138*(8), 2456–2462. https://doi.org/10.1039/c3an36775d
21. Durand, G., Dausse, E., Goux, E., Fiore, E., Peyrin, E., Ravelet, C., & Toulmé, J. J. (2016). A combinatorial approach to the repertoire of RNA kissing motifs; towards multiplex detection by switching hairpin aptamers. *Nucleic acids research*, *44*(9), 4450-4459.
22. Edwards, K. A., & Baeumner, A. J. (2007). Synthesis of a liposome incorporated 1-carboxyalkylxanthine-phospholipid conjugate and its recognition by an RNA aptamer. *Talanta*, *71*(1), 365–372. https://doi.org/10.1016/j.talanta.2006.04.031
23. Ellington, A. D. (1994). RNA Selection: Aptamers achieve the desired recognition. *Current Biology*, *4*(5), 427–429. https://doi.org/10.1016/S0960-9822(00)00093-2
24. Endoh, T., & Sugimoto, N. (2013). Selection of RNAs for Constructing "Lighting-UP" Biomolecular Switches in Response to Specific Small Molecules. *Plos one*, *8*(3), e60222.
25. Endoh, T., Shintani, R., Mie, M., Kobatake, E., Ohtsuki, T., & Sisido, M. (2009). Detection of bioactive small molecules by fluorescent resonance energy transfer (FRET) in RNA-protein conjugates. *Bioconjugate Chemistry*, *20*(12), 2242–2246. https://doi.org/10.1021/bc9002184
26. Espah Borujeni, A., Mishler, D. M., Wang, J., Huso, W., & Salis, H. M. (2016). Automated physics-based design of synthetic riboswitches from diverse RNA aptamers. *Nucleic Acids Research*, *44*(1), 1–13. https://doi.org/10.1093/nar/gkv1289
27. Feng, S., Che, X., Que, L., Chen, C., & Wang, W. (2016). Rapid detection of theophylline using aptamer-based nanopore thin film sensor. In *2016 IEEE SENSORS* (pp. 1–3). https://doi.org/10.1109/ICSENS.2016.7808959
28. Feng, S., Chen, C., Wang, W., & Que, L. (2018). An aptamer nanopore-enabled microsensor for detection of theophylline. *Biosensors and Bioelectronics*, *105*(November 2017), 36–41. https://doi.org/10.1016/j.bios.2018.01.016
29. Ferapontova, E. E., & Gothelf, K. V. (2009). Optimization of the Electrochemical RNA-Aptamer Based Biosensor for Theophylline by Using a Methylene Blue Redox Label. *Electroanalysis*, *21*(11), 1261–1266. https://doi.org/10.1002/elan.200804558
30. Ferapontova, E. E., & Gothelf, K. V. (2009). Effect of Serum on an RNA Aptamer-Based Electrochemical Sensor for Theophylline. *Langmuir*, *25*(8), 4279–4283. https://doi.org/10.1021/la804309j
31. Ferapontova, E. E., Olsen, E. M., & Gothelf, K. V. (2008). An RNA Aptamer-Based Electrochemical Biosensor for Detection of Theophylline in Serum. *Journal of the American Chemical Society*, *130*(13), 4256–4258. https://doi.org/10.1021/ja711326b
32. Fowler, C. C., Brown, E. D., & Li, Y. (2008). A FACS-based approach to engineering artificial riboswitches. *ChemBioChem*, *9*(12), 1906-1911.
33. Freedman, H., Huynh, L. P., Le, L., Cheatham, T. E., Tuszynski, J. A., & Truong, T. N. (2010). Explicitly Solvated Ligand Contribution to Continuum Solvation Models for Binding Free Energies: Selectivity of Theophylline Binding to an RNA Aptamer. *The Journal of Physical Chemistry B*, *114*(6), 2227–2237. https://doi.org/10.1021/jp9059664
34. Fujita, Y., Tanaka, T., Furuta, H., & Ikawa, Y. (2012). Functional roles of a tetraloop/receptor interacting module in a cyclic di-GMP riboswitch. *Journal of bioscience and bioengineering*, *113*(2), 141-145.
35. Gao, Y., & Guo, L. (2013). A sensitive theophylline sensor based on a single walled carbon nanotube-large mesoporous carbon/Nafion/glassy carbon electrode. *Analytical Methods*, *5*(20), 5785–5791. https://doi.org/10.1039/c3ay41236a
36. Gouda, H., Kuntz, I. D., Case, D. A., & Kollman, P. A. (2003). Free energy calculations for theophylline binding to an RNA aptamer: Comparison of MM-PBSA and thermodynamic integration methods. *Biopolymers*, *68*(1), 16–34. https://doi.org/10.1002/bip.10270
37. Groher, F., & Suess, B. (2014). Synthetic riboswitches—a tool comes of age. *Biochimica et Biophysica Acta (BBA)-Gene Regulatory Mechanisms*, *1839*(10), 964-973.
38. Güney, S., & Cebeci, F. (2015). Selective electrochemical sensor for theophylline based on an electrode modified with imprinted sol-gel film immobilized on carbon nanoparticle

layer. *Sensors and Actuators, B: Chemical*, *208*, 307–314. https://doi.org/10.1016/j.snb.2014.10.056
39. Hermann, T., & Patel, D. J. (2000). Adaptive Recognition by Nucleic Acid Aptamers. *Science*, *287*(5454), 820–825. https://doi.org/10.1126/science.287.5454.820
40. Harbaugh, S., Kelley-Loughnane, N., Davidson, M., Narayanan, L., Trott, S., Chushak, Y. G., & Stone, M. O. (2009). FRET-based optical assay for monitoring riboswitch activation. *Biomacromolecules*, *10*(5), 1055-1060.
41. Harvey, I., Garneau, P., & Pelletier, J. (2002). Inhibition of translation by RNA-small molecule interactions. *Rna*, *8*(4), 452-463.
42. Helm, M., Petermeier, M., Ge, B., Fiammengo, R., & Jäschke, A. (2005). Allosterically Activated Diels− Alder Catalysis by a Ribozyme. *Journal of the American Chemical Society*, *127*(30), 10492-10493.
43. Heus, H. A. (1997). RNA aptamers. *Nature structural biology*, *4*(8), 597-600.
44. Lau, J. L., Baksh, M. M., Fiedler, J. D., Brown, S. D., Kussrow, A., Bornhop, D. J., ... & Finn, M. G. (2011). Evolution and protein packaging of small-molecule RNA aptamers. *ACS nano*, *5*(10), 7722-7729.
45. Jenison, R. D., Gill, S. C., Pardi, A., & Polisky, B. (1994). High-resolution molecular discrimination by RNA. *Science*, *263*(5152), 1425-1429.
46. Jiang, H., Ling, K., Tao, X., & Zhang, Q. (2015). Theophylline detection in serum using a self-assembling RNA aptamer-based gold nanoparticle sensor. *Biosensors and Bioelectronics*, *70*, 299–303. https://doi.org/10.1016/j.bios.2015.03.054
47. Jo, J. J., & Shin, J. S. (2009). Construction of intragenic synthetic riboswitches for detection of a small molecule. *Biotechnology letters*, *31*(10), 1577-1581.
48. Jo, J. J., Kim, J. H., & Shin, J. S. (2012). Probing Translation Initiation through Ligand Binding to the 5′ mRNA Coding Region. *ChemBioChem*, *13*(14), 2048-2051.
49. Jose, A. M., Soukup, G. A., & Breaker, R. R. (2001). Cooperative binding of effectors by an allosteric ribozyme. *Nucleic acids research*, *29*(7), 1631-1637.
50. Jucker, F. M., Phillips, R. M., McCallum, S. A., & Pardi, A. (2003). Role of a heterogeneous free state in the formation of a specific RNA - Theophylline complex. *Biochemistry*, *42*(9), 2560–2567. https://doi.org/10.1021/bi027103+
51. Katiyar, N., Selvakumar, L. S., Patra, S., & Thakur, M. S. (2013). Gold nanoparticles based colorimetric aptasensor for theophylline. *Analytical Methods*, *5*(3), 653–659. https://doi.org/10.1039/c2ay26133b
52. Kiga, D., Futamura, Y., Sakamoto, K., & Yokoyama, S. (1998). An RNA aptamer to the xanthine/guanine base with a distinctive mode of purine recognition. *Nucleic Acids Research*, *26*(7), 1755–1760. https://doi.org/10.1093/nar/26.7.1755
53. Kawai, R., Kimoto, M., Ikeda, S., Mitsui, T., Endo, M., Yokoyama, S., & Hirao, I. (2005). Site-specific fluorescent labeling of RNA molecules by specific transcription using unnatural base pairs. *Journal of the American Chemical Society*, *127*(49), 17286-17295.
54. Kertsburg, A., & Soukup, G. A. (2002). A versatile communication module for controlling RNA folding and catalysis. *Nucleic acids research*, *30*(21), 4599-4606.
55. Kim, D. S., Gusti, V., Dery, K. J., & Gaur, R. K. (2008). Ligand-induced sequestering of branchpoint sequence allows conditional control of splicing. *BMC Molecular Biology*, *9*, 1–15. https://doi.org/10.1186/1471-2199-9-23
56. KIM, D. S., GUSTI, V., PILLAI, S. G., & GAUR, R. K. (2005). An artificial riboswitch for controlling pre-mRNA splicing. *Rna*, *11*(11), 1667-1677.
57. Kumar, D., An, C. I., & Yokobayashi, Y. (2009). Conditional RNA interference mediated by allosteric ribozyme. *Journal of the American Chemical Society*, *131*(39), 13906-13907.
58. Lam, B. J., & Joyce, G. F. (2009). Autocatalytic aptazymes enable ligand-dependent exponential amplification of RNA. *Nature biotechnology*, *27*(3), 288-292.
59. Lam, B. J., & Joyce, G. F. (2011). An isothermal system that couples ligand-dependent catalysis to ligand-independent exponential amplification. *Journal of the American Chemical Society*, *133*(9), 3191-3197.
60. Latham, M. P., Zimmermann, G. R., & Pardi, A. (2009). NMR Chemical Exchange as a Probe for Ligand-Binding Kinetics in a Theophylline-Binding RNA Aptamer. *Journal of the American Chemical Society*, *131*(14), 5052–5053. https://doi.org/10.1021/ja900695m
61. Lau, P. S., Coombes, B. K., & Li, Y. (2010). A General Approach to the Construction of Structure-Switching Reporters from RNA Aptamers. *Angewandte Chemie International Edition*, *49*(43), 7938–7942. https://doi.org/10.1002/anie.201002621
62. Lau, P. S., Lai, C. K., & Li, Y. (2013). Quality Control Certification of RNA Aptamer-Based Detection. *ChemBioChem*, *14*(8), 987–992. https://doi.org/10.1002/cbic.201300134
63. Le, T. T., Scott, S., & Cass, A. E. G. (2013). Streptavidin binding bifunctional aptamers and their interaction with low molecular weight ligands. *Analytica Chimica Acta*, *761*, 143–148. https://doi.org/10.1016/j.aca.2012.11.016
64. Lee, S. W., Zhao, L., Pardi, A., & Xia, T. (2010). Ultrafast Dynamics Show That the Theophylline and 3-Methylxanthine Aptamers Employ a Conformational Capture Mechanism for Binding Their Ligands. *Biochemistry*, *49*(13), 2943–2951. https://doi.org/10.1021/bi100106c
65. Li, M., Sato, Y., Nishizawa, S., Seino, T., Nakamura, K., & Teramae, N. (2009). 2-aminopurine-modified abasic-site-containing duplex DNA for highly selective detection of theophylline. *Journal of the American Chemical Society*, *131*(7), 2448–2449. https://doi.org/10.1021/ja8095625
66. Li, X., Song, J., Wang, Y., & Cheng, T. (2013). Cyclically amplified fluorescent detection of theophylline and thiamine pyrophosphate by coupling self-cleaving RNA ribozyme with endonuclease. *Analytica Chimica Acta*, *797*, 95–101. https://doi.org/10.1016/j.aca.2013.08.023
67. Liao, A. M., Pan, W., Benson, J. C., Wong, A. D., Rose, B. J., & Caltagirone, G. T. (2018). A Simple Colorimetric System for Detecting Target Antigens by a Three-Stage Signal Transformation-Amplification Strategy. *Biochemistry*, *57*(34), 5117–5126. https://doi.org/10.1021/acs.biochem.8b00523
68. Ling, K., Jiang, H., Li, Y., Tao, X., Qiu, C., & Li, F.-R. R. (2016). A self-assembling RNA aptamer-based graphene oxide sensor for the turn-on detection of theophylline in serum. *Biosensors and Bioelectronics*, *86*, 8–13. https://doi.org/10.1016/j.bios.2016.06.024
69. Lou, Y. F., Peng, Y. B., Luo, X., Yang, Z., Wang, R., Sun, D., … Cui, L. (2019). A universal aptasensing platform based on cryonase-assisted signal amplification and graphene oxide induced quenching of the fluorescence of labeled nucleic acid probes: application to the detection of theophylline and ATP. *Microchimica Acta*, *186*(8), 1–9. https://doi.org/10.1007/s00604-019-3596-1
70. Luzi, E., Minunni, M., Tombelli, S., & Mascini, M. (2003). New trends in affinity sensing: aptamers for ligand binding. *TrAC Trends in Analytical Chemistry*, *22*(11), 810–818. https://doi.org/10.1016/S0165-9936(03)01208-1
71. Lynch, S. A., Desai, S. K., Sajja, H. K., & Gallivan, J. P. (2007). A High-Throughput Screen for Synthetic Riboswitches Reveals Mechanistic Insights into Their Function. *Chemistry and Biology*, *14*(2), 173–184. https://doi.org/10.1016/j.chembiol.2006.12.008
72. Lynch, S. A., & Gallivan, J. P. (2009). A flow cytometry-based screen for synthetic riboswitches. *Nucleic Acids Research*, *37*(1), 184–192. https://doi.org/10.1093/nar/gkn924
73. Ma, X., Guo, Z., Mao, Z., Tang, Y., & Miao, P. (2018). Colorimetric theophylline aggregation assay using an RNA

aptamer and non-crosslinking gold nanoparticles. *Microchimica Acta*, *185*(1). https://doi.org/10.1007/s00604-017-2606-4
74. Mansouri Majd, S., Teymourian, H., Salimi, A., & Hallaj, R. (2013). Fabrication of electrochemical theophylline sensor based onmanganese oxide nanoparticles/ionic liquid/chitosan nanocomposite modified glassy carbon electrode. *Electrochimica Acta*, *108*, 707–716. https://doi.org/10.1016/j.electacta.2013.07.029
75. Michener, J. K., & Smolke, C. D. (2012). High-throughput enzyme evolution in Saccharomyces cerevisiae using a synthetic RNA switch. *Metabolic Engineering*, *14*(4), 306–316. https://doi.org/10.1016/j.ymben.2012.04.004
76. Nishizawa, S., Sato, Y., Xu, Z., Morita, K., Li, M., & Teramae, N. (2010). Abasic site-based DNA aptamers for analytical applications. *Supramolecular Chemistry*, *22*(7), 467–476. https://doi.org/10.1080/10610278.2010.484865
77. Nomura, Y., Zhou, L., Miu, A., & Yokobayashi, Y. (2013). Controlling mammalian gene expression by allosteric hepatitis delta virus ribozymes. *ACS Synthetic Biology*, *2*(12), 684–689. https://doi.org/10.1021/sb400037a
78. Ogawa, A. (2011). Rational design of artificial riboswitches based on ligand-dependent modulation of internal ribosome entry in wheat germ extract and their applications as label-free biosensors. *Rna*, *17*(3), 478–488. https://doi.org/10.1261/rna.2433111
79. Ogawa, A. (2012). Rational construction of eukaryotic OFF-riboswitches that downregulate internal ribosome entry site-mediated translation in response to their ligands. *Bioorganic and Medicinal Chemistry Letters*, *22*(4), 1639–1642. https://doi.org/10.1016/j.bmcl.2011.12.118
80. Ogawa, A., & Maeda, M. (2007). Aptazyme-based riboswitches as label-free and detector-free sensors for cofactors. *Bioorganic and Medicinal Chemistry Letters*, *17*(11), 3156–3160. https://doi.org/10.1016/j.bmcl.2007.03.033
81. Ogawa, A., Masuoka, H., & Ota, T. (2017). Artificial OFF-Riboswitches That Downregulate Internal Ribosome Entry without Hybridization Switches in a Eukaryotic Cell-Free Translation System. *ACS Synthetic Biology*, *6*(9), 1656–1662. https://doi.org/10.1021/acssynbio.7b00124
82. Ogawa, A., Murashige, Y., & Takahashi, H. (2018). Canonical translation-modulating OFF-riboswitches with a single aptamer binding to a small molecule that function in a higher eukaryotic cell-free expression system. *Bioorganic and Medicinal Chemistry Letters*, *28*(14), 2353–2357. https://doi.org/10.1016/j.bmcl.2018.06.041
83. Ogawa, A., & Tabuchi, J. (2015). Biofunction-assisted aptasensors based on ligand-dependent 3′ processing of a suppressor tRNA in a wheat germ extract. *Organic and Biomolecular Chemistry*, *13*(24), 6681–6685. https://doi.org/10.1039/c5ob00794a
84. Page, K., Shaffer, J., Lin, S., Zhang, M., & Liu, J. M. (2018). Engineering Riboswitches in Vivo Using Dual Genetic Selection and Fluorescence-Activated Cell Sorting. *ACS Synthetic Biology*, *7*(9), 2000–2006. https://doi.org/10.1021/acssynbio.8b00099
85. Patel, D. J., & Suri, A. K. (2000). Structure, recognition and discrimination in RNA aptamer complexes with cofactors, amino acids, drugs and aminoglycoside antibiotics. *Reviews in Molecular Biotechnology*, *74*(1), 39–60. https://doi.org/10.1016/s1389-0352(99)00003-3
86. Pei, R., & Stojanovic, M. N. (2008). Study of thiazole orange in aptamer-based dye-displacement assays. *Analytical and Bioanalytical Chemistry*, *390*(4), 1093–1099. https://doi.org/10.1007/s00216-007-1773-2
87. Petermeier, M., & Jäschke, A. (2009). New theophylline-activated Diels-Alderase ribozymes by molecular engineering. *Organic and Biomolecular Chemistry*, *7*(2), 288–292. https://doi.org/10.1039/b816726e
88. Qi, L., Lucks, J. B., Liu, C. C., Mutalik, V. K., & Arkin, A. P. (2012). Engineering naturally occurring trans-acting non-coding RNAs to sense molecular signals. *Nucleic Acids Research*, *40*(12), 5775–5786. https://doi.org/10.1093/nar/gks168
89. Rankin, C. J., Fuller, E. N., Hamor, K. H., Gabarra, S. A., & Shields, T. P. (2006). A Simple Fluorescent Biosensor for Theophylline Based on its RNA Aptamer. *Nucleosides, Nucleotides & Nucleic Acids*, *25*(12), 1407–1424. https://doi.org/10.1080/15257770600919084
90. Robertson, M. P., & Ellington, A. D. (2000). Design and optimization of effector-activated ribozyme ligases. *Nucleic Acids Research*, *28*(8), 1751–1759. https://doi.org/10.1093/nar/28.8.1751
91. Rudolph, M. M., Vockenhuber, M. P., & Suess, B. (2013). Synthetic riboswitches for the conditional control of gene expression in Streptomyces coelicolor. *Microbiology (United Kingdom)*, *159*(PART7), 1416–1422. https://doi.org/10.1099/mic.0.067322-0
92. Sarpong, K., & Datta, B. (2012). Nucleic-acid-binding chromophores as efficient indicators of aptamer-target interactions. *Journal of Nucleic Acids*, *2012*. https://doi.org/10.1155/2012/247280
93. Sato, Y., Zhang, Y., Nishizawa, S., Seino, T., Nakamura, K., Li, M., & Teramae, N. (2012). Competitive assay for theophylline based on an abasic site-containing DNA duplex aptamer and a fluorescent ligand. *Chemistry - A European Journal*, *18*(40), 12719–12724. https://doi.org/10.1002/chem.201201298
94. Sekella, P. T., Rueda, D., & Walter, N. G. (2002). A biosensor for theophylline based on fluorescence detection of ligand-induced hammerhead ribozyme cleavage. *RNA*, *8*(10), 1242–1252. https://doi.org/10.1017/S1355838202028066
95. Sharma, V., Nomura, Y., & Yokobayashi, Y. (2008). Engineering complex riboswitch regulation by dual genetic selection. *Journal of the American Chemical Society*, *130*(48), 16310–16315. https://doi.org/10.1021/ja805203w
96. Sibille, N., Pardi, A., Simorre, J., & Blackledge, M. (2001). Refinement of Local and Long-Range Structural Order in. *Journal of American Chemical Society*, *123*(9), 12135–12146.
97. Song, J., Lau, P. S., Liu, M., Shuang, S., Dong, C., & Li, Y. (2014). A general strategy to create RNA aptamer sensors using regulated graphene oxide adsorption. *ACS Applied Materials and Interfaces*, *6*(24), 21806–21812. https://doi.org/10.1021/am502138n
98. Soukup, G. A., & Breaker, R. R. (1999). Engineering precision RNA molecular switches. *Proceedings of the National Academy of Sciences of the United States of America*, *96*(7), 3584–3589. https://doi.org/10.1073/pnas.96.7.3584
99. Soukup, G. A., Emilsson, G. A. M., & Breaker, R. R. (2000). Altering molecular recognition of RNA aptamers by allosteric selection. *Journal of Molecular Biology*, *298*(4), 623–632. https://doi.org/10.1006/jmbi.2000.3704
100. Suess, B., Fink, B., Berens, C., Stentz, R., & Hillen, W. (2004). A theophylline responsive riboswitch based on helix slipping controls gene expression in vivo. *Nucleic Acids Research*, *32*(4), 1610–1614. https://doi.org/10.1093/nar/gkh321
101. Tang, J., & Breaker, R. R. (1997). Rational design of allosteric ribozymes. *Chemistry and Biology*, *4*(6), 453–459. https://doi.org/10.1016/S1074-5521(97)90197-6
102. Thompson, K. M., Syrett, H. A., Knudsen, S. M., & Ellington, A. D. (2002). Group I aptazymes as genetic regulatory switches. *BMC biotechnology*, *2*(1), 1-12.
103. Thompson, K. M., Syrett, H. A., Knudsen, S. M., & Ellington, A. D. (2002). Group I aptazymes as genetic regulatory switches. *BMC biotechnology*, *2*(1), 1-12.
104. Tanida, Y., Ito, M., & Fujitani, H. (2007). Calculation of absolute free energy of binding for theophylline and its analogs to RNA aptamer using nonequilibrium work values. *Chemical Physics*, *337*(1), 135–143. https://doi.org/10.1016/j.chemphys.2007.07.014
105. Topp, S., & Gallivan, J. P. (2008). Random walks to synthetic riboswitches - A high-throughput selection based on cell

motility. *ChemBioChem*, *9*(2), 210–213. https://doi.org/10.1002/cbic.200700546
106. Topp, S., & Gallivan, J. P. (2007). Guiding bacteria with small molecules and RNA. *Journal of the American Chemical Society*, *129*(21), 6807–6811. https://doi.org/10.1021/ja0692480
107. Topp, S., & Gallivan, J. P. (2008). Riboswitches in unexpected places - A synthetic riboswitch in a protein coding region. *Rna*, *14*(12), 2498–2503. https://doi.org/10.1261/rna.1269008
108. Townshend, B., Kennedy, A. B., Xiang, J. S., & Smolke, C. D. (2015). High-throughput cellular RNA device engineering. *Nature Methods*, *12*(10), 989–994. https://doi.org/10.1038/nmeth.3486
109. Tuleuova, N., An, C. Il, Ramanculov, E., Revzin, A., & Yokobayashi, Y. (2008). Modulating endogenous gene expression of mammalian cells via RNA-small molecule interaction. *Biochemical and Biophysical Research Communications*, *376*(1), 169–173. https://doi.org/10.1016/j.bbrc.2008.08.112
110. Wachsmuth, M., Domin, G., Lorenz, R., Serfling, R., Findeiß, S., Stadler, P. F., & Mörl, M. (2015). Design criteria for synthetic riboswitches acting on transcription. *RNA Biology*, *12*(2), 221–231. https://doi.org/10.1080/15476286.2015.1017235
111. Wachsmuth, M., Findeiß, S., Weissheimer, N., Stadler, P. F., & Mörl, M. (2013). De novo design of a synthetic riboswitch that regulates transcription termination. *Nucleic Acids Research*, *41*(4), 2541–2551. https://doi.org/10.1093/nar/gks1330
112. Wang, J., Cheng, W., Meng, F., Yang, M., Pan, Y., & Miao, P. (2018). Hand-in-hand RNA nanowire-based aptasensor for the detection of theophylline. *Biosensors and Bioelectronics*, *101*(June 2017), 153–158. https://doi.org/10.1016/j.bios.2017.10.025
113. Wang, S., Mortazavi, L., & White, K. A. (2008). Higher-order RNA structural requirements and small-molecule induction of tombusvirus subgenomic mRNA transcription. *Journal of virology*, *82*(8), 3864-3871.
114. Wang, S., & White, K. A. (2007). Riboswitching on RNA virus replication. *Proceedings of the National Academy of Sciences*, *104*(25), 10406-10411.
115. Warfield, B. M., & Anderson, P. C. (2017). Molecular simulations and Markov state modeling reveal the structural diversity and dynamics of a theophylline-binding RNA aptamer in its unbound state. *PLoS ONE*, *12*(4). https://doi.org/10.1371/journal.pone.0176229
116. Wieland, M., & Hartig, J. S. (2008). Improved aptazyme design and in vivo screening enable riboswitching in bacteria. *Angewandte Chemie - International Edition*, *47*(14), 2604–2607. https://doi.org/10.1002/anie.200703700
117. Wu, J. F., Gao, X., Ge, L., Zhao, G. C., & Wang, G. F. (2019). A fluorescence sensing platform of theophylline based on the interaction of RNA aptamer with graphene oxide. *RSC Advances*, *9*(34), 19813–19818. https://doi.org/10.1039/c9ra02475a
118. Xie, S., & Walton, S. P. (2009). Application and analysis of structure-switching aptamers for small molecule quantification. *Analytica Chimica Acta*, *638*(2), 213–219. https://doi.org/10.1016/j.aca.2009.02.018
119. Zhang, Y., Wang, J., Cheng, H., Sun, Y., Liu, M., Wu, Z., & Pei, R. (2017). Conditional control of suicide gene expression in tumor cells with theophylline-responsive ribozyme. *Gene Therapy*, *24*(2), 84–91. https://doi.org/10.1038/gt.2016.78
120. Zhao, G. C., & Yang, X. (2010). A label-free electrochemical RNA aptamer for selective detection of theophylline. *Electrochemistry Communications*, *12*(2), 300–302. https://doi.org/10.1016/j.elecom.2009.12.022
121. Zhou, Q., Xia, X., Luo, Z., Liang, H., & Shakhnovich, E. (2015). Searching the Sequence Space for Potent Aptamers Using SELEX in Silico. *Journal of Chemical Theory and Computation*, *11*(12), 5939–5946. https://doi.org/10.1021/acs.jctc.5b00707
122. Zimmermann, G. R., Jenison, R. D., Wick, C. L., Simorre, J., & Pardi, A. (1997). Interlocking structural motifs mediate molecular discrimination by a theophylline-binding RNA. *Nature Structural Biology*, *4*(8), 644–649.
123. Zimmermann, G. R., Shields, T. P., Jenison, R. D., Wick, C. L., & Pardi, A. (1998). A Semiconserved Residue Inhibits Complex Formation by Stabilizing Interactions in the Free State of a Theophylline-Binding RNA. *Biochemistry*, *37*(25), 9186–9192. https://doi.org/10.1021/bi980082s
124. Zimmermann, G. R., Wick, C. L., Shields, T. P., Jenison, R. D., & Pardi, A. (2000). Molecular interactions and metal binding in the theophylline-binding core of an RNA aptamer. *RNA*, *6*(5), 659–667. https://doi.org/10.1017/S1355838200000169

***S7: 80mer DNA anti-lysozyme aptamer (Apta1, Tran et al., 2010) References***

1. Alves, R. C., Barroso, M. F., González-García, M. B., Oliveira, M. B. P. P., & Delerue-Matos, C. (2016). New Trends in Food Allergens Detection: Toward Biosensing Strategies. *Critical Reviews in Food Science and Nutrition*, *56*(14), 2304–2319. https://doi.org/10.1080/10408398.2013.831026
2. Amaya-González, S., López-López, L., Miranda-Castro, R., de-los-Santos-Álvarez, N., Miranda-Ordieres, A. J., & Lobo-Castañón, M. J. (2015). Affinity of aptamers binding 33-mer gliadin peptide and gluten proteins: INFLUENCE of immobilization and labeling tags. *Analytica Chimica Acta*, *873*, 63–70. https://doi.org/10.1016/j.aca.2015.02.053
3. Ardekani, L. S., Moghadam, T. T., Thulstrup, P. W., & Ranjbar, B. (2019). Design and Fabrication of a Silver Nanocluster-Based Aptasensor for Lysozyme Detection. *Plasmonics*, *14*(6), 1765–1774. https://doi.org/10.1007/s11468-019-00954-5
4. Atalay, Y. T., Vermeir, S., Witters, D., Vergauwe, N., Verbruggen, B., Verboven, P., … Lammertyn, J. (2011). Microfluidic analytical systems for food analysis. *Trends in Food Science and Technology*, *22*(7), 386–404. https://doi.org/10.1016/j.tifs.2011.05.001
5. Boushell, V., Pang, S., & He, L. (2017). Aptamer-Based SERS Detection of Lysozyme on a Food-Handling Surface. *Journal of Food Science*, *82*(1), 225–231. https://doi.org/10.1111/1750-3841.13582
6. Carstens, C., Deckwart, M., Webber-Witt, M., Schäfer, V., Eichhorn, L., Brockow, K., … Paschke-Kratzin, A. (2014). Evaluation of the efficiency of enological procedures on lysozyme depletion in wine by an indirect ELISA method. *Journal of Agricultural and Food Chemistry*, *62*(26), 6247–6253. https://doi.org/10.1021/jf405319j
7. Chen, L. (2011). *Application of High Throughput Sequencing in Selection of RNA Aptamers*. Graduate School of Syracuse University.
8. Chumphukam, O., Le, T. T., Piletsky, S., & Cass, A. E. G. (2015). Generation of a pair of independently binding DNA aptamers in a single round of selection using proximity ligation. *Chemical Communications*, *51*(43), 9050-9053.
9. de Jong, S. (2013). *Methods of Optimization for KCE-based Aptamer Selection*. York University.
10. Dong, Y., Xu, Y., Yong, W., Chu, X., & Wang, D. (2014). Aptamer and Its Potential Applications for Food Safety. *Critical Reviews in Food Science and Nutrition*, *54*(12), 1548–1561. https://doi.org/10.1080/10408398.2011.642905
11. Durney, B. C., Crihfield, C. L., & Holland, L. A. (2017). Capillary electrophoresis applied to DNA: Determining and harnessing sequence and structure to advance bioanalyses (2009-2014). *Analytical and Bioanalytical Chemistry*, *407*(23), 6923–6938. https://doi.org/10.1007/s00216-015-8703-5

12. Han, B., Zhao, C., Yin, J., & Wang, H. (2012). High performance aptamer affinity chromatography for single-step selective extraction and screening of basic protein lysozyme. *Journal of Chromatography B: Analytical Technologies in the Biomedical and Life Sciences*, *903*, 112–117. https://doi.org/10.1016/j.jchromb.2012.07.003
13. Hünniger, T., Wessels, H., Fischer, C., Paschke-Kratzin, A., & Fischer, M. (2014). Just in time -selection: A rapid semiautomated SELEX of DNA aptamers using magnetic separation and BEAMing. *Analytical Chemistry*, *86*(21), 10940–10947. https://doi.org/10.1021/ac503261b
14. Kaur, H., Bhagwat, S. R., Sharma, T. K., & Kumar, A. (2019). Analytical techniques for characterization of biological molecules - Proteins and aptamers/oligonucleotides. *Bioanalysis*, *11*(2), 103–117. https://doi.org/10.4155/bio-2018-0225
15. Khedri, M., Ramezani, M., Rafatpanah, H., & Abnous, K. (2018). Detection of food-born allergens with aptamer-based biosensors. *TrAC - Trends in Analytical Chemistry*, *103*, 126–136. https://doi.org/10.1016/j.trac.2018.04.001
16. Lam, R. (2018). Novel Fingermark Detection Methods Using Biomolecular Recognition by. *Dissertation*, (February).
17. Leblebici, P., Leirs, K., Spasic, D., & Lammertyn, J. (2019). Encoded particle microfluidic platform for rapid multiplexed screening and characterization of aptamers against influenza A nucleoprotein. *Analytica Chimica Acta*, *1053*, 70–80. https://doi.org/10.1016/j.aca.2018.11.055
18. Lee, M., Urata, S. M., Aguilera, J. A., Perry, C. C., & Milligan, J. R. (2012). Modeling the Influence of Histone Proteins on the Sensitivity of DNA to Ionizing Radiation. *Radiation Research*, *177*(2), 152–163. https://doi.org/10.1667/rr2812.1
19. Liu, J., You, M., Pu, Y., Liu, H., Ye, M., & Tan, W. (2011). Recent Developments in Protein and Cell-Targeted Aptamer Selection and Applications. *Current Medicinal Chemistry*, *18*(27), 4117–4125.
20. Liu, X., Hu, R., Gao, Z., & Shao, N. (2015). Photoluminescence mechanism of DNA-templated silver nanoclusters: Coupling between surface plasmon and emitter and sensing of lysozyme. *Langmuir*, *31*(21), 5859–5867. https://doi.org/10.1021/acs.langmuir.5b00589
21. Luu, T., Liu, M., Chen, Y., Hushiarian, R., Cass, A., Tang, B. Z., & Hong, Y. (2019). Aptamer-Based Biosensing with a Cationic AIEgen. *Australian Journal of Chemistry*, *72*(8), 620–626. https://doi.org/10.1071/CH19238
22. Meirinho, S. G., Dias, L. G., Peres, A. M., & Rodrigues, L. R. (2017). Electrochemical aptasensor for human osteopontin detection using a DNA aptamer selected by SELEX. *Analytica Chimica Acta*, *987*, 25–37. https://doi.org/10.1016/j.aca.2017.07.071
23. Meirinho, S. G., Dias, L. G., Peres, A. M., & Rodrigues, L. R. (2015). Development of an electrochemical RNA-aptasensor to detect human osteopontin. *Biosensors and Bioelectronics*, *71*, 332–341. https://doi.org/10.1016/j.bios.2015.04.050
24. Meyer, M. (2014). Applications of Aptamers in Flow Cytometry Assays. *Dissertation*, (September 1985).
25. Mihai, I., Vezeanu, A., Polonschii, C., David, S., Gaspar, S., Bucar, B., … Vasilescu, A. (2012). Low-fouling SPR detection of lysozyme and aggregates. *Analytical Methods*, *0*(1–3), 323–350. https://doi.org/10.1039/x0xx00000x
26. Mishra, R. K., Hayat, A., Mishra, G. K., Catanante, G., Sharma, V., & Marty, J. L. (2017). A novel colorimetric competitive aptamer assay for lysozyme detection based on superparamagnetic nanobeads. *Talanta*, *165*(December 2016), 436–441. https://doi.org/10.1016/j.talanta.2016.12.083
27. Musumeci, D., Platella, C., Riccardi, C., Moccia, F., & Montesarchio, D. (2017). Fluorescence sensing using DNA Aptamers in cancer research and clinical diagnostics. *Cancers*, *9*(12), 1–43. https://doi.org/10.3390/cancers9120174
28. Ocaña, C., Hayat, A., Mishra, R. K., Vasilescu, A., del Valle, M., & Marty, J. L. (2015). Label free aptasensor for Lysozyme detection: A comparison of the analytical performance of two aptamers. *Bioelectrochemistry*, *105*(October), 72–77. https://doi.org/10.1016/j.bioelechem.2015.05.009
29. Ocaña, C., Hayat, A., Mishra, R., Vasilescu, A., Del Valle, M., & Marty, J. L. (2015). A novel electrochemical aptamer-antibody sandwich assay for lysozyme detection. *Analyst*, *140*(12), 4148–4153. https://doi.org/10.1039/c5an00243e
30. Ogihara, K., Savory, N., Abe, K., Yoshida, W., Arakawa, M., Asahi, M., … Ikebukuro, K. (2015). Inhibition of an Allergen-Antibody Reaction Related to Japanese Cedar Pollinosis Using DNA Aptamers Against the Cry j 2 Allergen. *Nucleic Acid Therapeutics*, *25*(6), 311–316. https://doi.org/10.1089/nat.2015.0539
31. Prado, M., Ortea, I., Vial, S., Rivas, J., Calo-Mata, P., & Barros-Velázquez, J. (2016). Advanced DNA- and protein-based methods for the detection and investigation of food allergens. *Critical Reviews in Food Science and Nutrition*, *56*(15), 2511–2542. https://doi.org/10.1080/10408398.2013.873767
32. Rose, C. M., Hayes, M. J., Stettler, G. R., Hickey, S. F., Axelrod, T. M., Giustini, N. P., & Suljak, S. W. (2010). Capillary Electrophoretic Development of Aptamers for a Glycosylated VEGF Peptide Fragment. *Analyst*, *135*(11), 2945–2951. https://doi.org/10.1039/c0an00445f
33. Tabarzad, M., & Jafari, M. (2016). Trends in the Design and Development of Specific Aptamers Against Peptides and Proteins. *Protein Journal*, *35*(2), 81–99. https://doi.org/10.1007/s10930-016-9653-2
34. Tian, R. Y., Lin, C., Yu, S. Y., Gong, S., Hu, P., Li, Y. S., … Lu, S. Y. (2016). Preparation of a specific ssDNA aptamer for brevetoxin-2 using SELEX. *Journal of Analytical Methods in Chemistry*, *2016*. https://doi.org/10.1155/2016/9241860
35. Titoiu, A. M., Porumb, R., Fanjul-Bolado, P., Epure, P., Zamfir, M., & Vasilescu, A. (2019). Detection of Allergenic Lysozyme during Winemaking with an Electrochemical Aptasensor. *Electroanalysis*, *31*(11), 2262–2273. https://doi.org/10.1002/elan.201900333
36. Tran, D. T., Janssen, K. P. F., Pollet, J., Lammertyn, E., Anné, J., Van Schepdael, A., & Lammertyn, J. (2010). Selection and characterization of DNA aptamers for egg white lysozyme. *Molecules*, *15*(3), 1127–1140. https://doi.org/10.3390/molecules15031127
37. Tran, D. T., Knez, K., Janssen, K. P., Pollet, J., Spasic, D., & Lammertyn, J. (2013). Selection of aptamers against Ara h 1 protein for FO-SPR biosensing of peanut allergens in food matrices. *Biosensors and Bioelectronics*, *43*(1), 245–251. https://doi.org/10.1016/j.bios.2012.12.022
38. Truong, P. L., Choi, S. P., & Sim, S. J. (2013). Amplification of resonant rayleigh light scattering response using immunogold colloids for detection of lysozyme. *Small*, *9*(20), 3485–3492. https://doi.org/10.1002/smll.201202638
39. Vasilescu, A., Gaspar, S., Mihai, I., Tache, A., & Litescu, S. C. (2013). Development of a label-free aptasensor for monitoring the self-association of lysozyme. *Analyst*, *138*(12), 3530–3537. https://doi.org/10.1039/c3an00229b
40. Vasilescu, A., Vezeanu, A., & Badea, M. (2014). Electrochemical impedance spectroscopy investigations focused on food allergens. *Sensing in Electroanalysis.(K. …*, *8*, 59–83. Retrieved from http://dspace.upce.cz/handle/10195/58389
41. Vasilescu, A., Wang, Q., Li, M., Boukherroub, R., & Szunerits, S. (2016). Aptamer-based electrochemical sensing of lysozyme. *Chemosensors*, *4*(2), 1–20. https://doi.org/10.3390/chemosensors4020010
42. Wang, P. (2018). *DNA Enzymes for Tyrosine PEGylation and Azido-adenylation of peptide and protein substrates*. University of Illinois at Urbana-Chapaign.
43. Wang, X., Xu, Y., Chen, Y., Li, L., Liu, F., & Li, N. (2011). The gold-nanoparticle-based surface plasmon resonance light scattering and visual DNA aptasensor for lysozyme. *Analytical and Bioanalytical Chemistry*, *400*(7), 2085–2091. https://doi.org/10.1007/s00216-011-4943-1

44. Wang, Y., Pu, K. Y., & Liu, B. (2010). Anionic conjugated polymer with aptamer-functionalized silica nanoparticle for label-free naked-eye detection of lysozyme in protein mixtures. *Langmuir*, *26*(12), 10025–10030. https://doi.org/10.1021/la100139p
45. Weng, X., & Neethirajan, S. (2016). A Nanomaterial-enhanced Microfluidic Biosensor For Ara h 1 Detection. *CSBE/SCGAB 2016 Annual Conference*, (July), 3–12.
46. Weng, X., & Neethirajan, S. (2018). Paper-based microfluidic aptasensor for food safety. *Journal of Food Safety*, *38*(1), 1–8. https://doi.org/10.1111/jfs.12412
47. Wood, M. (2014). *A novel approach to latent fingermark detection using aptamer-based reagents*. *Dissertation*. University of Technology, Syndey. Retrieved from https://opus.lib.uts.edu.au/bitstream/10453/24200/2/02whole.pdf
48. Wood, M., Maynard, P., Spindler, X., Lennard, C., & Roux, C. (2012). Visualization of latent fingermarks using an aptamer-based reagent. *Angewandte Chemie - International Edition*, *51*(49), 12272–12274. https://doi.org/10.1002/anie.201207394
49. Wu, K., Ma, C., Zhao, H., He, H., & Chen, H. (2018). Label-free G-quadruplex aptamer fluorescence assay for Ochratoxin A using a thioflavin T probe. *Toxins*, *10*(5). https://doi.org/10.3390/toxins10050198
50. Xie, S., Qiu, L., Cui, L., Liu, H., Sun, Y., Liang, H., … Tan, W. (2017). Reversible and Quantitative Photoregulation of Target Proteins. *Chem*, *3*(6), 1021–1035. https://doi.org/10.1016/j.chempr.2017.11.008
51. Zhang, X. X., Zhu, S., Xiong, Y., Deng, C., & Zhang, X. X. (2013). Development of a MALDI-TOF MS strategy for the high-throughput analysis of biomarkers: On-target aptamer immobilization and laser-accelerated proteolysis. *Angewandte Chemie - International Edition*, *52*(23), 6055–6058. https://doi.org/10.1002/anie.201300566
52. ZHOU, Y. Y., DU, Y. M., BIAN, X. J., & YAN, J. (2019). Preparation of Aptamer-functionalized Au@pNTP@SiO2 Core-Shell Surface-enhanced Raman Scattering Probes for Raman Imaging Study of Adhesive Tape Transferred-Latent Fingerprints. *Chinese Journal of Analytical Chemistry*, *47*(7), 998–1005. https://doi.org/10.1016/S1872-2040(19)61171-0
53. Zhu, C., Li, L., Yang, G., Fang, S., Liu, M., Ghulam, M., … Qu, F. (2019). Online reaction based single-step capillary electrophoresis-systematic evolution of ligands by exponential enrichment for ssDNA aptamers selection. *Analytica Chimica Acta*, *1070*, 112–122. https://doi.org/10.1016/j.aca.2019.04.034
54. Zhu, C., Yang, G., Ghulam, M., Li, L., & Qu, F. (2019). Evolution of multi-functional capillary electrophoresis for high-efficiency selection of aptamers. *Biotechnology Advances*, *37*(8), 107432. https://doi.org/10.1016/j.biotechadv.2019.107432
55. Zou, M., Chen, Y., Xu, X., Huang, H., Liu, F., & Li, N. (2012). The homogeneous fluorescence anisotropic sensing of salivary lysozyme using the 6-carboxyfluorescein-labeled DNA aptamer. *Biosensors and Bioelectronics*, *32*(1), 148–154. https://doi.org/10.1016/j.bios.2011.11.05

***S8: 26mer DNA anti-nucleolin aptamer (AS1411/AGRO100, Bates et al., 1999) References***

1. Ai, J., Xu, Y., Lou, B., Li, D., & Wang, E. (2014). Multifunctional AS1411-functionalized fluorescent gold nanoparticles for targeted cancer cell imaging and efficient photodynamic therapy. *Talanta*, *118*, 54–60. https://doi.org/10.1016/j.talanta.2013.09.062
2. Alibolandi, M., Abnous, K., Ramezani, M., Hosseinkhani, H., & Hadizadeh, F. (2014). Synthesis of AS1411-aptamer-conjugated CdTe quantum dots with high fluorescence strength for probe labeling tumor cells. *Journal of Fluorescence*, *24*(5), 1519–1529. https://doi.org/10.1007/s10895-014-1437-5
3. Aravind, A., Jeyamohan, P., Nair, R., & Veeranarayanan, S. (2012). AS1411 Aptamer Tagged PLGA-Lecithin-PEG Nanoparticles for Tumor Cell Targeting and Drug Delivery, *109*(11), 2920–2931. https://doi.org/10.1002/bit.24558/abstract
4. Balinsky, C. A., Schmeisser, H., Ganesan, S., Singh, K., Pierson, T. C., & Zoon, K. C. (2013). Nucleolin Interacts with the Dengue Virus Capsid Protein and Plays a Role in Formation of Infectious Virus Particles. *Journal of Virology*, *87*(24), 13094–13106. https://doi.org/10.1128/jvi.00704-13
5. Bates, P. J., Laber, D. a, Miller, D. M., Thomas, S. D., & Trent, J. O. (2010). Discovery and Development of the G-rich Oligonucleotide AS1411 as a Novel Treatment for Cancer. *Experimental and Molecular Pathology*, *86*(3), 151–164. https://doi.org/10.1016/j.yexmp.2009.01.004.Discovery
6. Bates, P. J., Kahlon, J. B., Thomas, S. D., Trent, J. O., & Miller, D. M. (1999). Antiproliferative activity of G-rich oligonucleotides correlates with protein binding. *Journal of Biological Chemistry*, *274*(37), 26369–26377. https://doi.org/10.1074/jbc.274.37.26369
7. Bates, P. J., Reyes-Reyes, E. M., Malik, M. T., Murphy, E. M., O'Toole, M. G., & Trent, J. O. (2017). G-quadruplex oligonucleotide AS1411 as a cancer-targeting agent: Uses and mechanisms. *Biochimica et Biophysica Acta - General Subjects*, *1861*(5), 1414–1428. https://doi.org/10.1016/j.bbagen.2016.12.015
8. Belyaeva, T., Nicol, C., Cesur, Ö., Travé, G., Blair, G., & Stonehouse, N. (2014). An RNA Aptamer Targets the PDZ-Binding Motif of the HPV16 E6 Oncoprotein. *Cancers*, *6*(3), 1553–1569. https://doi.org/10.3390/cancers6031553
9. Bunka, D. H. J., Platonova, O., & Stockley, P. G. (2010). Development of aptamer therapeutics. *Current Opinion in Pharmacology*, *10*(5), 557–562. https://doi.org/10.1016/j.coph.2010.06.009
10. Cao, Z., Tong, R., Mishra, A., Xu, W., Wong, G. C. L., Cheng, J., & Lu, Y. (2009). Reversible cell-specific drug delivery with aptamer-functionalized liposomes. *Angewandte Chemie - International Edition*, *48*(35), 6494–6498. https://doi.org/10.1002/anie.200901452
11. Carvalho, J., Paiva, A., Cabral Campello, M. P., Paulo, A., Mergny, J. L., Salgado, G. F., … Cruz, C. (2019). Aptamer-based Targeted Delivery of a G-quadruplex Ligand in Cervical Cancer Cells. *Scientific Reports*, *9*(1), 1–12. https://doi.org/10.1038/s41598-019-44388-9
12. Cerchia, L., & de Franciscis, V. (2010). Targeting cancer cells with nucleic acid aptamers. *Trends in Biotechnology*, *28*(10), 517–525. https://doi.org/10.1016/j.tibtech.2010.07.005
13. Charoenphol, P., & Bermudez, H. (2014). Aptamer-targeted DNA nanostructures for therapeutic delivery. *Molecular Pharmaceutics*, *11*(5), 1721–1725. https://doi.org/10.1021/mp500047b
14. Chauhan, R., Allen, N. C., Keynton, R. S., Bates, P. J., Malik, M. T., & Toole, M. G. O. (2019). Engineering sequence and stimuli dependent doxorubicin release from anti-nucleolin aptamer coated gold nanoparticles. *2018 IEEE International Symposium on Signal Processing and Information Technology, ISSPIT 2018*, 114–117. https://doi.org/10.1109/ISSPIT.2018.8642659
15. Cheng, Y., Zhao, G., Zhang, S., Nigim, F., Zhou, G., Yu, Z., … Li, Y. (2016). AS1411-induced growth inhibition of glioma cells by up-regulation of p53 and down-regulation of Bcl-2 and Akt1 via nucleolin. *PLoS ONE*, *11*(12), 1–21. https://doi.org/10.1371/journal.pone.0167094
16. Cho, Y., Lee, Y. Bin, Lee, J. H., Lee, D. H., Cho, E. J., Yu, S. J., … Yoon, J. H. (2016). Modified AS1411 Aptamer Suppresses Hepatocellular Carcinoma by Up-Regulating Galectin-14. *PLoS ONE*, *11*(8), 1–14. https://doi.org/10.1371/journal.pone.0160822
17. Choi, J. H., Chen, K. H., Han, J. H., Chaffee, A. M., & Strano, M. S. (2009). DNA aptamer-passivated nanocrystal synthesis: A facile approach for nanoparticle-based cancer cell growth inhibition. *Small*, *5*(6), 672–675. https://doi.org/10.1002/smll.200801821

18. Cong, R., Das, S., & Bouvet, P. (2011). The multiple properties and functions of nucleolin. In *The nucleolus* (pp. 185-212). Springer, New York, NY.
19. Courtenay-Luck, N., & Miller, D. M. (2008). AS1411: Development of an Anticancer Aptamer. *Therapeutic Oligonucleotides*, *12*, 189.
20. Dapić, V., Bates, P. J., Trent, J. O., Rodger, A., Thomas, S. D., & Miller, D. M. (2002). Antiproliferative activity of G-quartet-forming oligonucleotides with backbone and sugar modifications. *Biochemistry*, *41*(11), 3676–3685. https://doi.org/10.1021/bi0119520
21. Farin, K., & Pinkas-Kramarski, R. (2009). The crosstalk between ErbB1 and nucleolin. *Communicative and Integrative Biology*, *2*(6), 523–525. https://doi.org/10.4161/cib.2.6.9563
22. Farin, K., Schokoroy, S., Haklai, R., Cohen-Or, I., Elad-Sfadia, G., Reyes-Reyes, M. E., … Pinkas-Kramarski, R. (2011). Oncogenic synergism between ErbB1, nucleolin, and mutant ras. *Cancer Research*, *71*(6), 2140–2151. https://doi.org/10.1158/0008-5472.CAN-10-2887
23. Farzin, L., Shamsipur, M., Samandari, L., & Sheibani, S. (2018). Signalling probe displacement electrochemical aptasensor for malignant cell surface nucleolin as a breast cancer biomarker based on gold nanoparticle decorated hydroxyapatite nanorods and silver nanoparticle labels. *Microchimica Acta*, *185*(2). https://doi.org/10.1007/s00604-018-2700-2
24. Feng, L., Chen, Y., Ren, J., & Qu, X. (2011). A graphene functionalized electrochemical aptasensor for selective label-free detection of cancer cells. *Biomaterials*, *32*(11), 2930–2937. https://doi.org/10.1016/j.biomaterials.2011.01.002
25. Feng, L., Li, W., Ren, J., & Qu, X. (2015). Electrochemically and DNA-triggered cell release from ferrocene/β-cyclodextrin and aptamer modified dualfunctionalized graphene substrate. *Nano Research*, *8*(3), 887–899. https://doi.org/10.1007/s12274-014-0570-4
26. Girvan, A. C., Teng, Y., Casson, L. K., Thomas, S. D., Jüliger, S., Ball, M. W., … Bates, P. J. (2006). AGRO100 inhibits activation of nuclear factor-κB (NF-κB) by forming a complex with NF-κB essential modulator (NEMO) and nucleolin. *Molecular Cancer Therapeutics*, *5*(7), 1790–1799. https://doi.org/10.1158/1535-7163.MCT-05-0361
27. Goldshmit, Y., Trangle, S. S., Kloog, Y., & Pinkas-Kramarski, R. (2014). Interfering with the interaction between ErbB1, nucleolin and Ras as a potential treatment for glioblastoma. *Oncotarget*, *5*(18), 8602–8613. https://doi.org/10.18632/oncotarget.2343
28. Hong, E. J., Kim, Y. S., Choi, D. G., & Shim, M. S. (2018). Cancer-targeted photothermal therapy using aptamer-conjugated gold nanoparticles. *Journal of Industrial and Engineering Chemistry*, *67*, 429–436. https://doi.org/10.1016/j.jiec.2018.07.017
29. Hu, J., Zhao, Z., Liu, Q., Ye, M., Hu, B., Wang, J., & Tan, W. (2015). Study of the function of G-rich aptamers selected for lung adenocarcinoma. *Chemistry - An Asian Journal*, *10*(7), 1519–1525. https://doi.org/10.1002/asia.201500187
30. Hwang, D. W., Ko, H. Y., Lee, J. H., Kang, H., Ryu, S. H., Song, I. C., … Kim, S. (2010). A Nucleolin-Targeted Multimodal Nanoparticle Imaging Probe for Tracking Cancer Cells Using an Aptamer. *Journal of Nuclear Medicine*, *51*(1), 98–105. https://doi.org/10.2967/jnumed.109.069880
31. Ireson, C. R., & Kelland, L. R. (2006). Discovery and development of anticancer aptamers. *Molecular Cancer Therapeutics*, *5*(12), 2957–2962. https://doi.org/10.1158/1535-7163.MCT-06-0172
32. Ishimaru, D., Zuraw, L., Ramalingam, S., Sengupta, T. K., Bandyopadhyay, S., Reuben, A., … Spicer, E. K. (2010). Mechanism of regulation of bcl-2 mRNA by nucleolin and A+U-rich element-binding factor 1 (AUF1). *Journal of Biological Chemistry*, *285*(35), 27182–27191. https://doi.org/10.1074/jbc.M109.098830
33. Jiang, F., Liu, B., Lu, J., Li, F., Li, D., Liang, C., … Zhang, G. (2015). Progress and challenges in developing aptamer-functionalized targeted drug delivery systems. *International Journal of Molecular Sciences*, *16*(10), 23784–23822. https://doi.org/10.3390/ijms161023784
34. Kang, W. J., Ko, M. H., Lee, D. S., & Kim, S. (2009). Bioimaging of geographically adjacent proteins in a single cell by quantum dot-based fluorescent resonance energy transfer. *Proteomics - Clinical Applications*, *3*(12), 1383–1388. https://doi.org/10.1002/prca.200900077
35. Ko, M. H., Kim, S., Kang, W. J., Lee, J. H., Kang, H., Moon, S. H., … Lee, D. S. (2009). In vitro derby imaging of cancer biomarkers using quantum dots. *Small*, *5*(10), 1207–1212. https://doi.org/10.1002/smll.200801580
36. Kruspe, S., & Giangrande, P. H. (2017). Aptamer-siRNA chimeras: Discovery, progress, and future prospects. *Biomedicines*, *5*(3), 1–20. https://doi.org/10.3390/biomedicines5030045
37. Lai, P.-S., Pai, C.-L., Hsu, C.-Y., & Shieh, M.-J. (2010). AS1411 Aptamer-Conjugated Polymeric Micelle for Targetable Cancer Therapy. *NSTI-Nanotech*, *3*, 330–333.
38. Leaderer, D., Cashman, S. M., & Kumar-Singh, R. (2015). Topical application of a G-Quartet aptamer targeting nucleolin attenuates choroidal neovascularization in a model of age-related macular degeneration. *Experimental Eye Research*, *140*, 171–178. https://doi.org/10.1016/j.exer.2015.09.005
39. Lee, J. H., Yigit, M. V., Mazumdar, D., & Lu, Y. (2010). Molecular diagnostic and drug delivery agents based on aptamer-nanomaterial conjugates. *Advanced drug delivery reviews*, *62*(6), 592-605.
40. Lee, K. Y., Kang, H., Ryu, S. H., Lee, D. S., Lee, J. H., & Kim, S. (2010). Bioimaging of nucleolin aptamer-containing 5-(N -benzylcarboxyamide)- 2′-deoxyuridine more capable of specific binding to targets in cancer cells. *Journal of Biomedicine and Biotechnology*, *2010*. https://doi.org/10.1155/2010/168306
41. Li, H., Yang, S., Yu, G., Shen, L., Fan, J., Xu, L., … Zu, Y. (2017). Aptamer internalization via endocytosis inducing s-phase arrest and priming maver-1 lymphoma cells for cytarabine chemotherapy. *Theranostics*, *7*(5), 1204–1213. https://doi.org/10.7150/thno.17069
42. Li, J., Zheng, H., Bates, P. J., Malik, T., Li, X. F., Trent, J. O., & Ng, C. K. (2014). Aptamer imaging with Cu-64 labeled AS1411: Preliminary assessment in lung cancer. *Nuclear Medicine and Biology*, *41*(2), 179–185. https://doi.org/10.1016/j.nucmedbio.2013.10.008
43. Li, L. Le, Yin, Q., Cheng, J., & Lu, Y. (2012). Polyvalent mesoporous silica nanoparticle-aptamer bioconjugates target breast cancer cells. *Advanced Healthcare Materials*, *1*(5), 567–572. https://doi.org/10.1002/adhm.201200116
44. Li, L., Hou, J., Liu, X., Guo, Y., Wu, Y., Zhang, L., & Yang, Z. (2014). Nucleolin-targeting liposomes guided by aptamer AS1411 for the delivery of siRNA for the treatment of malignant melanomas. *Biomaterials*, *35*(12), 3840–3850. https://doi.org/10.1016/j.biomaterials.2014.01.019
45. Lian, S., Zhang, P., Gong, P., Hu, D., Shi, B., Zeng, C., & Cai, L. (2012). A universal quantum dots-aptamer probe for efficient cancer detection and targeted imaging. *Journal of Nanoscience and Nanotechnology*, *12*(10), 7703–7708. https://doi.org/10.1166/jnn.2012.6622
46. Liao, Z. X., Chuang, E. Y., Lin, C. C., Ho, Y. C., Lin, K. J., Cheng, P. Y., … Sung, H. W. (2015). An AS1411 aptamer-conjugated liposomal system containing a bubble-generating agent for tumor-specific chemotherapy that overcomes multidrug resistance. *Journal of Controlled Release*, *208*, 42–51. https://doi.org/10.1016/j.jconrel.2015.01.032
47. Litchfield, L. M., Riggs, K. A., Hockenberry, A. M., Oliver, L. D., Barnhart, K. G., Cai, J., … Klinge, C. M. (2012). Identification and characterization of nucleolin as a COUP-TFII coactivator of retinoic acid receptor β transcription in breast cancer cells. *PLoS ONE*, *7*(5), 1–14. https://doi.org/10.1371/journal.pone.0038278
48. Lorents, A., Säälik, P., Langel, Ü., & Pooga, M. (2018). Arginine-Rich Cell-Penetrating Peptides Require Nucleolin and Cholesterol-Poor Subdomains for Translocation across Membranes.

*Bioconjugate Chemistry*, *29*(4), 1168–1177. https://doi.org/10.1021/acs.bioconjchem.7b00805
49. Luo, Z., Yan, Z., Jin, K., Pang, Q., Jiang, T., Lu, H., … Jiang, X. (2017). Precise glioblastoma targeting by AS1411 aptamer-functionalized poly (L-γ-glutamylglutamine)–paclitaxel nanoconjugates. *Journal of Colloid and Interface Science*, *490*, 783–796. https://doi.org/10.1016/j.jcis.2016.12.004
50. Mahlknecht, G., Maron, R., Mancini, M., Schechter, B., Sela, M., & Yarden, Y. (2013). Aptamer to ErbB-2/HER2 enhances degradation of the target and inhibits tumorigenic growth. *Proceedings of the National Academy of Sciences*, *110*(20), 8170-8175.
51. Maier, K. E., & Levy, M. (2016). From selection hits to clinical leads: progress in aptamer discovery. *Molecular Therapy - Methods and Clinical Development*, *3*(November 2015), 16014. https://doi.org/10.1038/mtm.2016.14
52. Malik, M. T., O'Toole, M. G., Casson, L. K., Thomas, S. D., Bardi, G. T., Reyes-Reyes, E. M., … Bates, P. J. (2015). AS1411-conjugated gold nanospheres and their potential for breast cancer therapy. *Oncotarget*, *6*(26), 22270–22281. https://doi.org/10.18632/oncotarget.4207
53. Maremanda, N. G., Roy, K., Kanwar, R. K., Shyamsundar, V., Ramshankar, V., Krishnamurthy, A., … Kanwar, J. R. (2015). Quick chip assay using locked nucleic acid modified epithelial cell adhesion molecule and nucleolin aptamers for the capture of circulating tumor cells. *Biomicrofluidics*, *9*(5), 1–20. https://doi.org/10.1063/1.4930983
54. Métifiot, M., Amrane, S., Mergny, J. L., & Andreola, M. L. (2015). Anticancer molecule AS1411 exhibits low nanomolar antiviral activity against HIV-1. *Biochimie*, *118*, 173–175. https://doi.org/10.1016/j.biochi.2015.09.009
55. Motaghi, H., Mehrgardi, M. A., & Bouvet, P. (2017). Carbon Dots-AS1411 Aptamer Nanoconjugate for Ultrasensitive Spectrofluorometric Detection of Cancer Cells. *Scientific Reports*, *7*(1), 1–8. https://doi.org/10.1038/s41598-017-11087-2
56. Ni, X., Castanares, M., Mukherjee, A., & Lupold, S. E. (2011). Nucleic acid aptamers: clinical applications and promising new horizons. *Current medicinal chemistry*, *18*(27), 4206-4214.
57. Noaparast, Z., Hosseinimehr, S. J., Piramoon, M., & Abedi, S. M. (2015). Tumor targeting with a 99mTc-labeled AS1411 aptamer in prostate tumor cells. *Journal of Drug Targeting*, *23*(6), 497–505. https://doi.org/10.3109/1061186X.2015.1009075
58. Oh, S. S., Lee, B. F., Leibfarth, F. A., Eisenstein, M., Robb, M. J., Lynd, N. A., … Soh, H. T. (2014). Synthetic aptamer-polymer hybrid constructs for programmed drug delivery into specific target cells. *Journal of the American Chemical Society*, *136*(42), 15010–15015. https://doi.org/10.1021/ja5079464
59. Ozalp, V. C., Eyidogan, F., & Oktem, H. A. (2011). Aptamer-gated nanoparticles for smart drug delivery. *Pharmaceuticals*, *4*(8), 1137–1157. https://doi.org/10.3390/ph4081137
60. Park, J. Y., Cho, Y. L., Chae, J. R., Moon, S. H., Cho, W. G., Choi, Y. J., … Kang, W. J. (2018). Gemcitabine-Incorporated G-Quadruplex Aptamer for Targeted Drug Delivery into Pancreas Cancer. *Molecular Therapy - Nucleic Acids*, *12*(September), 543–553. https://doi.org/10.1016/j.omtn.2018.06.003
61. Park, S., Hwang, D., & Chung, J. (2012). Cotinine-conjugated aptamer/anti-cotinine antibody complexes as a novel affinity unit for use in biological assays. *Experimental and Molecular Medicine*, *44*(9), 554–561. https://doi.org/10.3858/emm.2012.44.9.063
62. Pasternak, A., Hernandez, F. J., Rasmussen, L. M., Vester, B., & Wengel, J. (2011). Improved thrombin binding aptamer by incorporation of a single unlocked nucleic acid monomer. *Nucleic Acids Research*, *39*(3), 1155–1164. https://doi.org/10.1093/nar/gkq823
63. Perrone, R., Butovskaya, E., Lago, S., Garzino-Demo, A., Pannecouque, C., Palù, G., & Richter, S. N. (2016). The G-quadruplex-forming aptamer AS1411 potently inhibits HIV-1 attachment to the host cell. *International Journal of Antimicrobial Agents*, *47*(4), 311–316. https://doi.org/10.1016/j.ijantimicag.2016.01.016
64. Reyes-Reyes, E. M., Šalipur, F. R., Shams, M., Forsthoefel, M. K., & Bates, P. J. (2015). Mechanistic studies of anticancer aptamer AS1411 reveal a novel role for nucleolin in regulating Rac1 activation. *Molecular Oncology*, *9*(7), 1392–1405. https://doi.org/10.1016/j.molonc.2015.03.012
65. Reyes-Reyes, E. M., Teng, Y., & Bates, P. J. (2010). A new paradigm for aptamer therapeutic AS1411 action: Uptake by macropinocytosis and its stimulation by a nucleolin-dependent mechanism. *Cancer Research*, *70*(21), 8617–8629. https://doi.org/10.1158/0008-5472.CAN-10-0920
66. Reyes-Reyes, E. M., & Bates, P. J. (2019). Characterizing Oligonucleotide Uptake in Cultured Cells: A Case Study Using AS1411 Aptamer. In *Methods in Molecular Biology* (Vol. 2036, pp. 173–186). https://doi.org/10.1007/978-1-4939-9670-4_10
67. Schokoroy, S., Juster, D., Kloog, Y., & Pinkas-Kramarski, R. (2013). Disrupting the Oncogenic Synergism between Nucleolin and Ras Results in Cell Growth Inhibition and Cell Death. *PLoS ONE*, *8*(9), 22–24. https://doi.org/10.1371/journal.pone.0075269
68. Sharma, V. R., Thomas, S. D., Miller, D. M., & Rezzoug, F. (2018). Nucleolin overexpression confers increased sensitivity to the anti-nucleolin aptamer, AS1411. *Cancer Investigation*, *36*, 475–491. https://doi.org/10.1080/07357907.2018.1527930
69. Shiao, Y. S., Chiu, H. H., Wu, P. H., & Huang, Y. F. (2014). Aptamer-functionalized gold nanoparticles as photoresponsive nanoplatform for Co-drug delivery. *ACS Applied Materials and Interfaces*, *6*(24), 21832–21841. https://doi.org/10.1021/am5026243
70. Shieh, M. J., Shieh, Y. A., & Lai, P. S. (2009). AS1411 aptamer for targetable photosensitizer delivery. *2nd International Conference on Biomedical and Pharmaceutical Engineering, ICBPE 2009 - Conference Proceedings*, (1), 1–5. https://doi.org/10.1109/ICBPE.2009.5384070
71. Shieh, Y., Yang, S., Wei, M., & Shieh, M. (2010). Aptamer-Based Tumor-Targeted Drug, *4*(3), 1433–1442.
72. Soundararajan, S., Chen, W., Spicer, E. K., Courtenay-Luck, N., & Fernandes, D. J. (2008). The nucleolin targeting aptamer AS1411 destabilizes Bcl-2 messenger RNA in human breast cancer cells. *Cancer Research*, *68*(7), 2358–2365. https://doi.org/10.1158/0008-5472.CAN-07-5723
73. Soundararajan, S., Wang, L. L., Sridharan, V., Chen, W., Courtenay-Luck, N., Jones, D., … Fernandes, D. J. (2009). Plasma Membrane Nucleolin Is a Receptor for the Anticancer Aptamer AS1411 in MV4-11 Leukemia Cells. *Molecular Pharmacology*, *76*(5), 984–991. https://doi.org/10.1124/mol.109.055947
74. Taghavi, S., Nia, A. H., Abnous, K., & Ramezani, M. (2017). Polyethylenimine-functionalized carbon nanotubes tagged with AS1411 aptamer for combination gene and drug delivery into human gastric cancer cells. *International Journal of Pharmaceutics*, *516*(1–2), 301–312. https://doi.org/10.1016/j.ijpharm.2016.11.027
75. Teng, Y., Girvan, A. C., Casson, L. K., Pierce, W. M., Qian, M., Thomas, S. D., & Bates, P. J. (2007). AS1411 alters the localization of a complex containing protein arginine methyltransferase 5 and nucleolin. *Cancer Research*, *67*(21), 10491–10500. https://doi.org/10.1158/0008-5472.CAN-06-4206
76. Tosoni, E., Frasson, I., Scalabrin, M., Perrone, R., Butovskaya, E., Nadai, M., … Richter, S. N. (2015). Nucleolin stabilizes G-quadruplex structures folded by the LTR promoter and silences HIV-1 viral transcription. *Nucleic Acids Research*, *43*(18), 8884–8897. https://doi.org/10.1093/nar/gkv897
77. Trinh, T. Le, Zhu, G., Xiao, X., Puszyk, W., Sefah, K., Wu, Q., … Liu, C. (2015). A synthetic aptamer-drug adduct for targeted liver cancer therapy. *PLoS ONE*, *10*(11), 1–17. https://doi.org/10.1371/journal.pone.0136673
78. Wang, W., Luo, J., Xiang, F., Liu, X., Jiang, M., Liao, L., & Hu, J. (2014). Nucleolin down-regulation is involved in ADP-induced cell cycle arrest in S phase and cell apoptosis in vascular

endothelial cells. *PLoS ONE*, *9*(10), 1–12. https://doi.org/10.1371/journal.pone.0110101
79. Wolfson, E., Goldenberg, M., Solomon, S., Frishberg, A., & Pinkas-Kramarski, R. (2016). Nucleolin-binding by ErbB2 enhances tumorigenicity of ErbB2-positive breast cancer. *Oncotarget*, *7*(40), 65320–65334. https://doi.org/10.18632/oncotarget.11323
80. Wolfson, E., Solomon, S., Schmukler, E., Goldshmit, Y., & Pinkas-Kramarski, R. (2018). Nucleolin and ErbB2 inhibition reduces tumorigenicity of ErbB2-positive breast cancer article. *Cell Death and Disease*, *9*(2). https://doi.org/10.1038/s41419-017-0067-7
81. Wu, J., Song, C., Jiang, C., Shen, X., Qiao, Q., & Hu, Y. (2013). Nucleolin targeting AS1411 modified protein nanoparticle for antitumor drugs delivery. *Molecular Pharmaceutics*, *10*(10), 3555–3563. https://doi.org/10.1021/mp300686g
82. Xing, H., Tang, L., Yang, X., Hwang, K., Wang, W., Yin, Q., … Lu, Y. (2013). Selective Delivery of an Anticancer Drug with Aptamer- Functionalized Liposomes to Breast Cancer Cells in Vitro and in Vivo Hang. *Journal of Materials Chemistry B*, *1*(39), 5288–5297. https://doi.org/10.1039/C3TB20412J
83. Xu, X., Hamhouyia, F., Thomas, S. D., Burke, T. J., Girvan, A. C., McGregor, W. G., … Bates, P. J. (2001). Inhibition of DNA Replication and Induction of S Phase Cell Cycle Arrest by G-rich Oligonucleotides. *Journal of Biological Chemistry*, *276*(46), 43221–43230. https://doi.org/10.1074/jbc.M104446200
84. Yan, A. C., & Levy, M. (2018). Aptamer-Mediated Delivery and Cell-Targeting Aptamers: Room for Improvement. *Nucleic Acid Therapeutics*, *28*(3), 194–199. https://doi.org/10.1089/nat.2018.0732
85. Yazdian-Robati, R., Bayat, P., Oroojalian, F., Zargari, M., Ramezani, M., Taghdisi, S. M., & Abnous, K. (2019). Therapeutic applications of AS1411 aptamer, an update review. *International Journal of Biological Macromolecules*, (xxxx). https://doi.org/10.1016/j.ijbiomac.2019.11.118
86. Yüce, M., Ullah, N., & Budak, H. (2015). Trends in aptamer selection methods and applications. *Analyst*. https://doi.org/10.1039/c5an00954e
87. Zhou, J., & Rossi, J. J. (2011). Cell-Specific aptamer-mediated targeted drug delivery. *Oligonucleotides*, *21*(1), 1–10. https://doi.org/10.1089/oli.2010.0264
88. Zhou, J., & Rossi, J. J. (2014). Cell-type-specific, aptamer-functionalized agents for targeted disease therapy. *Molecular Therapy - Nucleic Acids*, *3*(March), e169. https://doi.org/10.1038/mtna.2014.21
89. Zhou, L., Li, Z., Ju, E., Liu, Z., Ren, J., & Qu, X. (2013). Aptamer-directed synthesis of multifunctional lanthanide-doped porous nanoprobes for targeted imaging and drug delivery. *Small*, *9*(24), 4262–4268. https://doi.org/10.1002/smll.201301239
90. Zhu, G., & Chen, X. (2018). Aptamer-based targeted therapy. *Advanced Drug Delivery Reviews*, *134*, 65–78. https://doi.org/10.1016/j.addr.2018.08.005

***S9: 37mer DNA anti-Immunoglobulin E (IgE) aptamer (D17.4m Wiegand et al., 1996) References***

1. Adhikari, M., Strych, U., Kim, J., Goux, H., Dhamane, S., Poongavanam, M. V., … Willson, R. C. (2015). Aptamer-Phage Reporters for Ultrasensitive Lateral Flow Assays. *Analytical Chemistry*, *87*(23), 11660–11665. https://doi.org/10.1021/acs.analchem.5b00702
2. Bai, Y., & Zhao, Q. (2017). Rapid fluorescence detection of immunoglobulin e using an aptamer switch based on a binding-induced pyrene excimer. *Analytical Methods*, *9*(26), 3962–3967. https://doi.org/10.1039/c7ay01308f
3. Bai, Y., Li, Y., Zhang, D., Wang, H., & Zhao, Q. (2017). Enhancing the Affinity of Anti-Human α-Thrombin 15-mer DNA Aptamer and Anti-Immunoglobulin E Aptamer by PolyT Extension. *Analytical Chemistry*, *89*(17), 9467-9473.
4. Balamurugan, S., Obubuafo, A., Soper, S. A., & Spivak, D. A. (2008). Surface immobilization methods for aptamer diagnostic applications. *Analytical and Bioanalytical Chemistry*, *390*(4), 1009–1021. https://doi.org/10.1007/s00216-007-1587-2
5. Basnar, B., Elnathan, R., & Willner, I. (2006). Following aptamer−thrombin binding by force measurements. *Analytical chemistry*, *78*(11), 3638-3642.
6. Buchanan, D. D., Jameson, E. E., Perlette, J., Malik, A., & Kennedy, R. T. (2003). Effect of buffer, electric field, and separation time on detection of aptamer-ligand complexes for affinity probe capillary electrophoresis. *Electrophoresis*, *24*(9), 1375–1382. https://doi.org/10.1002/elps.200390176
7. Cheow, L. F., & Han, J. (2011). Continuous signal enhancement for sensitive aptamer affinity probe electrophoresis assay using electrokinetic concentration. *Analytical Chemistry*, *83*(18), 7086–7093. https://doi.org/10.1021/ac201307d
8. Cho, E. J., Collett, J. R., Szafranska, A. E., & Ellington, A. D. (2006). Optimization of aptamer microarray technology for multiple protein targets. *Analytica Chimica Acta*, *564*(1), 82–90. https://doi.org/10.1016/j.aca.2005.12.038
9. Cole, J. R., Dick, L. W., Morgan, E. J., & McGown, L. B. (2007). Affinity capture and detection of immunoglobulin E in human serum using an aptamer-modified surface in matrix-assisted laser desorption/ionization mass spectrometry. *Analytical Chemistry*, *79*(1), 273–279. https://doi.org/10.1021/ac061256b
10. Collett, J. R., Eun, J. C., & Ellington, A. D. (2005). Production and processing of aptamer microarrays. *Methods*, *37*(1), 4–15. https://doi.org/10.1016/j.ymeth.2005.05.009
11. Ding, S., Gao, C., & Gu, L. Q. (2009). Capturing single molecules of immunoglobulin and ricin with an aptamer-encoded glass nanopore. *Analytical Chemistry*, *81*(16), 6649–6655. https://doi.org/10.1021/ac9006705
12. Feng, K., Sun, C., Jiang, J., & Yu, R. (2011). An aptamer-based competitive fluorescence quenching assay for IgE. *Analytical letters*, *44*(7), 1301-1309.
13. Feng, K., Kang, Y., Zhao, J. J., Liu, Y. L., Jiang, J. H., Shen, G. L., & Yu, R. Q. (2008). Electrochemical immunosensor with aptamer-based enzymatic amplification. *Analytical Biochemistry*, *378*(1), 38–42. https://doi.org/10.1016/j.ab.2008.03.047
14. Fischer, N. O., & Tarasow, T. M. (2011). Identification and optimization of DNA aptamer binding regions using DNA microarrays. In *Protein Microarray for Disease Analysis* (pp. 57-66). Humana Press.
15. Fischer, N. O., Tok, J. B. H., & Tarasow, T. M. (2008). Massively parallel interrogation of aptamer sequence, structure and function. *PLoS ONE*, *3*(7), 1–10. https://doi.org/10.1371/journal.pone.0002720
16. Fukasawa, M., Yoshida, W., Yamazaki, H., Sode, K., & Ikebukuro, K. (2009). An aptamer-based bound/free separation system for protein detection. *Electroanalysis*, *21*(11), 1297–1302. https://doi.org/10.1002/elan.200804555
17. German, I., Buchanan, D. D., & Kennedy, R. T. (1998). Aptamers as ligands in affinity probe capillary electrophoresis. *Analytical Chemistry*, *70*(21), 4540–4545. https://doi.org/10.1021/ac980638h
18. Gokulrangan, G., Unruh, J. R., Holub, D. F., Ingram, B., Johnson, C. K., & Wilson, G. S. (2005). DNA aptamer-based bioanalysis of IgE by fluorescence anisotropy. *Analytical Chemistry*, *77*(7), 1963–1970. https://doi.org/10.1021/ac0483926
19. Gong, M., Wehmeyer, K. R., Limbach, P. A., & Heineman, W. R. (2007). Frontal analysis in microchip CE: A simple and accurate method for determination of protein-DNA dissociation constant. *Electrophoresis*, *28*, 837–842. https://doi.org/10.1002/elps.200600398

20. He, J. L., Wu, Z. S., Zhang, S. B., Shen, G. L., & Yu, R. Q. (2009). Novel fluorescence enhancement IgE assay using a DNA aptamer. *Analyst*, *134*(5), 1003–1007. https://doi.org/10.1039/b812450g
21. Hu, J., & Easley, C. J. (2011). A simple and rapid approach for measurement of dissociation constants of DNA aptamers against proteins and small molecules via automated microchip electrophoresis. *Analyst*, *136*(17), 3461–3468. https://doi.org/10.1039/c0an00842g
22. Jiang, B., Li, F., Yang, C., Xie, J., Xiang, Y., & Yuan, R. (2015). Aptamer pseudoknot-functionalized electronic sensor for reagentless and single-step detection of immunoglobulin e in human serum. *Analytical Chemistry*, *87*(5), 3094–3098. https://doi.org/10.1021/acs.analchem.5b00041
23. Jiang, Y., Fang, X., & Bai, C. (2004). Signaling aptamer/protein binding by a molecular light switch complex. *Analytical Chemistry*, *76*(17), 5230–5235. https://doi.org/10.1021/ac049565u
24. Jiang, Y., Zhu, C., Ling, L., Wan, L., Fang, X., & Bai, C. (2003). Specific aptamer-protein interaction studied by atomic force microscopy. *Analytical Chemistry*, *75*(9), 2112–2116. https://doi.org/10.1021/ac026182s
25. Jing, M., & Bowser, M. T. (2011). Isolation of DNA aptamers using micro free flow electrophoresis. *Lab on a Chip*, *11*(21), 3703–3709. https://doi.org/10.1039/c1lc20461k
26. Katilius, E., Flores, C., & Woodbury, N. W. (2007). Exploring the sequence space of a DNA aptamer using microarrays. *Nucleic Acids Research*, *35*(22), 7626–7635. https://doi.org/10.1093/nar/gkm922
27. Kim, S., Lee, J., Lee, S. J., & Lee, H. J. (2010). Ultra-sensitive detection of IgE using biofunctionalized nanoparticle-enhanced SPR. *Talanta*, *81*(4–5), 1755–1759. https://doi.org/10.1016/j.talanta.2010.03.036
28. Kim, Y. H., Kim, J. P., Han, S. J., & Sim, S. J. (2009). Aptamer biosensor for lable-free detection of human immunoglobulin E based on surface plasmon resonance. *Sensors and Actuators, B: Chemical*, *139*(2), 471–475. https://doi.org/10.1016/j.snb.2009.03.013
29. Lautnera, G., Gyurcsanyia, R., Mercierb, K., & Ly-Morinb, E. (2013). Monitoring of interactions between aptamers and human IgE by Surface Plasmon Resonance imaging. *Horiba Application Note 31*, 1–3. Retrieved from http://www.horiba.com/fileadmin/uploads/Scientific/Documents/SPRi/SPR31_-_Monitoring_of_interactions_LR.pdf
30. Lee, C. Y., Wu, K. Y., Su, H. L., Hung, H. Y., & Hsieh, Y. Z. (2013). Sensitive label-free electrochemical analysis of human IgE using an aptasensor with cDNA amplification. *Biosensors and Bioelectronics*, *39*(1), 133–138. https://doi.org/10.1016/j.bios.2012.07.009
31. Lee, J. O., So, H. M., Jeon, E. K., Chang, H., Won, K., & Kim, Y. H. (2008). Aptamers as molecular recognition elements for electrical nanobiosensors. *Analytical and Bioanalytical Chemistry*, *390*(4), 1023–1032. https://doi.org/10.1007/s00216-007-1643-y
32. Li, H., Qiang, W., Vuki, M., Xu, D., & Chen, H. Y. (2011). Fluorescence enhancement of silver nanoparticle hybrid probes and ultrasensitive detection of IgE. *Analytical Chemistry*, *83*(23), 8945–8952. https://doi.org/10.1021/ac201574s
33. Lin, L., Wang, H., Liu, Y., Yan, H., & Lindsay, S. (2006). Recognition imaging with a DNA aptamer. *Biophysical Journal*, *90*(11), 4236–4238. https://doi.org/10.1529/biophysj.105.079111
34. Liss, M., Petersen, B., Wolf, H., & Prohaska, E. (2002). An aptamer-based quartz crystal protein biosensor. *Analytical Chemistry*, *74*(17), 4488–4495. https://doi.org/10.1021/ac011294p
35. Liu, Y. M., Cao, J. T., Liu, Y. Y., Zhang, J. J., Zhou, M., Huang, K. J., … Ren, S. W. (2015). Aptamer-based detection and quantitative analysis of human immunoglobulin E in capillary electrophoresis with chemiluminescence detection. *Electrophoresis*, *36*(19), 2413–2418. https://doi.org/10.1002/elps.201500158
36. Luo, X., Lee, I., Huang, J., Yun, M., & Cui, X. T. (2011). Ultrasensitive protein detection using an aptamer-functionalized single polyaniline nanowire. *Chemical Communications*, *47*(22), 6368–6370. https://doi.org/10.1039/c1cc11353d
37. Maehashi, K., Katsura, T., Kerman, K., Takamura, Y., Matsumoto, K., & Tamiya, E. (2007). Label-free protein biosensor based on aptamer-modified carbon nanotube field-effect transistors. *Analytical Chemistry*, *79*(2), 782–787. https://doi.org/10.1021/ac060830g
38. Maehashi, K., Matsumoto, K., Takamur, Y., & Tamiya, E. (2009). Aptamer-based label-free immunosensors using carbon nanotube field-effect transistors. *Electroanalysis*, *21*(11), 1285–1290. https://doi.org/10.1002/elan.200804552
39. Mendonsa, S. D., & Bowser, M. T. (2004). In vitro selection of high-affinity DNA ligands for human IgE using capillary electrophoresis. *Analytical Chemistry*, *76*(18), 5387–5392. https://doi.org/10.1021/ac049857v
40. Nam, E. J., Kim, E. J., Wark, A. W., Rho, S., Kim, H., & Lee, H. J. (2012). Highly sensitive electrochemical detection of proteins using aptamer-coated gold nanoparticles and surface enzyme reactions. *Analyst*, *137*(9), 2011–2016. https://doi.org/10.1039/c2an15994e
41. Oh, S. S., Lee, B. F., Leibfarth, F. A., Eisenstein, M., Robb, M. J., Lynd, N. A., … Soh, H. T. (2014). Synthetic Aptamer-Polymer Hybrid Constructs for Programmed Drug Delivery into Specific Target Cells. *Journal of the American Chemical Society*, *136*, 15010–15015. https://doi.org/10.1021/ja5079464
42. Ohno, Y., Maehashi, K., Inoue, K., & Matsumoto, K. (2011). Label-free aptamer-based immunoglobulin sensors using graphene field-effect transistors. *Japanese Journal of Applied Physics*, *50*(7 PART 1). https://doi.org/10.1143/JJAP.50.070120
43. Papamichael, K. I., Kreuzer, M. P., & Guilbault, G. G. (2007). Viability of allergy (IgE) detection using an alternative aptamer receptor and electrochemical means. *Sensors and Actuators, B: Chemical*, *121*(1), 178–186. https://doi.org/10.1016/j.snb.2006.09.024
44. Park, M. K., Kee, J. S., Quah, J. Y., Netto, V., Song, J., Fang, Q., … Lo, G. Q. (2013). Label-free aptamer sensor based on silicon microring resonators. *Sensors and Actuators, B: Chemical*, *176*, 552–559. https://doi.org/10.1016/j.snb.2012.08.078
45. Pei, R., & Stojanovic, M. N. (2008). Study of thiazole orange in aptamer-based dye-displacement assays. *Analytical and Bioanalytical Chemistry*, *390*(4), 1093–1099. https://doi.org/10.1007/s00216-007-1773-2
46. Peng, Q., Cao, Z., Lau, C., Kai, M., & Lu, J. (2011). Aptamer-barcode based immunoassay for the instantaneous derivatization chemiluminescence detection of IgE coupled to magnetic beads. *Analyst*, *136*(1), 140–147. https://doi.org/10.1039/c0an00448k
47. Pollet, J., Delport, F., Thi, D. T., Weversy, M., & Lammertyn, J. (2008). Aptamer-Based Surface Plasmon Resonance Probe. *Proceedings of IEEE Sensors*, 1187–1190. https://doi.org/10.1109/ICSENS.2008.4716654
48. Pollet, J., Strych, U., & Willson, R. C. (2012). A peroxidase-active aptazyme as an isothermally amplifiable label in an aptazyme-linked oligonucleotide assay for low-picomolar IgE detection. *Analyst*, *137*(24), 5710–5712. https://doi.org/10.1039/c2an36201e
49. Poongavanam, M. V., Kisley, L., Kourentzi, K., Landes, C. F., & Willson, R. C. (2016). Ensemble and single-molecule biophysical characterization of D17.4 DNA aptamer-IgE interactions. *Biochimica et Biophysica Acta - Proteins and Proteomics*, *1864*(1), 154–164. https://doi.org/10.1016/j.bbapap.2015.08.008
50. Ruff, K. M., Snyder, T. M., & Liu, D. R. (2010). Enhanced functional potential of nucleic acid aptamer libraries patterned to increase secondary structure. *Journal of the American Chemical Society*, *132*(27), 9453–9464. https://doi.org/10.1021/ja103023m
51. Schachermeyer, S., Ashby, J., & Zhong, W. (2013). Aptamer-protein binding detected by asymmetric flow field flow fractionation. *Journal of Chromatography A*, *1295*, 107–113. https://doi.org/10.1016/j.chroma.2013.04.063
52. Sim, H. R., Wark, A. W., & Lee, H. J. (2010). Attomolar detection of protein biomarkers using biofunctionalized gold nanorods with

surface plasmon resonance. *Analyst*, *135*(10), 2528–2532. https://doi.org/10.1039/c0an00457j
53. Šnejdárková, M., Svobodová, L., Polohová, V., & Hianik, T. (2008). The study of surface properties of an IgE-sensitive aptasensor using an acoustic method. *Analytical and Bioanalytical Chemistry*, *390*(4), 1087–1091. https://doi.org/10.1007/s00216-007-1749-2
54. Spiridonova, V. A., Levashov, P. A., Ovchinnikova, E. D., Afanasieva, O. I., Glinkina, K. A., Adamova, I. Y., & Pokrovsky, S. N. (2014). DNA aptamer-based sorbents for binding human IgE. *Russian Journal of Bioorganic Chemistry*, *40*(2), 151–154. https://doi.org/10.1134/S1068162014020125
55. Stadtherr, K., Wolf, H., & Lindner, P. (2005). An aptamer-based protein biochip. *Analytical Chemistry*, *77*(11), 3437–3443. https://doi.org/10.1021/ac0483421
56. Tran, D. T., Vermeeren, V., Grieten, L., Wenmackers, S., Wagner, P., Pollet, J., … Lammertyn, J. (2011). Nanocrystalline diamond impedimetric aptasensor for the label-free detection of human IgE. *Biosensors and Bioelectronics*, *26*(6), 2987–2993. https://doi.org/10.1016/j.bios.2010.11.053
57. Turgeon, R. T., Fonslow, B. R., Jing, M., & Bowser, M. T. (2010). Measuring aptamer equilibria using gradient micro free flow electrophoresis. *Analytical Chemistry*, *82*(9), 3636–3641. https://doi.org/10.1021/ac902877v
58. Unruh, J. R., Gokulrangan, G., Lushington, G. H., Johnson, C. K., & Wilson, G. S. (2005). Orientational dynamics and dye-DNA interactions in a dye-labeled DNA aptamer. *Biophysical Journal*, *88*(5), 3455–3465. https://doi.org/10.1529/biophysj.104.054148
59. Unruh, J. R., Gokulrangan, G., Wilson, G. S., & Johnson, C. K. (2005). Fluorescence Properties of Fluorescein, Tetramethylrhodamine and Texas Red Linked to a DNA Aptamer. *Photochemistry and Photobiology*, *81*, 682–690.
60. Vinkenborg, J. L., Mayer, G., & Famulok, M. (2012). Aptamer-based affinity labeling of proteins. *Angewandte Chemie - International Edition*, *51*(36), 9176–9180. https://doi.org/10.1002/anie.201204174
61. Wang, H. Q., Wu, Z., Tang, L. J., Yu, R. Q., & Jiang, J. H. (2011). Fluorescence protection assay: A novel homogeneous assay platform toward development of aptamer sensors for protein detection. *Nucleic Acids Research*, *39*(18). https://doi.org/10.1093/nar/gkr559
62. Wang, J., Lv, R., Xu, J., Xu, D., & Chen, H. (2008). Characterizing the interaction between aptamers and human IgE by use of surface plasmon resonance. *Analytical and Bioanalytical Chemistry*, *390*(4), 1059–1065. https://doi.org/10.1007/s00216-007-1697-x
63. Wang, J., Munir, A., Li, Z., & Zhou, H. S. (2010). Aptamer-Au NPs conjugates-accumulated methylene blue for the sensitive electrochemical immunoassay of protein. *Talanta*, *81*(1–2), 63–67. https://doi.org/10.1016/j.talanta.2009.11.035
64. Wang, Z., Wilkop, T., Xu, D., Dong, Y., Ma, G., & Cheng, Q. (2007). Surface plasmon resonance imaging for affinity analysis of aptamer-protein interactions with PDMS microfluidic chips. *Analytical and Bioanalytical Chemistry*, *389*(3), 819–825. https://doi.org/10.1007/s00216-007-1510-x
65. Wiegand, T. W., Williams, P. B., Dreskin, S. C., Jouvin, M. H., Kinet, J. P., & Tasset, D. (1996). High-affinity oligonucleotide ligands to human IgE inhibit binding to Fc epsilon receptor I. *Journal of Immunology (Baltimore, Md. : 1950)*, *157*(1), 221–230. Retrieved from http://www.ncbi.nlm.nih.gov/pubmed/8683119
66. Wu, Z. S., Zheng, F., Shen, G. L., & Yu, R. Q. (2009). A hairpin aptamer-based electrochemical biosensing platform for the sensitive detection of proteins. *Biomaterials*, *30*(15), 2950–2955. https://doi.org/10.1016/j.biomaterials.2009.02.017
67. Xia, X., Piao, X., & Bong, D. (2014). Bifacial Peptide Nucleic Acid as an Allosteric Switch for Aptamer and Ribozyme Function. *Journal of the American Chemical Society*, *136*, 7265−7268. https://doi.org/10.1021/ja5032584
68. Xiao, Y., Uzawa, T., White, R. J., DeMartini, D., & Plaxco, K. W. (2009). On the signaling of electrochemical aptamer-based sensors: Collision- and folding-based mechanisms. *Electroanalysis*, *21*(11), 1267–1271. https://doi.org/10.1002/elan.200804564
69. Xu, D. D., Xu, D. D., Yu, X., Liu, Z., He, W., & Ma, Z. (2005). Label-free electrochemical detection for aptamer-based array electrodes. *Analytical Chemistry*, *77*(16), 5107–5113. https://doi.org/10.1021/ac050192m
70. Xu, D., Han, H., He, W., Liu, Z., Xu, D., & Liu, X. (2006). Electrically addressed fabrication of aptamer-based array electrodes. *Electroanalysis*, *18*(18), 1815–1820. https://doi.org/10.1002/elan.200603593
71. Yao, C., Qi, Y., Zhao, Y., Xiang, Y., Chen, Q., & Fu, W. (2009). Aptamer-based piezoelectric quartz crystal microbalance biosensor array for the quantification of IgE. *Biosensors and Bioelectronics*, *24*(8), 2499–2503. https://doi.org/10.1016/j.bios.2008.12.036
72. Yao, C., Zhu, T., Qi, Y., Zhao, Y., Xia, H., & Fu, W. (2010). Development of a quartz crystal microbalance biosensor with aptamers as bio-recognition element. *Sensors*, *10*(6), 5859–5871. https://doi.org/10.3390/s100605859
73. Yixian, W., Zunzhong, Y. E., Yibin, Y., Wang, Y., Ye, Z. Z., & Ying, Y. Bin. (2013). Detection of immunoglobulin E using an aptamer based dot-blot assay. *Chinese Science Bulletin*, *58*(24), 2938–2943. https://doi.org/10.1007/s11434-013-5702-9
74. Yoshida, W., Sode, K., & Ikebukuro, K. (2008). Label-free homogeneous detection of immunoglobulin E by an aptameric enzyme subunit. *Biotechnology Letters*, *30*(3), 421–425. https://doi.org/10.1007/s10529-007-9575-3
75. Zhang, X., & Yadavalli, V. K. (2011). Surface immobilization of DNA aptamers for biosensing and protein interaction analysis. *Biosensors and Bioelectronics*, *26*(7), 3142–3147. https://doi.org/10.1016/j.bios.2010.12.012

***S10: DNA anti-Ochratoxin A (OTA) aptamers (1.12 (2A, 61mer), 1.12.5 (1A, 36mer), and 1.12.8 (3A, 33mer), Cruz-Aguado et al., 2008) References***

1. Ali, W., & Pichon, V. (2014). Characterization of oligosorbents and application to the purification of ochratoxin A from wheat extracts. *Analytical and Bioanalytical Chemistry*, *406*(4), 1233–1240. https://doi.org/10.1007/s00216-013-7509-6
2. Baggiani, C., Giovannoli, C., & Anfossi, L. (2015). Man-made synthetic receptors for capture and analysis of ochratoxin A. *Toxins*, *7*(10), 4083–4098. https://doi.org/10.3390/toxins7104083
3. Barthelmebs, L., Hayat, A., Limiadi, A. W., Marty, J. L., & Noguer, T. (2011). Electrochemical DNA aptamer-based biosensor for OTA detection, using superparamagnetic nanoparticles. *Sensors and Actuators, B: Chemical*, *156*(2), 932–937. https://doi.org/10.1016/j.snb.2011.03.008
4. Barthelmebs, L., Jonca, J., Hayat, A., Prieto-Simon, B., & Marty, J. L. (2011). Enzyme-Linked Aptamer Assays (ELAAs), based on a competition format for a rapid and sensitive detection of Ochratoxin A in wine. *Food Control*, *22*(5), 737–743. https://doi.org/10.1016/j.foodcont.2010.11.005
5. Bianco, M., Sonato, A., De Girolamo, A., Pascale, M., Romanato, F., Rinaldi, R., & Arima, V. (2017). An aptamer-based SPR-polarization platform for high sensitive OTA

detection. *Sensors and Actuators, B: Chemical*, *241*, 314–320. https://doi.org/10.1016/j.snb.2016.10.056
6. Bonel, L., Vidal, J. C., Duato, P., & Castillo, J. R. (2011). An electrochemical competitive biosensor for ochratoxin A based on a DNA biotinylated aptamer. *Biosensors and Bioelectronics*, *26*(7), 3254–3259. https://doi.org/10.1016/j.bios.2010.12.036
7. Brothier, F., & Pichon, V. (2014). Miniaturized DNA aptamer-based monolithic sorbent for selective extraction of a target analyte coupled on-line to nanoLC. *Analytical and Bioanalytical Chemistry*, *406*(30), 7875–7886. https://doi.org/10.1007/s00216-014-8256-z
8. Cai, J., Hao, C., Sun, M., Ma, W., Xu, C., & Kuang, H. (2018). Chiral Shell Core–Satellite Nanostructures for Ultrasensitive Detection of Mycotoxin. *Small*, *14*(13), 1–8. https://doi.org/10.1002/smll.201703931
9. Castillo, G., Lamberti, I., Mosiello, L., & Hianik, T. (2012). Impedimetric DNA Aptasensor for Sensitive Detection of Ochratoxin A in Food. *Electroanalysis*, *24*(3), 512–520. https://doi.org/10.1002/elan.201100485
10. Chapuis-Hugon, F., Du Boisbaudry, A., Madru, B., & Pichon, V. (2011). New extraction sorbent based on aptamers for the determination of ochratoxin A in red wine. *Analytical and Bioanalytical Chemistry*, *400*(5), 1199–1207. https://doi.org/10.1007/s00216-010-4574-y
11. Chen, J., Fang, Z., Liu, J., & Zeng, L. (2012). A simple and rapid biosensor for ochratoxin A based on a structure-switching signaling aptamer. *Food Control*, *25*(2), 555–560. https://doi.org/10.1016/j.foodcont.2011.11.039
12. Chen, J., Zhang, X., Cai, S., Wu, D., Chen, M., Wang, S., & Zhang, J. (2014). A fluorescent aptasensor based on DNA-scaffolded silver-nanocluster for ochratoxin A detection. *Biosensors and Bioelectronics*, *57*, 226–231. https://doi.org/10.1016/j.bios.2014.02.001
13. Chen, Y. Y., Chen, M., Chi, J., Yu, X., Chen, Y. Y., Lin, X., & Xie, Z. (2018). Aptamer-based polyhedral oligomeric silsesquioxane (POSS)-containing hybrid affinity monolith prepared via a "one-pot" process for selective extraction of ochratoxin A. *Journal of Chromatography A*, *1563*, 37–46. https://doi.org/10.1016/j.chroma.2018.05.044
14. Chen, Y., Ding, X., Zhu, D., Lin, X., & Xie, Z. (2019). Preparation and evaluation of highly hydrophilic aptamer-based hybrid affinity monolith for on-column specific discrimination of ochratoxin A. *Talanta*, *200*(September 2018), 193–202. https://doi.org/10.1016/j.talanta.2019.03.053
15. Chen, Y., Yang, M., Xiang, Y., Yuan, R., & Chai, Y. (2014). Binding-induced autonomous disassembly of aptamer-DNAzyme supersandwich nanostructures for sensitive electrochemiluminescence turn-on detection of ochratoxin A. *Nanoscale*, *6*(2), 1099–1104. https://doi.org/10.1039/c3nr05499c
16. Chrouda, A., Sbartai, A., Baraket, A., Renaud, L., Maaref, A., & Jaffrezic-Renault, N. (2015). An aptasensor for ochratoxin A based on grafting of polyethylene glycol on a boron-doped diamond microcell. *Analytical Biochemistry*, *488*, 36–44. https://doi.org/10.1016/j.ab.2015.07.012
17. Costantini, F., Sberna, C., Petrucci, G., Reverberi, M., Domenici, F., Fanelli, C., … Caputo, D. (2016). Aptamer-based sandwich assay for on chip detection of Ochratoxin A by an array of amorphous silicon photosensors. *Sensors and Actuators, B: Chemical*, *230*, 31–39. https://doi.org/10.1016/j.snb.2016.02.036
18. Cruz-Aguado, J. A., & Penner, G. (2008). Determination of ochratoxin A with a DNA aptamer. *Journal of Agricultural and Food Chemistry*, *56*(22), 10456–10461. https://doi.org/10.1021/jf801957h
19. Dai, S., Wu, S., Duan, N., & Wang, Z. (2016). A near-infrared magnetic aptasensor for Ochratoxin A based on near-infrared upconversion nanoparticles and magnetic nanoparticles. *Talanta*, *158*, 246–253. https://doi.org/10.1016/j.talanta.2016.05.063
20. Dai, S., Wu, S., Duan, N., Chen, J., Zheng, Z., & Wang, Z. (2017). An ultrasensitive aptasensor for Ochratoxin A using hexagonal core/shell upconversion nanoparticles as luminophores. *Biosensors and Bioelectronics*, *91*(October 2016), 538–544. https://doi.org/10.1016/j.bios.2017.01.009
21. De Girolamo, A. De, Le, L., Penner, G., Schena, R., & Visconti, A. (2012). Analytical performances of a DNA-ligand system using time-resolved fluorescence for the determination of ochratoxin A in wheat. *Analytical and Bioanalytical Chemistry*, *403*(9), 2627–2634. https://doi.org/10.1007/s00216-012-6076-6
22. De Girolamo, A., McKeague, M., Miller, J. D., Derosa, M. C., & Visconti, A. (2011). Determination of ochratoxin A in wheat after clean-up through a DNA aptamer-based solid phase extraction column. *Food Chemistry*, *127*(3), 1378–1384. https://doi.org/10.1016/j.foodchem.2011.01.107
23. Deng, R., Dong, Y., Xia, X., Dai, Y., Zhang, K., He, Q., … Li, J. (2018). Recognition-Enhanced Metastably Shielded Aptamer for Digital Quantification of Small Molecules. *Analytical Chemistry*, *90*(24), 14347–14354. research-article. https://doi.org/10.1021/acs.analchem.8b03763
24. Deore, P. S., Gray, M. D., Chung, A. J., & Manderville, R. A. (2019). Ligand-Induced G-Quadruplex Polymorphism: A DNA Nanodevice for Label-Free Aptasensor Platforms. *Journal of the American Chemical Society*, *141*(36), 14288–14297. research-article. https://doi.org/10.1021/jacs.9b06533
25. Dou, X., Chu, X., Kong, W., Luo, J., & Yang, M. (2016). An indirect competitive fluorescence assay for ochratoxin A based on molecular beacon. *RSC Advances*, *6*(11), 8791–8796. https://doi.org/10.1039/c5ra23966d
26. Duan, N., Wu, S.-J., & Wang, Z.-P. (2011). An Aptamer-based Fluorescence Assay for Ochratoxin A. *Chinese Journal of Analytical Chemistry*, *39*(3), 300–304. https://doi.org/10.1016/S1872-2040(10)60423-9
27. Duan, N., Wu, S., Ma, X., Chen, X., Huang, Y., & Wang, Z. (2012). Gold Nanoparticle-Based Fluorescence Resonance Energy Transfer Aptasensor for Ochratoxin A Detection. *Analytical Letters*, *45*(7), 714–723. https://doi.org/10.1080/00032719.2011.653899
28. Evtugyn, G., Porfireva, A., Sitdikov, R., Evtugyn, V., Stoikov, I., Antipin, I., & Hianik, T. (2013). Electrochemical aptasensor for the determination of ochratoxin A at the Au electrode modified with Ag nanoparticles decorated with macrocyclic ligand. *Electroanalysis*, *25*(8), 1847–1854. https://doi.org/10.1002/elan.201300164
29. Evtugyn, G., Porfireva, A., Stepanova, V., Kutyreva, M., Gataulina, A., Ulakhovich, N., … Hianik, T. (2013). Impedimetric aptasensor for ochratoxin a determination based on Au nanoparticles stabilized with hyper-branched polymer. *Sensors (Switzerland)*, *13*(12), 16129–16145. https://doi.org/10.3390/s131216129
30. Galarreta, B. C., Tabatabaei, M., Guieu, V., Peyrin, E., & Lagugné-Labarthet, F. (2013). Microfluidic channel with embedded SERS 2D platform for the aptamer detection of ochratoxin A. *Analytical and Bioanalytical Chemistry*, *405*(5), 1613–1621. https://doi.org/10.1007/s00216-012-6557-7
31. Ganbold, E. O., Lee, C. M., Cho, E. M., Son, S. J., Kim, S., Joo, S. W., & Yang, S. I. (2014). Subnanomolar detection of ochratoxin A using aptamer-attached silver nanoparticles and surface-enhanced Raman scattering. *Analytical Methods*, *6*(11), 3573–3577. https://doi.org/10.1039/c4ay00440j
32. Geng, X., Zhang, D., Wang, H., & Zhao, Q. (2013). Screening interaction between ochratoxin A and aptamers by fluorescence anisotropy approach ABC Highlights: Authored by Rising Stars and Top Experts. *Analytical and Bioanalytical Chemistry*, *405*(8), 2443–2449. https://doi.org/10.1007/s00216-013-6736-1
33. Gu, C., Long, F., Zhou, X., & Shi, H. (2016). Portable detection of ochratoxin A in red wine based on a structure-switching aptamer using a personal glucometer. *RSC Advances*, *6*(35), 29563–29569. https://doi.org/10.1039/C5RA27880E

34. Guo, Z., Ren, J., Wang, J., & Wang, E. (2011). Single-walled carbon nanotubes based quenching of free FAM-aptamer for selective determination of ochratoxin A. *Talanta*, *85*(5), 2517–2521. https://doi.org/10.1016/j.talanta.2011.08.015
35. Hao, N., Jiang, L., Qian, J., & Wang, K. (2016). Ultrasensitive electrochemical Ochratoxin A aptasensor based on CdTe quantum dots functionalized graphene/Au nanocomposites and magnetic separation. *Journal of Electroanalytical Chemistry*, *781*, 332–338. https://doi.org/10.1016/j.jelechem.2016.09.053
36. Hayat, A., Andreescu, S., & Marty, J. L. (2013). Design of PEG-aptamer two piece macromolecules as convenient and integrated sensing platform: Application to the label free detection of small size molecules. *Biosensors and Bioelectronics*, *45*(1), 168–173. https://doi.org/10.1016/j.bios.2013.01.059
37. Hayat, A., Haider, W., Rolland, M., & Marty, J. L. (2013). Electrochemical grafting of long spacer arms of hexamethyldiamine on a screen printed carbon electrode surface: Application in target induced ochratoxin A electrochemical aptasensor. *Analyst*, *138*(10), 2951–2957. https://doi.org/10.1039/c3an00158j
38. Hayat, A., Mishra, R. K., Catanante, G., & Marty, J. L. (2015). Development of an aptasensor based on a fluorescent particles-modified aptamer for ochratoxin A detection. *Analytical and Bioanalytical Chemistry*, *407*(25), 7815–7822. https://doi.org/10.1007/s00216-015-8952-3
39. Hayat, A., Sassolas, A., Marty, J. L., & Radi, A. E. (2013). Highly sensitive ochratoxin A impedimetric aptasensor based on the immobilization of azido-aptamer onto electrografted binary film via click chemistry. *Talanta*, *103*, 14–19. https://doi.org/10.1016/j.talanta.2012.09.048
40. He, Y., Tian, F., Zhou, J., & Jiao, B. (2019). A fluorescent aptasensor for ochratoxin A detection based on enzymatically generated copper nanoparticles with a polythymine scaffold. *Microchimica Acta*, *186*(3). https://doi.org/10.1007/s00604-019-3314-z
41. Huang, L., Chen, K., Zhang, W., Zhu, W., Liu, X., Wang, J., … Wang, J. (2018). ssDNA-tailorable oxidase-mimicking activity of spinel MnCo2O4 for sensitive biomolecular detection in food sample. *Sensors and Actuators, B: Chemical*, *269*, 79–87. https://doi.org/10.1016/j.snb.2018.04.150
42. Huang, L., Wu, J., Zheng, L., Qian, H., Xue, F., Wu, Y., … Chen, W. (2013). Rolling chain amplification based signal-enhanced electrochemical aptasensor for ultrasensitive detection of ochratoxin A. *Analytical Chemistry*, *85*(22), 10842–10849. https://doi.org/10.1021/ac402228n
43. Hun, X., Liu, F., Mei, Z., Ma, L., Wang, Z., & Luo, X. (2013). Signal amplified strategy based on target-induced strand release coupling cleavage of nicking endonuclease for the ultrasensitive detection of ochratoxin A. *Biosensors and Bioelectronics*, *39*(1), 145–151. https://doi.org/10.1016/j.bios.2012.07.005
44. Ji, W., Zhang, Z., Tian, Y., Yang, Z., Cao, Z., Zhang, L., … Wang, H. (2019). Shape Coding Microhydrogel for a Real-Time Mycotoxin Detection System Based on Smartphones. *ACS Applied Materials and Interfaces*, *11*(8), 8584–8590. research-article. https://doi.org/10.1021/acsami.8b21851
45. Jiang, L., Qian, J., Yang, X., Yan, Y., Liu, Q., Wang, K., & Wang, K. (2014). Amplified impedimetric aptasensor based on gold nanoparticles covalently bound graphene sheet for the picomolar detection of ochratoxin A. *Analytica Chimica Acta*, *806*, 128–135. https://doi.org/10.1016/j.aca.2013.11.003
46. Jin, B., Yang, Y., He, R., Park, Y. Il, Lee, A., Bai, D., … Lin, M. (2018). Lateral flow aptamer assay integrated smartphone-based portable device for simultaneous detection of multiple targets using upconversion nanoparticles. *Sensors and Actuators, B: Chemical*, *276*(May), 48–56. https://doi.org/10.1016/j.snb.2018.08.074
47. Jo, E. J., Byun, J. Y., Mun, H., Bang, D., Son, J. H., Lee, J. Y., … Kim, M. G. (2018). Single-Step LRET Aptasensor for Rapid Mycotoxin Detection. *Analytical Chemistry*, *90*(1), 716–722. https://doi.org/10.1021/acs.analchem.7b02368
48. Jo, E. J., Mun, H., Kim, S. J., Shim, W. B., & Kim, M. G. (2016). Detection of ochratoxin A (OTA) in coffee using chemiluminescence resonance energy transfer (CRET) aptasensor. *Food Chemistry*, *194*, 1102–1107. https://doi.org/10.1016/j.foodchem.2015.07.152
49. Karimi, A., Hayat, A., & Andreescu, S. (2017). Biomolecular detection at ssDNA-conjugated nanoparticles by nano-impact electrochemistry. *Biosensors and Bioelectronics*, *87*, 501–507. https://doi.org/10.1016/j.bios.2016.08.108
50. Kidd, A., Guieu, V., Perrier, S., Ravelet, C., & Peyrin, E. (2011). Fluorescence polarization biosensor based on an aptamer enzymatic cleavage protection strategy. *Analytical and Bioanalytical Chemistry*, *401*(10), 3229–3234. https://doi.org/10.1007/s00216-011-5434-0
51. Kuang, H., Chen, W., Xu, D., Xu, L., Zhu, Y., Liu, L., … Zhu, S. (2010). Fabricated aptamer-based electrochemical " signal-off" sensor of ochratoxin A. *Biosensors and Bioelectronics*, *26*(2), 710–716. https://doi.org/10.1016/j.bios.2010.06.058
52. Lee, B., Park, J. H., Byun, J. Y., Kim, J. H., & Kim, M. G. (2018). An optical fiber-based LSPR aptasensor for simple and rapid in-situ detection of ochratoxin A. *Biosensors and Bioelectronics*, *102*(November 2017), 504–509. https://doi.org/10.1016/j.bios.2017.11.062
53. Lee, J., Jeon, C. H., Ahn, S. J., & Ha, T. H. (2014). Highly stable colorimetric aptamer sensors for detection of ochratoxin A through optimizing the sequence with the covalent conjugation of hemin. *Analyst*, *139*(7), 1622–1627. https://doi.org/10.1039/c3an01639k
54. Li, D. L., Zhang, X., Ma, Y., Deng, Y., Hu, R., & Yang, Y. (2018). Preparation of an OTA aptasensor based on a metal-organic framework. *Analytical Methods*, *10*(26), 3273–3279. https://doi.org/10.1039/c8ay00758f
55. Li, Y., & Zhao, Q. (2019). Aptamer Structure Switch Fluorescence Anisotropy Assay for Small Molecules Using Streptavidin as an Effective Signal Amplifier Based on Proximity Effect. *Analytical Chemistry*, *91*(11), 7379–7384. research-article. https://doi.org/10.1021/acs.analchem.9b01253
56. Li, Y., Sun, L., & Zhao, Q. (2019). Aptamer-Structure Switch Coupled with Horseradish Peroxidase Labeling on a Microplate for the Sensitive Detection of Small Molecules. *Analytical Chemistry*, 4–8. https://doi.org/10.1021/acs.analchem.8b05606
57. Liu, B., Huang, R., Yu, Y., Su, R., Qi, W., & He, Z. (2018). Gold nanoparticle-aptamer-based LSPR sensing of Ochratoxin A at a widened detection range by double calibration curve method. *Frontiers in Chemistry*, *6*(APR), 1–9. https://doi.org/10.3389/fchem.2018.00094
58. Liu, F., Ding, A., Zheng, J., Chen, J., & Wang, B. (2018). A label-free aptasensor for ochratoxin a detection based on the structure switch of aptamer. *Sensors (Switzerland)*, *18*(6). https://doi.org/10.3390/s18061769
59. Liu, L. hua, Zhou, X. hong, & Shi, H. chang. (2015). Portable optical aptasensor for rapid detection of mycotoxin with a reversible ligand-grafted biosensing surface. *Biosensors and Bioelectronics*, *72*, 300–305. https://doi.org/10.1016/j.bios.2015.05.033
60. Liu, L., Tanveer, Z. I., Jiang, K., Huang, Q., Zhang, J., Wu, Y., & Han, Z. (2019). Label-free fluorescent aptasensor for ochratoxin—A detection based on CdTe quantum dots and (N-methyl-4-pyridyl) porphyrin. *Toxins*, *11*(8). https://doi.org/10.3390/toxins11080447
61. Liu, R., Huang, Y., Ma, Y., Jia, S., Gao, M., Li, J., … Yang, C. (2015). Design and synthesis of target-responsive aptamer-cross-linked hydrogel for visual quantitative detection of ochratoxin A. *ACS Applied Materials and Interfaces*, *7*(12), 6982–6990. https://doi.org/10.1021/acsami.5b01120
62. Liu, Y., Yu, J., Wang, Y., Liu, Z., & Lu, Z. (2016). An ultrasensitive aptasensor for detection of Ochratoxin A based on shielding effect-induced inhibition of fluorescence resonance energy transfer. *Sensors and Actuators, B: Chemical*, *222*, 797–803. https://doi.org/10.1016/j.snb.2015.09.007

63. Loo, A. H., Bonanni, A., & Pumera, M. (2015). Mycotoxin Aptasensing Amplification by using Inherently Electroactive Graphene-Oxide Nanoplatelet Labels. *ChemElectroChem*, *2*(5), 743–747. https://doi.org/10.1002/celc.201402403
64. Lu, Z., Chen, X., & Hu, W. (2017). A fluorescence aptasensor based on semiconductor quantum dots and MoS2 nanosheets for ochratoxin A detection. *Sensors and Actuators, B: Chemical*, *246*, 61–67. https://doi.org/10.1016/j.snb.2017.02.062
65. Luan, Y., Chen, J., Li, C., Xie, G., Fu, H., Ma, Z., & Lu, A. (2015). Highly sensitive colorimetric detection of ochratoxin a by a label-free aptamer and gold nanoparticles. *Toxins*, *7*(12), 5377–5385. https://doi.org/10.3390/toxins7124883
66. Lv, L., Cui, C., Liang, C., Quan, W., Wang, S., & Guo, Z. (2016). Aptamer-based single-walled carbon nanohorn sensors for ochratoxin A detection. *Food Control*, *60*, 296–301. https://doi.org/10.1016/j.foodcont.2015.08.002
67. Lv, L., Jin, Y., Kang, X., Zhao, Y., Cui, C., & Guo, Z. (2018). PVP-coated gold nanoparticles for the selective determination of ochratoxin A via quenching fluorescence of the free aptamer. *Food Chemistry*, *249*(December 2017), 45–50. https://doi.org/10.1016/j.foodchem.2017.12.087
68. Lv, L., Li, D., Cui, C., Zhao, Y., & Guo, Z. (2017). Nuclease-aided target recycling signal amplification strategy for ochratoxin A monitoring. *Biosensors and Bioelectronics*, *87*, 136–141. https://doi.org/10.1016/j.bios.2016.08.024
69. Lv, L., Li, D., Liu, R., Cui, C., & Guo, Z. (2017). Label-free aptasensor for ochratoxin A detection using SYBR Gold as a probe. *Sensors and Actuators, B: Chemical*, *246*, 647–652. https://doi.org/10.1016/j.snb.2017.02.143
70. Lv, X., Zhang, Y., Liu, G., Du, L., & Wang, S. (2017). Aptamer-based fluorescent detection of ochratoxin A by quenching of gold nanoparticles. *RSC Advances*, *7*(27), 16290–16294. https://doi.org/10.1039/c7ra01474k
71. Lv, Z., Chen, A., Liu, J., Guan, Z., Zhou, Y., Xu, S., … Li, C. (2014). A simple and sensitive approach for ochratoxin a detection using a label-free fluorescent aptasensor. *PLoS ONE*, *9*(1), 2–6. https://doi.org/10.1371/journal.pone.0085968
72. Ma, L., Xu, B., Liu, L., & Tian, W. (2018). A Label-free Fluorescent Aptasensor for Turn-on Monitoring Ochratoxin A Based on AIE-active Probe and Graphene Oxide. *Chemical Research in Chinese Universities*, *34*(3), 363–368. https://doi.org/10.1007/s40242-018-8072-7
73. Ma, W., Yin, H., Xu, L., Xu, Z., Kuang, H., Wang, L., & Xu, C. (2013). Femtogram ultrasensitive aptasensor for the detection of Ochratoxin A. *Biosensors and Bioelectronics*, *42*(1), 545–549. https://doi.org/10.1016/j.bios.2012.11.024
74. Marechal, A., Jarrosson, F., Randon, J., Dugas, V., & Demesmay, C. (2015). In-line coupling of an aptamer based miniaturized monolithic affinity preconcentration unit with capillary electrophoresis and Laser Induced Fluorescence detection. *Journal of Chromatography A*, *1406*, 109–117. https://doi.org/10.1016/j.chroma.2015.05.073
75. Mazaafrianto, D. N., Ishida, A., Maeki, M., Tani, H., & Tokeshi, M. (2018). Label-Free Electrochemical Sensor for Ochratoxin A Using a Microfabricated Electrode with Immobilized Aptamer. *ACS Omega*, *3*(12), 16823–16830. research-article. https://doi.org/10.1021/acsomega.8b01996
76. McKeague, M., De Girolamo, A., Valenzano, S., Pascale, M., Ruscito, A., Velu, R., … DeRosa, M. C. (2015). Comprehensive Analytical Comparison of Strategies Used for Small Molecule Aptamer Evaluation. *Analytical Chemistry*, *87*(17), 8608–8612. https://doi.org/10.1021/acs.analchem.5b02102
77. McKeague, M., Velu, R., Hill, K., Bardóczy, V., Mészáros, T., & DeRosa, M. C. (2014). Selection and characterization of a novel DNA aptamer for label-free fluorescence biosensing of ochratoxin A. *Toxins*, *6*(8), 2435–2452. https://doi.org/10.3390/toxins6082435
78. Mejri-Omrani, N., Miodek, A., Zribi, B., Marrakchi, M., Hamdi, M., Marty, J. L., & Korri-Youssoufi, H. (2016). Direct detection of OTA by impedimetric aptasensor based on modified polypyrrole-dendrimers. *Analytica Chimica Acta*, *920*, 37–46. https://doi.org/10.1016/j.aca.2016.03.038
79. Mishra, R. K., Hayat, A., Catanante, G., Ocaña, C., & Marty, J. L. (2015). A label free aptasensor for Ochratoxin A detection in cocoa beans: An application to chocolate industries. *Analytica Chimica Acta*, *889*, 106–112. https://doi.org/10.1016/j.aca.2015.06.052
80. Modh, H., Scheper, T., & Walter, J. G. (2017). Detection of ochratoxin A by aptamer-assisted real-time PCR-based assay (Apta-qPCR). *Engineering in Life Sciences*, *17*(8), 923–930. https://doi.org/10.1002/elsc.201700048
81. Motyčka, J., Mach, P., Melicherčík, M., & Urban, J. (2014). DFT and MD study of the divalent-cation-mediated interaction of ochratoxin a with DNA nucleosides. *Journal of Molecular Modeling*, *20*(6). https://doi.org/10.1007/s00894-014-2274-9
82. Mun, H., Jo, E. J., Li, T., Joung, H. A., Hong, D. G., Shim, W. B., … Kim, M. G. (2014). Homogeneous assay of target molecules based on chemiluminescence resonance energy transfer (CRET) using DNAzyme-linked aptamers. *Biosensors and Bioelectronics*, *58*, 308–313. https://doi.org/10.1016/j.bios.2014.02.008
83. Muthamizh, S., Ribes, A., Anusuyajanakiraman, M., Narayanan, V., Soto, J., Martínez-Máñez, R., & Aznar, E. (2017). Implementation of oligonucleotide-gated supports for the electrochemical detection of Ochratoxin A. *Supramolecular Chemistry*, *29*(11), 776–783. https://doi.org/10.1080/10610278.2017.1390238
84. Nekrasov, N., Kireev, D., Emelianov, A., & Bobrinestskiy, I. (2019). Graphene-Based Sensing Platform for On-Chip Ochratoxin A Detection. *Toxins*, *11*(550). https://doi.org/10.3390/toxins11100550
85. Ni, J., Yang, W., Wang, Q., Luo, F., Guo, L., Qiu, B., … Yang, H. (2018). Homogeneous and label-free electrochemiluminescence aptasensor based on the difference of electrostatic interaction and exonuclease-assisted target recycling amplification. *Biosensors and Bioelectronics*, *105*(December 2017), 182–187. https://doi.org/10.1016/j.bios.2018.01.043
86. Niazi, S., Khan, I. M., Yan, L., Khan, M. I., Mohsin, A., Duan, N., … Wang, Z. (2019). Simultaneous detection of fumonisin B 1 and ochratoxin A using dual-color, time-resolved luminescent nanoparticles (NaYF 4 : Ce, Tb and NH 2 -Eu/DPA@SiO 2 ) as labels. *Analytical and Bioanalytical Chemistry*, *411*(7), 1453–1465. https://doi.org/10.1007/s00216-019-01580-0
87. Park, J. H., Byun, J. Y., Mun, H., Shim, W. B., Shin, Y. B., Li, T., & Kim, M. G. (2014). A regeneratable, label-free, localized surface plasmon resonance (LSPR) aptasensor for the detection of ochratoxin A. *Biosensors and Bioelectronics*, *59*, 321–327. https://doi.org/10.1016/j.bios.2014.03.059
88. Perrier, S., Zhu, Z., Fiore, E., Ravelet, C., Guieu, V., & Peyrin, E. (2014). Capillary gel electrophoresis-coupled aptamer enzymatic cleavage protection strategy for the simultaneous detection of multiple small analytes. *Analytical Chemistry*, *86*(9), 4233–4240. https://doi.org/10.1021/ac5010234
89. Phanchai, W., Srikulwong, U., Chompoosor, A., Sakonsinsiri, C., & Puangmali, T. (2018). Insight into the Molecular Mechanisms of AuNP-Based Aptasensor for Colorimetric Detection: A Molecular Dynamics Approach. *Langmuir*, *34*(21), 6161–6169. research-article. https://doi.org/10.1021/acs.langmuir.8b00701
90. Prabhakar, N., Matharu, Z., & Malhotra, B. D. (2011). Polyaniline Langmuir-Blodgett film based aptasensor for ochratoxin A detection. *Biosensors and Bioelectronics*, *26*(10), 4006–4011. https://doi.org/10.1016/j.bios.2011.03.014
91. Prieto-Simón, B., & Samitier, J. (2014). “Signal Off” Aptasensor Based on Enzyme Inhibition Induced By Conformational Switch. *Analytical Chemistry*, *86*(3), 1437–1444. https://doi.org/10.1021/ac402258x
92. Qian, J., Ren, C., Wang, C., Chen, W., Lu, X., Li, H., … Wang, K. (2018). Magnetically controlled fluorescence aptasensor for

simultaneous determination of ochratoxin A and aflatoxin B1. *Analytica Chimica Acta*, *1019*, 119–127. https://doi.org/10.1016/j.aca.2018.02.063

93. Qian, J., Wang, K., Wang, C., Hua, M., Yang, Z., Liu, Q., … Wang, K. (2015). A FRET-based ratiometric fluorescent aptasensor for rapid and onsite visual detection of ochratoxin A. *Analyst*, *140*(21), 7434–7442. https://doi.org/10.1039/c5an01403d
94. Qing, Y., Li, X., Chen, S., Zhou, X. P., Luo, M., Xu, X., … Qiu, J. F. (2017). Differential pulse voltammetric ochratoxin A assay based on the use of an aptamer and hybridization chain reaction. *Microchimica Acta*, *184*(3), 863–870. https://doi.org/10.1007/s00604-017-2080-z
95. Rhouati, A., Hayat, A., Hernandez, D. B., Meraihi, Z., Munoz, R., & Marty, J. L. (2013). Development of an automated flow-based electrochemical aptasensor for on-line detection of Ochratoxin A. *Sensors and Actuators, B: Chemical*, *176*, 1160–1166. https://doi.org/10.1016/j.snb.2012.09.111
96. Rivas, L., Mayorga-Martinez, C. C., Quesada-González, D., Zamora-Gálvez, A., De La Escosura-Muñiz, A., & Merkoçi, A. (2015). Label-free impedimetric aptasensor for ochratoxin-A detection using iridium oxide nanoparticles. *Analytical Chemistry*, *87*(10), 5167–5172. https://doi.org/10.1021/acs.analchem.5b00890
97. Samokhvalov, A. V., Safenkova, I. V., Eremin, S. A., Zherdev, A. V., & Dzantiev, B. B. (2017). Use of anchor protein modules in fluorescence polarisation aptamer assay for ochratoxin A determination. *Analytica Chimica Acta*, *962*, 80–87. https://doi.org/10.1016/j.aca.2017.01.024
98. Samokhvalov, A. V., Safenkova, I. V., Eremin, S. A., Zherdev, A. V., & Dzantiev, B. B. (2018). Measurement of (Aptamer-Small Target) KD Using the Competition between Fluorescently Labeled and Unlabeled Targets and the Detection of Fluorescence Anisotropy. *Analytical Chemistry*, *90*(15), 9189–9198. research-article. https://doi.org/10.1021/acs.analchem.8b01699
99. Samokhvalov, A. V., Safenkova, I. V., Zherdev, A. V., & Dzantiev, B. B. (2018). The registration of aptamer–ligand (ochratoxin A) interactions based on ligand fluorescence changes. *Biochemical and Biophysical Research Communications*, *505*(2), 536–541. https://doi.org/10.1016/j.bbrc.2018.09.109
100. Sanzani, S. M., Reverberi, M., Fanelli, C., & Ippolito, A. (2015). Detection of ochratoxin a using molecular beacons and real-time PCR thermal cycler. *Toxins*, *7*(3), 812–820. https://doi.org/10.3390/toxins7030812
101. Schax, E., Lönne, M., Scheper, T., Belkin, S., & Walter, J. G. (2015). Aptamer-based depletion of small molecular contaminants: A case study using ochratoxin A. *Biotechnology and Bioprocess Engineering*, *20*(6), 1016–1025. https://doi.org/10.1007/s12257-015-0486-1
102. Shao, B., Ma, X., Zhao, S., Lv, Y., Hun, X., Wang, H., & Wang, Z. (2018). Nanogapped Au(core) @ Au-Ag(shell) structures coupled with Fe3O4 magnetic nanoparticles for the detection of Ochratoxin A. *Analytica Chimica Acta*, *1033*, 165–172. https://doi.org/10.1016/j.aca.2018.05.058
103. Sharma, A., Hayat, A., Mishra, R. K., Catanante, G., Bhand, S., & Marty, J. L. (2015). Titanium dioxide nanoparticles (TiO2) quenching based aptasensing platform: Application to ochratoxin a detection. *Toxins*, *7*(9), 3771–3784. https://doi.org/10.3390/toxins7093771
104. Sharma, A., Hayat, A., Mishra, R. K., Catanante, G., Shahid, S. A., Bhand, S., & Marty, J. L. (2016). Design of a fluorescence aptaswitch based on the aptamer modulated nano-surface impact on the fluorescence particles. *RSC Advances*, *6*(70), 65579–65587. https://doi.org/10.1039/c6ra10942j
105. Shen, P., Li, W., Ding, Z., Deng, Y., Liu, Y., Zhu, X., … Zheng, T. (2018). A competitive aptamer chemiluminescence assay for ochratoxin A using a single silica photonic crystal microsphere. *Analytical Biochemistry*, *554*(May), 28–33. https://doi.org/10.1016/j.ab.2018.05.025
106. Shen, P., Li, W., Liu, Y., Ding, Z., Deng, Y., Zhu, X., … Zheng, T. (2017). High-Throughput Low-Background G-Quadruplex Aptamer Chemiluminescence Assay for Ochratoxin A Using a Single Photonic Crystal Microsphere. *Analytical Chemistry*, *89*(21), 11862–11868. https://doi.org/10.1021/acs.analchem.7b03592
107. Sheng, L., Ren, J., Miao, Y., Wang, J., & Wang, E. (2011). PVP-coated graphene oxide for selective determination of ochratoxin A via quenching fluorescence of free aptamer. *Biosensors and Bioelectronics*, *26*(8), 3494–3499. https://doi.org/10.1016/j.bios.2011.01.032
108. Simão, E. P., Cao-Milán, R., Costa-Pedro, G., De Melo, C. P., Cao, R., Oliveira, M. D. L., & Andrade, C. A. S. (2017). Simple and Fast Picomolar Detection of Ochratoxin A Using a Reusable Label Free Aptasensor Built with a Layer-by-layer Procedure. *Electroanalysis*, *29*(10), 2268–2275. https://doi.org/10.1002/elan.201700290
109. Somerson, J., & Plaxco, K. W. (2018). Electrochemical aptamer-based sensors for rapid point-of-use monitoring of the mycotoxin ochratoxin a directly in a food stream. *Molecules*, *23*(4). https://doi.org/10.3390/molecules23040912
110. Song, C., Hong, W., Zhang, X., & Lu, Y. (2018). Label-free and sensitive detection of Ochratoxin A based on dsDNA-templated copper nanoparticles and exonuclease-catalyzed target recycling amplification. *Analyst*, *143*(8), 1829–1834. https://doi.org/10.1039/c8an00158h
111. Song, D., Yang, R., Fang, S., Liu, Y., Long, F., & Zhu, A. (2018). SERS based aptasensor for ochratoxin A by combining Fe3O4@Au magnetic nanoparticles and Au-DTNB@Ag nanoprobes with multiple signal enhancement. *Microchimica Acta*, *185*(10). https://doi.org/10.1007/s00604-018-3020-2
112. Sui, Z., Wu, W., Komiyama, M., & Liang, X. (2018). Highly sensitive and selective detection of food toxin using three functional DNA hairpins. *Chemistry Letters*, *47*(8), 1026–1028. https://doi.org/10.1246/cl.180357
113. Sun, A. L., Zhang, Y. F., Sun, G. P., Wang, X. N., & Tang, D. (2017). Homogeneous electrochemical detection of ochratoxin A in foodstuff using aptamer–graphene oxide nanosheets and DNase I-based target recycling reaction. *Biosensors and Bioelectronics*, *89*, 659–665. https://doi.org/10.1016/j.bios.2015.12.032
114. Tan, Y., Wei, X., Zhang, Y., Wang, P., Qiu, B., Guo, L., … Yang, H. H. (2015). Exonuclease-Catalyzed Target Recycling Amplification and Immobilization-free Electrochemical Aptasensor. *Analytical Chemistry*, *87*(23), 11826–11831. https://doi.org/10.1021/acs.analchem.5b03314
115. Tian, F., Zhou, J., Jiao, B., & He, Y. (2019). A nanozyme-based cascade colorimetric aptasensor for amplified detection of ochratoxin A. *Nanoscale*, *11*(19), 9547–9555. https://doi.org/10.1039/c9nr02872b
116. Tian, J., Wei, W., Wang, J., Ji, S., Chen, G., & Lu, J. (2018). Fluorescence resonance energy transfer aptasensor between nanoceria and graphene quantum dots for the determination of ochratoxin A. *Analytica Chimica Acta*, *1000*, 265–272. https://doi.org/10.1016/j.aca.2017.08.018
117. Tong, P., Zhang, L., Xu, J. J., & Chen, H. Y. (2011). Simply amplified electrochemical aptasensor of Ochratoxin A based on exonuclease-catalyzed target recycling. *Biosensors and Bioelectronics*, *29*(1), 97–101. https://doi.org/10.1016/j.bios.2011.07.075
118. Tong, P., Zhao, W. W., Zhang, L., Xu, J. J., & Chen, H. Y. (2012). Double-probe signal enhancing strategy for toxin aptasensing based on rolling circle amplification. *Biosensors and Bioelectronics*, *33*(1), 146–151. https://doi.org/10.1016/j.bios.2011.12.042
119. Wang, C., Dong, X., Liu, Q., & Wang, K. (2015). Label-free colorimetric aptasensor for sensitive detection of ochratoxin A

utilizing hybridization chain reaction. *Analytica Chimica Acta*, *860*, 83–88. https://doi.org/10.1016/j.aca.2014.12.031
120. Wang, C., Qian, J., An, K., Huang, X., Zhao, L., Liu, Q., … Wang, K. (2017). Magneto-controlled aptasensor for simultaneous electrochemical detection of dual mycotoxins in maize using metal sulfide quantum dots coated silica as labels. *Biosensors and Bioelectronics*, *89*(September 2016), 802–809. https://doi.org/10.1016/j.bios.2016.10.010
121. Wang, C., Qian, J., Wang, K., Hua, M., Liu, Q., Hao, N., … Huang, X. (2015). Nitrogen-Doped Graphene Quantum Dots@SiO2 Nanoparticles as Electrochemiluminescence and Fluorescence Signal Indicators for Magnetically Controlled Aptasensor with Dual Detection Channels. *ACS Applied Materials and Interfaces*, *7*(48), 26865–26873. https://doi.org/10.1021/acsami.5b09300
122. Wang, C., Qian, J., Wang, K., Wang, K., Liu, Q., Dong, X., … Huang, X. (2015). Magnetic-fluorescent-targeting multifunctional aptasensorfor highly sensitive and one-step rapid detection of ochratoxin A. *Biosensors and Bioelectronics*, *68*, 783–790. https://doi.org/10.1016/j.bios.2015.02.008
123. Wang, C., Qian, J., Wang, K., Yang, X., Liu, Q., Hao, N., … Huang, X. (2016). Colorimetric aptasensing of ochratoxin A using Au@Fe3O4 nanoparticles as signal indicator and magnetic separator. *Biosensors and Bioelectronics*, *77*, 1183–1191. https://doi.org/10.1016/j.bios.2015.11.004
124. Wang, C., Tan, R., & Chen, D. (2018). Fluorescence method for quickly detecting ochratoxin A in flour and beer using nitrogen doped carbon dots and silver nanoparticles. *Talanta*, *182*(November 2017), 363–370. https://doi.org/10.1016/j.talanta.2018.02.007
125. Wang, J., Wang, Y., Liu, S., Wang, H., Zhang, X., Song, X., … Huang, J. (2019). Primer remodeling amplification-activated multisite-catalytic hairpin assembly enabling the concurrent formation of Y-shaped DNA nanotorches for the fluorescence assay of ochratoxin A. *Analyst*, *144*(10), 3389–3397. https://doi.org/10.1039/c9an00316a
126. Wang, L., Chen, W., Ma, W., Liu, L., Ma, W., Zhao, Y., … Xu, C. (2011). Fluorescent strip sensor for rapid determination of toxins. *Chemical Communications*, *47*(5), 1574–1576. https://doi.org/10.1039/c0cc04032k
127. Wang, L., Ma, W., Chen, W., Liu, L., Ma, W., Zhu, Y., … Xu, C. (2011). An aptamer-based chromatographic strip assay for sensitive toxin semi-quantitative detection. *Biosensors and Bioelectronics*, *26*(6), 3059–3062. https://doi.org/10.1016/j.bios.2010.11.040
128. Wang, R., Xiang, Y., Zhou, X., Liu, L. hua, & Shi, H. (2015). A reusable aptamer-based evanescent wave all-fiber biosensor for highly sensitive detection of Ochratoxin A. *Biosensors and Bioelectronics*, *66*, 11–18. https://doi.org/10.1016/j.bios.2014.10.079
129. Wang, X., Shan, Y., Gong, M., Jin, X., Lv, liangrui, Jiang, M., & Xu, J. (2019). A novel electrochemical sensor for ochratoxin A based on the hairpin aptamer and double report DNA via multiple signal amplification strategy. *Sensors and Actuators, B: Chemical*, *281*(October 2018), 595–601. https://doi.org/10.1016/j.snb.2018.10.148
130. Wang, Z., Duan, N., Hun, X., & Wu, S. (2010). Electrochemiluminescent aptamer biosensor for the determination of ochratoxin A at a gold-nanoparticles-modified gold electrode using N-(aminobutyl)-N-ethylisoluminol as a luminescent label. *Analytical and Bioanalytical Chemistry*, *398*(5), 2125–2132. https://doi.org/10.1007/s00216-010-4146-1
131. Wei, M., Wang, C., Xu, E., Chen, J., Xu, X., Wei, W., & Liu, S. (2019). A simple and sensitive electrochemiluminescence aptasensor for determination of ochratoxin A based on a nicking endonuclease-powered DNA walking machine. *Food Chemistry*, *282*(September 2018), 141–146. https://doi.org/10.1016/j.foodchem.2019.01.011
132. Wei, Y., Zhang, J., Wang, X., & Duan, Y. (2015). Amplified fluorescent aptasensor through catalytic recycling for highly sensitive detection of ochratoxin A. *Biosensors and Bioelectronics*, *65*, 16–22. https://doi.org/10.1016/j.bios.2014.09.100
133. Wu, J., Chu, H., Mei, Z., Deng, Y., Xue, F., Zheng, L., & Chen, W. (2012). Ultrasensitive one-step rapid detection of ochratoxin A by the folding-based electrochemical aptasensor. *Analytica Chimica Acta*, *753*, 27–31. https://doi.org/10.1016/j.aca.2012.09.036
134. Wu, K., Ma, C., Zhao, H., Chen, M., & Deng, Z. (2019). Sensitive aptamer-based fluorescene assay for ochratoxin A based on RNase H signal amplification. *Food Chemistry*, *277*, 273–278. https://doi.org/10.1016/j.foodchem.2018.10.130
135. Wu, K., Ma, C., Zhao, H., He, H., & Chen, H. (2018). Label-free G-quadruplex aptamer fluorescence assay for Ochratoxin A using a thioflavin T probe. *Toxins*, *10*(5). https://doi.org/10.3390/toxins10050198
136. Wu, S., Duan, N., Ma, X., Xia, Y., Wang, H., Wang, Z., & Zhang, Q. (2012). Multiplexed fluorescence resonance energy transfer aptasensor between upconversion nanoparticles and graphene oxide for the simultaneous determination of mycotoxins. *Analytical Chemistry*, *84*(14), 6263–6270. https://doi.org/10.1021/ac301534w
137. Wu, S., Duan, N., Wang, Z., & Wang, H. (2011). Aptamer-functionalized magnetic nanoparticle-based bioassay for the detection of ochratoxin A using upconversion nanoparticles as labels. *Analyst*, *136*(11), 2306–2314. https://doi.org/10.1039/c0an00735h
138. Wu, S., Liu, L., Duan, N., Wang, W., Yu, Q., & Wang, Z. (2018). A test strip for ochratoxin A based on the use of aptamer-modified fluorescence upconversion nanoparticles. *Microchimica Acta*, *185*(11). https://doi.org/10.1007/s00604-018-3022-0
139. Wu, X., Hu, J., Zhu, B., Lu, L., Huang, X., & Pang, D. (2011). Aptamer-targeted magnetic nanospheres as a solid-phase extraction sorbent for determination of ochratoxin A in food samples. *Journal of Chromatography A*, *1218*(41), 7341–7346. https://doi.org/10.1016/j.chroma.2011.08.045
140. Xiao, M. W., Bai, X. L., Liu, Y. M., Yang, L., & Liao, X. (2018). Simultaneous determination of trace Aflatoxin B1 and Ochratoxin A by aptamer-based microchip capillary electrophoresis in food samples. *Journal of Chromatography A*, *1569*, 222–228. https://doi.org/10.1016/j.chroma.2018.07.051
141. Xu, G., Zhao, J., Liu, N., Yang, M., Zhao, Q., Li, C., & Liu, M. (2019). Structure-guided post-SELEX optimization of an ochratoxin A aptamer. *Nucleic Acids Research*, *47*(11), 5963–5972. https://doi.org/10.1093/nar/gkz336
142. Xu, J., Li, W., Shen, P., Li, Y., Li, Y., Deng, Y., … Zheng, T. (2017). Microfluidic fabrication of photonic encoding magnetized silica microspheres for aptamer-based enrichment of Ochratoxin A. *Microchimica Acta*, *184*(10), 3755–3763. https://doi.org/10.1007/s00604-017-2400-3
143. Xu, L., Zhang, Z., Zhang, Q., & Li, P. (2016). Mycotoxin determination in foods using advanced sensors based on antibodies or aptamers. *Toxins*, *8*(8), 1–16. https://doi.org/10.3390/toxins8080239
144. Xu, X., Xu, C., & Ying, Y. (2016). Aptasensor for the simple detection of ochratoxin A based on side-by-side assembly of gold nanorods. *RSC Advances*, *6*(56), 50437–50443. https://doi.org/10.1039/c6ra04439e
145. Yang, C., Lates, V., Prieto-Simón, B., Marty, J. L., & Yang, X. (2013). Rapid high-throughput analysis of ochratoxin A by the self-assembly of DNAzyme-aptamer conjugates in wine. *Talanta*, *116*, 520–526. https://doi.org/10.1016/j.talanta.2013.07.011
146. Yang, C., Lates, V., Prieto-Simón, B., Marty, J. L., & Yang, X. (2012). Aptamer-DNAzyme hairpins for biosensing of Ochratoxin A. *Biosensors and Bioelectronics*, *32*(1), 208–212. https://doi.org/10.1016/j.bios.2011.12.011
147. Yang, C., Wang, Y., Marty, J. L., & Yang, X. (2011). Aptamer-based colorimetric biosensing of Ochratoxin A using

unmodified gold nanoparticles indicator. *Biosensors and Bioelectronics*, *26*(5), 2724–2727. https://doi.org/10.1016/j.bios.2010.09.032

148. Yang, L., Zhang, Y., Li, R., Lin, C., Guo, L., Qiu, B., … Chen, G. (2015). Electrochemiluminescence biosensor for ultrasensitive determination of ochratoxin A in corn samples based on aptamer and hyperbranched rolling circle amplification. *Biosensors and Bioelectronics*, *70*, 268–274. https://doi.org/10.1016/j.bios.2015.03.067
149. Yang, M., Jiang, B., Xie, J., Xiang, Y., Yuan, R., & Chai, Y. (2014). Electrochemiluminescence recovery-based aptasensor for sensitive Ochratoxin A detection via exonuclease-catalyzed target recycling amplification. *Talanta*, *125*, 45–50. https://doi.org/10.1016/j.talanta.2014.02.061
150. Yang, X., Hu, Y., Kong, W., Chu, X., Yang, M., Zhao, M., & Ouyang, Z. (2014). Ultra-fast liquid chromatography with tandem mass spectrometry determination of ochratoxin A in traditional Chinese medicines based on vortex-assisted solid-liquid microextraction and aptamer-affinity column clean-up. *Journal of Separation Science*, *37*(21), 3052–3059. https://doi.org/10.1002/jssc.201400635
151. Yang, X., Kong, W., Hu, Y., Yang, M., Huang, L., Zhao, M., & Ouyang, Z. (2014). Aptamer-affinity column clean-up coupled with ultra high performance liquid chromatography and fluorescence detection for the rapid determination of ochratoxin A in ginger powder. *Journal of Separation Science*, *37*(7), 853–860. https://doi.org/10.1002/jssc.201301136
152. Yang, X., Qian, J., Jiang, L., Yan, Y., Wang, K., Liu, Q., & Wang, K. (2014). Ultrasensitive electrochemical aptasensor for ochratoxin A based on two-level cascaded signal amplification strategy. *Bioelectrochemistry*, *96*, 7–13. https://doi.org/10.1016/j.bioelechem.2013.11.006
153. Yang, Y., Li, W., Shen, P., Liu, R., Li, Y., Xu, J., … Zheng, T. (2017). Aptamer fluorescence signal recovery screening for multiplex mycotoxins in cereal samples based on photonic crystal microsphere suspension array. *Sensors and Actuators, B: Chemical*, *248*, 351–358. https://doi.org/10.1016/j.snb.2017.04.004
154. Yao, L., Chen, Y., Teng, J., Zheng, W., Wu, J., Adeloju, S. B., … Chen, W. (2015). Integrated platform with magnetic purification and rolling circular amplification for sensitive fluorescent detection of ochratoxin A. *Biosensors and Bioelectronics*, *74*, 534–538. https://doi.org/10.1016/j.bios.2015.06.056
155. Yin, J., Liu, Y., Wang, S., Deng, J., Lin, X., & Gao, J. (2018). Engineering a universal and label-free evaluation method for mycotoxins detection based on strand displacement amplification and G-quadruplex signal amplification. *Sensors and Actuators, B: Chemical*, *256*, 573–579. https://doi.org/10.1016/j.snb.2017.10.083
156. Yin, X., Wang, S., Liu, X., He, C., Tang, Y., Li, Q., … Dong, Y. (2017). Aptamer-based colorimetric biosensing of ochratoxin A in fortified white grape wine sample using unmodified gold nanoparticles. *Analytical Sciences*, *33*(6), 659–664. https://doi.org/10.2116/analsci.33.659
157. Yu, X., Lin, Y., Wang, X., Xu, L., Wang, Z., & Fu, F. F. (2018). Exonuclease-assisted multicolor aptasensor for visual detection of ochratoxin A based on G-quadruplex-hemin DNAzyme-mediated etching of gold nanorod. *Microchimica Acta*, *185*(5). https://doi.org/10.1007/s00604-018-2811-9
158. Yuan, Y., Wei, S., Liu, G., Xie, S., Chai, Y., & Yuan, R. (2014). Ultrasensitive electrochemiluminescent aptasensor for ochratoxin A detection with the loop-mediated isothermal amplification. *Analytica Chimica Acta*, *811*, 70–75. https://doi.org/10.1016/j.aca.2013.11.022
159. Yue, S., Jie, X., Wei, L., Bin, C., Dou Dou, W., Yi, Y., … Tiesong, Z. (2014). Simultaneous detection of ochratoxin a and fumonisin b1 in cereal samples using an aptamer-photonic crystal encoded suspension array. *Analytical Chemistry*, *86*(23), 11797–11802. https://doi.org/10.1021/ac503355n
160. Zhang, G., Zhu, C., Huang, Y., Yan, J., & Chen, A. (2018). A lateral flow strip based aptasensor for detection of Ochratoxin a in corn samples. *Molecules*, *23*(2). https://doi.org/10.3390/molecules23020291
161. Zhang, J., Chen, J., Zhang, X., Zeng, Z., Chen, M., & Wang, S. (2012). An electrochemical biosensor based on hairpin-DNA aptamer probe and restriction endonuclease for ochratoxin A detection. *Electrochemistry Communications*, *25*(1), 5–7. https://doi.org/10.1016/j.elecom.2012.09.006
162. Zhang, J., Zhang, X., Yang, G., Chen, J., & Wang, S. (2013). A signal-on fluorescent aptasensor based on Tb3+ and structure-switching aptamer for label-free detection of Ochratoxin A in wheat. *Biosensors and Bioelectronics*, *41*(1), 704–709. https://doi.org/10.1016/j.bios.2012.09.053
163. Zhang, Y. Y., Wang, S., Zhang, Y. Y., Pang, G., & Guo, S. (2017). Tuning the Aggregation/Disaggregation Behavior of Graphene Quantum Dots by Structure-Switching Aptamer for High-Sensitivity Fluorescent Ochratoxin A Sensor. *Analytical Chemistry*, *89*(3), 1704–1709. https://doi.org/10.1021/acs.analchem.6b03913
164. Zhang, Y., Yang, L., Lin, C., Guo, L., Qiu, B., Lin, Z., & Chen, G. (2015). Fluorescence aptasensor for Ochratoxin A in the food samples based on hyperbranched rolling circle amplification. *Analytical Methods*, *3*, 10715–10722. https://doi.org/10.1039/b000000x
165. Zhao, H., Xiang, X., Chen, M., & Ma, C. (2019). Aptamer-based fluorometric ochratoxin a assay based on photoinduced electron transfer. *Toxins*, *11*(2). https://doi.org/10.3390/toxins11020065
166. Zhao, Q., Geng, X., & Wang, H. (2013). Fluorescent sensing ochratoxin A with single fluorophore-labeled aptamer. *Analytical and Bioanalytical Chemistry*, *405*(19), 6281–6286. https://doi.org/10.1007/s00216-013-7047-2
167. Zhao, Q., Lv, Q., & Wang, H. (2014). Identification of allosteric nucleotide sites of tetramethylrhodamine- labeled aptamer for noncompetitive aptamer-based fluorescence anisotropy detection of a small molecule, ochratoxin A. *Analytical Chemistry*, *86*(2), 1238–1245. https://doi.org/10.1021/ac4035532
168. Zhao, X., Wu, X., Xu, L., Ma, W., Kuang, H., Wang, L., & Xu, C. (2015). Building heterogeneous core-satellite chiral assemblies for ultrasensitive toxin detection. *Biosensors and Bioelectronics*, *66*, 554–558. https://doi.org/10.1016/j.bios.2014.12.021
169. Zhu, Y., Xia, X., Deng, S., Yan, B., Dong, Y., Zhang, K., … He, Q. (2019). Label-free fluorescent aptasensing of mycotoxins via aggregation-induced emission dye. *Dyes and Pigments*, *170*(May), 107572. https://doi.org/10.1016/j.dyepig.2019.107572
170. Zhu, Z. Z., Feng, M., Zuo, L., Zhu, Z. Z., Wang, F., Chen, L., … Luo, S. Z. (2015). An aptamer based surface plasmon resonance biosensor for the detection of ochratoxin A in wine and peanut oil. *Biosensors and Bioelectronics*, *65*, 320–326. https://doi.org/10.1016/j.bios.2014.10.059
